# Genotyping-by-sequencing and SNP-arrays are complementary for detecting quantitative trait loci by tagging different haplotypes in association studies

**DOI:** 10.1101/476598

**Authors:** Sandra Silvia Negro, Emilie Millet, Delphine Madur, Cyril Bauland, Valérie Combes, Claude Welcker, François Tardieu, Alain Charcosset, Stéphane Dimitri Nicolas

**Author notes:** E-mail addresses.

## Abstract

**Background:** Single Nucleotide Polymorphism (SNP) array and re-sequencing technologies have different properties (*e.g.* calling rate, minor allele frequency profile) and drawbacks (*e.g.* ascertainment bias). This lead us to study their complementarity and the consequences of using them separately or combined in diversity analyses and Genome-Wide Association Studies (GWAS). We performed GWAS on three traits (grain yield, plant height and male flowering time) measured in 22 environments on a panel of 247 F1 hybrids obtained by crossing 247 diverse dent maize inbred lines with a same flint line. The 247 lines were genotyped using three genotyping technologies (Genotyping-By-Sequencing, Illumina Infinium 50K and Affymetrix Axiom 600K arrays).

**Results:** The effects of ascertainment bias of the 50K and 600K arrays were negligible for deciphering global genetic trends of diversity and for estimating relatedness in this panel. We developed an original approach based on linkage disequilibrium (LD) extent in order to determine whether SNPs significantly associated with a trait and that are physically linked should be considered as a single Quantitative Trait Locus (QTL) or several independent QTLs. Using this approach, we showed that the combination of the three technologies, which have different SNP distributions and densities, allowed us to detect more QTLs (gain in power) and potentially refine the localization of the causal polymorphisms (gain in resolution).

**Conclusions:** Conceptually different technologies are complementary for detecting QTLs by tagging different haplotypes in association studies. Considering LD, marker density and the combination of different technologies (SNP-arrays and re-sequencing), the genotypic data available were most likely enough to well represent polymorphisms in the centromeric regions, whereas using more markers would be beneficial for telomeric regions.

## Background

Understanding the genetic bases of complex traits involved in the adaptation to biotic and abiotic stress in plants is a pressing concern, with world-wide drought due to climate change as a major source of human food and agriculture threats. Recent progress in next generation sequencing and genotyping array technologies contribute to a better understanding of the genetic basis of quantitative trait variation by allowing Genome-Wide Association Studies (GWAS) on large diversity panels [1]. Single Nucleotide Polymorphism (SNP)-based techniques became the most commonly used genotyping methods for GWAS because SNPs are cheap, numerous, codominant and can be automatically analysed with SNP-arrays or produced by genotyping-by-sequencing (GBS), or sequencing [2–4]. The decreasing cost of genotyping technologies has led to an exponential increase in the number of markers used for the GWAS in association panels, thereby raising the question of computation time to perform the association tests. Computational issues were addressed by using either approximate methods by avoiding re-estimating variance components for each SNP [5] or exact methods using mathematical tools for sparing time in matrix inversion [6, 7]. It is noteworthy that using approximate computation in GWAS can produce inaccurate p-values when the SNP effect size is large or/and when the sample structure is strong [8].

Several causes may impact the power of Quantitative Trait Locus (QTL, locus involved in quantitative trait variation) detection in GWAS. Highly diverse panels have in general undergone multiple historical recombinations, leading to a low extent of linkage disequilibrium (LD). However, these panels can present different average and local patterns of LD [9–11]. A high marker density and a proper distribution of SNPs are therefore essential to capture causal polymorphisms. Furthermore, minor allele frequencies (MAF), population stratification and cryptic relatedness are three other important parameters affecting power and false positive detection [12, 14]. These last two factors are substantial in several cultivated species such as maize [15] and grapevine [16], and their impact on LD can be statistically evaluated [17]. Population structure and kinship can be estimated using molecular markers [18–21] and can be modelled to efficiently detect marker-trait associations due to linkage only [12, 22, 23]. These advances have largely increased the power and effectiveness of linear mixed models that can now efficiently account for population structure and relatedness in GWAS [8, 12].

In maize, an Illumina Infinium HD 50,000 SNP-array, named MaizeSNP50 (hereafter 50 K) was developed by Ganal *et al.* [3] and has been used extensively for diversity and association studies [24, 25]. For example, GWAS were conducted to unravel the genetic architecture of phenology, yield component traits and to identify several flowering time QTLs linked to adaptation of tropical maize to temperate climate [26, 27]. In the same way, Rincent *et al.* [11] showed that LD occurs over a longer distance in a dent than in a flint panel, with appreciable effects on the power of QTL detection. Low LD extent and relationship between allelic frequencies with population and pedigree structure at some SNPs reduce the power of GWAS [14, 27]. Therefore, higher marker densities are desirable because the maize genome size is large (2.4 Gb), the level of diversity is high (more than one substitution per hundred nucleotides), and LD extent is low [28]. As a consequence, an Affymetrix Axiom 600,000 SNP-array (hereafter 600 K) was developed and used in association genetics [29, 30] and detection of selective sweeps [4]. Another possibility is whole genome sequencing, but this is currently impractical for large genomes such as maize because of the associated cost. Hence, a Genotyping-By-Sequencing (GBS) procedure has been developed [2] that targets low-copy genomic regions by using restriction enzymes. Genotyping-by-sequencing technology is cost-effective and has been successfully used in maize for genomic prediction [31]. Romay *et al.* [32] and Gouesnard *et al.* [33] highlighted the interest of the GBS for (i) deciphering and comparing the genetic diversity of the inbred lines in seedbanks and (ii) identifying QTLs by GWAS for kernel colour, sweet corn and flowering time.

Few studies in plants have compared datasets from different high-throughput genotyping technologies [34–36]. Elbasyoni et al. [34] used GBS and a 90 K SNP-array in winter wheat. They highlighted strong positive correlations between the population structure matrices and kinships identified by both technologies. They showed that GBS-SNPs led to higher genomic prediction accuracy compared to Array-SNPs. Torkamaneh and Belzile [36] used GBS and a 50 K SNP-array in soybean. They estimated *ca.* 98% accuracy of genotype called by their GBS pipeline and showed that the accuracy of imputation for missing genotypes was hardly affected by the chosen the MAF and only moderately affected by the rate of missing values. Li et al. [35] created a reliable integrated variation map using a 600 K and 50 K SNP-array, GBS and RNA sequencing to dissect regulatory causality and its link to maize kernel variation. These authors used a fixed physical distance (<10 kb) for grouping associated SNPs into QTLs despite the variable LD pattern along the genome. None of these studies compared QTL detection between the different technologies.

The main drawback of the DNA arrays is that they do not allow the discovery of new SNPs. This possibly leads to some ascertainment bias in diversity analysis when the SNPs selected for building arrays come from (i) the sequencing of a set of individuals that did not represent well the diversity explored in the studied panel, (ii) a subset of SNPs that skews the allelic frequency profile towards the intermediate frequencies [27, 37]. Ascertainment bias can compromise the ability of the SNP-arrays to reveal an exact view of the genetic diversity [37]. Genotyping-by-sequencing can overcome ascertainment bias since it is based on sequencing and therefore allows the discovery of alleles in the diversity panel analysed. It can be generalized to any species at a low cost providing that numerous individuals have been sequenced in order to build a representative library of short haplotypes to call SNPs [38]. Non-repetitive regions of genomes can be targeted with two- to three-fold higher efficiency, thereby considerably reducing the computationally challenging problems associated with alignment in species with high repeat content. However, GBS may have low coverage leading to a high missing data rate (65% in both studies; [32, 33]) and heterozygote under-calling, depending on genome size and structure, and on the multiplexing level per sequencing flow-cell. Furthermore, GBS requires the establishment of demanding bioinformatic pipelines and imputation algorithms [39]. Pipelines have been developed to call SNP genotypes from raw GBS sequence data and to impute the missing data from a haplotype library [38, 39].

Here, we investigated the impact of using GBS and SNP-arrays on the quality of the genotyping data, together with the biological properties of data generated by each technology, and the potential complementarity of these approaches. In particular, we analyzed the impact of marker density and genotyping technologies (sequencing *vs* array) on (i) the estimates of relatedness and population structure, and (ii) the detection of QTLs (power). To address these issues, we performed a GWAS based on genotypic datasets obtained using either GBS or SNP-arrays with low (50k) or high (600k) densities on a diversity panel of maize hybrids obtained by crossing a panel of dent lines with a common flint tester lines. Three traits were considered, namely grain yield, plant height and male flowering time (day to anthesis), measured in 22 different environments (sites × years × treatments) over Europe. We developed an original approach based on LD extent in order to determine whether SNPs significantly associated with a trait should be considered as a single QTL or several independent QTLs.

## Results

### Combining Tassel and Beagle imputations improved the genotyping quality for GBS

We estimated the genotyping and imputation concordance of the GBS based on common markers with the 50 K or 600 K arrays (Additional file 1: Figure S1 and Table 1). The genotyping concordance of the 600 K with the 50 K was extremely high (99.50%), although slightly lower for residual heterozygotes (92.88%). After SNP calling from sequencing reads using AllZeaGBSv2.7 database (direct reads, GBS_1_, Additional file 1: Figure S1), the call rate was 33.81% for the common SNPs with the 50 K, *vs* 37% for the whole GBS dataset. The genotyping concordance rate between the direct reads of GBS and the 50 K was 98.88% (Table 1). After imputation using *TASSEL* by Cornell Institute (GBS_2_), the concordance rate was 96.04% on the common markers with the 50 K and 11.91% of missing data remained for the whole GBS dataset. In GBS_3_, all missing data were imputed by *Beagle* and the remaining missing genotypes in GBS_2_ were excluded here to be comparable with *TASSEL*. This method yielded a lower concordance rate (93.04% and 92.84% with 50 K and 600 K, respectively). In an attempt to increase the concordance rate of the genotyping while removing missing data, we tested two additional methods, namely GBS_4_ where the missing data and heterozygotes of Cornell imputed data (GBS_2_) were replaced by Beagle imputation, and GBS_5_ where Cornell homozygous genotypes (GBS_2_) were completed by imputations from GBS_3_ (Additional file 1: Figure S1 and Table 1). GBS_5_ displayed a slightly better concordance rate than GBS_2_ (96.25% vs 96.04%) and predicted heterozygotes with a higher quality than GBS_4_. GBS_5_ was therefore used for all genetic analyses and named GBS hereafter.

**Table 1:** Percentage of GBS concordance and call rates (in parentheses).

|  | Reference | Total | Homozygotes | Heterozygotes |
| --- | --- | --- | --- | --- |
| GBS<sub>1</sub> Direct Read | 50K | 98.88 (33.81) | 99.03 (33.72) | 45.09 (0.09) |
|  | 600K | 98.99 (35.58) | 99.21 (35.47) | 28.67 (0.11) |
| GBS<sub>2</sub> Cornell Imputation | 50K | 96.04 (91.56) | 98.66 (88.79) | 12.51 (2.78) |
|  | 600K | 95.50 (93.41) | 98.69 (90.14) | 7.75 (3.28) |
| GBS<sub>3</sub> Beagle Imputation | 50K | 93.04 (*91.56) | 93.23 (91.30) | 30.54 (0.26) |
|  | 600K | 92.84 (*93.41) | 93.07 (93.12) | 22.50 (0.29) |
| GBS<sub>4</sub> Beagle Imputation on the missing data and heterozygotes after Tassel Imputation (GBS<sub>2</sub>) | 50K | 96.46 (*97.64) | 96.46 (97.63) | <0.01 (<0.01) |
|  | 600K | 96.21 (*99.97) | 96.21 (>99.99) | <0.01 (<0.01) |
| GBS<sub>5</sub> Compilation of Homoz. genotypes from Tassel Imputation (GBS<sub>2</sub>) and Imputation by Beagle for Other Data (GBS<sub>3</sub>) | 50K | 96.25 (*97.65) | 96.36 (97.47) | 39.07 (0.18) |
|  | 600K | 95.98 (*99.97) | 96.11 (99.78) | 32.02 (0.22) |

The 50K and 600K SNP-arrays were considered as reference genotypes. \* After Beagle inference of missing data, the call rate was 100%. Here the call rate is <100% because the comparison was made against the 50K and the 600K that include few missing data. For GBS<sub>3</sub>, the remaining missing genotypes in GBS<sub>2</sub> were also excluded to obtain comparable results.

### GBS displayed more rare alleles and lower call rate than SNP-arrays

The SNP call rate was higher for the SNP-arrays (average values of 96% and >99% for the 50 K and 600 K, respectively), than for the GBS (37% for the direct reads). The MAF distribution differed between the technologies (Additional file 2: Figure S2): while the use of SNP-arrays resulted in a near-uniform distribution, GBS resulted in an excess of rare alleles with a L-shaped distribution (22% of SNPs with MAF < 0.05 for the GBS *versus* 6% and 9% for the 50 K and 600 K, respectively). This can be explained by the fact that the 50 K was based on 27 sequenced lines for SNPs discovery [3], the 600 K was based on 30 lines [4], whereas GBS was based on 31,978 lines, thereby leading to higher discovery of rare alleles. Consistent with MAF distribution, the average gene diversity (*He*) was lower for GBS (0.27) than for arrays (0.35 and 0.34 for the 50 K and 600 K arrays, respectively). The distribution of SNP residual heterozygosity of inbred lines was similar for the three technologies, with a mean of 0.80%, 0.89% and 0.22% for the 50 K, 600 K and GBS, respectively. The residual heterozygosity of inbred lines was highly correlated between technologies with large coefficients of Spearman correlation: r_50K-600K_ = 0.90, r_50K-GBS_ = 0.76, r_600K-GBS_ = 0.83. The distribution of the SNPs along the genome was denser in the telomeres for the GBS and in the peri-centromeric regions for the 600 K, whereas the 50 K exhibited a more uniform distribution (Figure 1 and top graph in Additional file 3: Figure S3).

**Figure 1:**
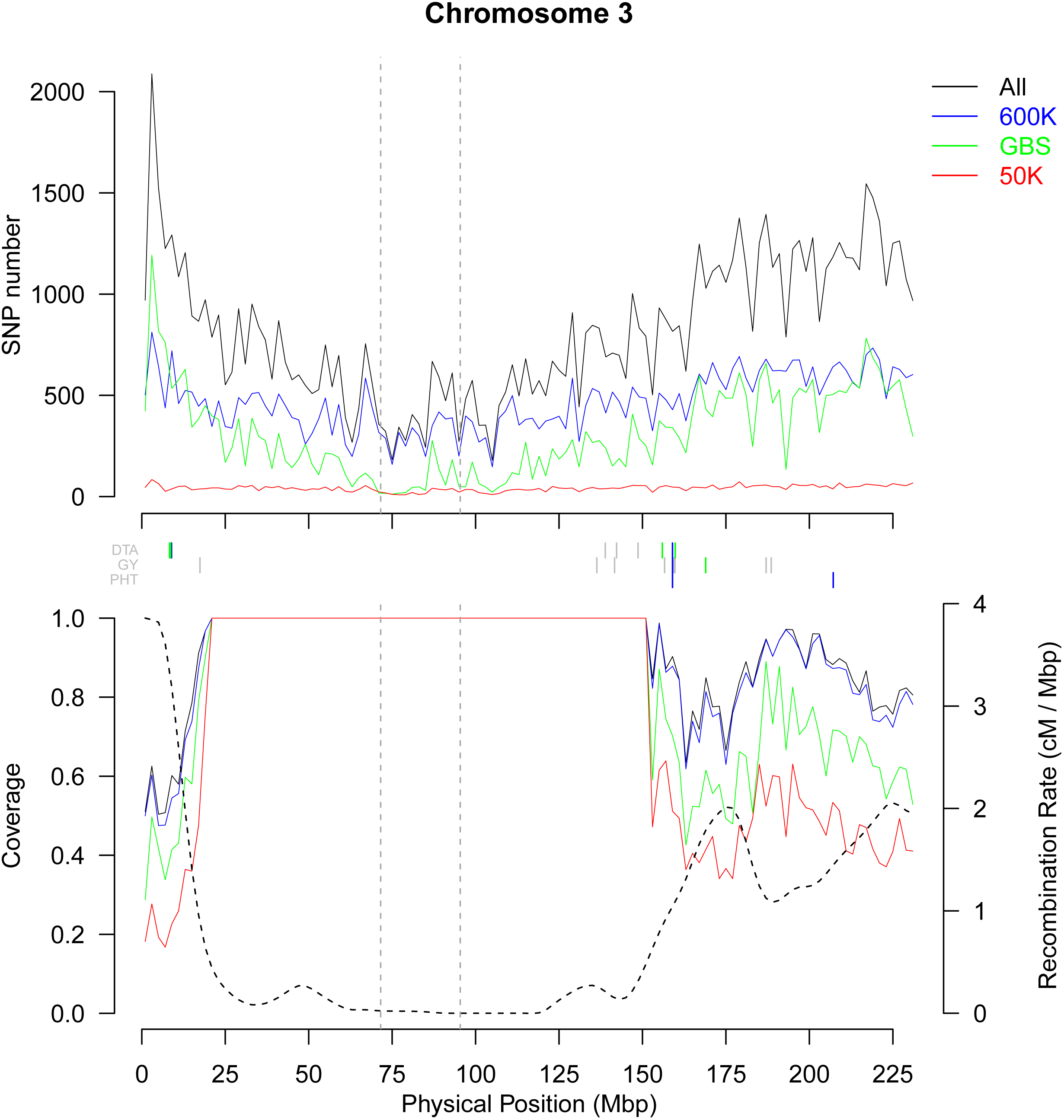
Variation of the markers density, the recombination rate and the genome coverage in non-overlapping 2 Mbp windows along chromosome 3. Markers have MAF above 5%. Top panel shows the variation of SNP number. In the bottom panel, dotted line represents the variation of recombination rate (cM / Mbp) and solid lines the proportion of genome covered by the SNPs using the cumulated length of physical LD windows around each SNP in each 2Mbp-windows. In these two panels, green, blue, red and black lines represent variation for GBS, 600 K, 50 K and combined technologies, respectively. Vertical dotted gray lines indicate limits of centromeric regions. Vertical lines between the two panels indicate the position of QTLs for flowering time (DTA), grain yield (GY) and Plant Height (PHT). Green, blue, red vertical lines indicate QTLs detected only by GBS, 600 K and 50 K technologies, respectively. Grey lines indicated QTLs detected by at least two technologies. Only QTL including a marker associated with −log_10_(pval) above 6 were shown.

### Population structure and relatedness were consistent between the three technologies

We used the ADMIXTURE software to analyse the genetic structure within the studied panel based on SNPs from the three technologies, by considering two to ten groups. Based on a K- fold cross-validation, the clustering in four genetic groups (*N_Q_* = 4) was identified as the best for the three datasets. Considering a threshold of 0.5 for ancestral fraction, the assignation to the four groups was identical except for a few admixed inbred lines (Additional file 4: Figure S4). Based on the 50 K, the four groups were constituted by (i) 39 lines in the Non Stiff Stalk (Iodent) family traced by PH207, (ii) 46 lines in the Lancaster family traced by Mo17 and Oh43, and (iii) 55 lines in theStiff Stalk family traced by B73 and (iv) 107 lines that did not fit into these three primary heterotic groups, such as W117 and F7057. This organization appeared consistent with the organization of breeding programs into heterotic groups, generally related to few key founder lines.

We compared two estimators of relatedness between inbred lines, IBS (Identity-By-State) and *K_Freq* (Identity-By-Descent), calculated per technology. For IBS, pairs of individuals were on average more related using GBS than SNP-arrays (mean IBS: 0.66, 0.67 and 0.73 for 50 K, 600 K and GBS, respectively). As expected, mean IBD was close for the three technologies (*K_Freq*: −0.004). Relatedness estimates with the two SNP-arrays were highly correlated: r = 0.95 and 0.98 for IBS and *K_Freq*, respectively (Additional file 5: Figure S5b and d). Likewise, relatedness estimates between arrays and GBS were strongly correlated (between 0.94 and 0.98, Additional file 5: Figure S5b and d).

We further carried out diversity analyses by performing Principal Coordinate Analyses (PCoA) on IBD (*K_Freq*, weights by allelic frequency) estimated from the three technologies (Figure 2). The three first PCoA axes explained 12.9%, 15.6% and 16.3% of the variability for the GBS, 50 K and 600 K, respectively (Figure 2). The same pattern was observed regardless of the technology with the first axis separating the Stiff Stalk from all other groups (Iodent and Lancaster lines, see illustration with the 50 K kinship, Figure 2). Key founder lines of the three heterotic groups (Iodent: PH207, Stiff Stalk: B73, Lancaster: Mo17) were found at extreme positions along the axes, which was consistent with the admixture groups previously described.

**Figure 2:**
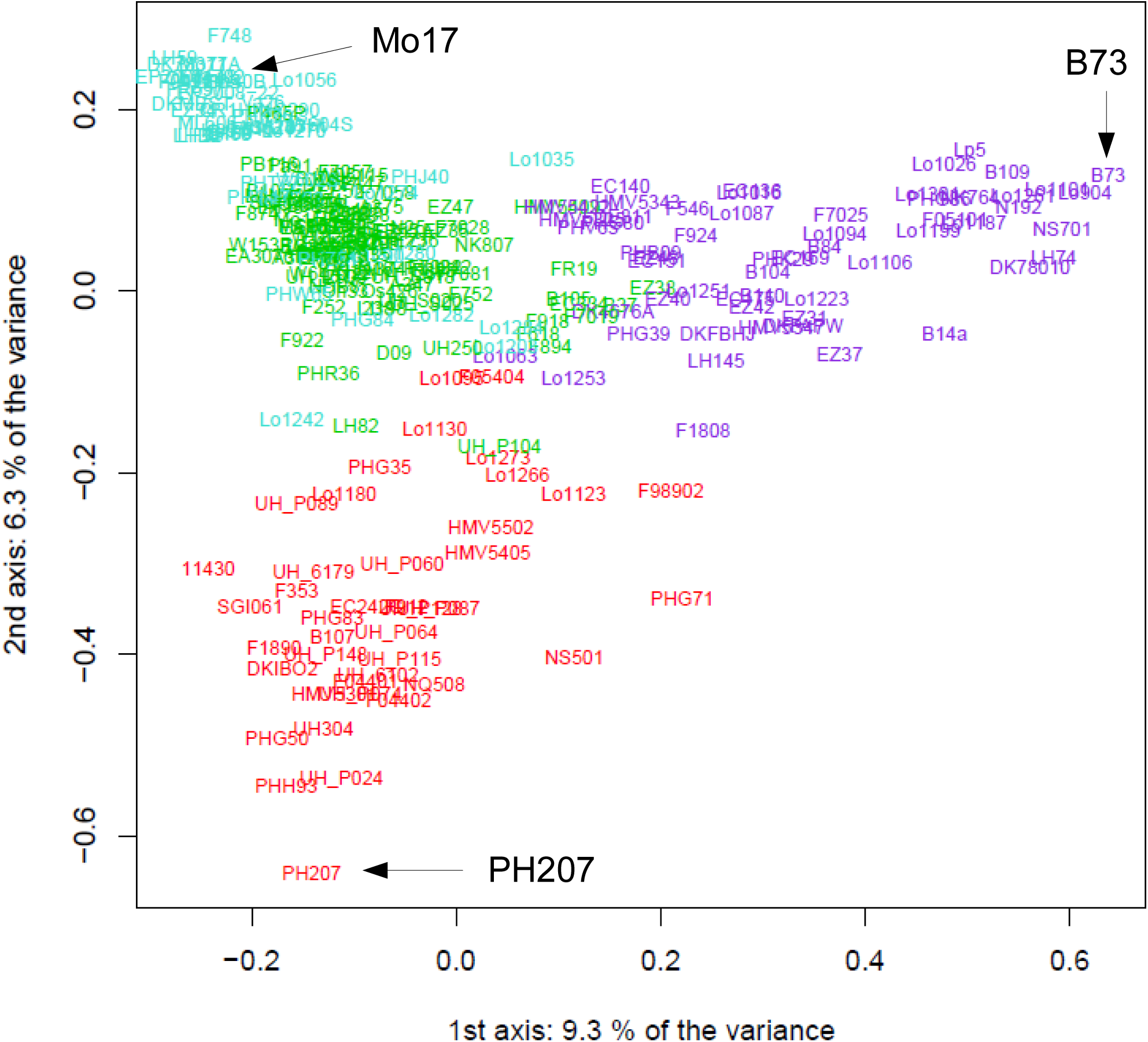
Principal coordinate analyses (PCoA) of the DROPS panel. The PCoA were based on the covariance matrix *K_Freq* estimated from the 50 K Illumina array. The genetic groups identified by ADMIXTURE (*N_Q_* = 4) are colored. Three key founders are indicated (Iodent: PH207 in red, Stiff Stalk: B73 in blueviolet, Lancaster: Mo17 in turquoise).

### Long distance linkage disequilibrium was removed by taking into account population structure or relatedness

In order to evaluate the effect of kinship and the genetic structure on linkage disequilibrium (LD), we studied genome-wide LD between 29,257 PANZEA markers from the 50 K within and between chromosomes before and after taking into account the kinship (*K_Freq* estimated from the 50 K), structure (Number of groups = 4) or both (Additional file 6: Figure S6). Whereas inter-chromosomal LD was only partially removed when the genetic structure was taken into account, it was mostly removed when either the kinship or both kinship and structure were considered (Additional file 6: Figure S6b and c). Accordingly, long distance intra-chromosomal LD was almost totally removed for all chromosomes by accounting for the kinship, structure or both. Interestingly, some pairs of loci located on different chromosomes or very distant on a same chromosome remained in high LD despite correction for genetic structure and kinship (Additional file 6: Figure S6). This can be explained either by genome assembly errors, by chromosomal rearrangements such as translocations or by strong epistatic interactions. Linkage disequilibrium decreased with genetic or physical distance Figure 3. The majority of pairs of loci with high LD (*r^2^K*>0.4) in spite of long physical distance (>30Mbp), were close genetically (<3cM), notably on chromosome 3, 5, 7 and to a lesser extent 9 and 10 (data not shown). These loci were located in centromeric and peri-centromeric regions that displayed low recombination rate, suggesting that this pattern was due to variation of recombination rate along the chromosome. Only very few pairs of loci in high LD were genetically distant (>5cM) but physically close (<2Mbp). Linkage disequilibrium (*r^2^K* and *r^2^KS*) was negligible beyond 1 cM since 99% of LD values were less than 0.12 in this case. Note that some unplaced SNPs remained in LD after taking into account the kinship and structure with some SNPs with known positions on chromosome 1, 3 and 4 (Additional File 6: Figure S6). Therefore, LD measurement corrected by the kinship can help to map unplaced SNPs.

**Figure 3:**
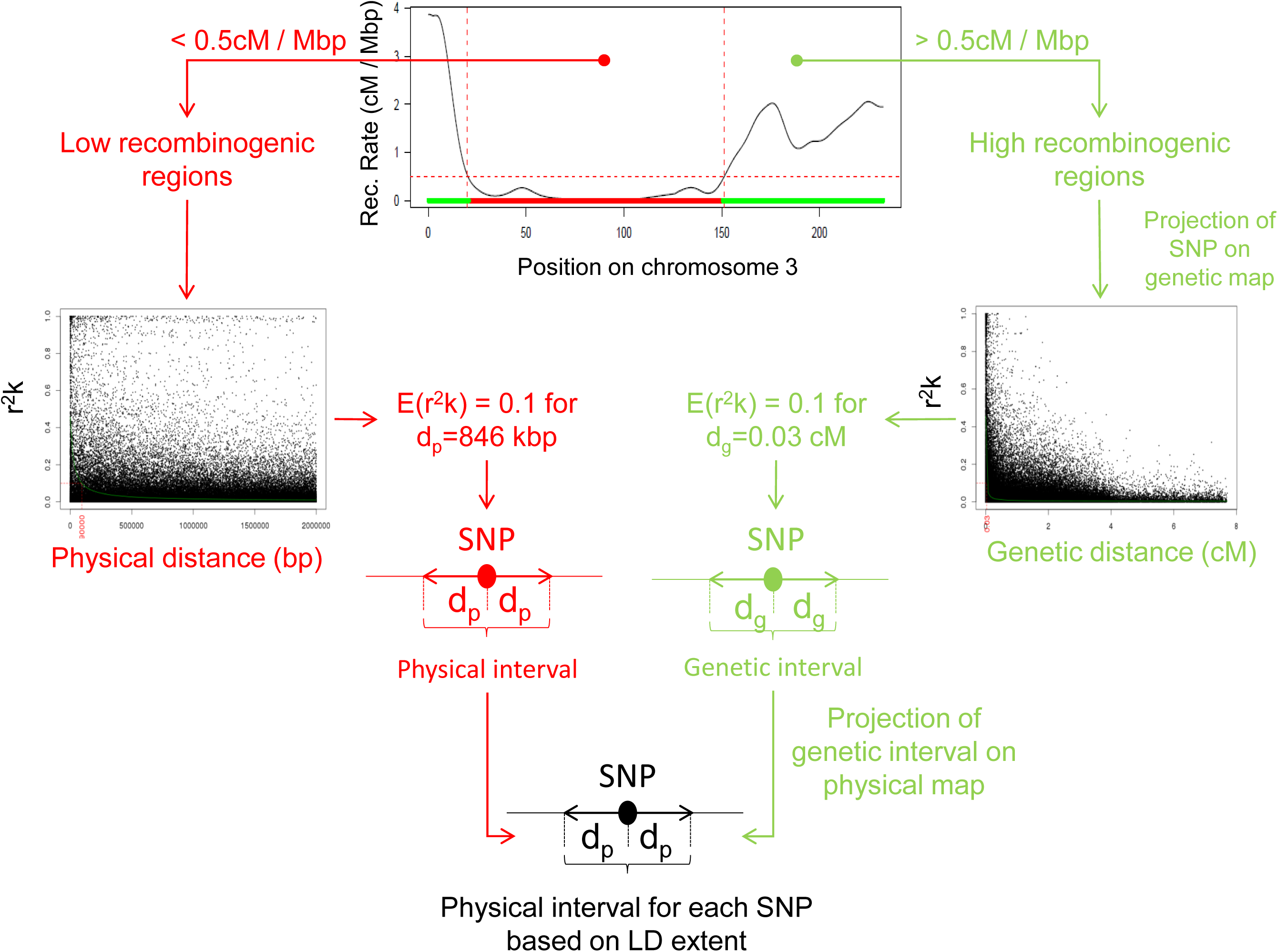
Linkage disequilibrium based approach to delineate a physical window around each SNP, exemplified with chromosome 3. Linkage disequilibrium (LD) windows were defined for each SNP based on physical LD extent in low recombinogenic regions (left part) and based on genetic LD extent in high recombinogenic regions (right part). These LD windows were used (i) to group significant SNPs into QTLs when they overlapped, (ii) to estimate genome coverage region covered by LD windows around SNPs, and (iii) identify putative genes underlying QTLs involved in trait variations.

### Linkage disequilibrium strongly differed between and within chromosomes

We combined the three technologies together to calculate the *r^2^K* for all pairs of SNPs which were genetically distant by less than 1 cM. For any chromosome region, LD extent in terms of genetic and physical distance showed a limited variation over the 100 sets of 500,000 loci pairs (cf. Material). This suggests that the estimation of LD extent did not strongly depend on our set of loci. LD extent varied significantly between chromosomes for both high recombinogenic (>0.5 cM/Mbp) and low recombinogenic regions (<0.5 cM/Mbp, Table 2). Chromosome 1 had the highest LD extent in high recombination regions (0.062 ±0.007 cM) and chromosome 9 the highest LD extent in low recombinogenic regions (898.6±21.7 kbp) (Table 2). Linkage disequilibrium extent relative to genetic and physical distances was highly and positively correlated in high recombinogenic regions (r = 0.86), whereas it was not in low recombinogenic regions (r = −0.64).

**Table 2:** Variation of LD extent, and percentage of genome covered.

|  |  | Chromosome |  |  |  |  |  |  |  |  |  | Whole Genome |
| --- | --- | --- | --- | --- | --- | --- | --- | --- | --- | --- | --- | --- |
|  |  | 1 | 2 | 3 | 4 | 5 | 6 | 7 | 8 | 9 | 10 |  |
| Physical Size (Mbp) |  | 301 | 237 | 232 | 241 | 217 | 169 | 176 | 175 | 156 | 150 | 2,058 |
| Genetic Size (cM) |  | 268 | 211 | 188 | 150 | 205 | 129 | 148 | 182 | 145 | 139 | 1766 |
| Physical LD extent (kbp) in low recombination regions |  | 306 | 491 | 846 | 808 | 658 | 418 | 547 | 497 | 899 | 815 | 629 |
| Genetic LD Extent (cM) in high recombination regions |  | 0.062 | 0.027 | 0.033 | 0.022 | 0.031 | 0.019 | 0.012 | 0.038 | 0.023 | 0.019 | 0.029 |
| Percent of physical genome covered | 50K | 81% | 72% | 76% | 77% | 74% | 67% | 71% | 73% | 71% | 71% | 74% |
|  | 600K | 98% | 88% | 91% | 89% | 90% | 84% | 81% | 90% | 87% | 84% | 89% |
|  | GBS | 92% | 81% | 84% | 83% | 83% | 77% | 77% | 81% | 79% | 76% | 82% |
|  | ALL | 98% | 90% | 92% | 90% | 91% | 87% | 83% | 92% | 88% | 85% | 90% |
| Percent of genetic map covered | 50K | 72% | 41% | 44% | 38% | 41% | 32% | 24% | 46% | 32% | 27% | 42% |
|  | 600K | 96% | 71% | 76% | 68% | 72% | 62% | 47% | 78% | 63% | 53% | 71% |
|  | GBS | 86% | 58% | 61% | 53% | 61% | 48% | 37% | 63% | 48% | 40% | 58% |
|  | ALL | 97% | 74% | 78% | 72% | 74% | 65% | 51% | 81% | 66% | 57% | 74% |

Genetic and Physical LD extent were obtained by adjusting Hill and Weir model's on 100 different sets of 500,000 loci randomly sampled in high ( $>0.5$ cM / Mbp) and low ( $<0.5$ cM / Mbp) recombination regions, respectively. The value represented the average across these 100 sets. The percentage of genome coverage was estimated using markers with $MAF > 5\%$ and $E(r^2k) = 0.1$ , for each technology and for the three technologies combined (ALL: GBS+600K+50K).

### Large differences in genome coverage between technologies

We estimated the percentage of the genome that was covered by LD windows around SNPs, calculated by using either physical or genetic distances (Figure 3, Table 2). We observed a strong difference in coverage between the three technologies at both genome-wide and chromosome scale, as illustrated in Figure 1 on chromosome 3 (Table 2, and Additional file 3: Figure S3). For a LD extent of *r^2^K* = 0.1, 74%, 82% and 89% of the physical map, and 42%, 58% and 71% of the genetic map were covered by the 50 K, the GBS and the 600 K, respectively (Table 2). For the combined data (ALL: 50 K + 600 K + GBS), the coverage strongly varied between chromosomes, ranging from 83% (chromosome 7) to 98% (chromosome 1) of the physical map, and from 51% (chromosome 7) to 97% (chromosome 1) of the genetic map (Table 2). For the physical map, increasing the LD extent threshold to *r^2^K*=0.4 reduced the genome coverage from 89% to 49% for 600 K, 82% to 28% for GBS, 74% to 20% for 50 K and 90% to 52% for the combined data. Increasing the MAF threshold reduced slightly the genome coverage, with smaller reduction for the physical map than genetic map. Surprisingly, increasing the SNP number by combining the markers from the arrays and GBS did not strongly increase the genome coverage as compared to the 600 K, regardless of the threshold for LD extent (Figure 1 and Additional file 3: Figure S3).

We observed a strong variation of genome coverage along each chromosome with contrasted patterns in low and high recombinogenic regions (Figure 1 Additional file 3: Figure S3). While low recombinogenic regions were totally covered with all the technologies (except for few intervals using the 50 K), the genome coverage in high recombinogenic regions varied depending on both technology and SNP distribution. 47% of the 2Mbp intervals in high recombination regions were better covered by the 600 K than the GBS against only 1%, which were better covered by GBS than 600 K. When exploring smaller window sizes (20, 100, 500 kb), the number of intervals better covered by 600 K than GBS decreased strongly when the intervals were shortened (17.1% of 20kbp-intervals *vs* 47.1% of 2Mbp-intervals). In the contrary, the intervals better covered by GBS than 600 K increased slightly (4.1% *vs* 1.1% of 2Mbp-intervals). The number of interval with no or weak coverage differences between GBS and 600 K increased strongly: 84.5% of 20kbp-intervals *vs* 68% of 2Mbp-intervals with coverage differences inferior to 10%. Interestingly, the proportion of interval with strong coverage differences (>50%) increased when the intervals were shortened (7.8% of 20kbp-intervals *vs* 0% of 2Mbp-intervals).

### Number of QTLs detected using genome-wide association studies increases with markers density

We observed a strong variation in the number of SNP significantly associated with the three traits across the 22 environments (Table 3). The mean number of significant SNPs per environment and trait was 3.7, 44.7, 17.9 and 62.4 for the 50 K, 600 K, GBS and the three technologies combined, respectively (Table 4). Considering the p-value threshold used, 28, 303 and 204 false positives were expected among the 243, 2,953 and 1,182 associations detected for 50 K, 600 K and GBS, respectively. False discovery rate appeared therefore higher for GBS (17.2%) than for DNA arrays (11.5% and 10.2% for 50 K and 600 K, respectively). It could be explained by the higher genotyping error rate of GBS due to imputation and/or by its higher number of markers with a low MAF. Both reduce the power of GBS compared to DNA arrays and therefore lead to a higher false discovery rate. Proportionally to the SNP number, the 50 K and 600 K resulted in 1.5- and 1.7-fold more associated SNPs per situation (environment × trait) than GBS (p-value<2×10^−6^, Table 4). This difference between SNP- arrays and GBS was higher for grain yield (GY) and plant height (plantHT) than for male flowering time (DTA, Table 4).

**Table 3:**
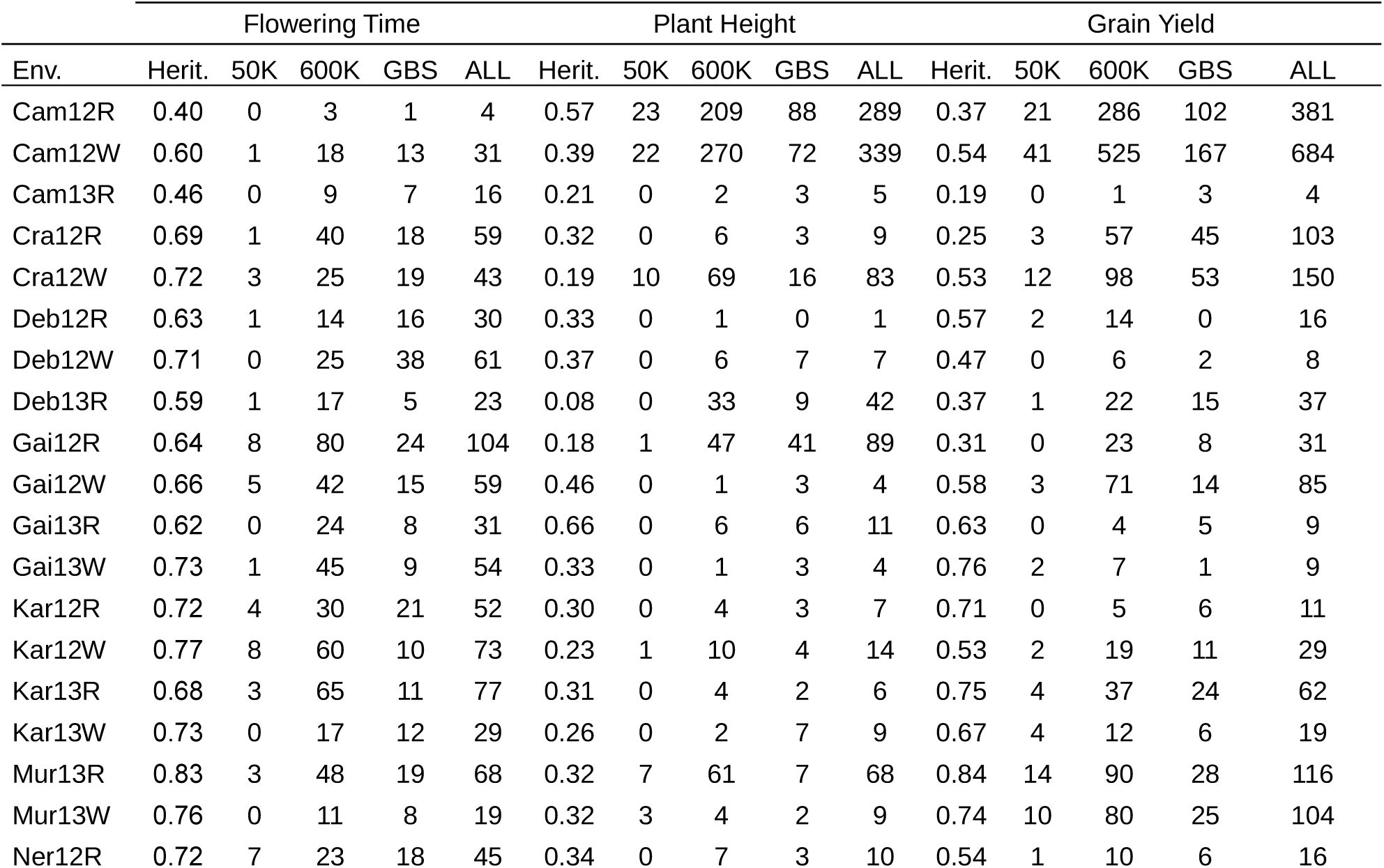
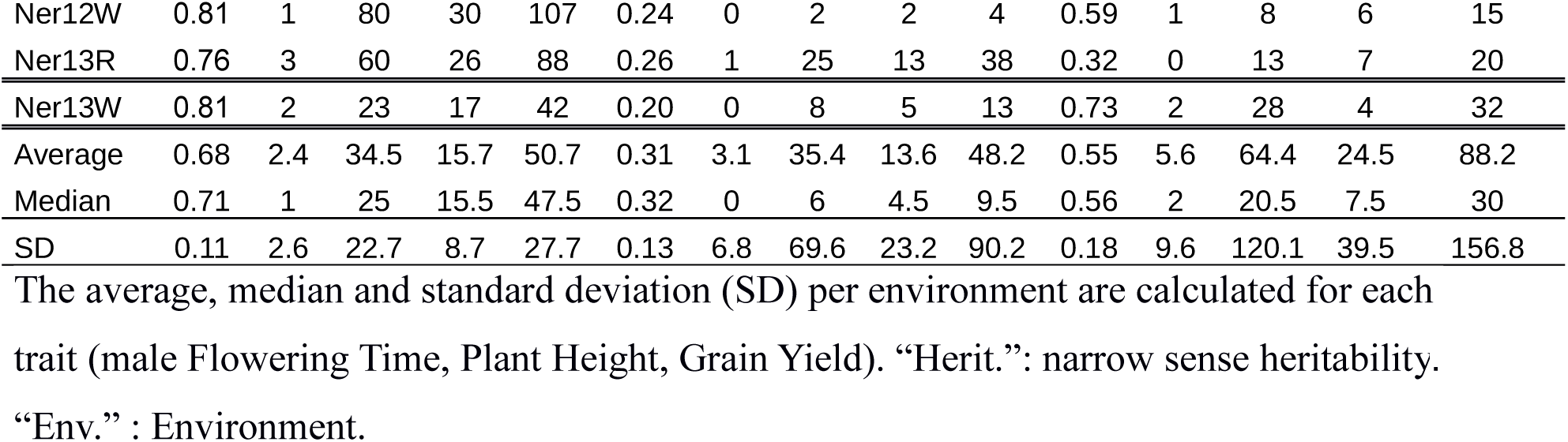
Number of significant SNPs per environment, per technology and for the combined technologies.

**Table 4:** Comparison of associated SNPs and QTLs detected between traits and three technologies.

|  |  | Significant SNPs |  |  |  | QTLs |  |  |  |
| --- | --- | --- | --- | --- | --- | --- | --- | --- | --- |
| Technology |  | 50K | 600K | GBS | ALL | 50K | 600K | GBS | ALL |
| Marker Nb |  | 42046 | 459191 | 308929 | 810580 | 42046 | 459191 | 308929 | 810580 |
| Total Nb | DTA | 52 | 759 | 345 | 1115 | 20 | 130 | 133 | 226 |
|  | plantHT | 68 | 778 | 299 | 1061 | 16 | 96 | 90 | 160 |
|  | GY | 123 | 1416 | 538 | 1941 | 33 | 166 | 120 | 238 |
|  | Per trait | 81 | 984 | 394 | 1372 | 23 | 131 | 114 | 208 |
| Average per env. | DTA | 2.4 | 34.5 | 15.7 | 50.7 | 0.9 | 5.9 | 6.0 | 10.3 |
|  | plantHT | 3.1 | 35.4 | 13.6 | 48.2 | 0.7 | 4.4 | 4.1 | 7.3 |
|  | GY | 5.6 | 64.4 | 24.5 | 88.2 | 1.5 | 7.5 | 5.5 | 10.8 |
|  | Per trait | 3.7 | 44.7 | 17.9 | 62.4 | 1.0 | 5.9 | 5.2 | 9.5 |

QTLs were obtained by grouping associated SNPs with overlapping LD windows ( $LD_{win}$ ) for the three traits (DTA: male flowering time; PlantHT: plant height; GY: grain yield). “Marker Nb” indicates the number of markers tested in GWAS. “Total number”: is the sum of associated SNPs or QTLs across environments. “Average per env” indicates the average number of QTLs obtained in 22 environments for three traits (66 trait-environments combinations). *Average per SNP tested* indicates the number of associated SNPs or QTLs detected divided by the number of SNP tested.

We used two approaches based on LD for grouping significant SNPs (Figure 3): (i) considering that all SNPs with overlapping LD windows for *r^2^K=0.1* belong to the same QTL (*LD_win*) and (ii) grouping significant SNPs that are adjacent on the physical map and are in LD (*r^2^K* > 0.5, *LD_adj*). The QTLs defined by using the two approaches were globally consistent since significant SNPs within QTLs were in high LD whereas SNPs from different adjacent QTLs were not (Additional file 7: Figure S7-LD-Adjacent and Additional file 8: Figure S8-LD-Windows). *LD_adj* detected more QTLs than *LD_win* for flowering time (242 vs 226), plant height (240 vs 160) and grain yield (433 vs 237). The number of QTLs detected with the *LD_adj* approach increased strongly when the LD threshold was set above 0.5. Differences in QTL groupings between the two methods were observed for specific LD and recombination patterns. This occurred for instance on chromosome 6 for grain yield (Additional file 7: Figure S7-LD-Adjacent and Additional file 8: Figure S8-LD-Windows). Within this region, the recombination rate was low and the LD pattern between associated SNPs was complex (Additional file 1: Figure S1). While *LD_adj* splitted several SNPs in high LD into different QTLs (for instance QTL 232, 235, 237, 249), *LD_win* grouped together associated SNPs that are genetically close but displayed a low LD (Additional file 7: Figure S7-LD-Adjacent and Additional file 8: Figure S8-LD-Windows). Reciprocally, for flowering time, we observed different cases where *LD_win* separated distant SNPs in high LD into different QTLs whereas *LD_adj* grouped them (QTL 25 and 26, 51 to 53, 95 to 97, 208 and 209, 218 and 219). As these differences were specific to complex LD and recombination patterns, we used the *LD_win* approach for the rest of the analyses.

Although a large difference in number of associated SNPs was observed between 600 K and GBS, little difference was observed between QTL number after grouping SNPs (Table 3, Table 4). The mean number of QTLs was indeed 1.0, 5.9, 5.2 and 9.5 for the 50 K, 600 K, GBS, and the three technologies combined, respectively (Table 4). Note that the number of QTLs continued to increase with marker density when SNPs from GBS, 50 K and 600 K were combined (Additional file 9: Figure S9). The number of SNPs associated with each QTL varied according to the technology (on average 3.7, 7.6, 3.4 and 6.6 significant SNPs for the 50 K, 600 K, GBS, and the combined technologies, respectively). The total number of QTLs detected over all environments by using the 600 K and GBS was close for flowering time (130 *vs* 133) and plant height (96 *vs* 90). It was 1.4-fold higher for the 600 K than GBS for grain yield (166 *vs* 120).

### The 600 K and GBS were highly complementary for association mapping

Seventy eight percent, 76% and 71% of the QTLs of flowering time, plant height and grain yield were specifically detected by 600 K or GBS, respectively (Figure 4). On the contrary, 50 K displayed very few specific QTLs. When we combined the GBS and 600 K markers, 7% of their common QTLs had -*log_10_(Pval)* increased by 2 and 21% by 1, potentially indicating a gain in accuracy of the position of the causal polymorphism (Additional file 10: Table S1).

**Figure 4:**
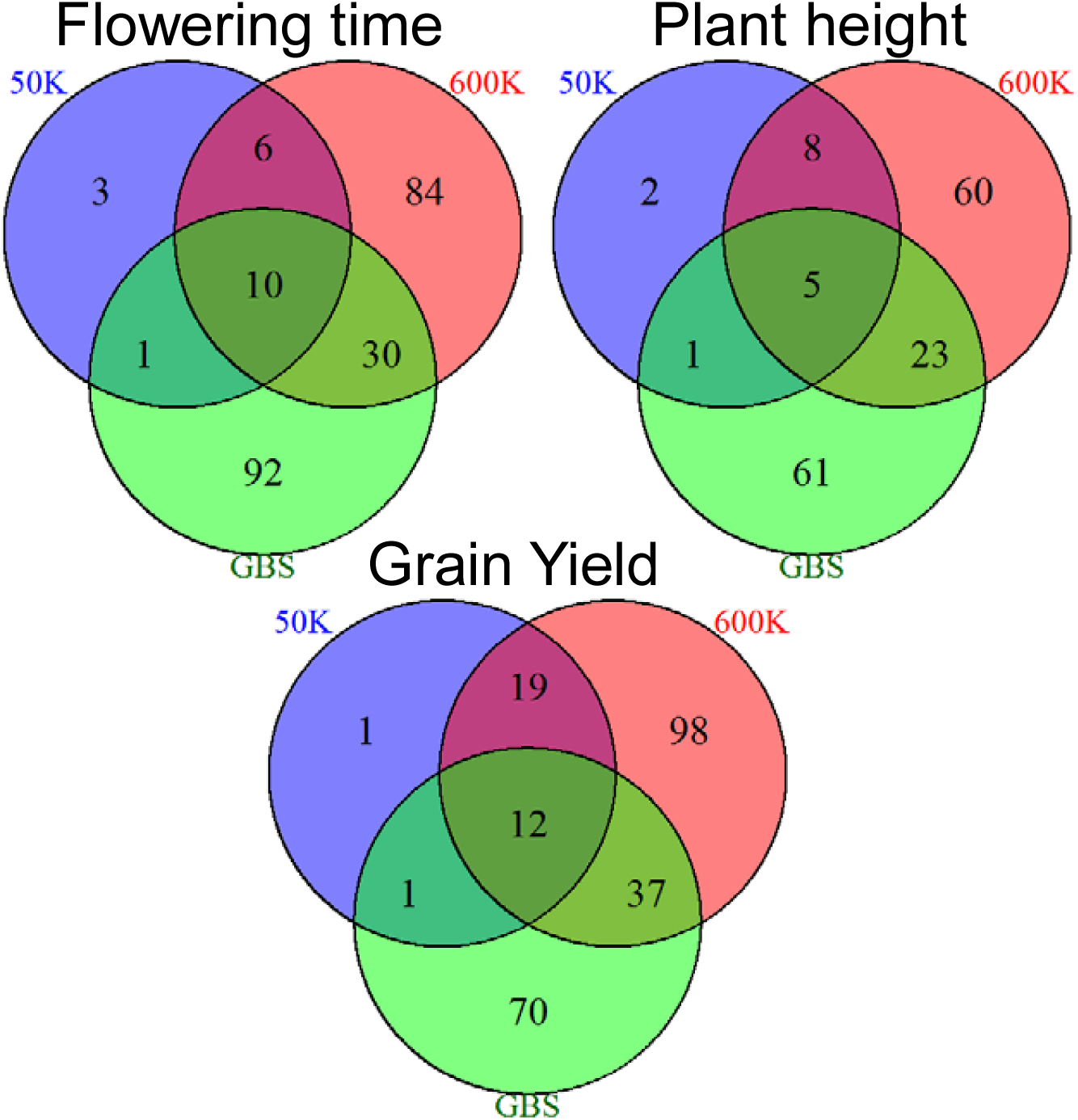
Complementarity of the three technologies to detect QTLs. The numbers of specific QTLs detected by each technology for the three traits (flowering time, plant height, grain yield) are shown.

This complementarity between GBS and 600 K is well exemplified with two strong association peaks for flowering time on chromosome 1 (QTL32) and 3 (QTL95) detected in several environments (Additional file 10: Table S1 and Figure 5a). In order to better understand the origin of the complementarity between GBS and 600 K technologies for GWAS, we scrutinized the LD between SNPs and the haplotypes within these two QTLs (Figure 5b and c, and Additional file 11: Figure S10 for other examples). QTL95 showed a gain in power. It was only identified by the 600 K although the region included numerous SNPs from GBS close to the associated peak. None of these SNPs was in high LD with the most associated marker of the QTL95 (Figure 5b). QTL32 was detected by 1 to 10 GBS markers in 9 environments with *–log(p-value)* ranging from 5 to 7.6, whereas it was detected by only two 600 K markers in one environment (Ner12W) with *–log(p-value)* slightly above the significance threshold (Additional file 10: Table S1 and Figure 5b).

**Figure 5:**
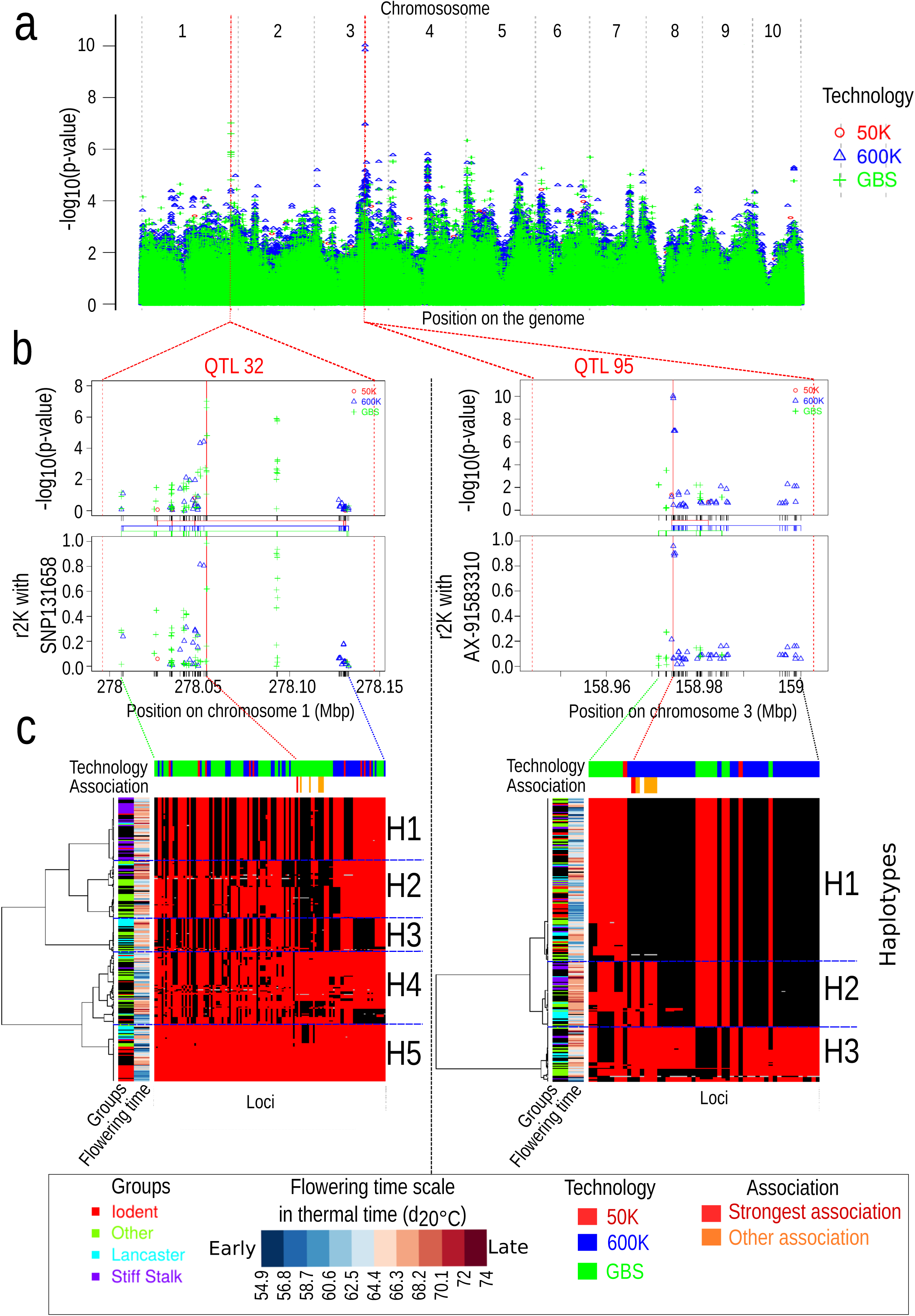
Complementarity of QTLs detection between the 600 K array and the GBS for two regions (QTL 32/QTL95). (**a**) Manhattan plot of the -*log_10_*(*p*-value) along the genome. Dotted red lines correspond to QTL32 and QTL95 located on chromosome 1 and 3, respectively, for the flowering time in one environment (Ner13R). (**b**) Local manhattan plot of the *-log_10_*(*p*-*value*) (top) and linkage disequilibrium corrected by the kinship (*r^2^K*) (bottom) of all SNPs with the strongest associated marker within QTL 32 (left) and QTL 95 (right). Colored vertical lines between manhattan plot and linkage disequilibrium plot represents the distribution of markers for different technologies. Dotted lines between panels b and c linked the first marker, the most associated marker, and the last marker of each QTL (**c**) Local haplotypes displayed by all SNPs within the QTLs 32 (left) and 95 (right) with MAF>5%. Inbred lines are in rows and SNPs are in columns. Inbred lines were ordered by hierarchical clustering based on local dissimilarity estimated by all SNPs within each QTL. Genotyping matrix is colored according to their allelic dose at each SNP. Red and black represent homozygotes and gray represent heterozygotes. The associated peaks (red vertical lines) and other associated SNPs with *-log_10_*(*p-value*) > 5 (orange vertical lines) are indicated above the genotyping matrix. H1, H2, H3, H4, H5 represent the 5 and 3 haplotypes obtained by cutting the dendograms with the most 5 and 3 dissimilar clusters within QTL32 and QTL95, respectively.

Haplotype analyses showed that the SNPs from the GBS within QTL95 were not able to discriminate all haplotypes (Figure 5c). In QTL95, the 600 K markers discriminated the three main haplotypes (H1, H2, H3), whereas using the GBS markers did not discriminate H3 against H1 + H2. As H1 contributed to an earlier flowering time than H2 or H3, associations appeared more significant for the 600 K than for GBS (Figure 5c). In QTL32, the use of GBS markers identified late individuals that mostly displayed H1, H2 and H3 haplotypes, against early individuals that mostly displayed H4 and H5 haplotypes (Figure 5c). The gain of power for GBS markers as compared to 600 K markers for QTL32 originated from the ability to discriminate late individuals (black alleles) from early individuals (red alleles) within H4 haplotypes (Figure 5c).

To further decipher which differences between 600 K and GBS impacted GWAS, we used a resampling approach to explore the interplay between (i) MAF distribution and (ii) SNP distribution along the genome, at different SNP densities. We detected more SNP associations but less QTL with MAF distribution skewed towards low than high MAF. This difference increased as marker density increased (Additional file 12: Figure S11). As GBS has a MAF distribution skewed towards low MAF compared to 600 K, GBS detected more QTLs but less associated SNPs than 600 K. This discrepancy between association and QTL detection came from the fact that QTLs with low MAF were identified by less associated SNP than those with high MAF (Additional file 13: Figure S12).

Regarding distribution along the genome, SNPs distributed similarly to GBS detected more QTLs but less significant SNPs than those following the distribution of 600 K and 50 K, notably for the highest SNP density (Additional file 13: Figure S12). We observed that SNP evenly distributed according to the physical distance detected more associations but less QTLs than all other SNP distributions along the genome. It was the contrary for SNP evenly distributed according to genetic distance (Additional file 12: Figure S11 C and D). This is consistent with QTL distribution along the genome being more correlated to the genetic than physical distance (see below), and the fact that recombination is higher in gene rich regions, leading to less associated SNPs per QTL. Superiority of QTL detection by GBS distribution as compared to 600 K and 50 K SNP distributions came from the higher proportion of SNPs in high recombinogenic regions for GBS than for 600 K and 50 K (Figure 1). This suggests that the complementarity of 600 K and GBS in terms of QTL detected and SNP associations came also from their specificities for both SNP distributions along the genome and MAF distribution. In the end, we studied the impact of genomic coverage differences between 600 K and GBS on QTL detection along the genome. QTLs detected by both 600 K and GBS were located in intervals with large differences in coverage less frequently than their proportion in the entire genome (0.8% vs 7.8%, respectively). Intervals with specific QTLs showed an enrichment in such intervals with high differences in coverage (3.5%) but still below the proportion in the entire genome. It confirmed that most specific QTLs showed no strong genomic coverage differences between GBS and 600 K and therefore that complementarity of QTL detection between these two technologies came from ability to tag different haplotypes.

### Colocalization of QTLs between environments and traits and distribution of QTL along the genome

After combining the three technologies, we identified 226, 160, 238 QTLs for flowering time, plant height and grain yield, respectively (Table 4 and Additional file 10: Table S1). We highlighted 23 QTLs with the strongest effects on flowering time, plant height and grain yield (*-log_10_(Pval) ≥* 8, Table 5). The strongest association corresponded to the QTL95 for flowering time (*-log_10_(p-value)* = 10.03) on chromosome 3 (158,943,646–159,005,990 bp), the QTL135 for GY (*-log_10_(p-value)* = 18.7) on chromosome 6 (12,258,527–29,438,316 bp) and QTL78 on chromosome 6 (12,258,527–20,758,095 bp) for plant height (*-log_10_(p-value)* = 17.31). The QTL95 for flowering time trait was the most stable QTLs across environments since it was detected in 19 environments (Additional file 10: Table S1). Moreover, this QTL showed a colocalization with QTL74 for grain yield in 5 environments and QTL30 for plant height in 1 environment suggesting a pleiotropic effect. More globally, 472 QTLs appeared trait-specific whereas 70 QTLs overlapped between at least two traits (6,3%, 5.2% and 3.0% for GY and plantHT, GY and DTA, and DTA and plantHT, respectively) suggesting that some QTLs may be pleiotropic (Additional file 14: Figure S13). This was not surprising since average corresponding correlations within environments for these traits were moderate (0.47, 0.54 and 0.45, respectively). Only 0.7% overlapped between the three traits (Additional file 14: Figure S13). Twenty percent of QTLs were detected in at least two environments and 9% in at least three environments (Additional file 15: Table S2). We observed no significant differences of stability between the three traits (*p-value* = 0.2). However, 6 out 7 most stable QTLs (Number of environments >5) were found for flowering time. This was consistent with higher average correlations between environments observed flowering time than for plant height and grain yield (0.76, 0.43, 0.48, respectively). We observed that QTLs that displayed a significant effect in more than one environment had larger effects and *-log(p-value)* values than those significant in a single environment. This difference in *-log(p-value)* values was stronger for grain yield and plant height than flowering time.

**Table 5:**
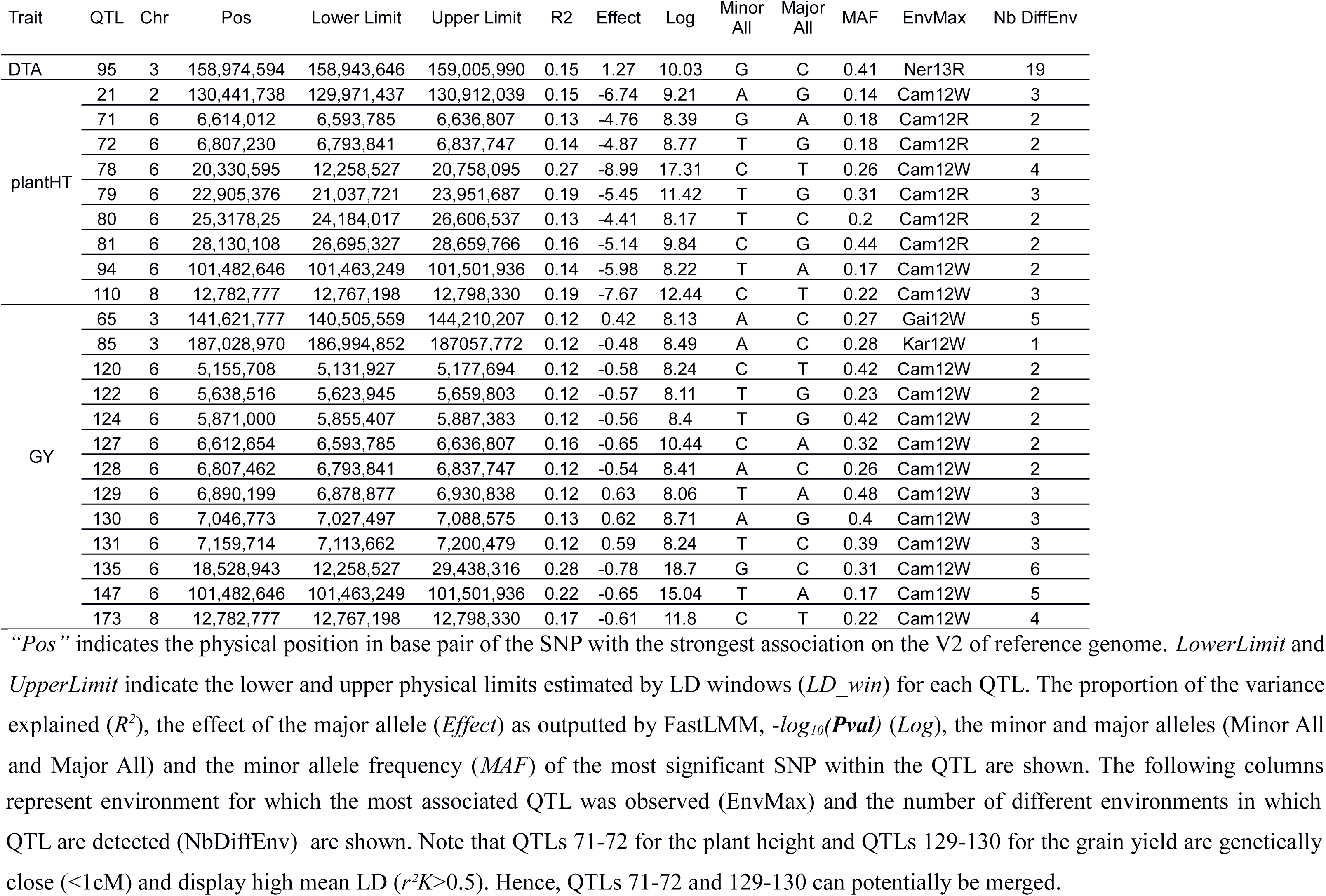
Summary of the main QTLs (*–log_10_(Pval) ≥* 8) identified for the three traits.

The distribution of QTLs was not homogeneous along the genome since 82%, 77% and 79% of flowering time, plant height and grain yield QTLs, respectively, were located in the high recombinogenic regions, whereas they represented 46% of the physical genome (Additional file 16: Table S3). The QTLs were more stable (*≥* 2 environments) in low than in high recombinogenic regions (12.8% vs 5.8%, *p-value* = 0.03).

## Discussion

### GBS required massive imputation but displayed similar global trends than DNA arrays for genetic diversity organization

In order to reduce genotyping cost, GBS is most often performed at low depth leading to a high proportion of missing data, thereby requiring imputation in order to perform GWAS. Imputation can produce genotyping errors that can cause false associations and introduce bias in diversity analysis [33]. We evaluated the quality of genotyping and imputation obtained by different approaches, taking the 50 K or 600 K as references. The best imputation method that yielded a fully genotyped matrix with the lowest error rate for the prediction of both heterozygotes and homozygotes was the approach merging the homozygous genotypes from TASSEL and the imputation of Beagle for the other data (GBS_5_ Table 1). The quality of imputation was high with 96% of allelic values consistent with those of the 50 K and 600 K. This level of concordance is identical to a study of USA national maize inbred seed bank by Romay *et al.* [32]. It is higher than in a diversity study of European flint maize collection (93%) by Gouesnard *et al.* [33], which was more distant from the reference AllZeaGBSv2.7 database than for the panel presented here. For further studies, integrating genotyping data from the three technologies may reduce imputation errors for missing data of GBS [35].

The ascertainment bias of SNP-arrays due to the limited number of lines used for SNP discovery was reinforced by counter-selection of rare alleles during the design process of DNA arrays [3, 4]. For GBS, the polymorphism database to call polymorphisms included thousands of diverse lines [38]. In our study, we used AllZeaGBSv2.7 database. After a first step of GBS imputation (GBS_2_), missing data dropped to 11.9% *i.e.* only slightly more than in Romay *et al.* (10%) [32]. This confirms that the polymorphism database (AllZeaGBSv2.7) covered adequately the genetic diversity of our genetic material.

Although, we observed differences of allelic frequency spectrum between GBS and DNA arrays, these technologies revealed similar trends in the organization of population structure and relatedness (Figure 2, Additional file 4: Figure S4) suggesting no strong ascertainment bias for deciphering global genetic structure trends in the panel. However, although highly correlated, level of relatedness differed between GBS and DNA arrays, especially when the lines were less related as showed by the deviation (to the left) of the linear regression from the bisector (Additional file 5: Figure S5).

### The extent of linkage disequilibrium strongly varied along and between chromosomes

Linkage disequilibrium extent in high recombinogenic regions varied to a large extent among chromosomes, ranging from 0.012 to 0.062 cM. Similar variation of genetic LD extent between maize chromosomes has been previously observed by Rincent et al. [14], but their classification of chromosomes was different from ours. This difference could be explained by the fact that we analyzed specifically high and low recombination regions. According to Hill and Weir Model [40], the physical LD extent in a genomic region increased when the local recombination rate decreased. As a consequence, chromosome 1 and 9 had the lowest and highest physical LD extent and displayed the highest and one of the lowest recombination rate in pericentromeric regions, respectively (0.26 vs 0.11 cM / Mbp, Table 2 and Additional file 16: Table S3). Unexpectedly, the genetic LD extent also correlated negatively with the recombination rate. It suggested that chromosomes with a low recombination rate also display a low effective population size. Background selection against deleterious alleles could explain this pattern since it reduces the genetic diversity in low recombinogenic regions [41, 42]. Finally, we observed a strong variation of the LD extent along each chromosome. As we used a consensus genetic map [43] that represents well the recombination within our population, it suggested, according to Hill and Weir’s model, that the number of ancestors contributing to genetic diversity varied strongly along the chromosomes. This likely reflects the selection of genomic regions for adaptation to environment or agronomic traits [41], that leads to a differential contribution of ancestors according to their allelic effects. Ancestors with strong favorable allele(s) in a genomic region may lead ultimately to large identical by descent genomic segments [44].

### SNPs were clustered into QTL highlighting interesting genomic regions

In previous GWAS studies, the closest associated SNPs were grouped into QTLs according to either a fixed physical distance [1] or a fixed genetic distance [30, 43]. These approaches suffer of two drawbacks. First, the physical LD extent can vary strongly along chromosomes according to the variation of recombination rate (Figure 1 and Additional file 3: Figure S3). Second, the genetic LD extent depends both on panel composition and the position along the genome (Table 2). These approaches may therefore strongly overestimate or underestimate the number of QTLs. To address both issues Cormier *et al.* [46] proposed to group associated SNPs by using a genetic window based on the genetic LD extent estimated by Hill and Weir model in the genomic regions around the associated peaks [40]. In our study, we improved this last approach (*LD_win*):

- First, we used *r^2^K* that corrected r^2^ for kinship rather than the classical r^2^ since *r^2^K* reflected the LD addressed in our GWAS mixed models to map QTL [17].

- Second, we took advantage of the availability of both physical and genetic maps of maize to project the genetic LD extent on the physical map. This physical window was useful to retrieve the annotation from B73 reference genome, decipher local haplotype diversity (Figure 5) and estimate physical genome coverage (Table 2, 1, Additional file 3: Figure S3).

- Third, we considered an average LD extent estimated separately in the high and low recombinogenic genomic regions. This average was estimated by using several large random sets of pairs of loci in these regions rather than the local LD extent in the genomic regions around each associated peaks.

We preferred this approach rather than using local LD extent in order to limit the effect of (i) the strong variation of marker density along the chromosome (Additional file 3: Figure S3), (ii) the local ascertainment bias due to the markers sampling (iii) the poor estimation of the local recombination rate using a genetic map, notably for low recombination regions [3, 44] (iv) errors in locus order due to assembly errors or chromosomal rearrangements.

We compared *LD_win* with *LD_adj,* another approach based on LD to group the SNPs associated to trait variation into QTL. The discrepancies between the two approaches can be explained by the local recombination rate and LD pattern. Since *LD_adj* approach was based on the grouping of contiguous SNPs according to their LD, this approach was highly sensitive to (i) error in marker order or position due to genome assembly errors or structural variations, which are important in maize [47] (ii) genotyping or imputation errors, which we estimated at *ca.* 1% and *ca.* 4%, respectively, for GBS (Table 1), (iii) presence of allelic series with contrasted effects in different experiments which are currently observed in maize [45], (iv) LD threshold used. On the other hand, *LD_win* lead either to inflate the number of QTLs in high recombinogenic regions in which SNPs were too distant genetically to be grouped, or deflated their number by grouping associated SNPs in low recombinogenic regions. Since *LD_win* considered the average LD extent, this method could conduct either to separate or group abusively SNPs when local LD extent was different than the global LD extent. Simulations will be carried out in further research to better understand the properties of *LD_win* and *LD_adj* approach.

Note that LD windows should not be considered as confidence intervals since the relationship between LD and recombination is complex due to demography, drift and selection in association panels, contrary to linkage based QTL mapping [17]. The magnitude of the effect of causal polymorphism in the estimation of these intervals, which is well established for linkage mapping, should be explored further [48]. Other approaches have been proposed to cluster SNPs according to LD [45, 46]. These approaches aim at segmenting the genome in different haplotype blocks separating by high recombination regions. These methods are difficult to use for estimating putative windows inside which the causal polymorphisms are located because such approaches are not centered on the associated SNP.

Several QTLs identified by *LD_win* in our study correspond to regions previously identified: in particular six regions associated with female flowering time [27] and 30 regions associated with different traits in the Cornfed dent panel [11]. Conversely, we did not identify in our study any QTL associated to the florigen *ZCN8*, which showed significant effect in these two previous studies. One of the explanation is that we narrowed the flowering time range in our study, in particular by eliminating early lines. This reduced the representation of the early allele at the Zcn8 locus, leading to a MAF of 0.27 in our study vs. 0.35 in Rincent *et al.* [11], which can slightly diminish the power of the tests [14]. Also, this effect may have strengthened by frequency evolution at loci involved in epistatic interactions with Zcn8 (see [47] for a recent demonstration of such effects).

### Complementarity of 600 K and GBS for QTL detection resulted mostly from the tagging of different haplotypes rather than the coverage of different genomic regions

Number of significant SNPs and QTLs increased with the increase in marker number (Table 4, Additional file 9: Figure S9). This could be explained partly by a better coverage of some genomic regions by SNPs, notably in high recombinogenic regions which showed a very short LD extent and were enriched in QTLs (Additional file 16 Table S3). Numerous new QTLs identified by the 600 K and GBS as compared with those identified by the 50 K were detected in high recombinogenic regions that were considerably less covered by the 50 K than the 600 K or GBS (Figure 1 and Additional file 3: Figure S3).

The high complementarity for QTL detection between GBS and 600 K was only explained to a limited extent by the difference of the SNP distribution and density along the genome, since these two technologies targeted similar regions as showed by coverage analysis (Figure 1 and Additional file 3: Figure S3). However, at a finer scale, SNPs from the 600 K and GBS could tag close but different genomic regions around genes. SNPs from the 600 K were mostly selected within coding regions of genes [4], whereas SNP from GBS targeted more largely low copy regions, which included coding but also regulatory regions of genes [32, 38]. To further analyse the complementarity of the technologies, we analysed local haplotypes and the effect of genome coverage differences between technologies on QTL detection. We showed that both technologies captured different haplotypes when similar genomic regions were targeted (Figure 5). In this figure, two QTLs were specifically detected by markers from either 600 K or GBS although there are several markers from other technology very close from the most associated marker, considering the size of LD windows around it. Additionally, we did not observe an enrichment of QTL specifically detected by one technology in 20kbp-intervals with high genomic coverage difference between 600 K and GBS Hence, we pinpointed that GBS and DNA arrays are highly complementary for QTL detection because they tagged different haplotypes rather than different regions (Figure 5). Based on the L-shaped MAF distribution, which suggests no ascertainment bias, and the high number of sequenced lines used for the GBS, we expect a closer representation of the variation present in our panel by this technology compared to the 600 K, but this comes to the cost of an enrichment in rare alleles. Both factors tend to counterbalance each other in terms of GWAS power (Additional file 13: Figure S12).

Our results suggest that we did not reach saturation with our *c.* 800,000 SNPs because (i) some haplotypes certainly remain not tagged (ii) the genome coverage was not complete, and (iii) the number of significant SNPs and QTLs continued to increase with marker density (Additional file 9: Figure S9). Considering LD and marker density, the genotypic data presently available were most likely enough to well represent polymorphisms in the centromeric regions, whereas using more markers would be beneficial for telomeric regions. New approaches based on resequencing of representative lines and imputation are currently developed to achieve this goal.

## Methods

### Plant Material and Phenotypic Data

The panel involves 247 maize inbred lines, further referred to as DROPS panel (Additional file 17: Table S4). They include 164 lines from a wider panel of lines from Europe and America [11] and 83 additional lines derived from public breeding programs in Hungary, Italy and Spain and recent lines free of patent from the USA. All lines belong to the dent genetic group, which can be subdivided in different sub-groups (see [11, 30]). Lines were selected within a restricted flowering time window (10 days) in order to limit the effect of drought escape due to flowering time variation in the identification of genomic regions involved in drought tolerance [30]. Candidate lines with poor sample quality, i.e. high level of heterozygosity, or high relatedness with other lines were discarded in this selection. The lines selection was also guided by pedigree to avoid as far as possible over-representation of some ancestral materials.

The 247 inbred lines were all crossed with a common line (UH007) from the Flint genetic groups to obtain 247 hybrids (hybrid panel). Dent and Flint genetic groups are known to be complementary to produce hybrids [48]. Further, as UH007 is unrelated to any line in the panel, no hybrid is affected by inbreeding depression. This guarantees that hybrids have a level of performance and an overall physiology comparable to that of varieties used in agriculture. Conversely, field evaluation of inbred lines per se would have diminished yield by more than 50%.

Experimental design and model used for obtaining adjusted means for male flowering time (Day To Anthesis, DTA), plant height (plantHT), and grain yield (GY) were previously described [30]. While DTA and GY were previously analyzed in [30], PlantHT was not. Briefly, the hybrid panel were evaluated for these three traits in 22 experiments (combination year x site x water regime), *i.e.* at seven sites in Europe, during two years (2012 and 2013), and for two water treatments (watered and rainfed) [30]. Experiments were designed as alpha-lattice designs with two and three replicates for watered and rain-fed regimes, respectively. Grain yield (t ha^−1^) was adjusted to 15% moisture. The adjusted mean (Best Linear Unbiased Estimation, BLUEs, https://doi.org/10.15454/IASSTN) of the three traits were estimated per environment (site × year × water regime) using a mixed model based on fixed hybrid and replicate effects, random spatial effects (rows and columns), and spatially correlated errors in order to take into account spatial variation of micro-environment in each field trial (see [30] for more details). The same model, but with random hybrids effects, was used to estimate variance components. Models were fitted with ASReml-R [49]. Narrow-sense heritability of each trait in each environment were also estimated as in [30] (Additional file 18: Table S5). As all hybrids share a common parent (UH007), adjusted means (BLUEs) of hybrids were combined with genotyping data of the corresponding dent inbred lines of the panel to perform GWAS, following a usual practice in maize genetics [11].

### Genotyping and Genotyping-By-Sequencing Data

The 247 inbred lines were genotyped using three technologies: a maize Illumina Infinium HD 50 K array [3], a maize Affymetrix Axiom 600 K array [4], and Genotyping-By-Sequencing [2, 38]. In the arrays, DNA fragments are hybridized with probes attached to the array (Additional file 19: Notes S1 for the description of the data from the two SNP-arrays). Genotyping-by-sequencing technology is based on multiplex resequencing of tagged DNA using restriction enzyme (Keygene N.V. owns patents and patent applications protecting its Sequence Based Genotyping technologies) [2]. Cornell Institute (NY, USA) processed raw sequence data using a multi-step Discovery and a one-step Production pipeline (*TASSEL-GBS*) in order to obtain genotypes (Additional file 19: Notes S1). An imputation step of missing genotypes was carried out by Cornell Institute [39], which utilized an algorithm that searches for the closest neighbour in small SNP windows across the haplotype library [38].

We applied different filters (heterozygosity rate, missing data rate, minor allele frequency) for a quality control of the genetic data before performing the diversity and association genetic analyses. For GBS data, the filters were applied after imputation using the method “Compilation of Cornell homozygous genotypes and Beagle genotypes” (GBS_5_ in Additional file 1: Figure S1; See section “Evaluating Genotyping and Imputation Quality”). We eliminated markers that had an average heterozygosity and missing data rate higher than 0.15 and 0.20, respectively, and a Minor Allele Frequency (MAF) lower than 0.01 for the diversity analyses and 0.05 for the GWAS (Additional file 20: Table S6). Individuals which had heterozygosity and/or missing data rate higher than 0.06 and 0.10, respectively, were eliminated.

### Evaluating Genotyping and Imputation Quality

Estimation of genotyping and imputation quality was performed using the entire panel except two inbred lines that had different seedlots between technologies. The 50 K and the 600 K were taken as reference to compare the concordance of genotyping (genotype matches) with the imputation of GBS based on their position. While SNP positions and orientation from GBS were called on the reference maize genome B73 AGP_v2 (release 5a) [50], flanking sequences of SNPs in the 50 K were primary aligned on the first maize genome reference assembly B73 AGP_v1 (release 4a.53) [51]. Both position and orientation scaffold carrying SNPs from the 50 K can be different in the AGP_v2, which could impair correct comparison of genotype between the 50 K and GBS. Hence, we aligned flanking sequences of SNPs from the 50 K on maize B73 AGP_v2 using the Basic Local Alignment Search Tool (BLAST) to retrieve both positions and genotype in the same and correct strand orientation (forward) to compare geno-typing. The number of common markers between the 50 K/600 K, 50 K/GBS, GBS/600 K and 50 K/600 K/GBS was 36,395, 7,018, 25,572 and 5,947 SNPs, respectively. The comparison of the genotyping and imputation quality between the 50 K/GBS, 50 K/600 K and 600 K/GBS was done on 5,336 and 24,286 and 26,154 common markers, respectively. The comparison for the 50 K involved PANZEA markers, prefixed as “PZE” [52]. In order to achieve these comparisons, we considered the direct reads from GBS (**GBS_1_**) and four approaches for imputation (GBS_2_ to GBS_5_, Additional file 1: Figure S1). **GBS_2_** approach consisted of one imputation step from the direct read by Cornell University, using *TASSEL* software, but missing data was still present. **GBS_3_** approach consisted of imputation by *Beagle v3* [13] of the missing data of GBS_1_. To compare data from GBS_3_ and GBS_2_ to those of the 50 K and 600 K, missing data in GBS_2_ were excluded from GBS_3_. In **GBS_4_,** genotype imputation by Beagle was performed on Cornell imputed data after replacing the heterozygous genotypes with missing data. **GBS_5_**, consisted of homozygous genotypes of GBS_2_ completed by values imputed in GBS_3_, no missing data remained (Additional file 1: Figure S1).

### Diversity Analyses

After excluding the unplaced SNPs and applying the filtering criteria for the diversity analyses (MAF > 0.01), we obtained the final genotyping data of the 247 lines with 44,729 SNPs from the 50 K, 506,662 SNPs from the 600 K array, and 395,024 SNPs from the GBS (Additional file 20: Table S6). All markers of the 600 K and GBS_5_ that passed the quality control were used to perform the diversity analyses (estimation of Q genetic groups and K kinships). For the 50 K, we used only the PANZEA markers (29,257 SNPs) [52] in order to reduce the ascertainment bias noted by Ganal *et al.* [3] when estimating Nei’s index of diversity [53] and relationship coefficients. Genotypic data generated by the three technologies were organized as *G* matrices with *N* rows and *L* columns, *N* and *L* being the panel size and number of markers, respectively. Genotype of individual *i* at marker *l* (*G_i,l_*) was coded as 0 (the homozygote for an arbitrarily chosen allele), 0.5 (heterozygote), or 1 (the other homozygote). Identity-By-Descent (IBD) was estimated according to Astle and Balding [19]:

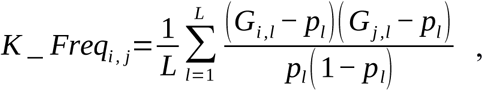

where *p_l_* is the frequency of the allele coded 1 of marker *l* in the panel of interest, *i* and *j* indicate the inbred lines for which the kinship was estimated. We also estimated the Identity-By-State (IBS) by estimating the proportion of shared alleles. For GWAS, we used *K_Chr* [14] that are computed using similar formula as *K_Freq*, but with the genotype data of all the chromosomes except the chromosome of the SNP tested. This formula provides an unbiased estimate of the kinship coefficient and weights by allelic frequency assuming Hardy-Weinberg equilibrium. Hence, relatedness is higher if two individuals share rare alleles than common alleles.

Genetic structure was analysed using the sofware *ADMIXTURE v1.22* [18] with a number of groups varying from 2 to 10 for the three technologies. We compared assignation by *ADMIXTURE* of inbred lines between the three technologies by estimating the proportion of inbred lines consistently assigned between technologies two by two (50 K *vs* GBS_5_, 50 K *vs* 600 K, 600 K *vs* GBS_5_) using a threshold of 0.5 for admixture.

Expected heterozygosity (He) [53] was estimated at each marker as *2p_l_(1 − p_l_)* and was averaged on all the markers for a global characterization of the panel for the three technologies. Principal Coordinate Analyses (PCoA) were performed on the genetic distance matrices [54], estimated as *1_N,N_ − K_Freq*, where *1_N,N_* is a matrix of ones of the same size as *K_Freq*.

### Linkage Disequilibrium Analyses

We first analyzed the effect of the genetic structure and kinship on linkage disequilibrium (LD) extent within and between chromosomes by estimating genome-wide linkage disequilibrium using the 29,257 PANZEA SNPs from the 50 K. Four estimates of LD were used: the squared correlation (*r^2^*) between allelic dose at two markers [55], the squared correlation taking into account global kinship with *K_Freq* estimator (*r^2^K*), the squared correlation taking into account population structure (*r^2^S*), and the squared correlation taking into account both (*r^2^KS*) [17].

To explore the variation of LD decay and the stability of LD extent along the chromosomes, we estimated LD between a non-redundant set of 810,580 loci from the GBS, the 50 K and 600 K. To save computation time, we calculated LD between loci within a sliding window of 1 cM. Genetic position was obtained by projecting the physical position of each locus using a *smooth.spline* function R calibrated on the genetic consensus map of the Cornfed Dent Nested Association Mapping (NAM) design [56]. We used the estimator *r^2^* and *r^2^K* using 10 different kinships *K_Chr*. This last estimator was calculated because it corresponds exactly to the LD used to map QTL in our GWAS model. It determines the power of GWAS to detect QTL considering that causal polymorphisms were in LD with some polymorphisms genotyped in our panel [17]. To study LD extent variation, we estimated LD extent by adjusting Hill and Weir’s model [40] using non-linear regression (*nls* function in R-package *nlme*) against both physical and genetic position within each chromosome. Since recombination rate (cM / Mbp) varied strongly along the genome (Figure 1 and Additional file 3: Figure S3), we defined high (>0.5 cM / Mbp) and low (<0.5 cM / Mbp) recombinogenic genomic regions within each chromosome. We adjusted Hill and Weir’s model [40] separately in low and high recombinogenic regions (Additional file 16: Table S3)by randomly sampling 100 sets of 500,000 pairs of loci distant from less than 1 cM. This random sampling avoided over-representation of pairs of loci from low recombinogenic regions due to the sliding-window approach (Figure 3). 500,000 pairs of loci represented 0.36% (Chromosome 3 / High rec) to 1.20% of all pairs of loci (Chromosome 8 / High rec).

For all analyses, we estimated LD extent by calculating the genetic and physical distance for the fitted curve of Hill and Weir’s Model that reached *r^2^K*=0.1, *r^2^K*=0.2 and *r^2^K*=0.4.

### Genome coverage estimation

In order to estimate the genomic regions in which the effect of an underlying causal polymorphisms could be captured by GWAS using LD with SNP from three technologies, we developed an approach to define LD windows around each SNP with MAF ≥ 5% based on LD extent (Figure 3). To set the LD window around each SNP, we used LD extent with *r^2^K*=0.1 (negligible LD), *r^2^K*=0.2 (intermediate LD) and *r^2^K*=0.4 (high LD) estimated in low and high recombinogenic regions for each chromosome. We used the global LD decay estimated for these large chromosomal regions rather than local LD extent (i) to avoid bias due to SNP sampling within small genomic regions, (ii) to reduce computational time, and (iii) to limit the impact of possible local error in genome assembly. In low recombinogenic regions, we used the physical LD extent, hypothesizing that recombination rate is constant along physical distance in these regions. In high recombinogenic regions, we used the genetic LD extent since there is a strong variation of recombination rate by base pair along the physical position (Figure 1 and Additional file 3: Figure S3). We then converted genetic LD windows into physical windows by projecting the genetic positions on the physical map using the *smooth.spline* function implemented in R, calibrated on the NAM dent consensus map [56]. Reciprocally, we obtained the genetic positions of LD windows in low recombinogenic regions by projecting the physical boundaries of LD windows on the genetic map.

To estimate coverage of the three technologies to detect QTLs based on their SNP distribution and density, we calculated cumulative genetic and physical lengths that are covered by LD windows around the markers, considering different LD extents for each chromosome (*r^2^K*=0.1, *r^2^K*=0.2, *r^2^K*=0.4). In order to explore variation of genome coverage along the chromosome, we estimated the proportion of genome covered using a sliding-windows approach based on variable physical distances (20, 100, 500, 2000 kbp) considering LD extent for a r2 K =0.1.

### Statistical Models for Association Mapping

We used four models to determine the statistical models that control best the confounding factors (*i.e.* population structure and relatedness) in GWAS (Additional file 21: Notes S2). We tested different software implementing either approximate (EMMAX) [8] or exact computation of standard test statistics (ASReml and FaST-LMM) [6, 49] for computational time and GWAS results differences. Single-trait, single-environment GWAS was performed for each marker for each environment and all traits using FaST-LMM. We selected the mixed model using *K_Chr*, estimated from PANZEA markers of the 50 K to perform GWAS on 66 situations (environment × trait) (Additional file 21: Notes S2). We developed a GWAS pipeline in *R v3.2.1* [55] calling FaST-LMM software and implementing [14] approaches to conduct single trait and single environment association tests.

To take into account multiple tests in GWAS and their dependence, we applied the methods of Moskvina and Schmidt [56] and Gao *et al.* [57, 58] to infer the number of independent tests to be considered in the Bonferroni formula. Using the Gao et al. [57, 58] approaches, we estimated the number of independent tests for GWAS at 15,780 for the 50 K, 92,752 for the 600 K, 109,117 for the GBS_5_ and 191,026 for the combined genetic data (i.e. merging of 50 K, 600 K, GBS), leading to different *-log_10_(p-value)* thresholds: 5.49, 6.27, 6.34 and 6.58, respectively. Because of these differences, we used two thresholds of *-log_10_(p-value) = 5* (less stringent) *and 8* (hightly conservative and slightly above Bonferroni) for comparing GWAS to avoid the differences of identification of significant SNPs between the technologies due to the choice of the threshold.

### Methods for grouping associated SNPs into QTLs

We used two approaches based on LD for grouping significant SNPs. The first approach (*LD_win*) used LD windows, previously described, to group significant SNPs into QTLs considering that all significant SNPs with overlapping LD windows of *r^2^K=0.1* belong to the same QTL (Figure 3). We hypothesized that significant SNPs with overlapping LD windows at *r^2^K=0.1* captured the same causal polymorphism and were therefore a single and unique QTL. For the second approach (*LD_adj*), significant SNPs were grouped into a same QTL if they were connected in terms of LD (*r^2^K* between adjacent significant SNPs superior to *0.5*). We used LD heatmaps for comparing the SNP grouping produced by the two approaches on the three different traits across all environments (Additional file 7: Figure S7-LD-Adjacent and Additional file 8: Figure S8-LD-Windows). All scripts are implemented in R software [59].

### Resampling approach to analyze effect of MAF distribution, SNP distribution along the genome, SNP density on QTL detection

To study the effect of SNP density, MAF distribution and SNP distribution along the genome on association and QTL detection, we used a resampling approach of several sets of SNPs displaying different MAF distribution and SNP distribution along the chromosome. We compared these modalities with different SNP densities (50,000, 100,000, 150,000, 200,000, 250,000 markers). In this resampling approach, we considered all markers together and that both associations and QTLs detected by the whole SNP sets are true. We selected only markers having MAF above 5%. To study the effect of MAF distribution on QTL detection SNPs were classified in 5 MAF classes (0-0.1, 0.1-0.2, 0.2-0.3, 0.3-0.4 and 0.4, 0.5) and SNP were randomly selected in each classes according to MAF distribution t1) similar to GBS (GBS_MAF) 2) similar to 600 K (600K_MAF) 3) with equal frequency for 5 MAF classes (Flat_MAF) 4) skewed towards high MAF (High_MAF) with SNP frequency of 0, 0, 0.2, 0.4, 0.4 in (0-0.1], (0.1-0.2], (0.2-0.3], (0.3-0.4], (0.4-0.5] MAF classes, respectively 5) skewed towards low MAF (Low_MAF) with SNP frequency of 0, 0, 0.2, 0.4, 0.4 in (0-0.1], (0.1-0.2], (0.2-0.3], (0.3-0.4], (0.4-0.5] MAF classes, respectively.

To study the effect of SNP distribution along the genome on QTL detection, we compared 5 different SNP distributions along the chromosome: 1) evenly distributed according to the physical distance (Dens_Phys), 2) evenly distributed according to the genetic distance (Dens_Gen), 3) distributed like GBS (Dens_GBS), 4) distributed similarly to 600 K (Dens_600 K) 5) distributed like 50 K (Dens_50 K). SNPs were sampled randomly according to the different densities in contiguous windows of 10Mbp.

## List of abreviations

DTA: Day to Anthesis
GY: Grain Yield adjusted at 15% moisture
plantHT: Plant Height
GBS: Genotyping By Sequencing
LD: Linkage disequilibrium
GWAS: Genome-Wide Association Studies
MAF: Minimum Allelic Frequency
SNP: Single Nucleotide Polymorphism
HRR: High Recombinogenic Regions
LRR: Low Recombinogenic Regions
QTL: Quantitative Trait Locus

## Declarations

### Ethics approval and consent to participate

Not applicable.

### Consent for publication

Not applicable.

### Availability of data and material

The following links toward the data will be available upon publication of this paper.

All the genotyping data used in this study can be found at https://doi.org/10.15454/AEC4BN.

*Genotyping data will become publicly available upon the publication with link above. Genotyping data will be accessible anonymously with following private link for reviewers:* https://data.inra.fr/privateurl.xhtml?token=dc792926-6996-4767-bbbe-bb347fd1edd4

The GWAS results can be found at https://doi.org/10.15454/6TL2N4.

*GWAS results will become publicly available upon the publication with link below. GWAS results will be accessible anonymously with following private link for reviewers:* https://data.inra.fr/privateurl.xhtml?token=1b99f286-94c8-40d1-a289-c18db1d6e646

The phenotypic dataset can be found at https://doi.org/10.15454/IASSTN.

### Authors’ contributions

S.S.N., S.D.N. and A.C., designed the studied and wrote the article. S.S.N. performed genotyping data quality control, imputation and genetic analyses. S.D.N. developed and performed LD analyses. A.C. designed the association panel with the help of S.D.N. and C.W. C.B. participated in assembling the dent inbred lines panel, organizing the germplasms and field work for seeds production. E.J.M., C.W. and F.T. collected and analysed the phenotypic data. V.C. and D.M. performed DNA extraction and prepared the samples. All authors critically reviewed and approved the final manuscript.

## Acknowledgements

We are grateful to key partners from the field: Pierre Dubreuil, Cécile Richard, Jérémy Lopez (Biogemma), Tamás Spitkó (MTA ATK), Therese Welz (KWS), Franco Tanzi, Ferenc Racz, Vincent Schlegel (Syngenta) and Maria Angela Canè (UNIBO). We also acknowledge Björn Usadel and Axel Nagel (MPI) for data management. We thank Willem Kruijer, Fred Van Eeuwijk (WUR), Tristan Mary-Huard and Laurence Moreau (INRA) for helpful discussions and statistical advice. We are grateful to Chris-Carolin Schön (TUM) for providing an early access to the Affymetrix Axiom 600 K array and Edward Buckler (USDA) for providing genotyping using GBS. We are also grateful to partners of the CornFed project, Univ. Hohenheim (Germany), CSIC (Spain), CRAG (Spain), MTA ATK (Hungary), NCRPIS (USA), CRB Maize (France) and CRA-MAC (Italy) who contributed to the genetic material.

## Authors’ information (optional)

Not applicable.

## Additional file legends

**Additional file 1 (.docx):**

**Figure S1:** Different approaches used to impute missing data of the GBS. We considered the direct reads from GBS (**GBS_1_**) and four approaches for imputation (GBS_2_ to GBS_5_). **GBS_2_** approach consisted in one imputation step from the direct read by Cornell University, using *TASSEL* software, but missing data was still present. **GBS_3_** approach consisted in a genotype imputation of the whole missing data of the direct read by *Beagle v3*. In **GBS_4_,** genotype imputation by Beagle was performed on Cornell imputed data after replacing the heterozygous genotypes into missing data. **GBS_5_**, consisted in homozygous genotypes of GBS_2_ completed by values imputed in GBS_3_.

**Additional file 2 (.docx):**

**Figure S2:** Comparison of genotyping data between 50 K and 600 K arrays, and GBS. (a) Distribution of minor allele frequency per SNP before filtering (monomorphic SNPs removed). (b) Distribution of SNP missing data proportion for the 50 K array, 600 K array, GBS direct reads (GBS_1_) and GBS after imputation by Cornell Institute (GBS_2_, note that the scale of the x-axes is different). (c) Relatedness distribution (Identity-By-State, IBS) after QC filtering with MAF≥1% (IBS using GBS_1_ was not estimated because of the low calling rate).

**Additional file 3 (.pdf):**

**Figure S3:** Variation of the markers density, the recombination rate and the genome coverage in non-overlapping 2 Mbp windows along each chromosome except chromosome 3 (presented in Figure 1). Markers have MAF above 5%. Top panel shows the variation of SNP number. In the bottom panel, dotted line represents the variation of recombination rate (cM / Mbp) and solid lines the proportion of genome covered by the SNPs using the cumulated length of physical LD windows around each SNP in each 2Mbp-windows. In these two panel, green, blue, red and black lines represent variation for GBS, 600 K, 50 K and combined technologies, respectively. Vertical dotted gray lines indicate limits of centromeric regions. Vertical lines between the two panels indicate the position of QTLs for flowering time (DTA), grain yield (GY) and Plant Height (PHT). Green, blue, red vertical lines indicate QTLs detected only by GBS, 600 K and 50 K technologies, respectively. Grey vertical lines indicate QTL detected by at least two technologies. Only QTL including a marker associated with −log_10_(pval) above 6 were shown.

**Additional file 4 (.docx):**

**Figure S4:** Contribution of four ancestral populations to 247 inbred lines after ADMIXTURE analysis. Markers from the 50 K (top), 600 K (middle) and GBS (bottom) were used. One vertical bar corresponds to one individual. Lines were ordered according to contributions observed for the 50 K. From left to right, we have Stiff Stalk lines type B73 and B14a (blue), Iodent lines type PH207 (red), Lancaster lines type Mo17 and Oh43 (turquoise), a group of lines assembling W117, F7057 type lines (green).

**Additional file 5 (.docx):**

**Figure S5:** Correlation between kinship matrix estimated by different technologies. Correlation (*r*) between the IBS and IBD (*K_Freq*) for each technology (A) and correlation of IBD (B) and IBS (D) between the three technologies (after imputation). (C) Correlation of IBD between the three technologies after removing the excess of rare alleles in the GBS to have the same distribution of MAF as in the 50 K and the 600 K. The red line is the bisector.

**Additional file 6 (.docx):**

**Figure S6:** Heatmap of genome-wide linkage disequilibrium (LD) between all markers within and between chromosomes using PANZEA SNPs from the 50 K. All SNPs were ordered according to their position on the genome. Dots represent LD between two loci and were colored according to their strength. Classical LD measurement r^2^ between loci were represented within triangle below the diagonal. Linkage disequilibrium corrected for structure (*r^2^S*, A), relatedness (*r^2^K*, B) or both (*r^2^KS*, C) were represented within triangle above the diagonal.

**Additional file 7 (.pdf):**

**Figure S7:** QTL limits obtained by the *LD_Adj* approach projected on heatmaps representing the level of LD between associated SNPs for each trait (DTA: male flowering time, plantHT: plant height and GY: grain yield) for each chromosome. Upper and lower triangles on the heatmaps represent the r^2^ and r^2^K values between associated SNPs, respectively. Linkage disequilibrium between loci was colored according to values from weak LD (yellow) to high LD (red). The significant markers were ordered according to their physical positions on the chromosome and were represented by ticks on the four sides of the heatmaps. Limits of QTLs were displayed by gray dotted lines. QTL numbers were indicated in gray on the top and the right of each heatmap.

**Additional file 8 (.pdf):**

**Figure S8-LD_Windows:** QTL limits obtained by the *LD_win* approach projected on heatmaps representing the level of LD between associated SNPs for each trait (DTA: male flowering time, plantHT: plant height and GY: grain yield) and each chromosome. Upper and lower triangles on the heatmaps represented the r^2^ and r^2^K values between associated SNPs, respectively. Linkage disequilibrium between loci was colored according to values from weak LD (yellow) to high LD (red). The significant markers were ordered according to their physical positions on the chromosome and were represented by ticks on the four sides of the heatmaps. Limits of QTLs were displayed by gray dotted lines. QTL numbers were indicated in gray on the top and the right of each heatmap.

**Additional file 9 (.docx):**

**Figure S9:** Number of significant SNPs (blue line) and QTLs (red line) identified as a function of SNP density (x-axis) for the three traits (DTA, male flowering time; plantHT, plant height; GY, grain yield).

**Additional file 10 (.csv):**

**Table S1:** Summary of all the QTLs identified for the male flowering time (DTA), plant height (plantHT) and grain yield (GY). “LowerLimit” and “UpperLimit” columns are the lower and upper physical limits for each QTL. The “Rec” column indicates if the QTL is located in a high or low region of recombination. “NbSNP50”, “LogPvaMax50”, “NbSNP600”, “LogPvaMax600”, “NbSNPGBS”, “LogPvaMaxGBS” are the number of significant SNPs and the most significant *–log_10_(Pval)* within the QTL for each technology across all environments. The physical position (“PosMax”), the proportion of the variance explained (“R2_LDMax”) and the effect (“EffectMax”) of the most significant SNP within the QTL is shown. “NbDiffEnv” gives the number of different situations that detected the QTL.

**Additional file 11 (.docx):**

**Figure S10:** Examples of QTL detection on Chromosome 3, 6 and 8 for the different traits. The top panel represents the distribution of the QTLs along the chromosome of interest, for the different technologies. The vertical red line in this panel localizes the SNP chosen as reference for the QTL (marker with the strongest association). The middle panel is a zoom in the vicinity of the reference SNP, showing the Local distribution of the -*log_10_*(*p-value*). The bottom panel is the same zoom as the middle panel and shows the local linkage disequilibrium corrected by the kinship (*r^2^k*) of all SNPs, within this region, within the reference SNP. Ticks on different x-axes show the marker density of the three technologies (red for the 50 K, blue for the 600 K and green for the GBS).

**Additional File 12 (.pdf)**

**Figure S11:** Effect of minor allelic frequency distribution, SNP distributions along the genome and SNP densities on the number of associated SNP and QTL detected. Boxplot were drawn on 100 sets of 50 000 to 250 000 markers sampled according to different MAF distributions (A, B) and different SNP distributions along the genome (C, D). A, C: number of SNP associated; B, D: Number of QTL detected. In A and B, 600K_MAF (yellow), GBS_MAF (green), Low_MAF (cyan), Flat_MAF (blue), High_MAF (pink) on x axis indicate boxplots corresponding to MAF distribution similar to 600 K, similar to GBS, skewed towards low MAF, flat MAF and skewed toward high MAF, respectively. In C and D, Dens_50 K (red), Dens_600 K (yellow), Dens_GBS (cyan), Dens_Gen (blue), Dens_Phys (pink) on x axis indicate distribution of SNPs along the genome corresponding to 50 K, GBS, 600 K, even genetic and physical distances, respectively. For A, B, C and D, modalities indicated as “Random” in x axis correspond to random sample of SNP. Number of markers for each boxplot are indicated after the point.

**Additional file 13 (.pdf)**

**Figure S12:** Distribution of markers, associations and QTLs according to MAF for 50 K, 600 K GBS, and ALL technologies. A) Number of markers, B) Proportion of markers, C) Proportion of Association, D) Proportion of QTLs

**Additional file 14 (.docx):**

**Figure S13:** Colocalization of QTLs between the traits. Number of QTLs specific and shared by the three traits across all environments. Note that several QTLs from one trait were sometimes included in a single QTL of another trait.

**Additional file 15:**

**Table S2:** Stability of QTLs across environments for the three traits (DTA: male flowering time, plantHT: Plant Height, GY: Grain Yield) and all traits. “Env. Nb” indicates the number of environment in which a QTL was detected. Next four columns indicate the number of QTL corresponding to each cartegory

**Additional file 16:**

**Table S3:** Proportion of low and high recombination regions, recombination rate and percentage of QTLs located in these regions for the three traits.

“Chr” indicates the chromosome. Physical and genetic size columns indicated the size of each chromosome in bp and cM, respectively. Average recombination rate (“RecRate”) and proportion of the physical (“Phys”) and genetic (“Genetic) map in high recombination regions (“HighRec”, >0.5 cM / Mbp) for each chromosome are shown. Percentage of QTL in high recombination regions were displayed for three traits (DTA PlantHT, GY)

**Additional file 17 (.xlsx):**

**Table S4:** Description of inbred lines. Variety and accession along with the breeders, seeds providers and genetic groups obtained using ADMIXTURE for K=4 (Stiff Stalk, Iodent, Lancaster, Other).

**Additional file 18 (.docx):**

**Table S5:** Narrow sense heritability (h^2^) and variance components (V_g_, genetic variance; V_e_, residual variance). The heritability and variance components were estimated for all traits (grain yield, male flowering time and plant height) using the R package Heritability [1].

**Additional file 19(.docx):**

**Notes S1**: Differences between arrays and GBS discrovery / pipelines and.

**Additional file 20:**

**Table S6:** Number of SNPs called, after QC filtering (MAF>1%) and useful for GWAS (MAF≥5%). Note that GBS1 have SNPs with 100% missing genotypes which were removed while GBS2 used external haplotype library which allow to impute loci with 100% missing data. It conducted to a smaller number of SNPs for GBS1 than GBS2.

**Additional file 21:**

**Notes S2**: GWAS statistical models and effects of confounding factors on GWAS.

