## Additional file 1: Fig S1 for "Genotyping-by-sequencing and SNP-arrays are complementary for detecting quantitative trait loci by tagging different haplotypes in association studies"

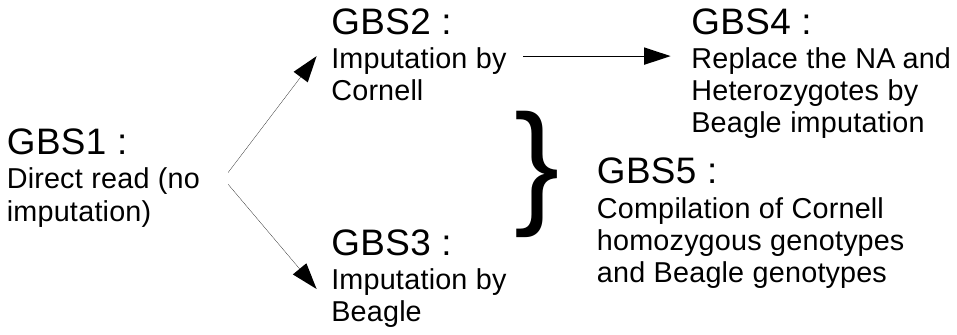

**Figure S1:** Different approaches used to impute missing data of the GBS. We considered the direct reads from GBS (**GBS_1_**) and four approaches for imputation (GBS_2_ to GBS_5_). **GBS_2_** approach consisted in one imputation step from the direct read by Cornell University, using *TASSEL* software, but missing data was still present. **GBS_3_** approach consisted in a genotype imputation of the whole missing data of the direct read by *Beagle v3*. In **GBS_4_,** genotype imputation by Beagle was performed on Cornell imputed data after replacing the heterozygous genotypes into missing data. **GBS_5_**, consisted in homozygous genotypes of GBS_2_ completed by values imputed in GBS_3_.
