## Additional file 2: Fig S2 for "Genotyping-by-sequencing and SNP-arrays are complementary for detecting quantitative trait loci by tagging different haplotypes in association studies"

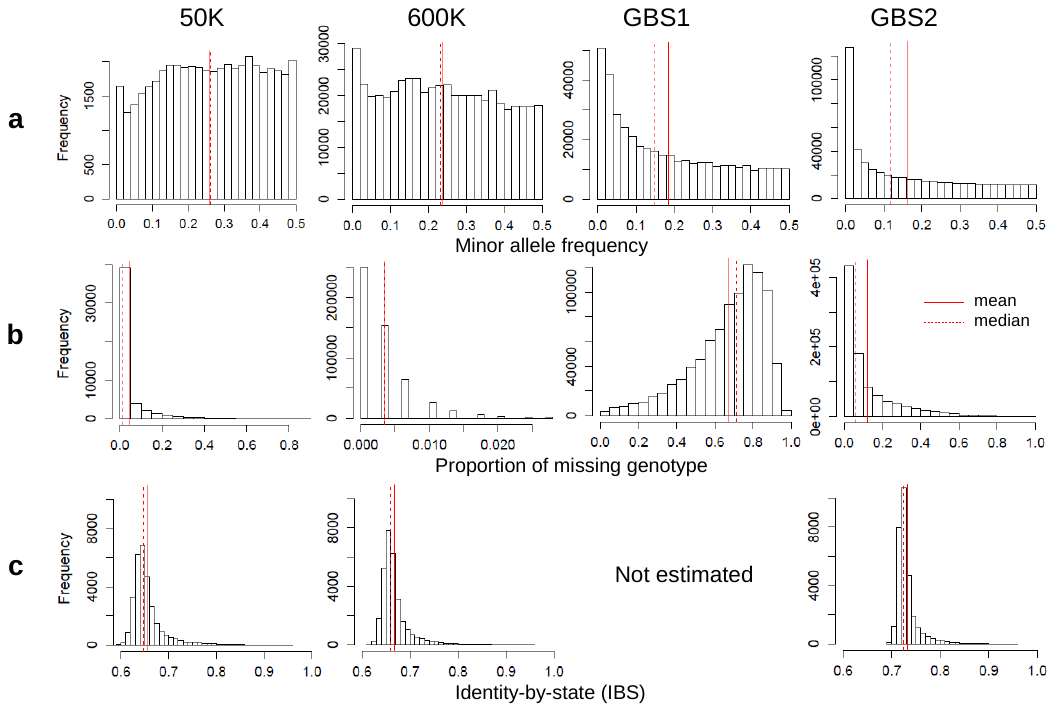

**Figure S2**: Comparison of genotyping data between 50K and 600K arrays, and GBS. (a**)** Distribution of minor allele frequency per SNP before filtering (monomorphic SNPs removed). (b**)** Distribution of SNP missing data proportion for the 50K array, 600K array, GBS direct reads (GBS_1_) and GBS after imputation by Cornell Institute (GBS_2_, note that the scale of the x-axes is different). (**c**) Relatedness distribution (Identity-By-State, IBS) after QC filtering with MAF≥1% (IBS using GBS_1_ was not estimated because of the low calling rate).
