## Additional file 3: Fig S3 for "Genotyping-by-sequencing and SNP-arrays are complementary for detecting quantitative trait loci by tagging different haplotypes in association studies"

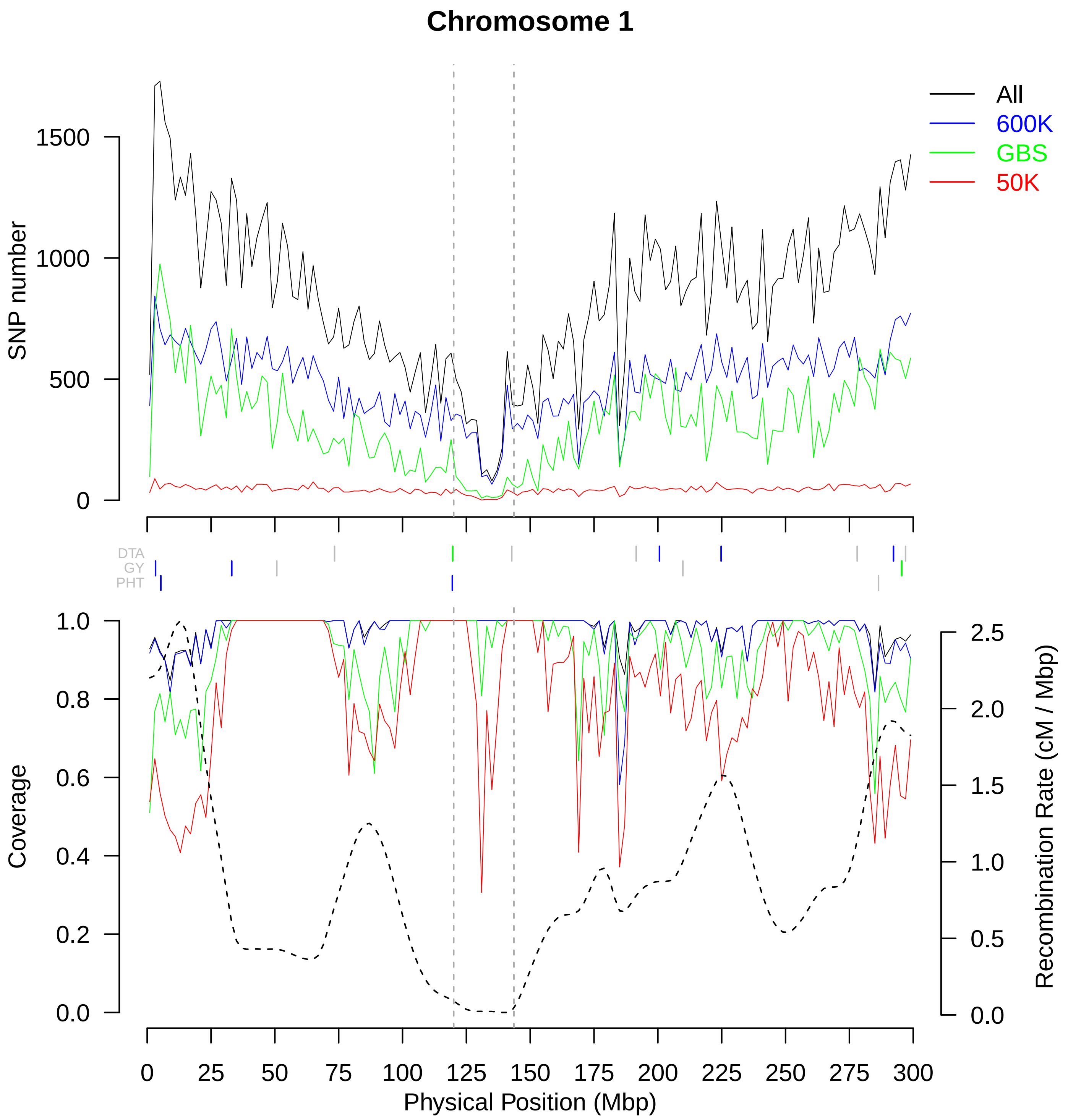

Chromosome 1
All
600K
GBS
50K
SNP number
1500
1000
500
0
DTA
GY
PHT
Coverage
1.0
0.8
0.6
0.4
0.2
0.0
Recombination Rate (cM / Mbp)
2.5
2.0
1.5
1.0
0.5
0.0
0 25 50 75 100 125 150 175 200 225 250 275 300
Physical Position (Mbp)

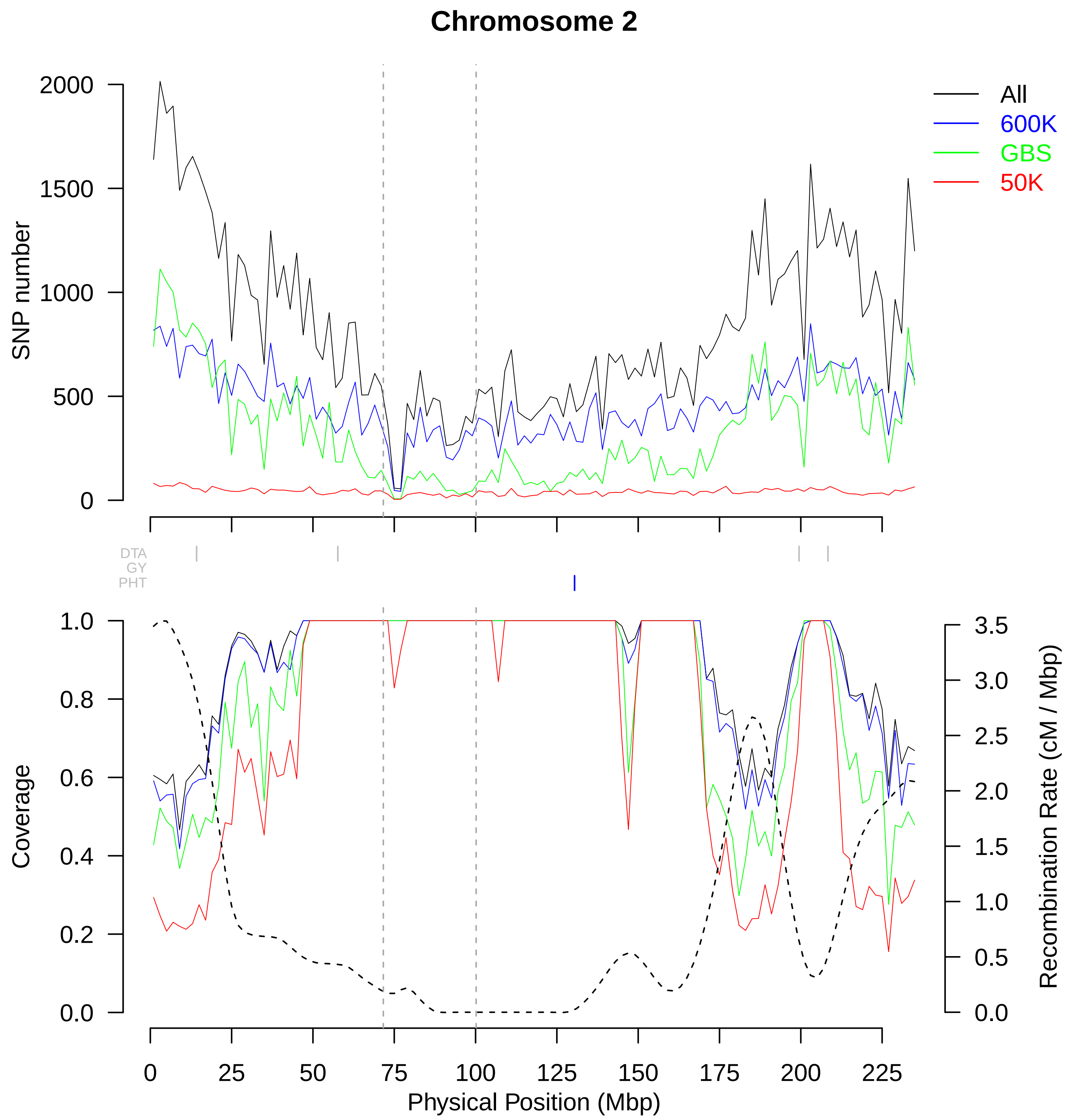
Chromosome 2
All
600K
GBS
50K
SNP number
2000
1500
1000
500
0
DTA
GY
PHT
Coverage
1.0
0.8
0.6
0.4
0.2
0.0
Recombination Rate (cM / Mbp)
3.5
3.0
2.5
2.0
1.5
1.0
0.5
0.0
0
25
50
75
100
125
150
175
200
225
Physical Position (Mbp)

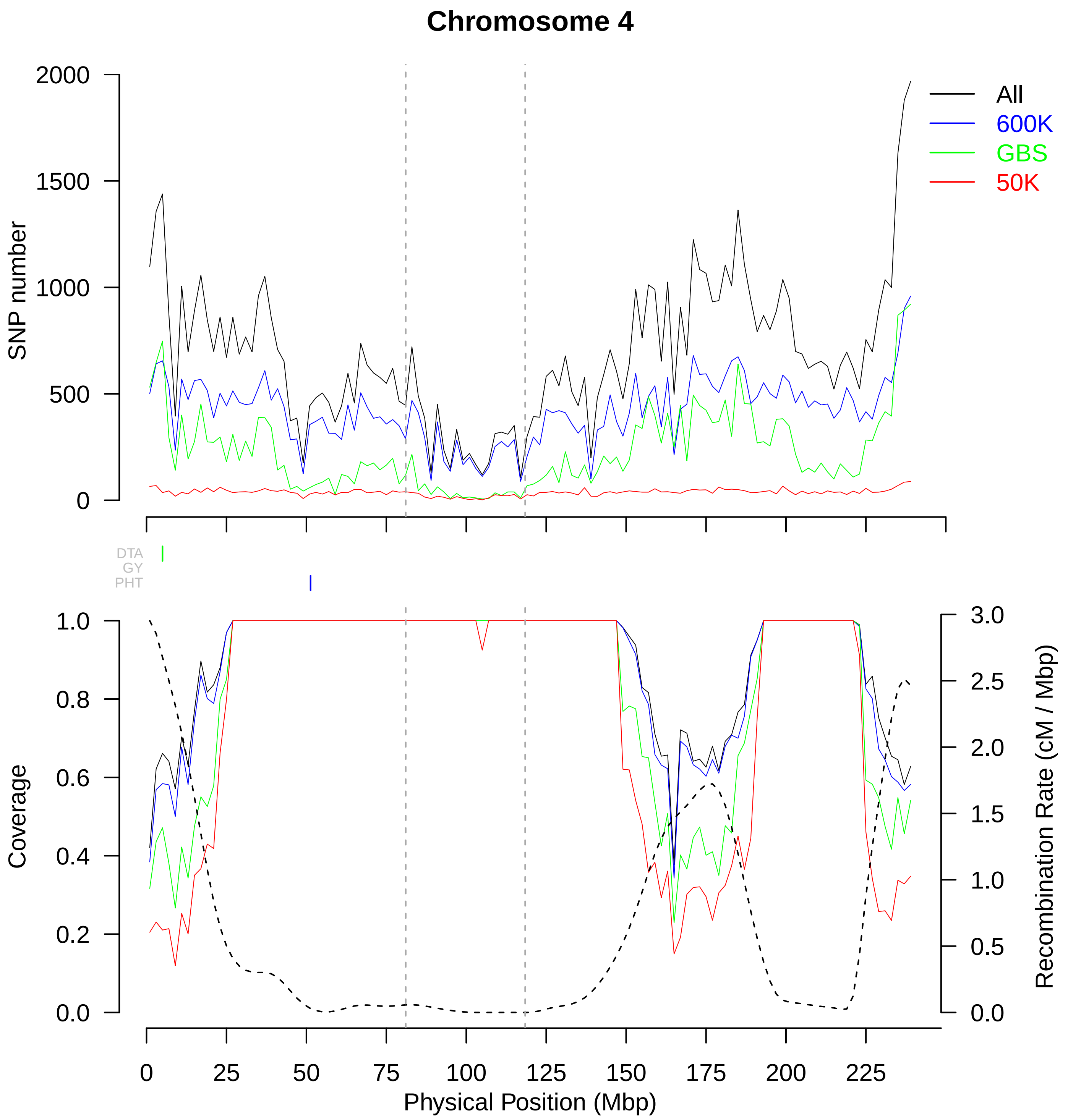

Chromosome 4
SNP number
2000
1500
1000
500
0
All
600K
GBS
50K
DTA
GY
PHT
Coverage
1.0
0.8
0.6
0.4
0.2
0.0
Recombination Rate (cM / Mbp)
3.0
2.5
2.0
1.5
1.0
0.5
0.0
0
25
50
75
100
125
150
175
200
225
Physical Position (Mbp)

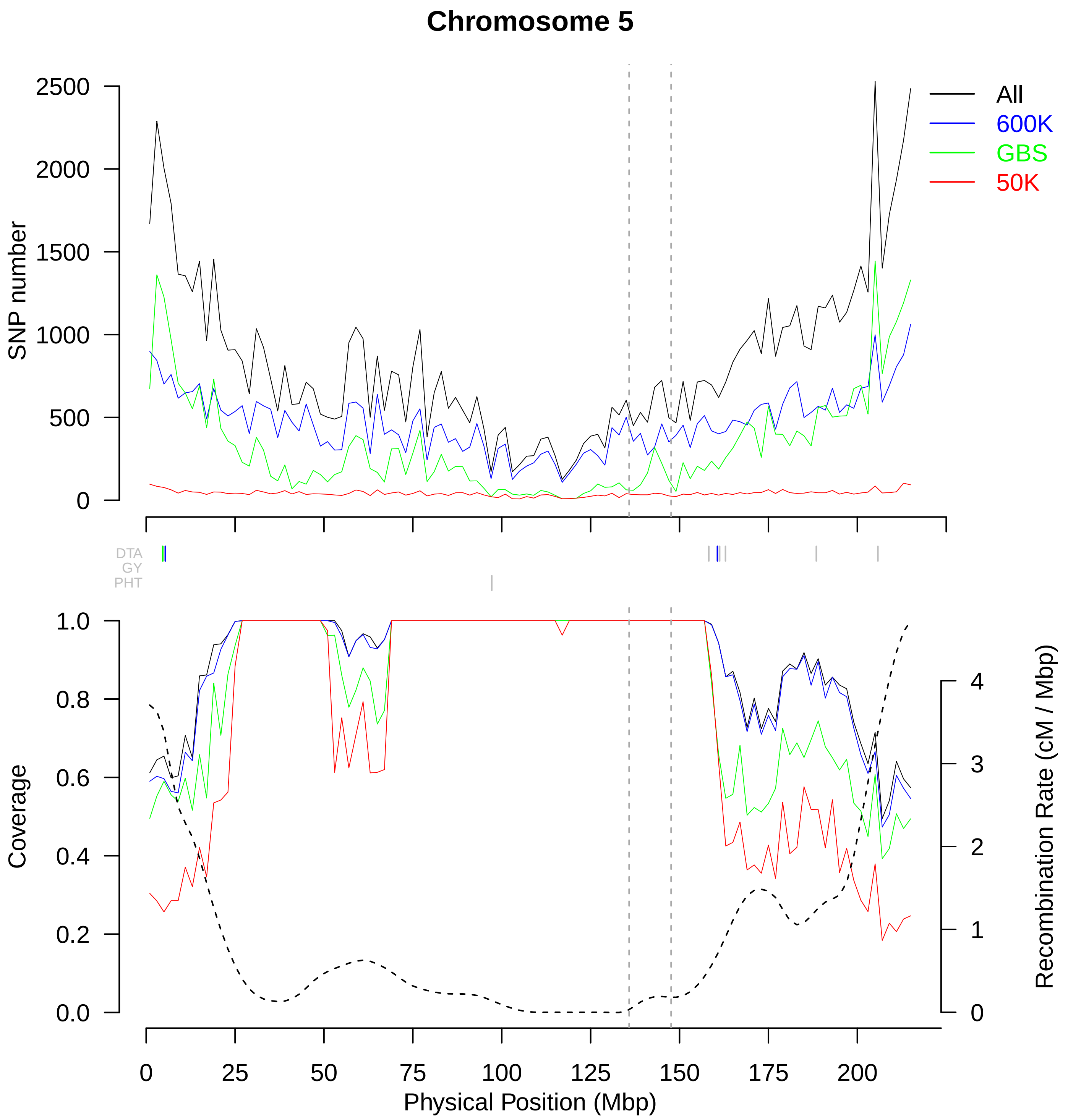

Chromosome 5
SNP number
2500
2000
1500
1000
500
0
All
600K
GBS
50K
DTA
GY
PHT
Coverage
1.0
0.8
0.6
0.4
0.2
0.0
Recombination Rate (cM / Mbp)
4
3
2
1
0
0
25
50
75
100
125
150
175
200
Physical Position (Mbp)

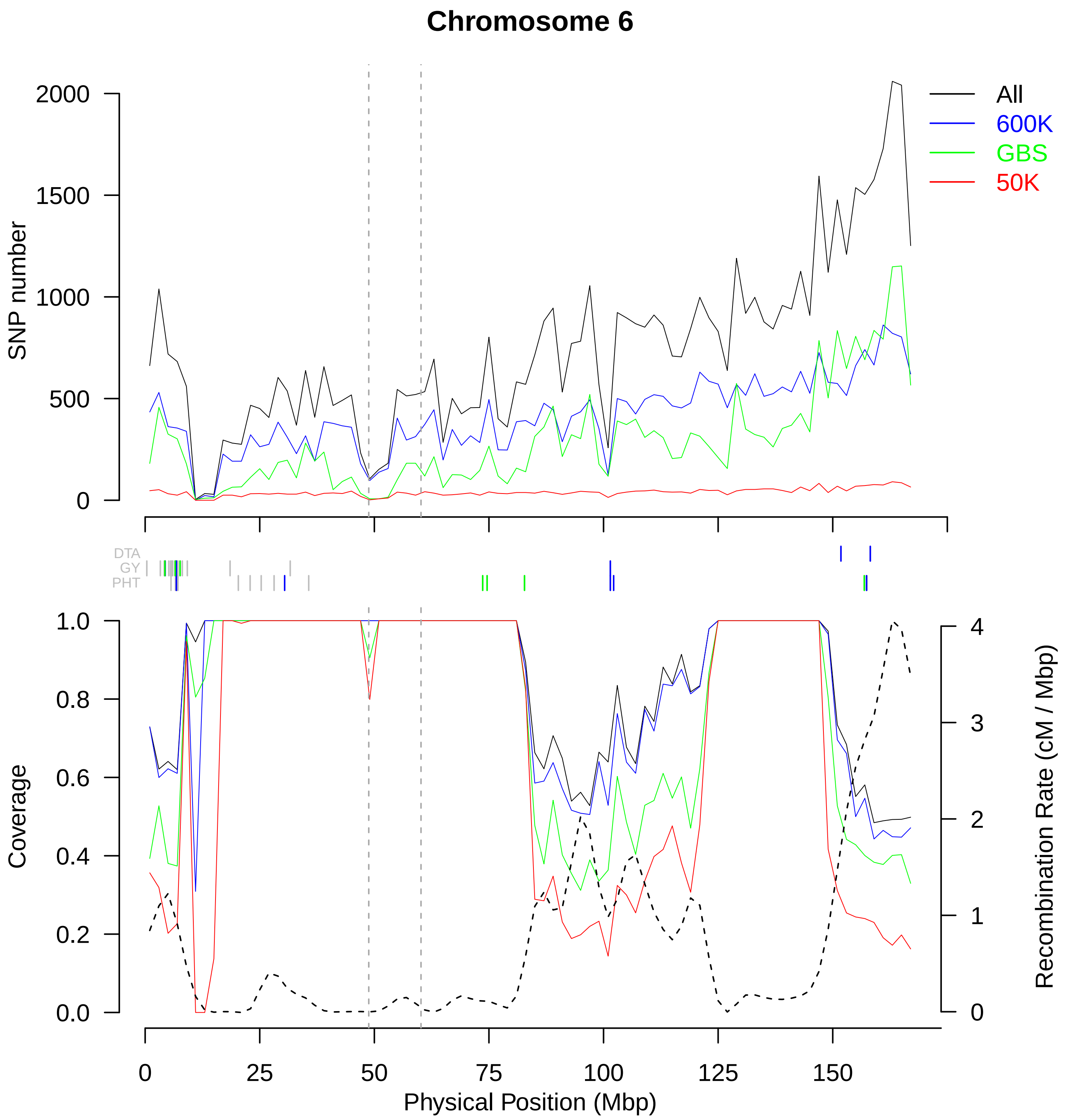
Chromosome 6
SNP number
2000
1500
1000
500
0
All
600K
GBS
50K
DTA
GY
PHT
Coverage
1.0
0.8
0.6
0.4
0.2
0.0
Recombination Rate (cM / Mbp)
4
3
2
1
0
0
25
50
75
100
125
150
Physical Position (Mbp)

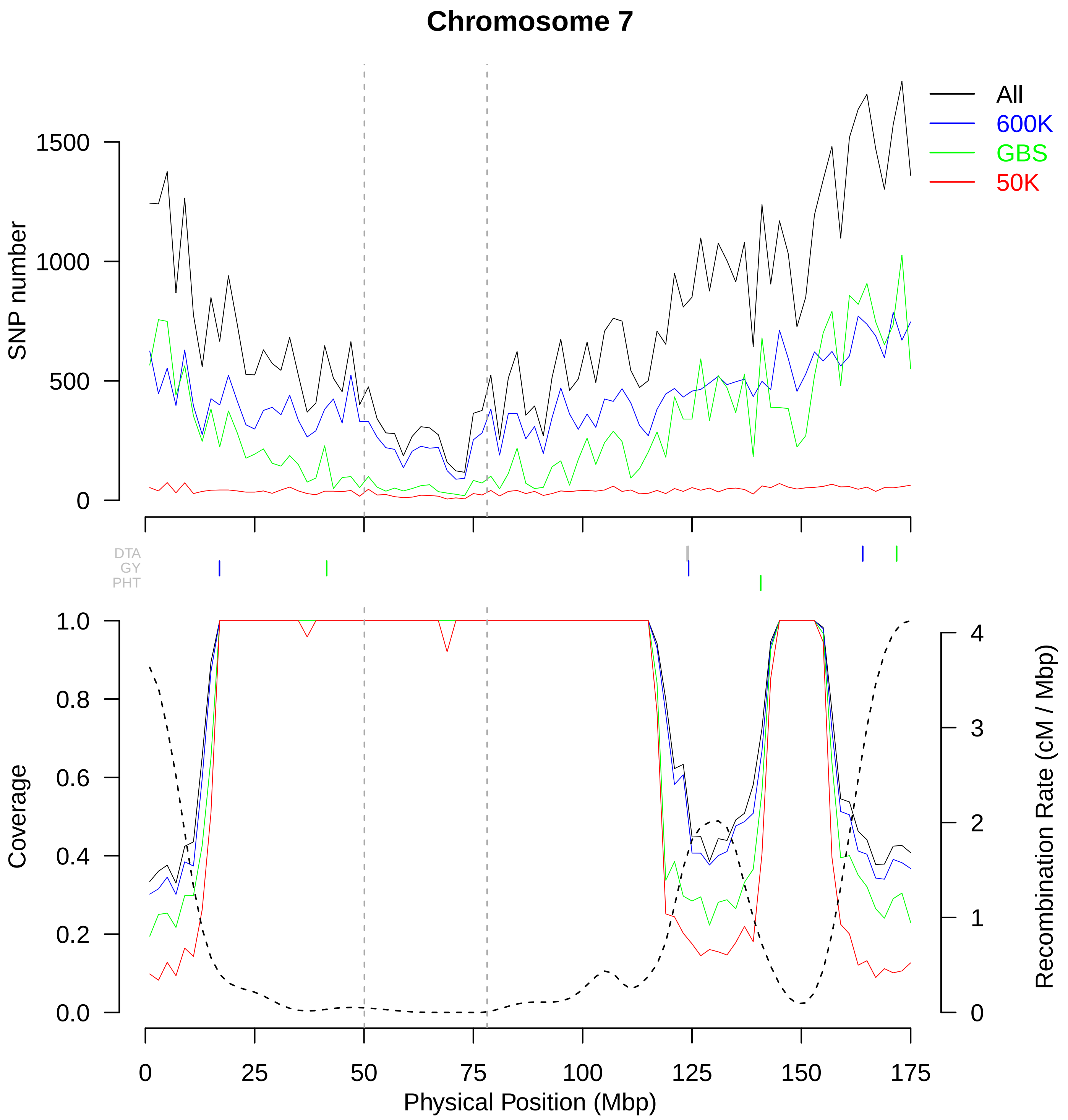

Chromosome 7
All
600K
GBS
50K
SNP number
1500
1000
500
0
DTA
GY
PHT
Coverage
1.0
0.8
0.6
0.4
0.2
0.0
Recombination Rate (cM / Mbp)
4
3
2
1
0
25
50
75
100
125
150
175
Physical Position (Mbp)

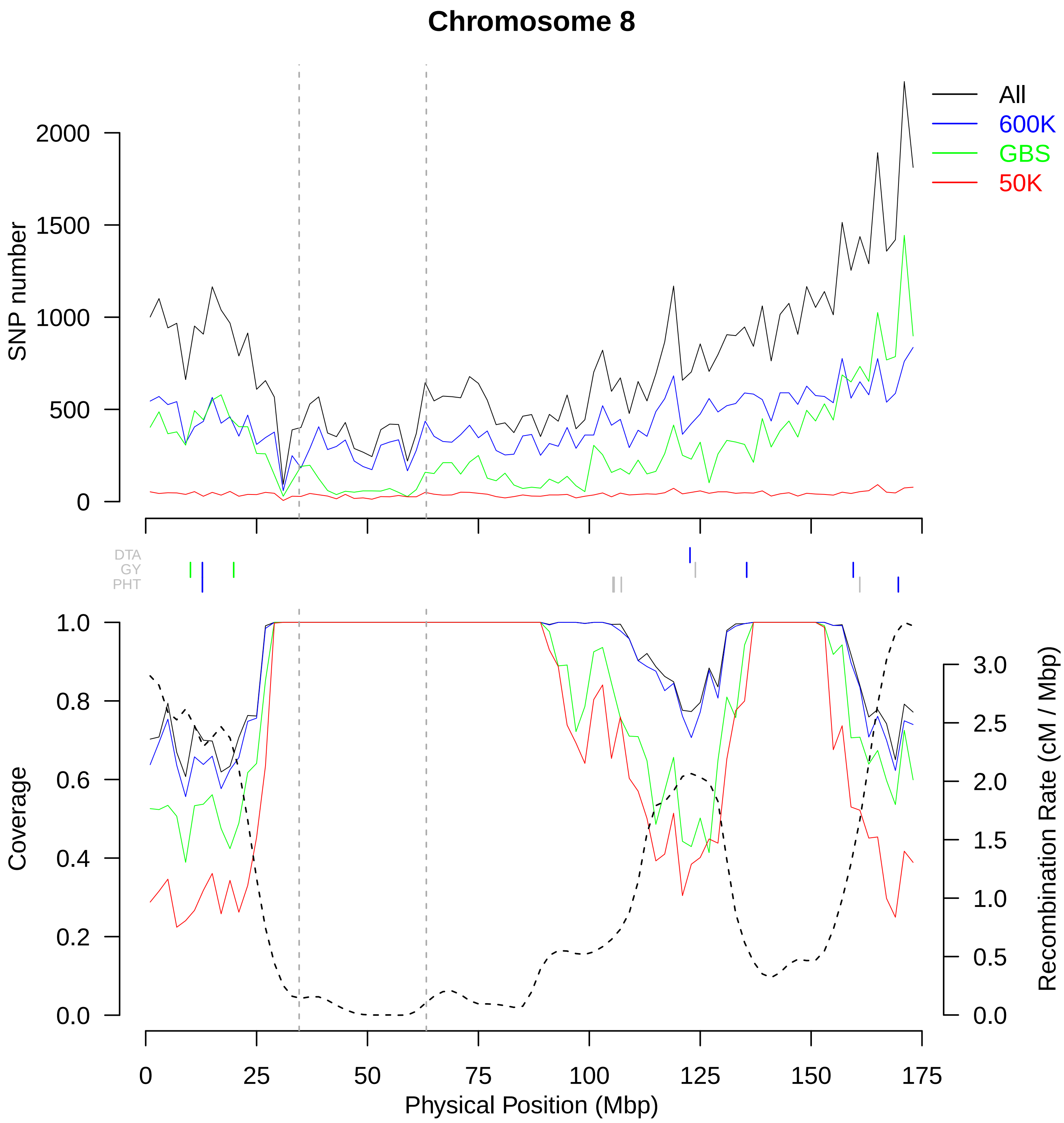
Chromosome 8
All
600K
GBS
50K
SNP number
2000
1500
1000
500
0
DTA
GY
PHT
Coverage
1.0
0.8
0.6
0.4
0.2
0.0
Recombination Rate (cM / Mbp)
3.0
2.5
2.0
1.5
1.0
0.5
0.0
0
25
50
75
100
125
150
175
Physical Position (Mbp)

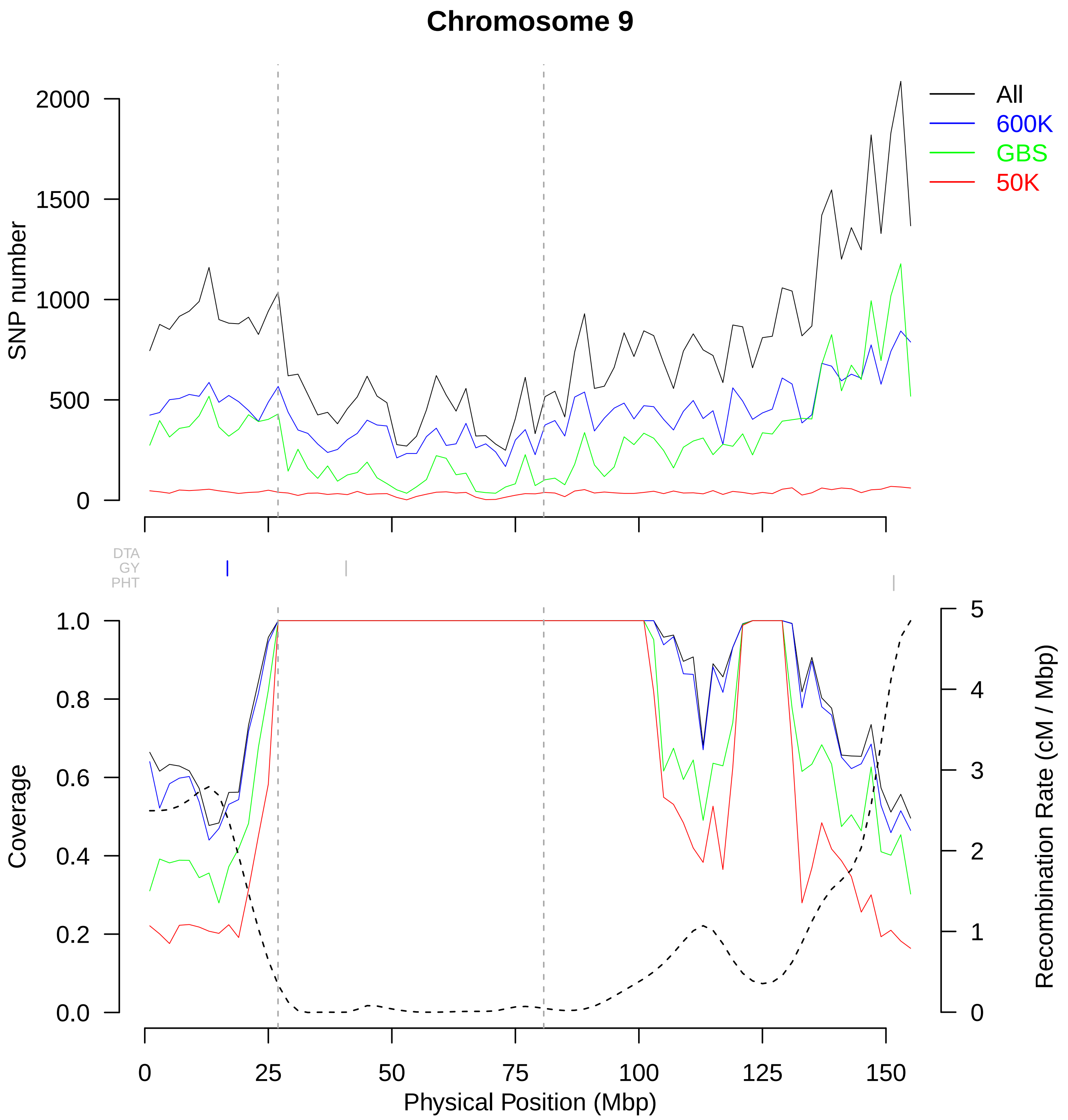

Chromosome 9
All
600K
GBS
50K
SNP number
2000
1500
1000
500
0
DTA
GY
PHT
Coverage
1.0
0.8
0.6
0.4
0.2
0.0
Recombination Rate (cM / Mbp)
5
4
3
2
1
0
0
25
50
75
100
125
150
Physical Position (Mbp)

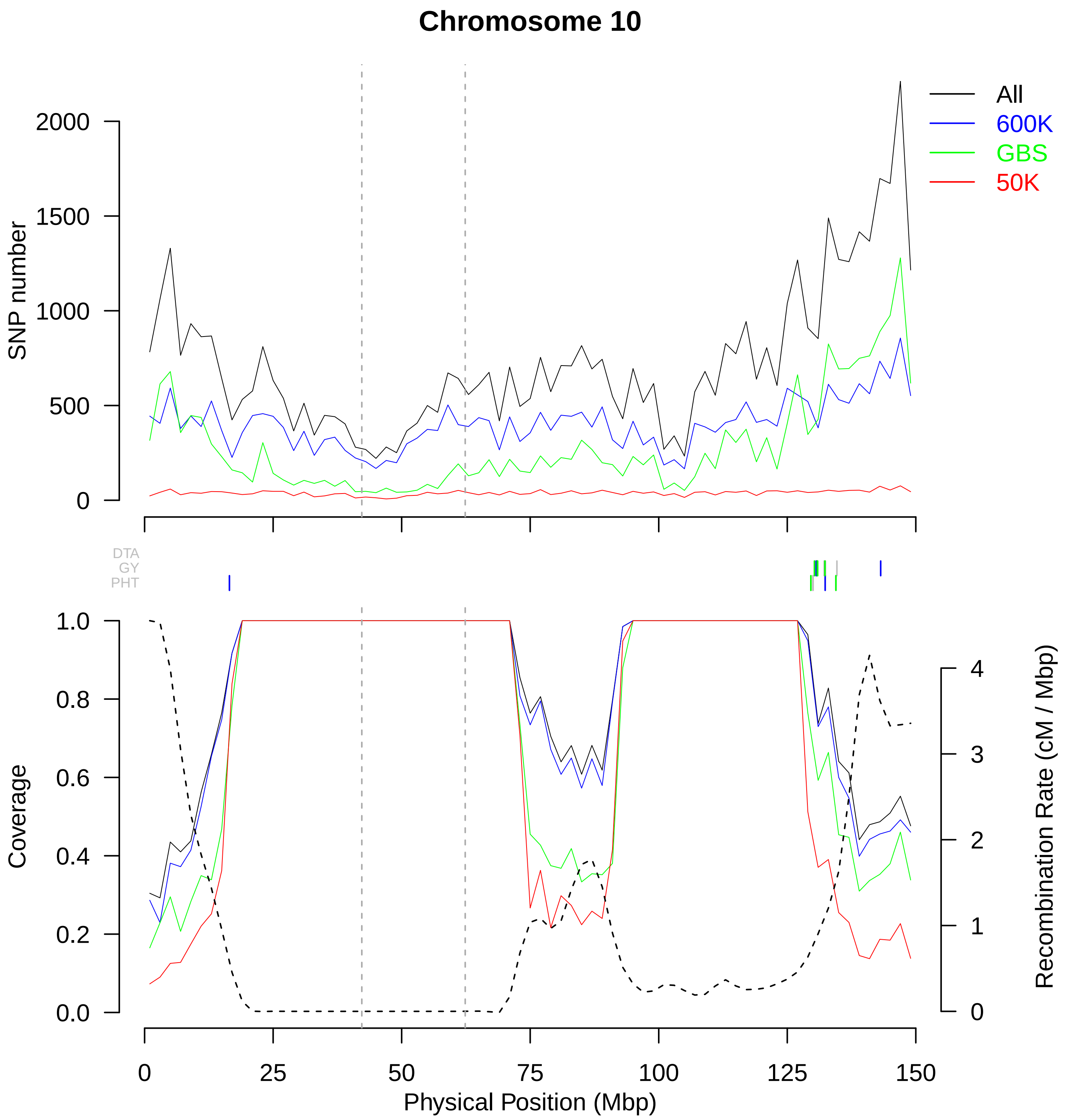

Chromosome 10
All
600K
GBS
50K
SNP number
2000
1500
1000
500
0
DTA
GY
PHT
Coverage
1.0
0.8
0.6
0.4
0.2
0.0
Recombination Rate (cM / Mbp)
4
3
2
1
0
25
50
75
100
125
150
Physical Position (Mbp)

**Additional file 3 (.pdf)**

**Figure S3:** Variation of the markers density, the recombination rate and the genome coverage in non-overlapping 2 Mbp windows along each chromosome except chromosome 3 (presented in Figure 1). Markers have MAF above 5%. Top panel shows the variation of SNP number. In the bottom panel, dotted line represents the variation of recombination rate (cM / Mbp) and solid lines the proportion of genome covered by the SNPs using the cumulated length of physical LD windows around each SNP in each 2Mbp-windows. In these two panel, green, blue, red and black lines represent variation for GBS, 600K, 50K and combined technologies, respectively. Vertical dotted gray lines indicate limits of centromeric regions. Vertical lines between the two panels indicate the position of QTLs for flowering time (DTA), grain yield (GY) and Plant Height (PHT). Green, blue, red vertical lines indicate QTLs detected only by GBS, 600K and 50K technologies, respectively. Grey vertical lines indicate QTL detected by at least two technologies. Only QTL including a marker associated with $-\log_{10}$(pval) above 6 were shown.
