## Additional file 4: Fig S4 for "Genotyping-by-sequencing and SNP-arrays are complementary for detecting quantitative trait loci by tagging different haplotypes in association studies"

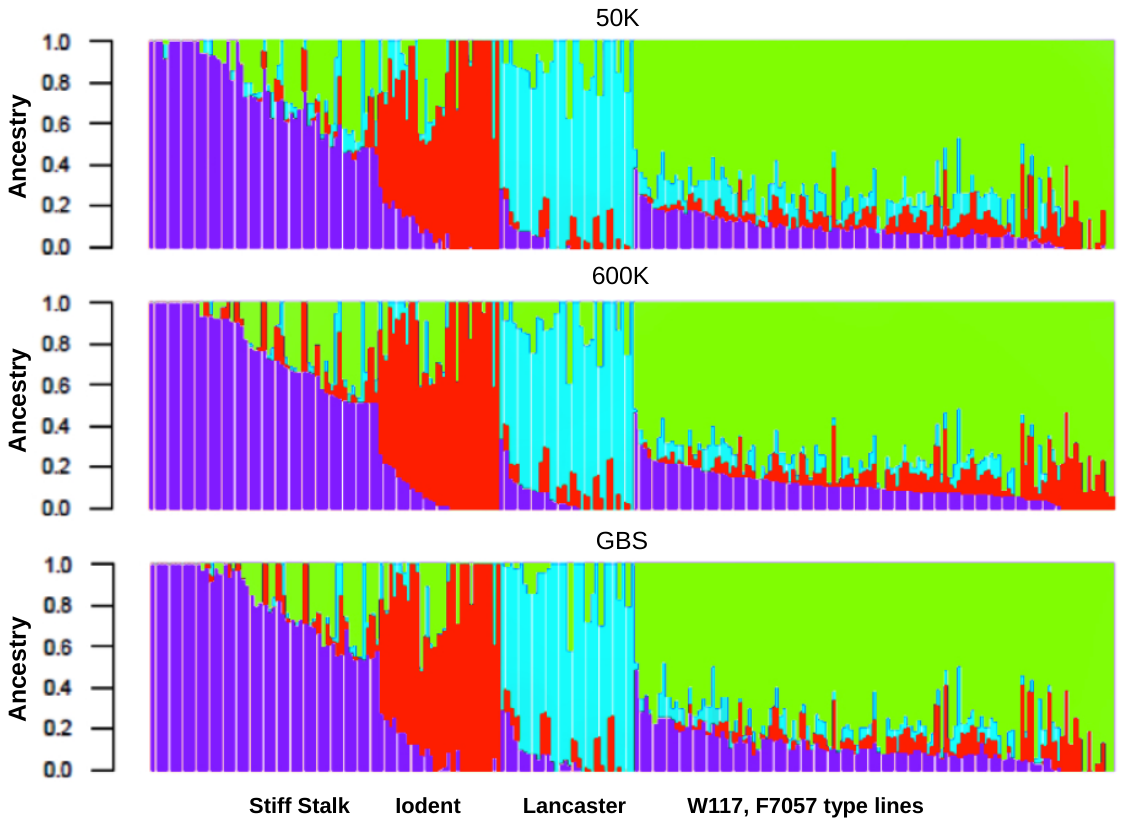

**Figure S4:** Contribution of four ancestral populations to 247 inbred lines after ADMIXTURE analysis. Markers from the 50K (top), 600K (middle) and GBS (bottom) were used. One vertical bar corresponds to one individual. Lines were ordered according to contributions observed for the 50K. From left to right, we have Stiff Stalk lines type B73 and B14a (blue), Iodent lines type PH207 (red), Lancaster lines type Mo17 and Oh43 (turquoise), a group of lines assembling W117, F7057 type lines (green).
