## Additional file 5: Fig S5 for "Genotyping-by-sequencing and SNP-arrays are complementary for detecting quantitative trait loci by tagging different haplotypes in association studies"

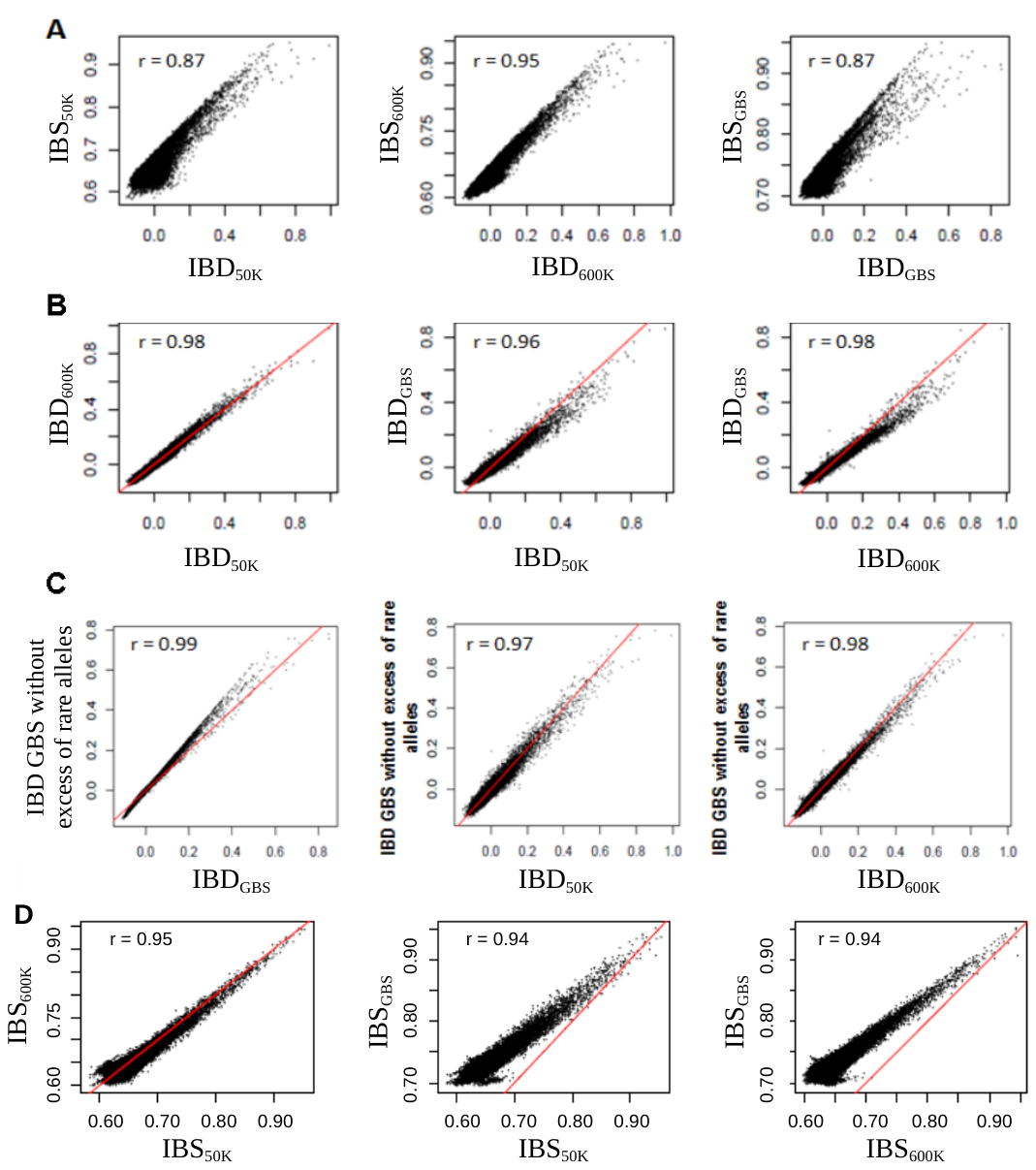

**Figure S5:** Correlation between kinship matrix estimated from different technologies. Correlation (*r*) between the IBS and IBD (*K_Freq*) for each technology (A). Correlation of IBD (B) and IBS (D) between the three technologies (after imputation). (C) Correlation of IBD between the three technologies after removing the excess of rare alleles in the GBS to have the same distribution of MAF as in the 50K and the 600K. The red line is the bisector.
