## Additional file 6: Fig S6 for "Genotyping-by-sequencing and SNP-arrays are complementary for detecting quantitative trait loci by tagging different haplotypes in association studies"

**Figure S6:** Heatmap of genome-wide linkage disequilibrium (LD) between all markers within and between chromosomes using PANZEA SNPs from the 50K. All SNPs were ordered according to their position on the genome. “Unpl” after chromosome 10 refers to unplaced SNPs, in an arbitrary order. Dots represented LD between two loci and were colored according to their strength. Classical LD measurement r^2^ between loci were represented within triangle below the diagonal. Linkage disequilibrium corrected for structure (*r^2^S*, A), relatedness (*r^2^K*, B) or both (*r^2^KS*, C) were represented within triangle above the diagonal.
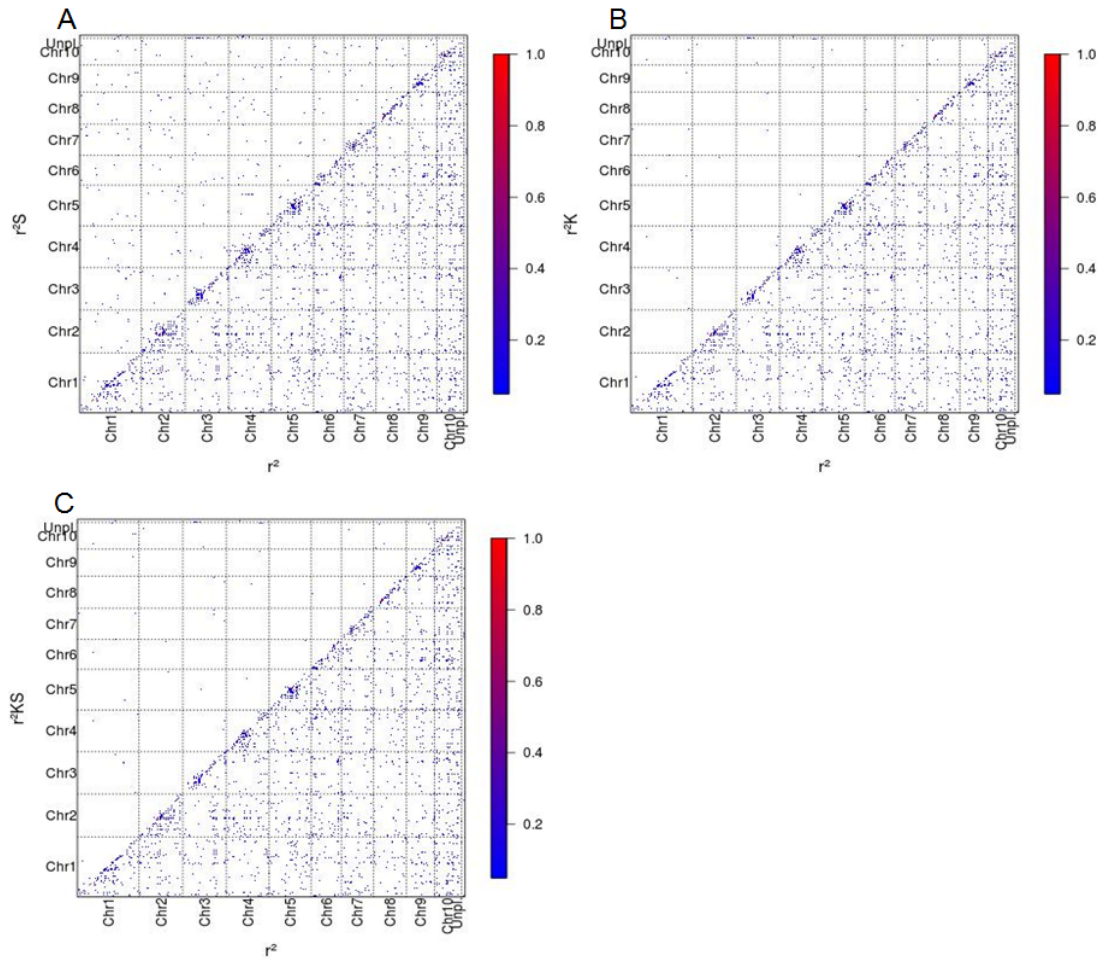
