## Additional file 7: Fig S7 for "Genotyping-by-sequencing and SNP-arrays are complementary for detecting quantitative trait loci by tagging different haplotypes in association studies"

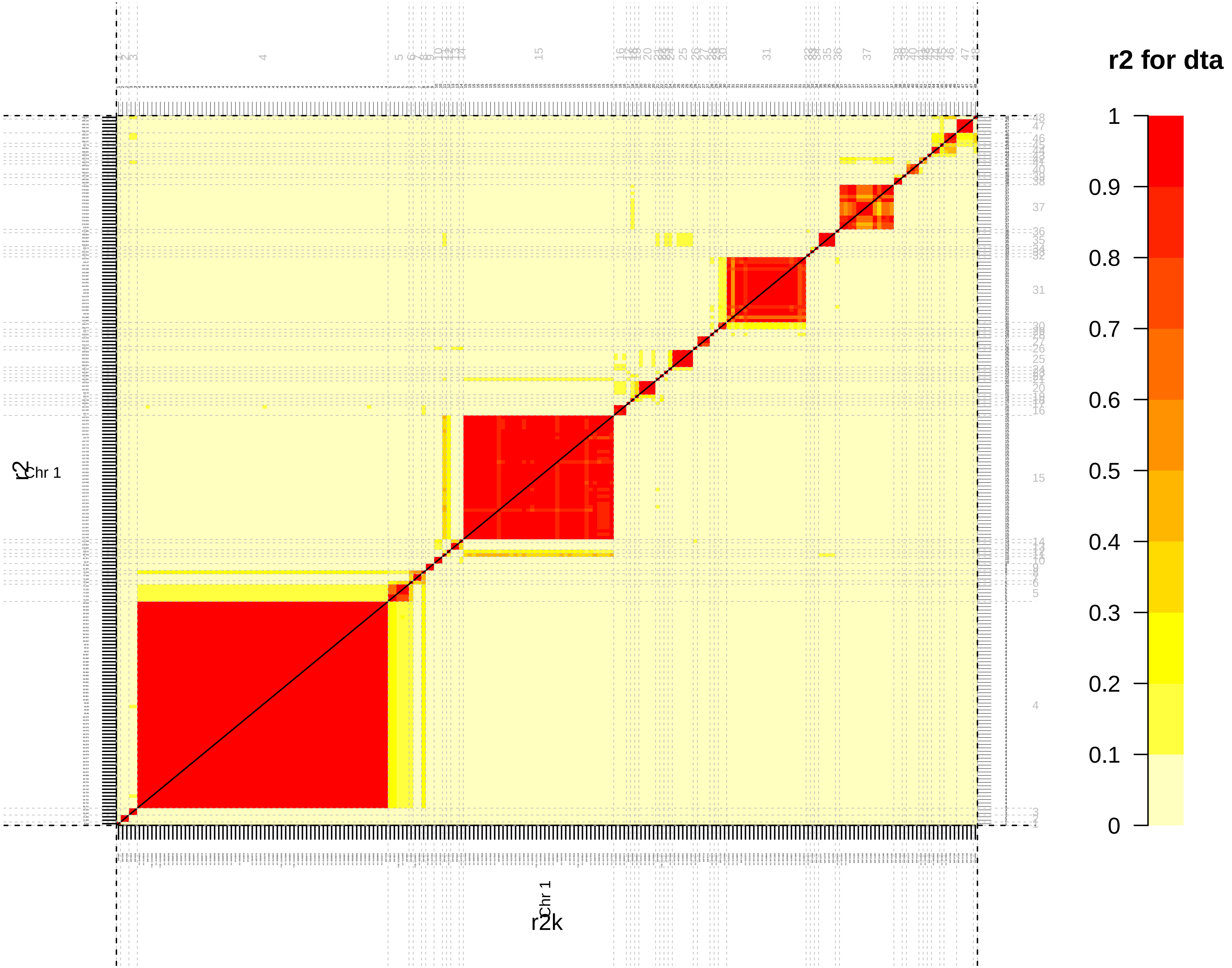
r2 for dta
1
0.9
0.8
0.7
0.6
0.5
0.4
0.3
0.2
0.1
0
Chr 1
r2k
r2

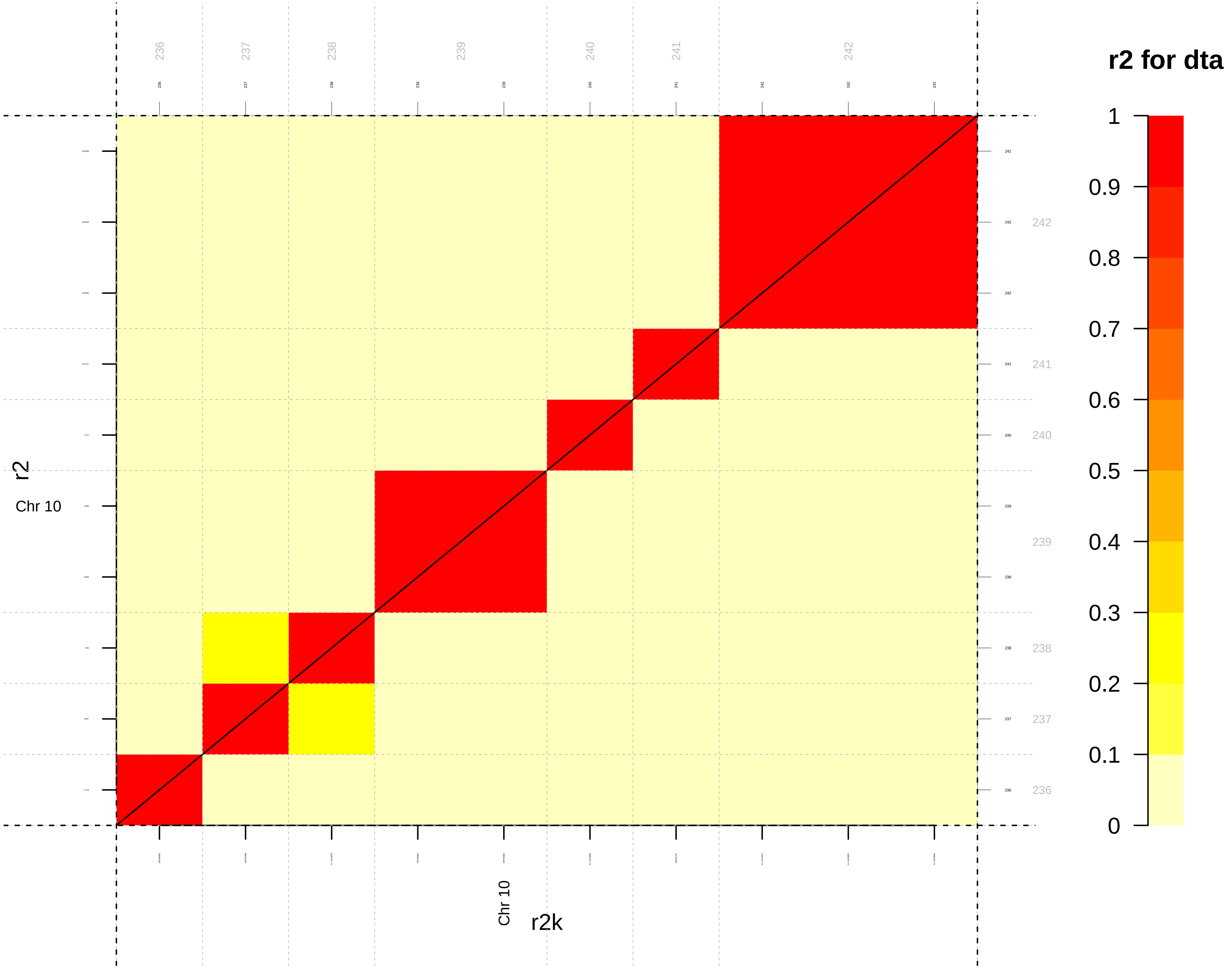

r2 for dta
236
237
238
239
240
241
242
r2
Chr 10
r2k
Chr 10
1
0.9
0.8
0.7
0.6
0.5
0.4
0.3
0.2
0.1
0

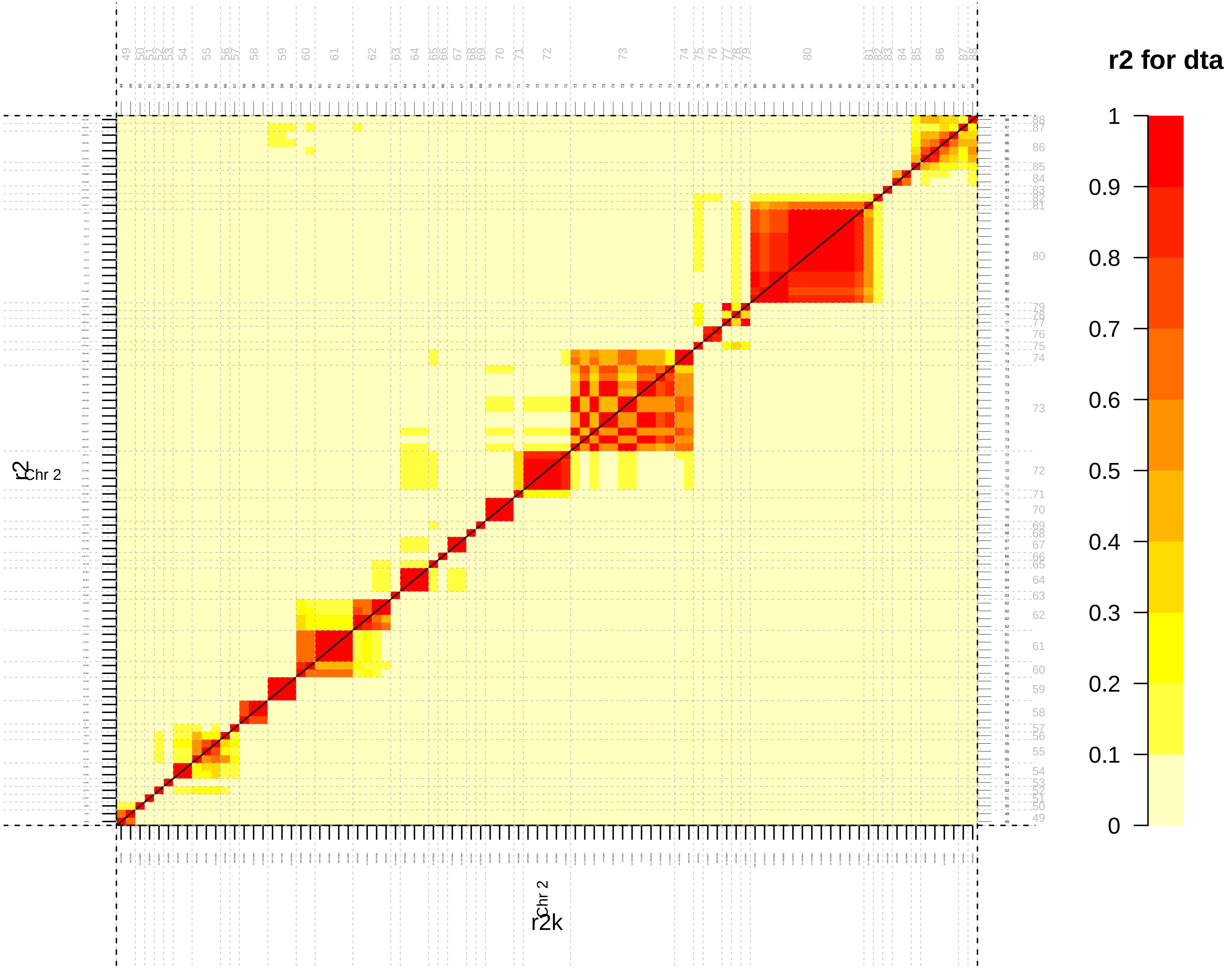
r2 for dta
1
0.9
0.8
0.7
0.6
0.5
0.4
0.3
0.2
0.1
0
r2
Chr 2
r2k
Chr 2

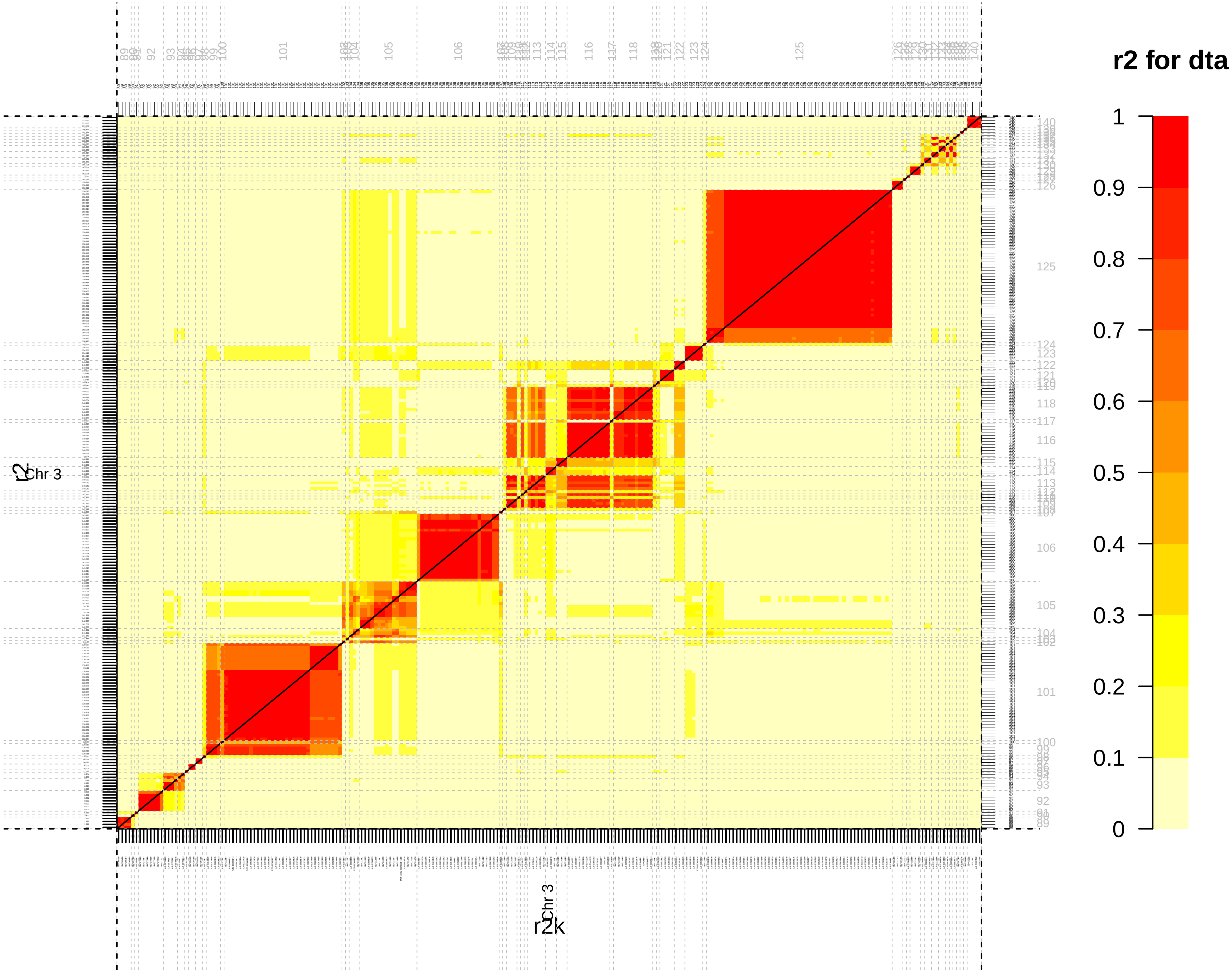
r2 for dta
1
0.9
0.8
0.7
0.6
0.5
0.4
0.3
0.2
0.1
0
Chr 3
r2
Chr 3
r2k

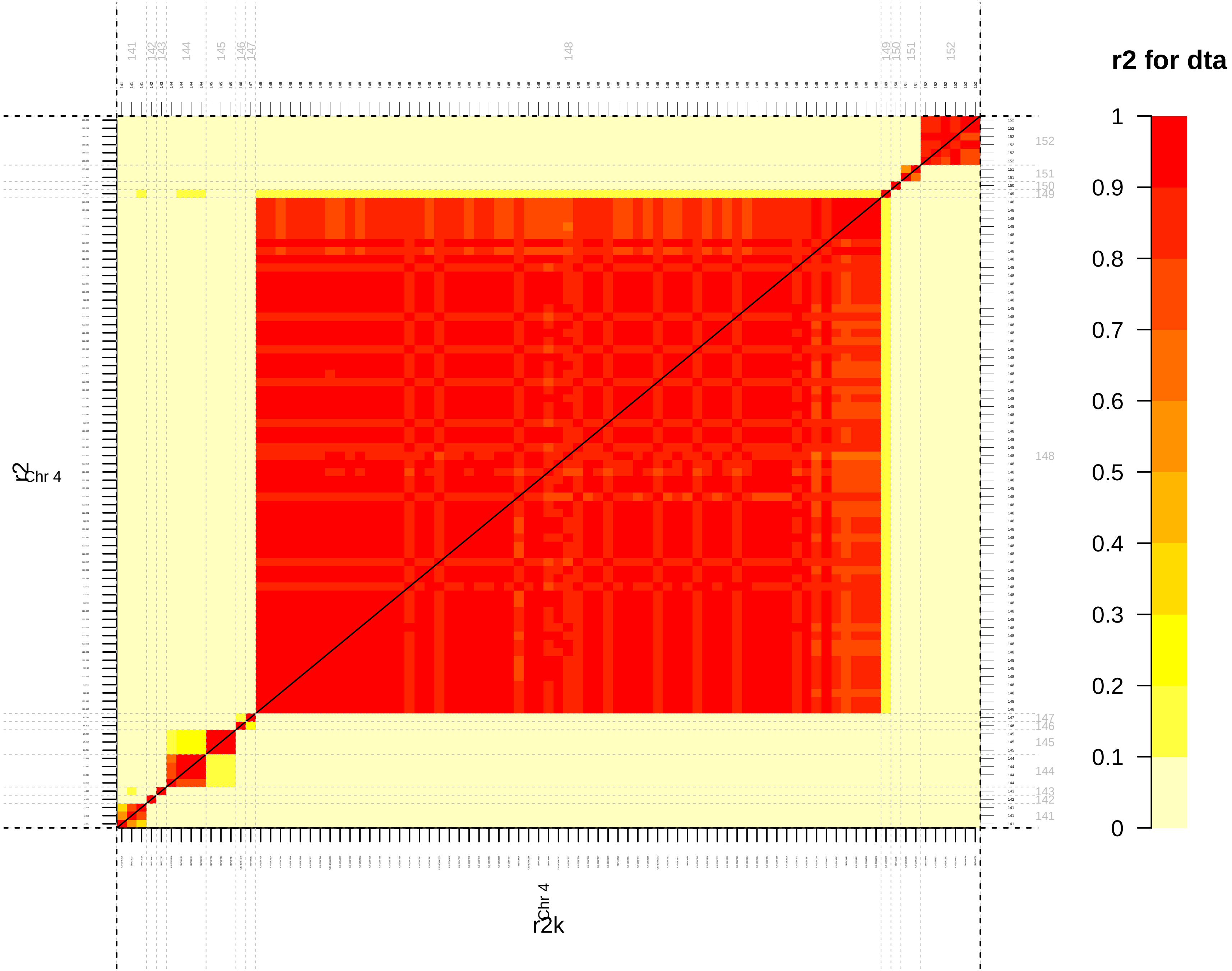

r2 for dta
1
0.9
0.8
0.7
0.6
0.5
0.4
0.3
0.2
0.1
0
Chr 4
r2
Chr 4
r2k

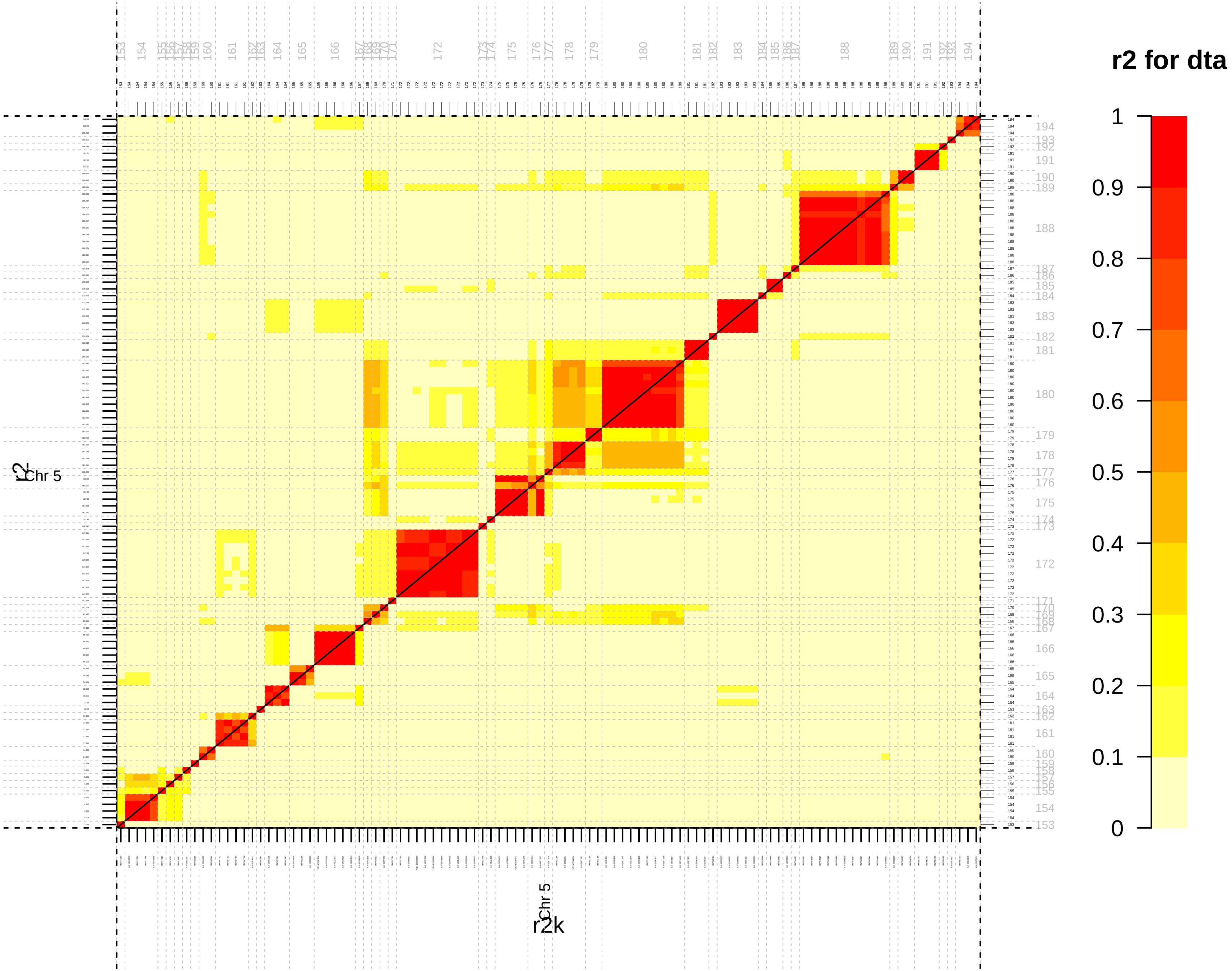

r2 for dta
1
0.9
0.8
0.7
0.6
0.5
0.4
0.3
0.2
0.1
0
Chr 5
r2
Chr 5
r2k

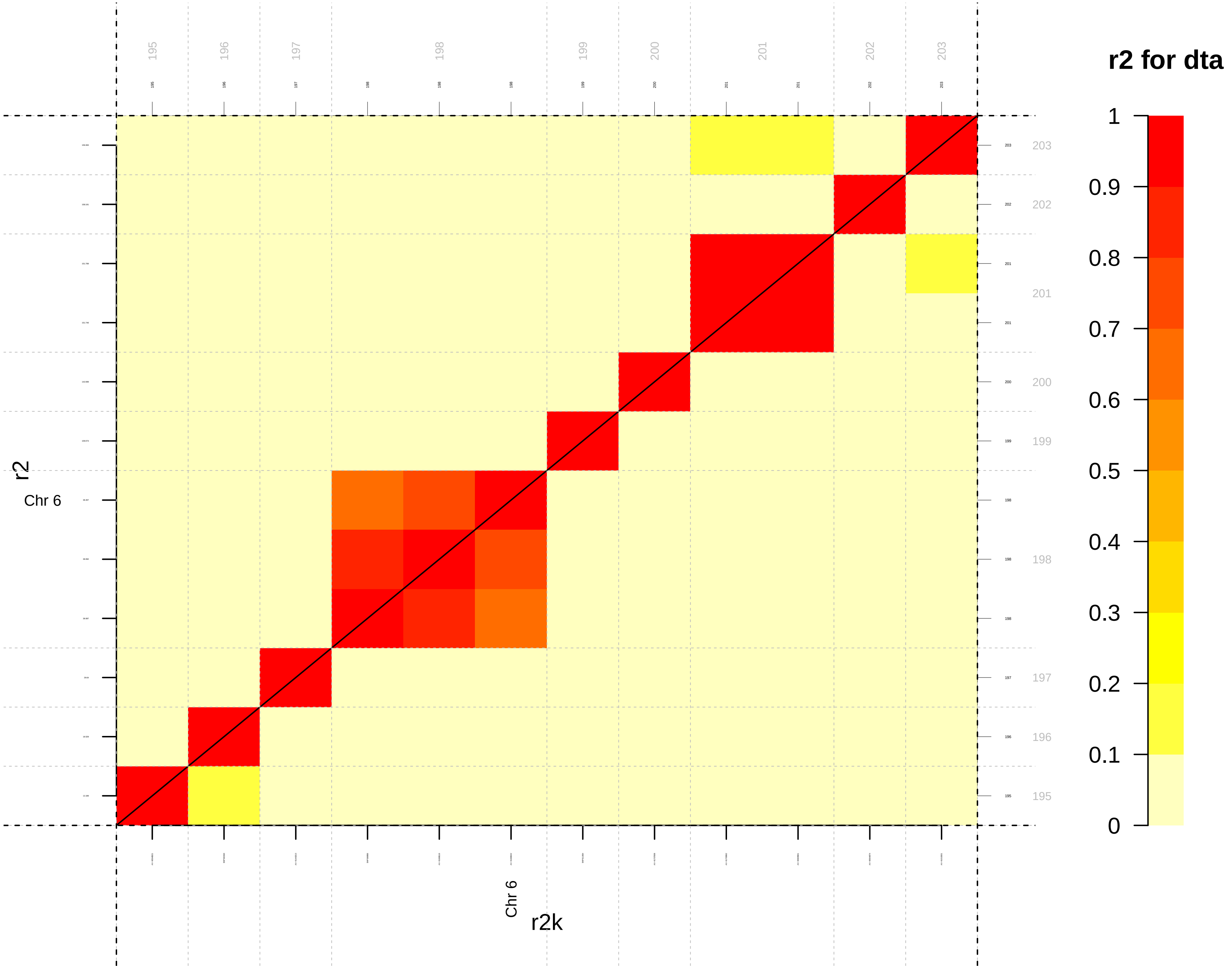

r2 for dta
1
0.9
0.8
0.7
0.6
0.5
0.4
0.3
0.2
0.1
0
195
196
197
198
199
200
201
202
203
r2
Chr 6
r2k
Chr 6

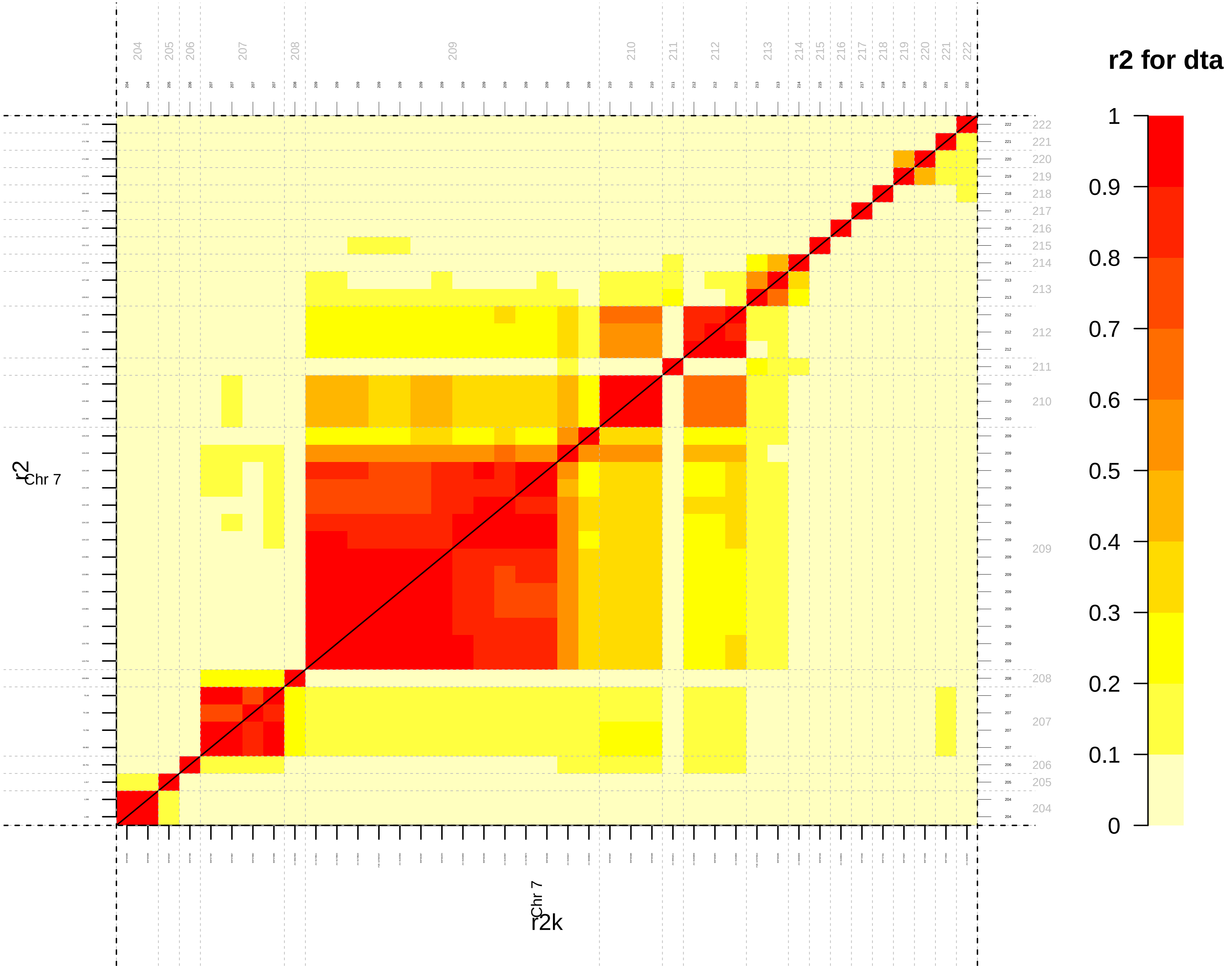
r2 for dta
1
0.9
0.8
0.7
0.6
0.5
0.4
0.3
0.2
0.1
0
Chr 7
r2
r2k

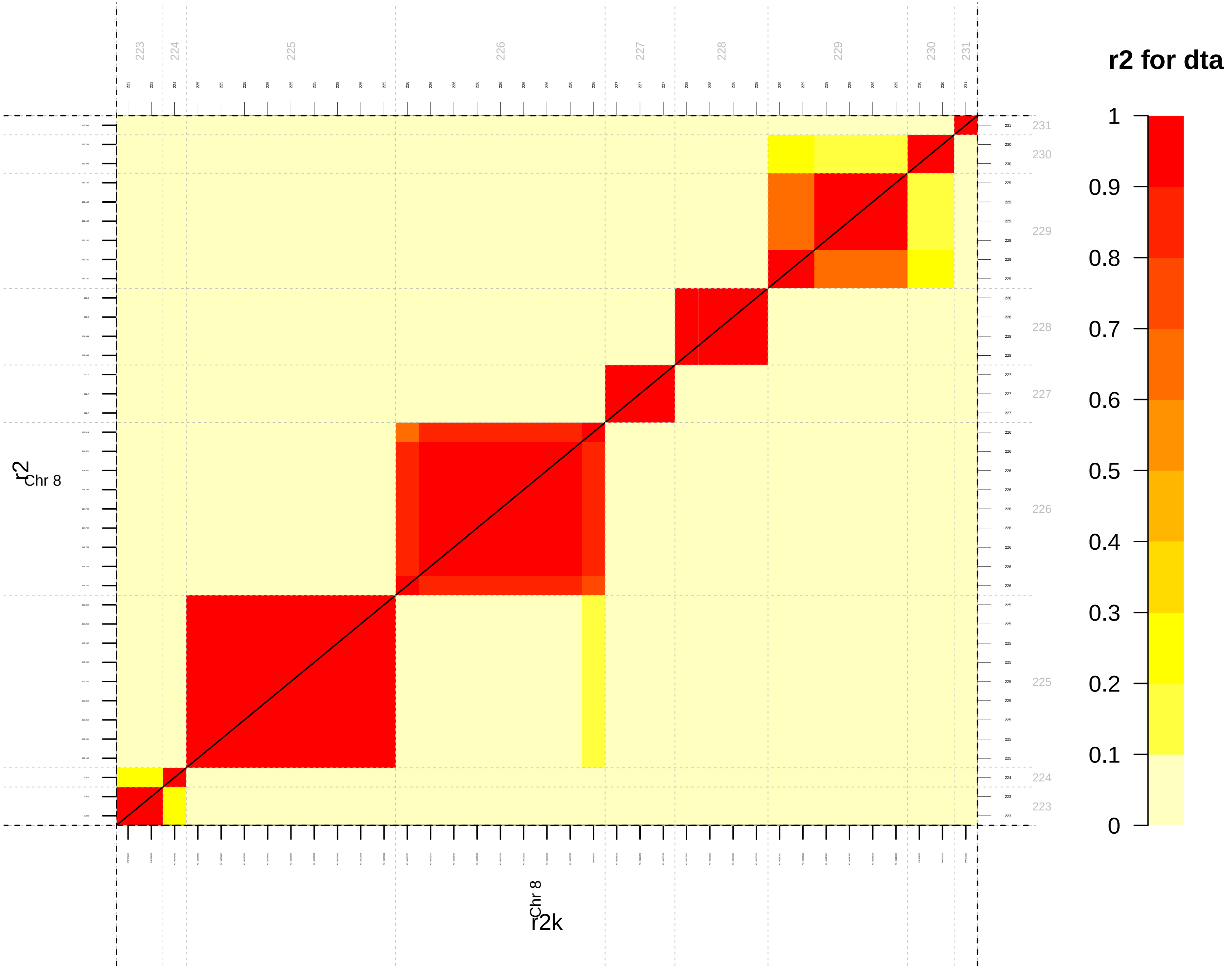
r2 for dta
1
0.9
0.8
0.7
0.6
0.5
0.4
0.3
0.2
0.1
0
223
224
225
226
227
228
229
230
231
Chr 8
r2
r2k

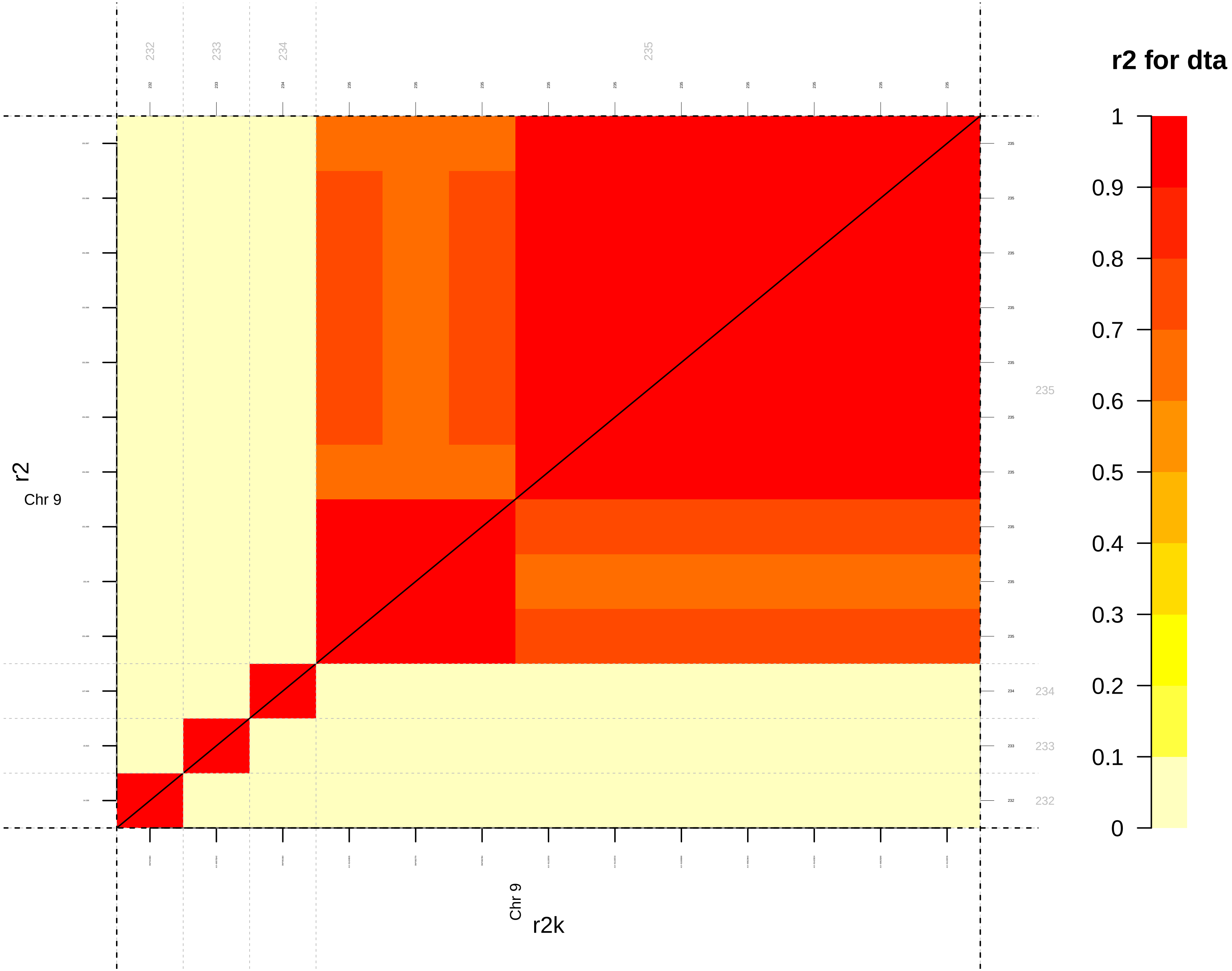
r2 for dta
1
0.9
0.8
0.7
0.6
0.5
0.4
0.3
0.2
0.1
0
232
233
234
235
r2
Chr 9
Chr 9
r2k

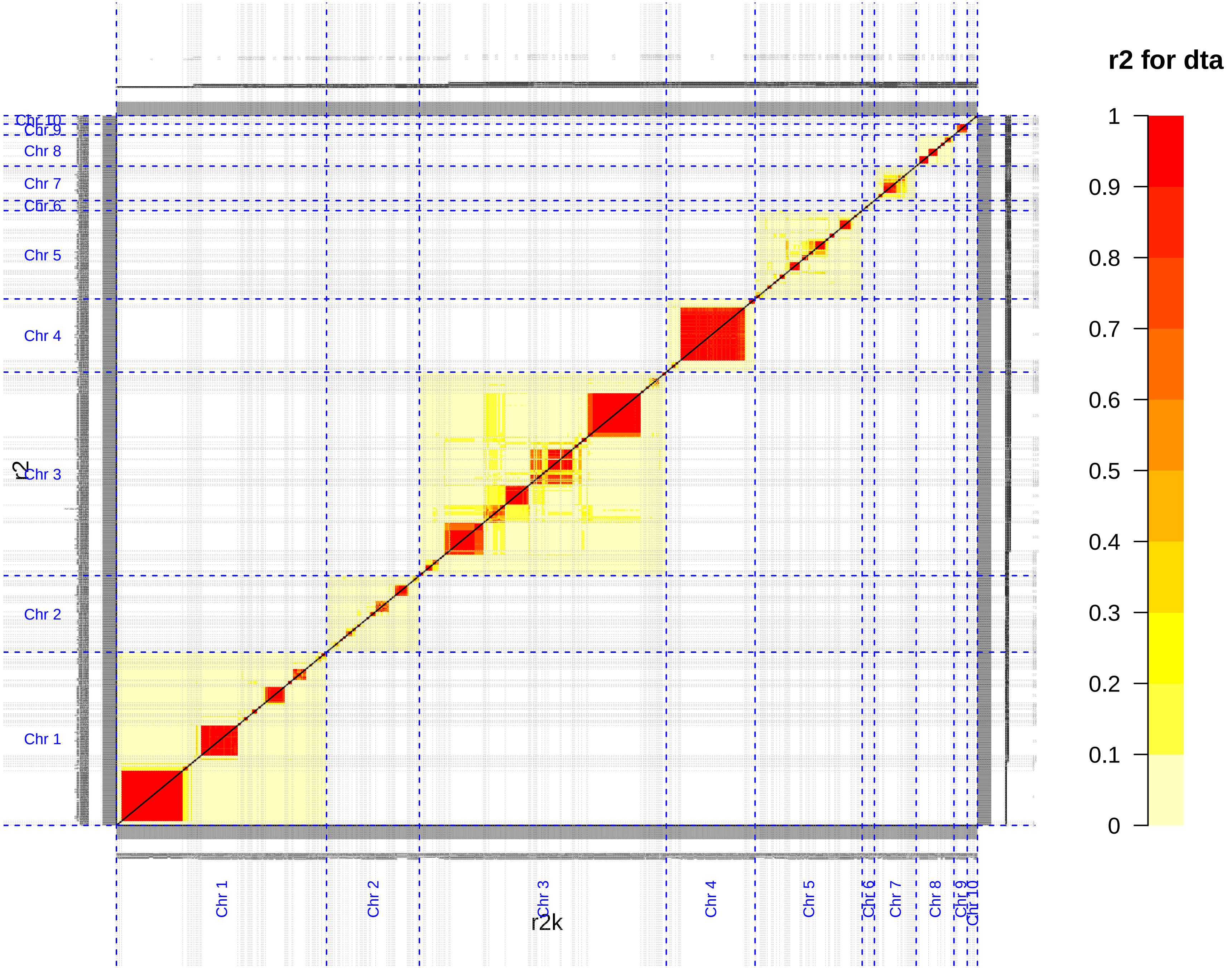
r2 for dta
1
0.9
0.8
0.7
0.6
0.5
0.4
0.3
0.2
0.1
0
Chr 10
Chr 9
Chr 8
Chr 7
Chr 6
Chr 5
Chr 4
Chr 3
Chr 2
Chr 1
r2
r2k

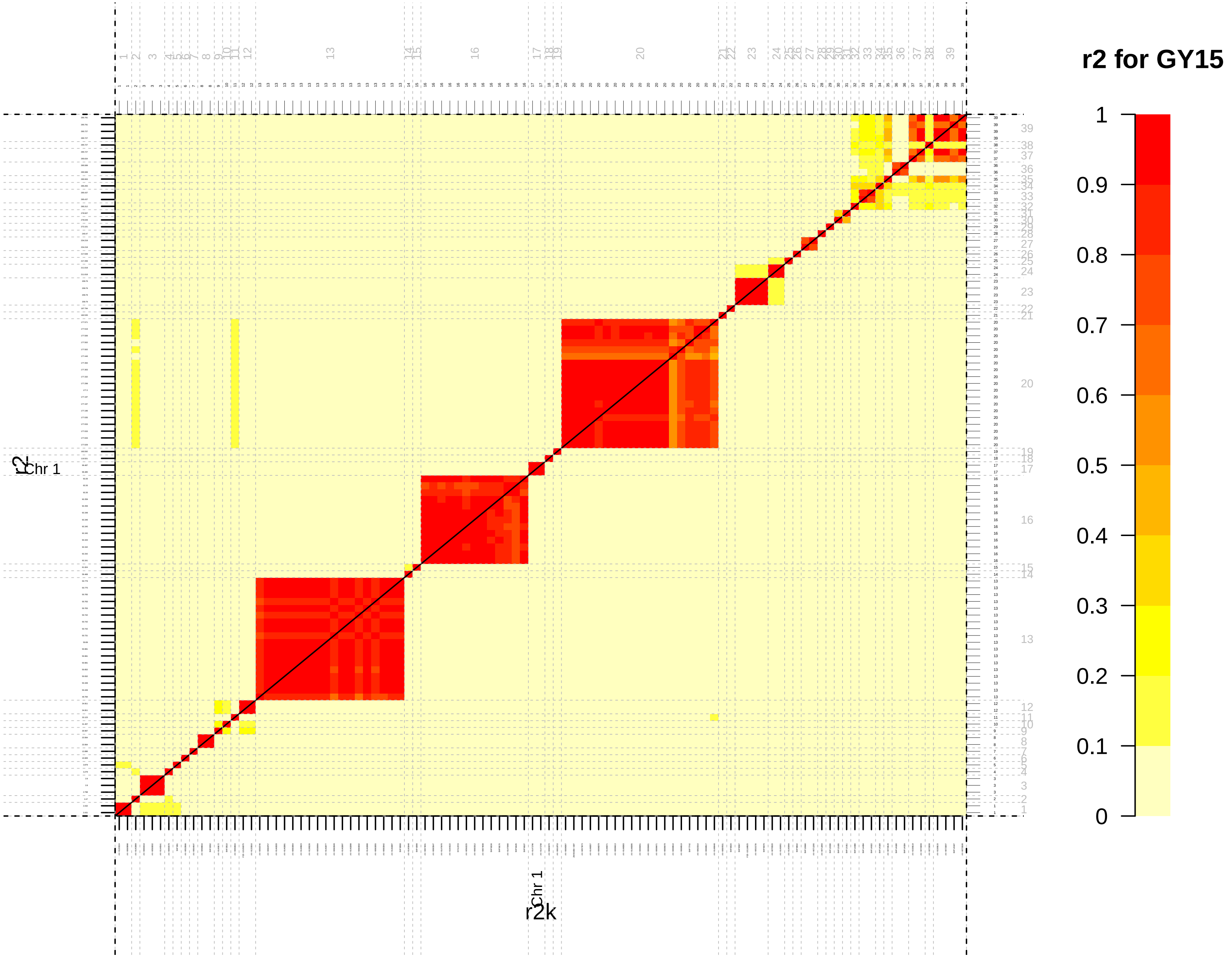
r2 for GY15
1
0.9
0.8
0.7
0.6
0.5
0.4
0.3
0.2
0.1
0
Chr 1
r2
r2k

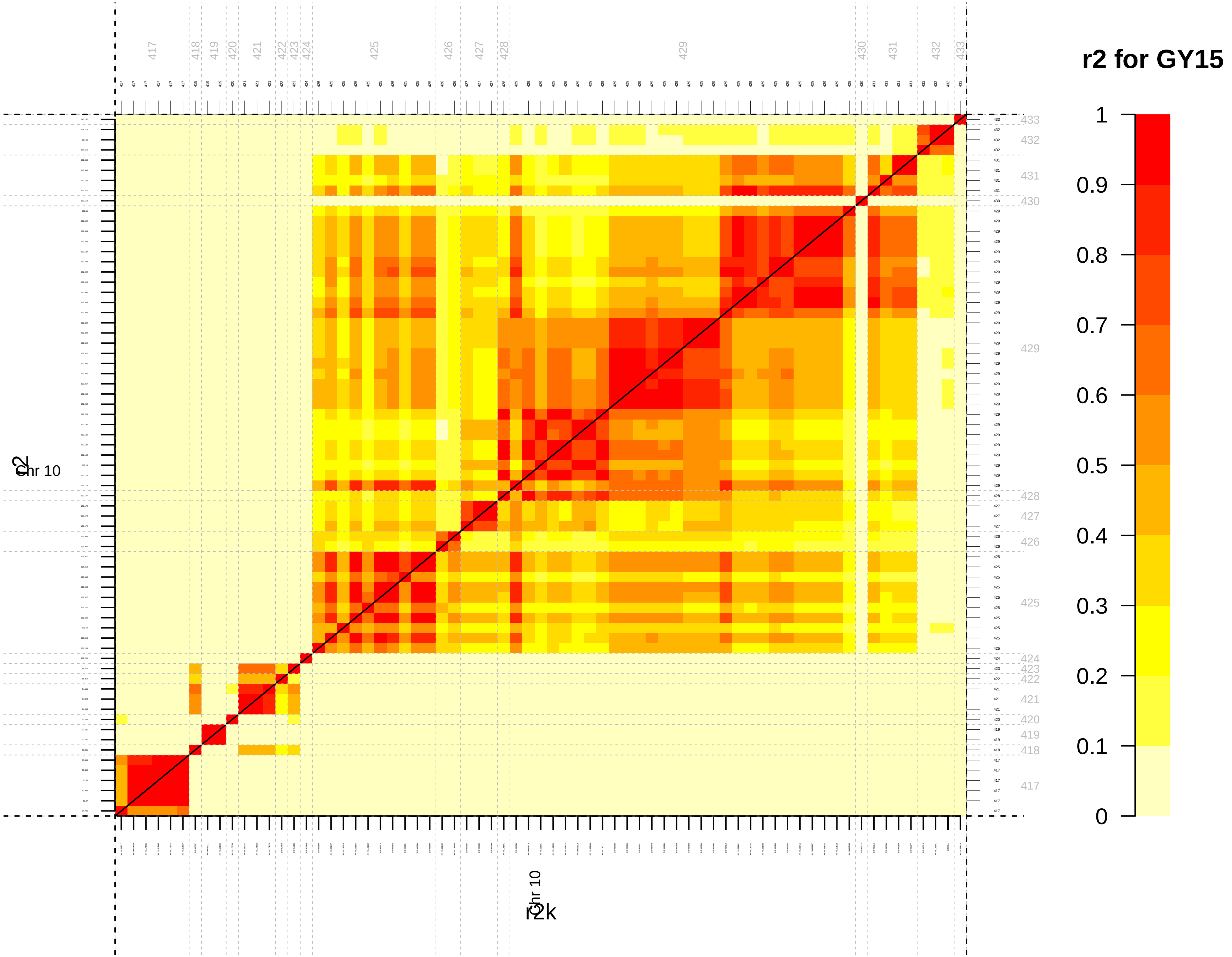
r2 for GY15
1
0.9
0.8
0.7
0.6
0.5
0.4
0.3
0.2
0.1
0
Chr 10
r2
r2k

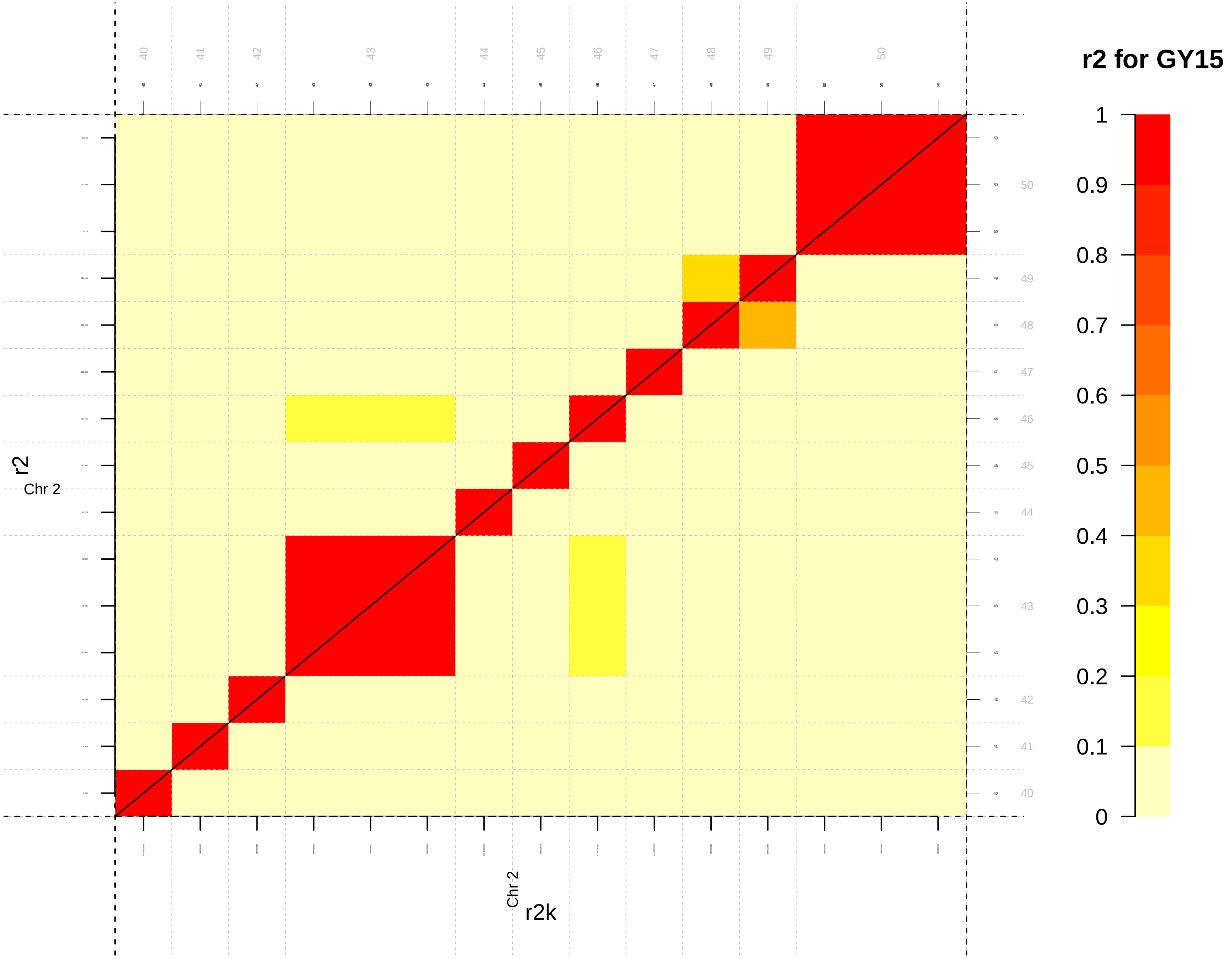
r2 for GY15
1
0.9
0.8
0.7
0.6
0.5
0.4
0.3
0.2
0.1
0
r2
Chr 2
r2k

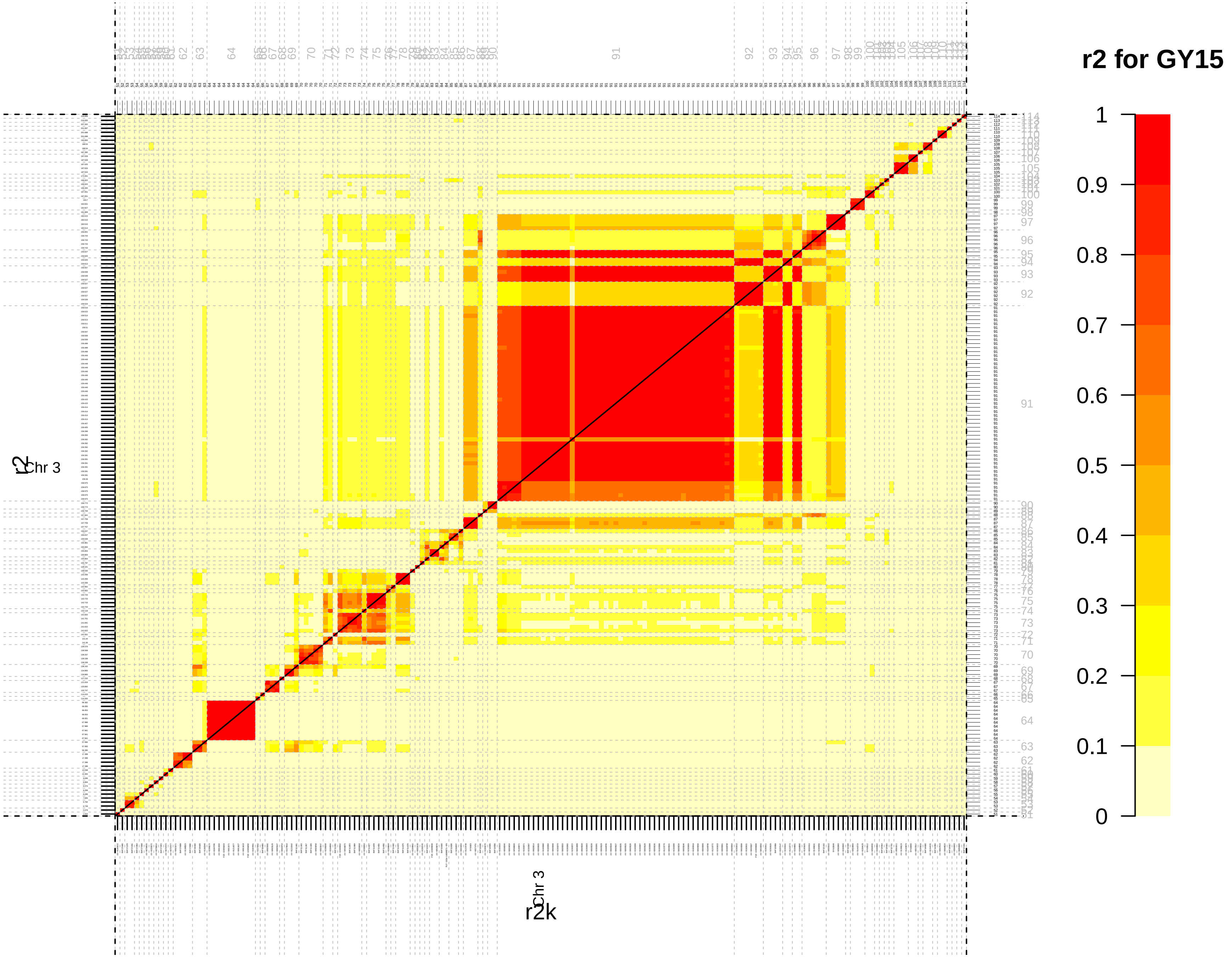

r2 for GY15
Chr 3
r2
r2k
1
0.9
0.8
0.7
0.6
0.5
0.4
0.3
0.2
0.1
0

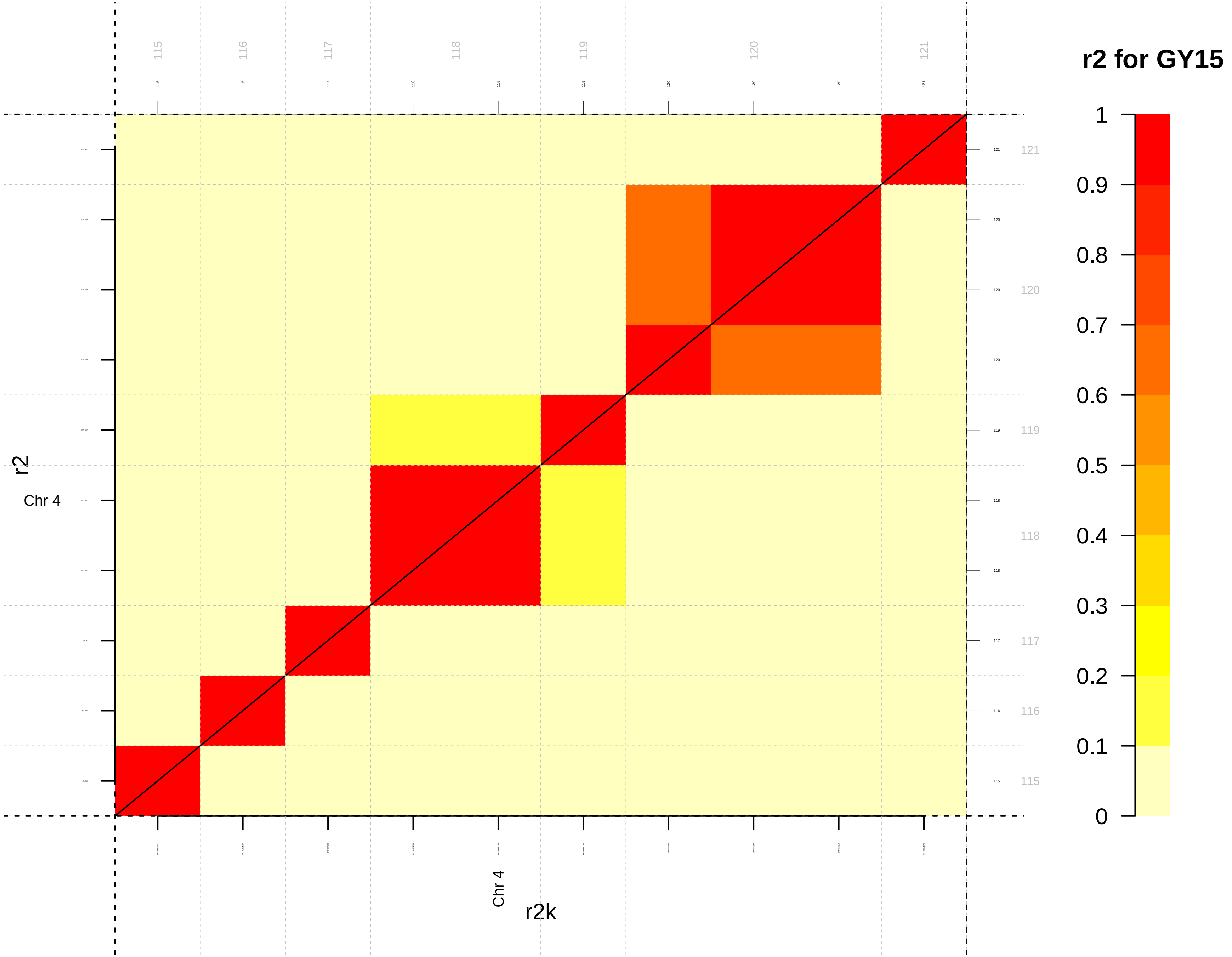

r2 for GY15
1
0.9
0.8
0.7
0.6
0.5
0.4
0.3
0.2
0.1
0
115
116
117
118
119
120
121
r2
Chr 4
r2k

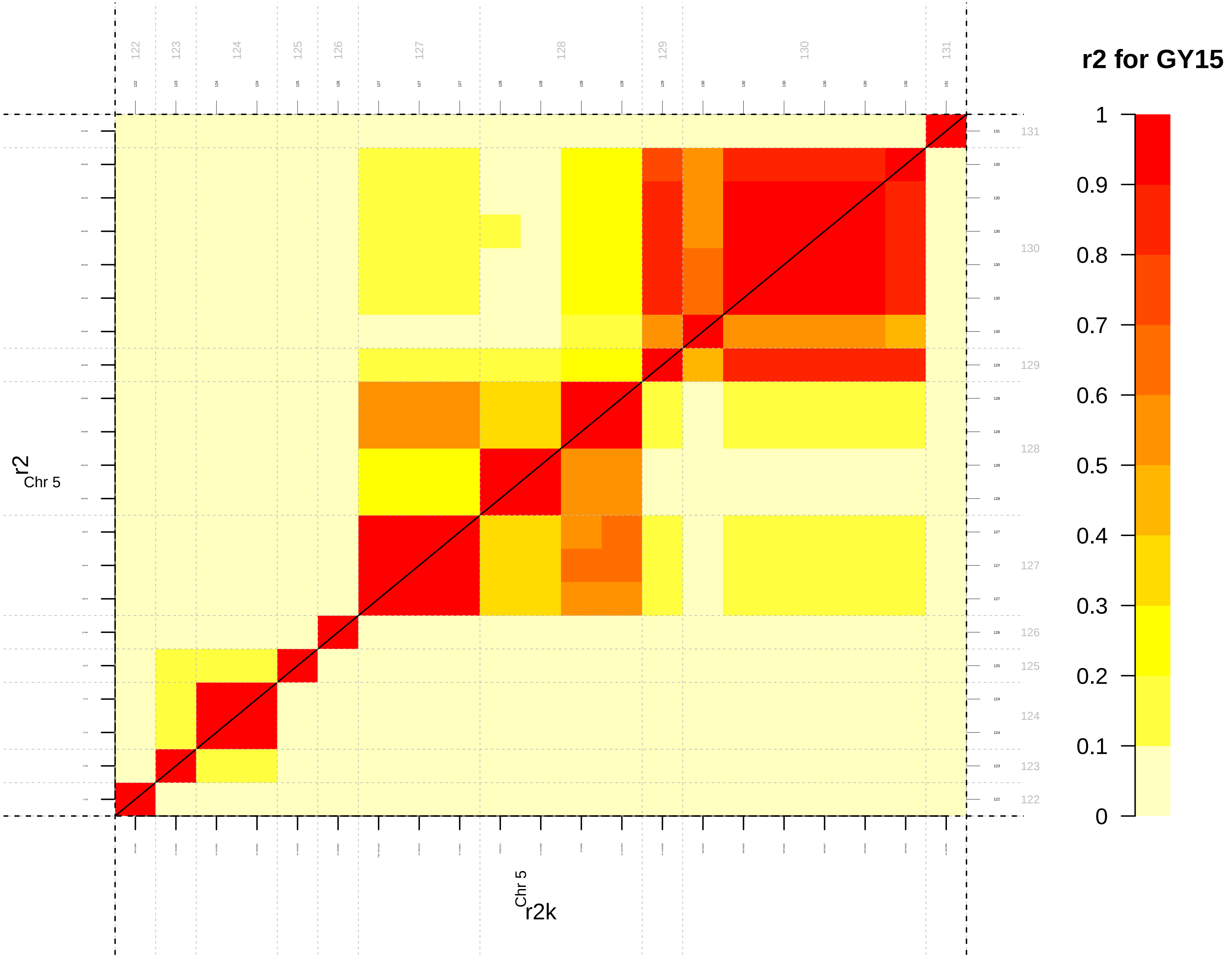

r2 for GY15
1
0.9
0.8
0.7
0.6
0.5
0.4
0.3
0.2
0.1
0
r2
Chr 5
r2k
Chr 5

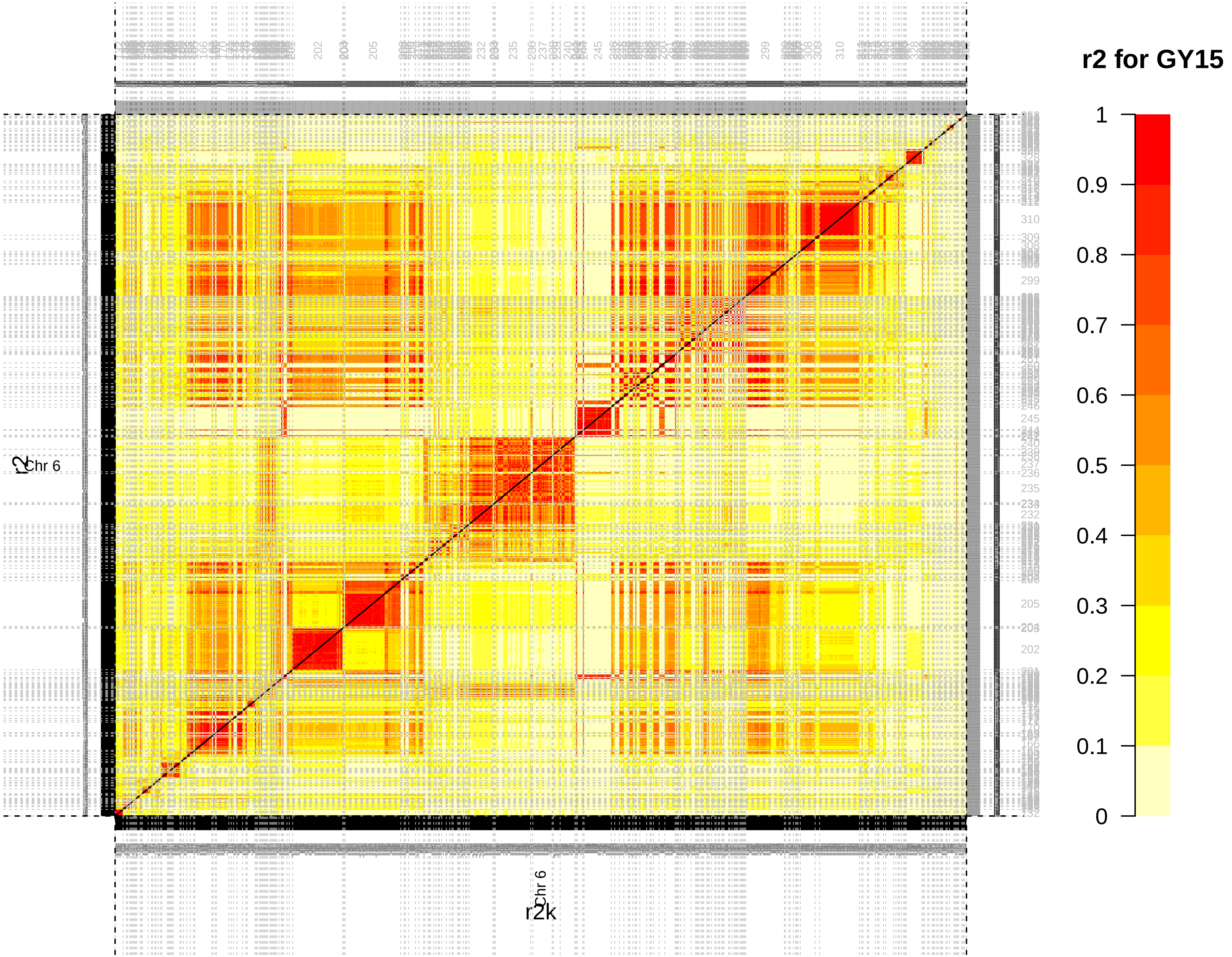
r2 for GY15
1
0.9
0.8
0.7
0.6
0.5
0.4
0.3
0.2
0.1
0
Chr 6
r2
r2k

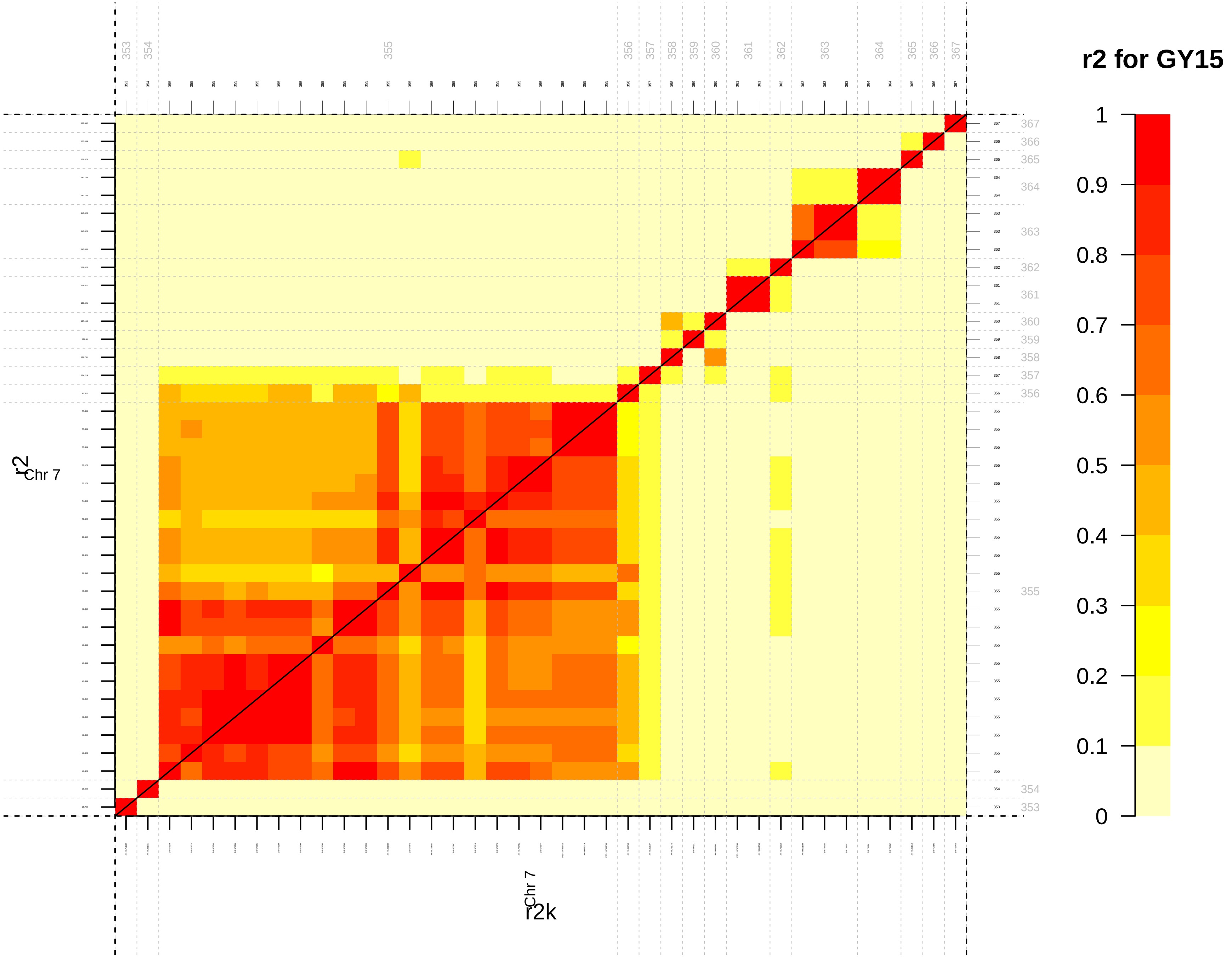

r2 for GY15
1
0.9
0.8
0.7
0.6
0.5
0.4
0.3
0.2
0.1
0
r2
Chr 7
r2k

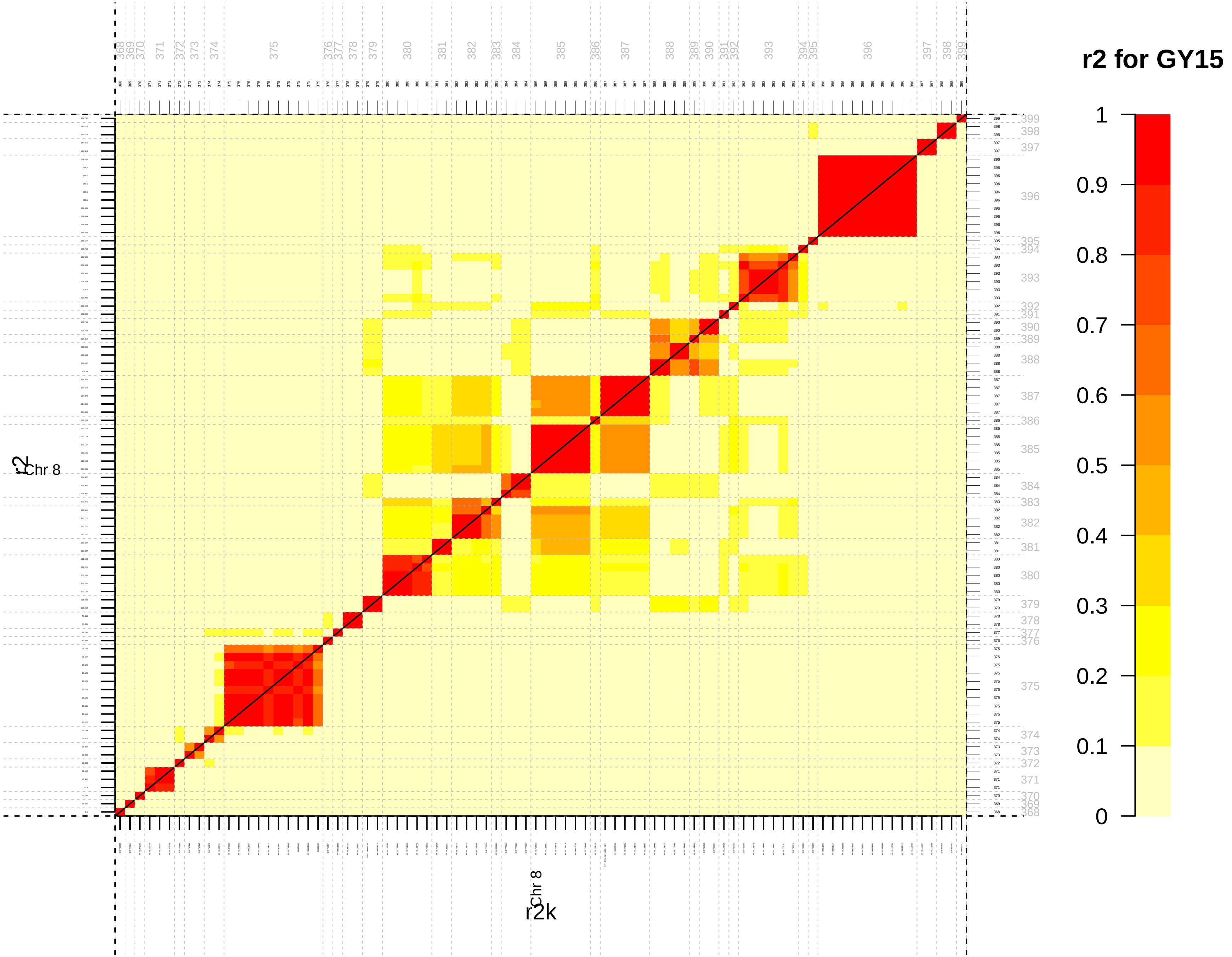

r2 for GY15
1
0.9
0.8
0.7
0.6
0.5
0.4
0.3
0.2
0.1
0
Chr 8
r2
Chr 8
r2k

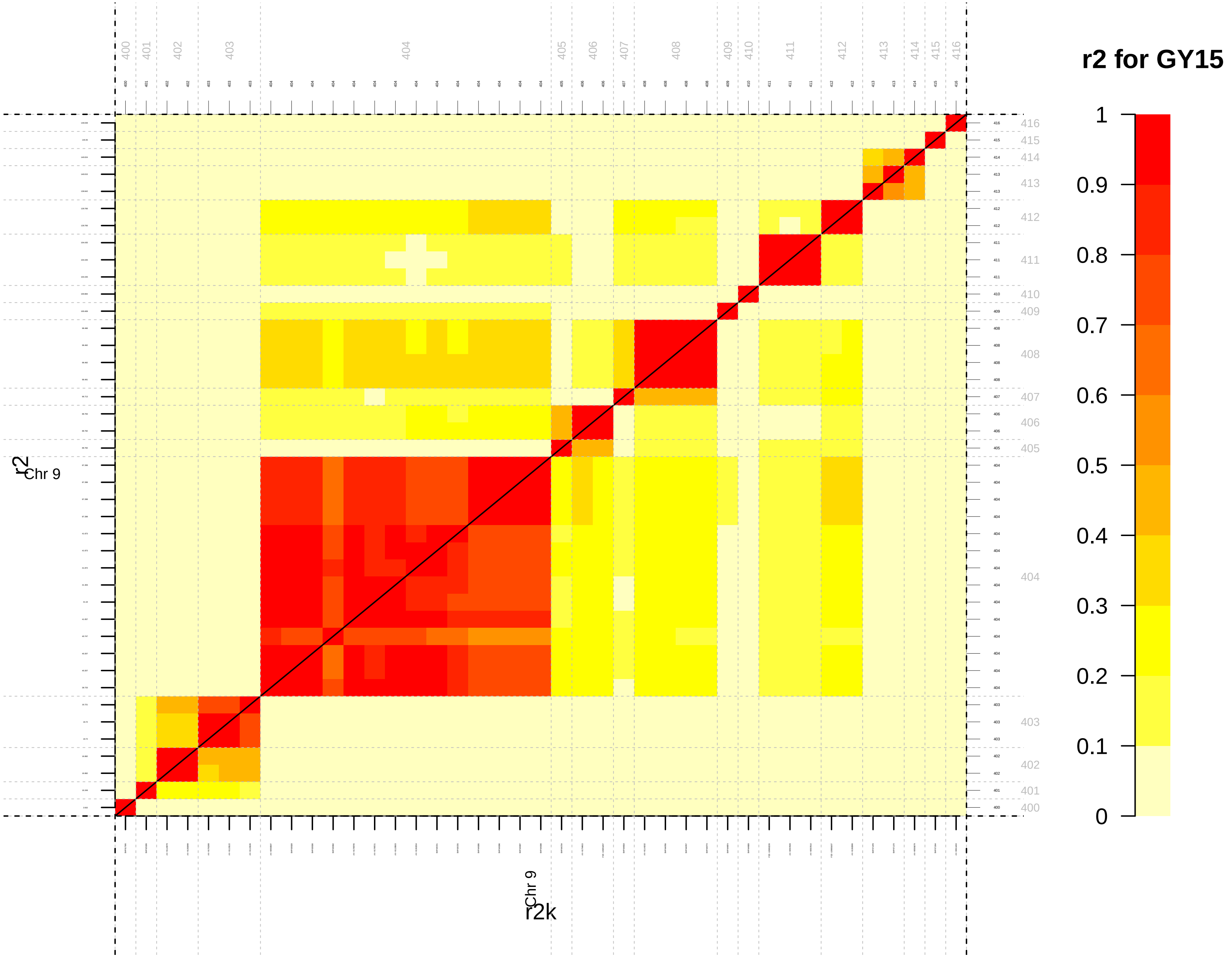

r2 for GY15
1
0.9
0.8
0.7
0.6
0.5
0.4
0.3
0.2
0.1
0
Chr 9
r2
Chr 9
r2k

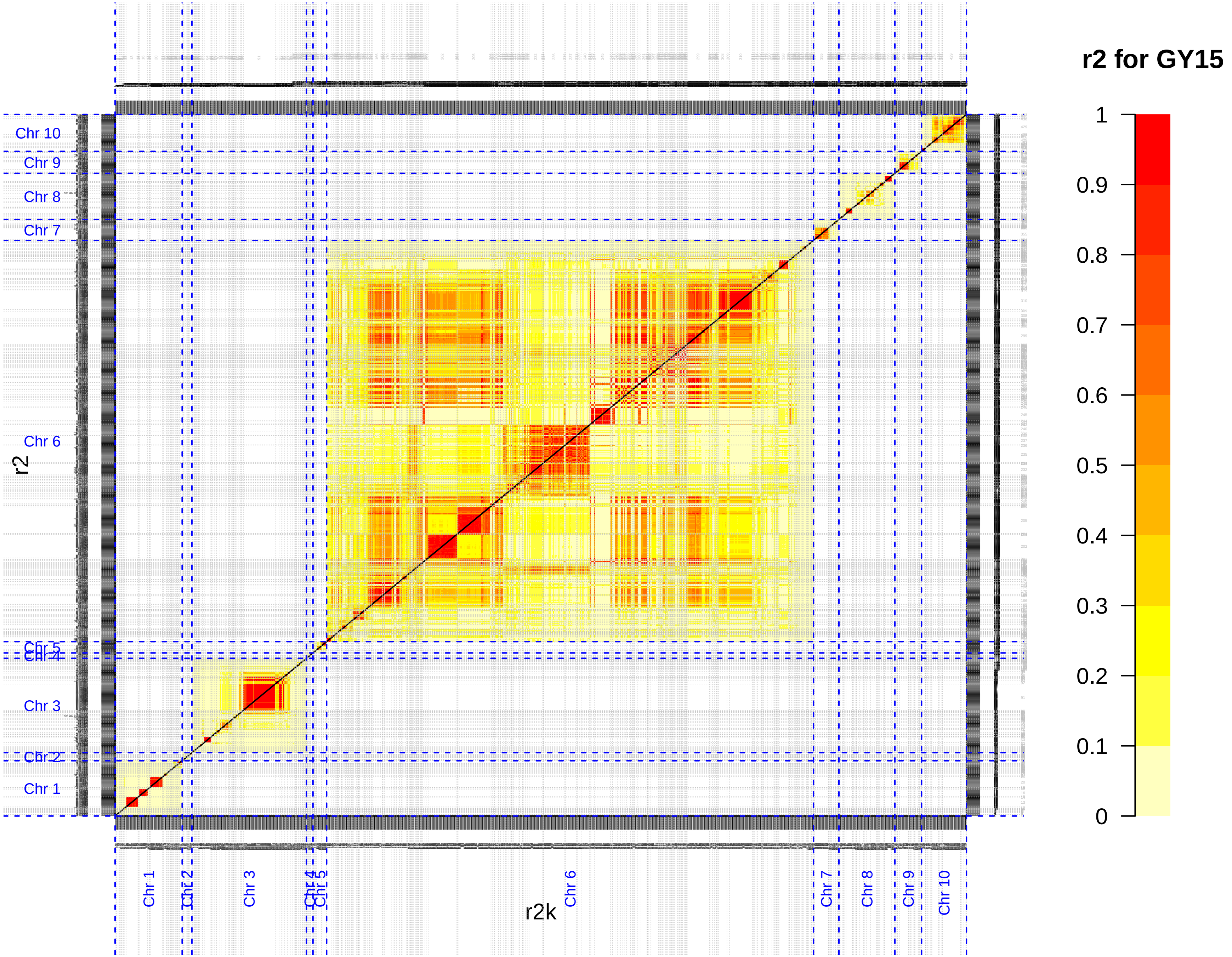
r2 for GY15
1
0.9
0.8
0.7
0.6
0.5
0.4
0.3
0.2
0.1
0
Chr 10
Chr 9
Chr 8
Chr 7
Chr 6
Chr 5
Chr 4
Chr 3
Chr 2
Chr 1
r2
r2k

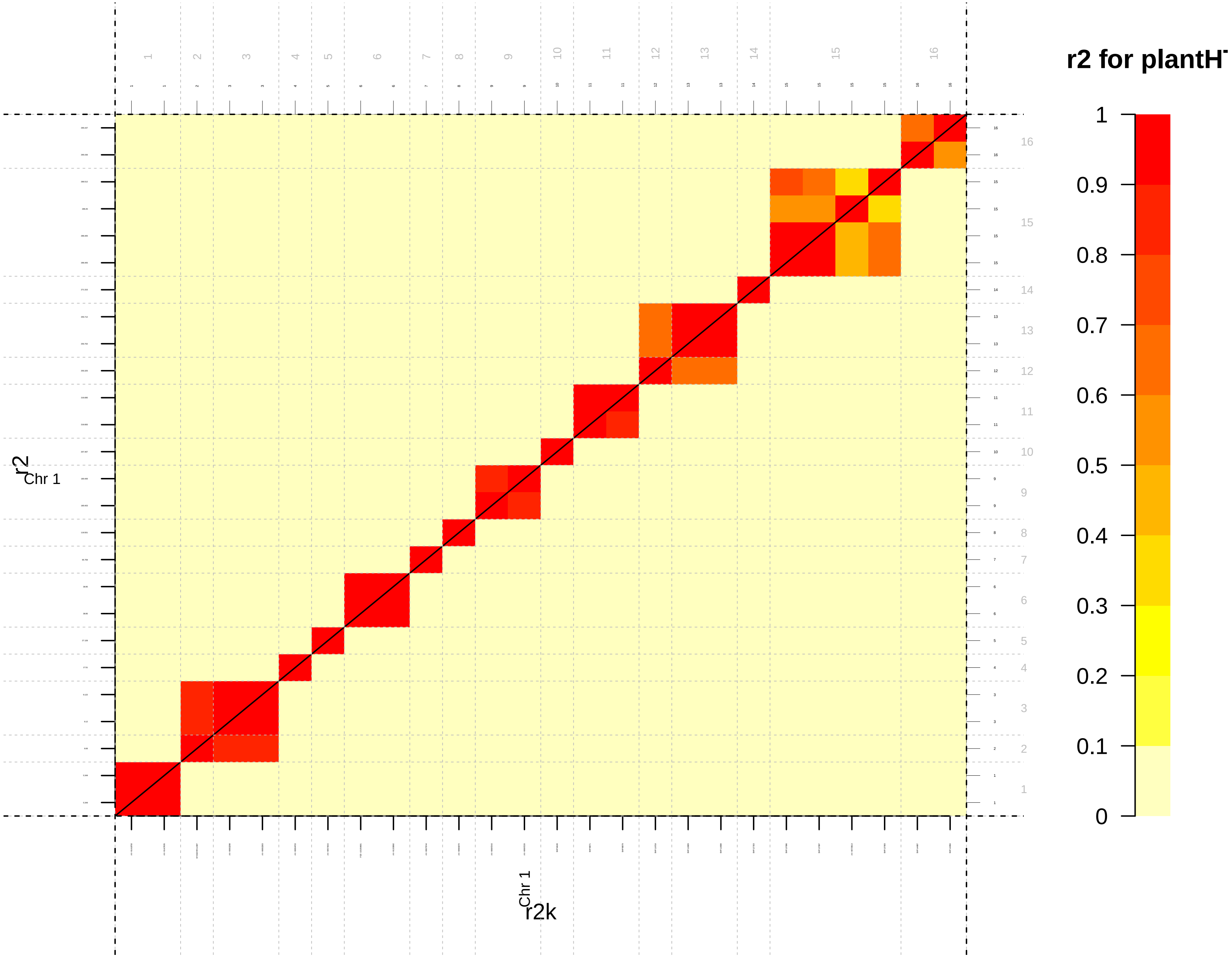
r2 for plantH
1
0.9
0.8
0.7
0.6
0.5
0.4
0.3
0.2
0.1
0
r2
Chr 1
r2k
Chr 1

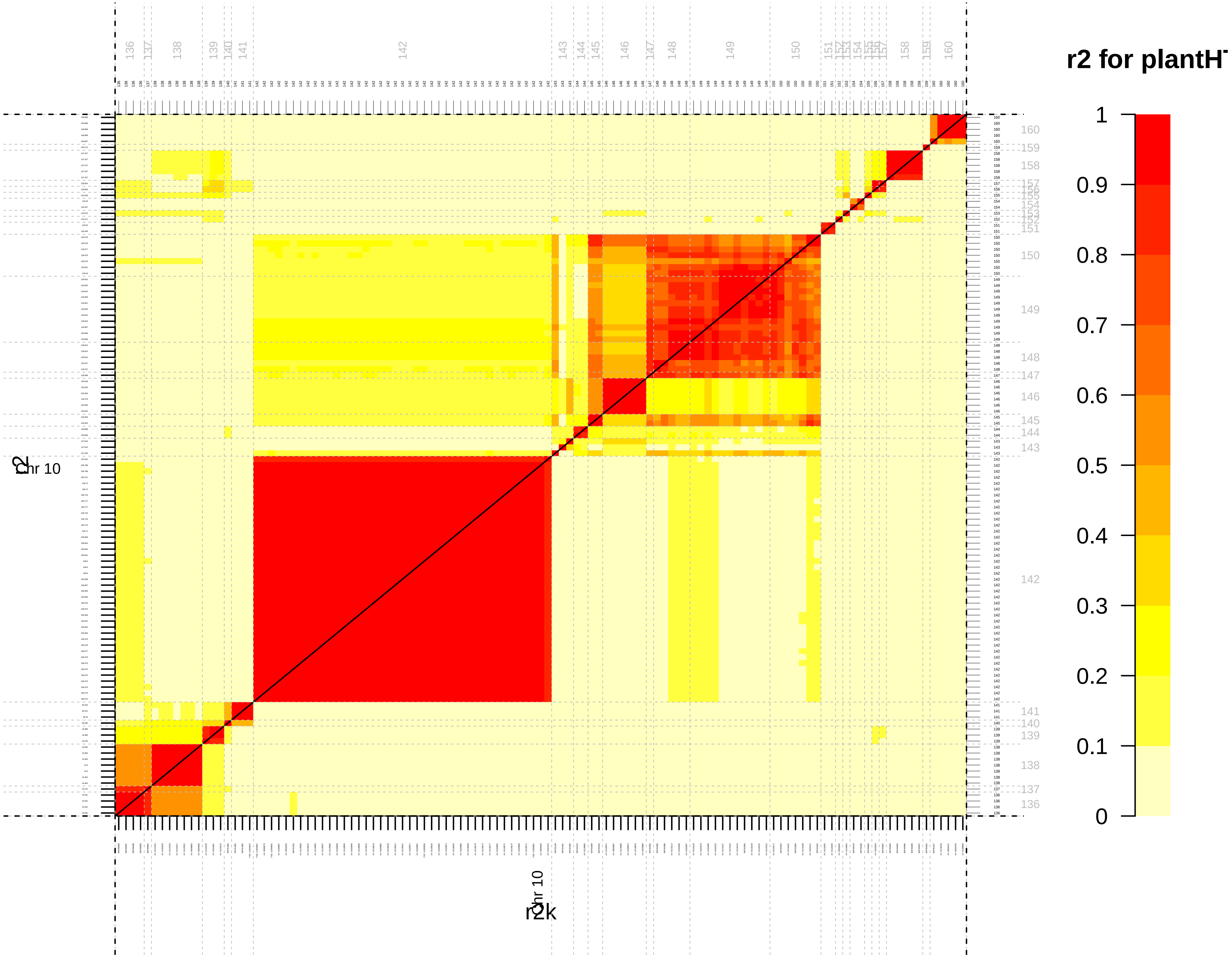

r2 for plantH
1
0.9
0.8
0.7
0.6
0.5
0.4
0.3
0.2
0.1
0
chr 10
r2
r2k

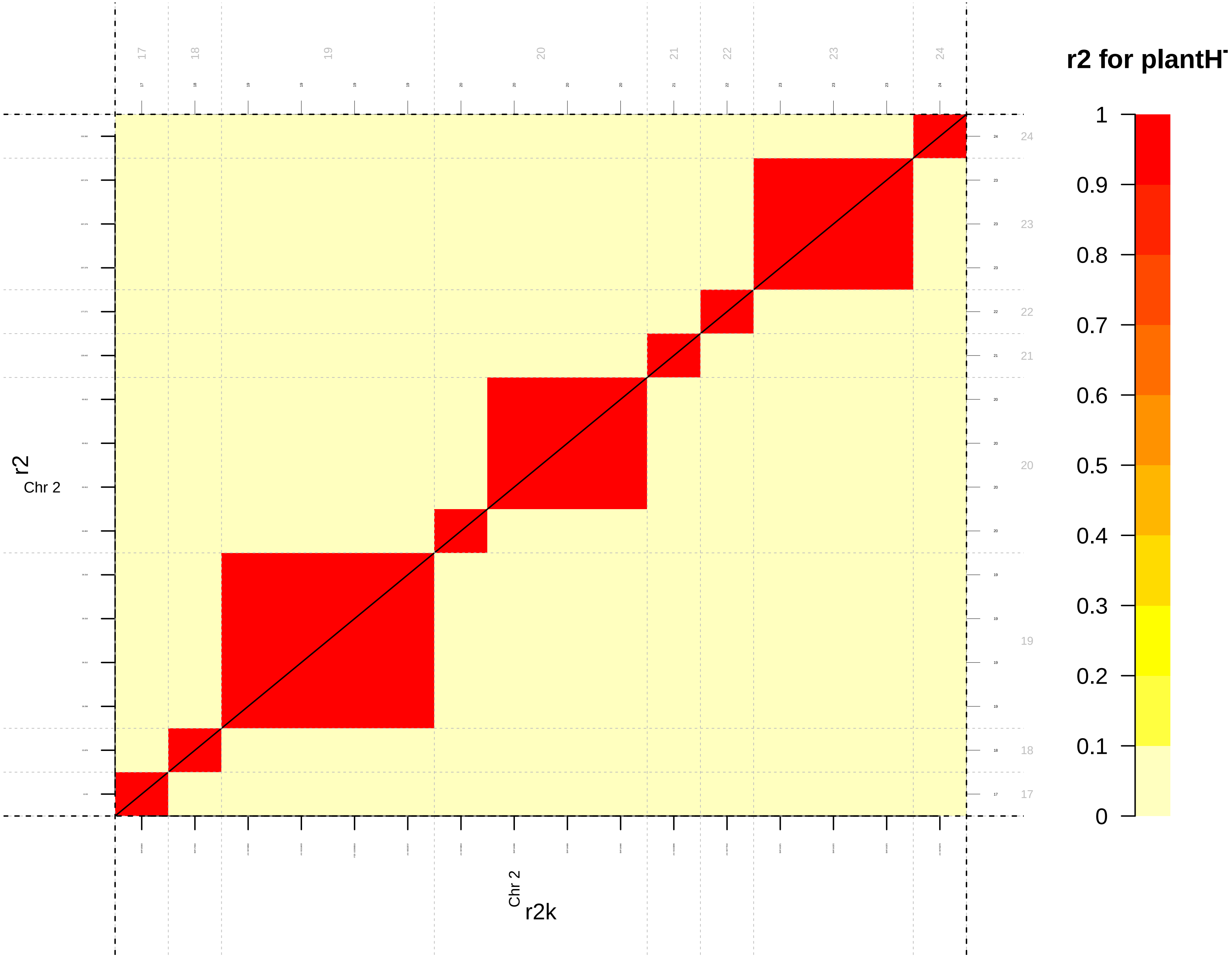
r2 for plantH
1
0.9
0.8
0.7
0.6
0.5
0.4
0.3
0.2
0.1
0
r2
Chr 2
r2k
Chr 2
17
18
19
20
21
22
23
24

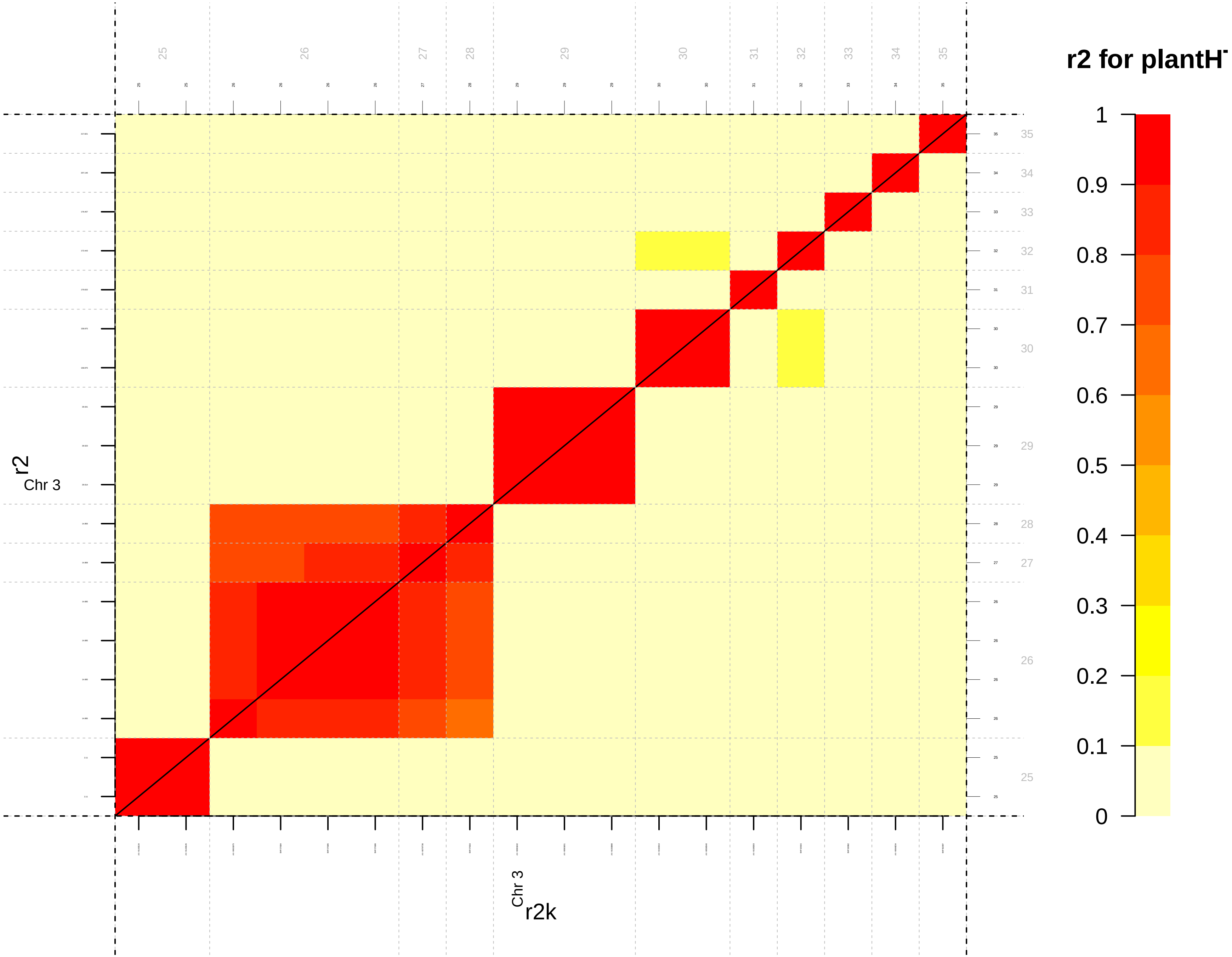

r2 for plantH
1
0.9
0.8
0.7
0.6
0.5
0.4
0.3
0.2
0.1
0
r2
Chr 3
r2k
Chr 3

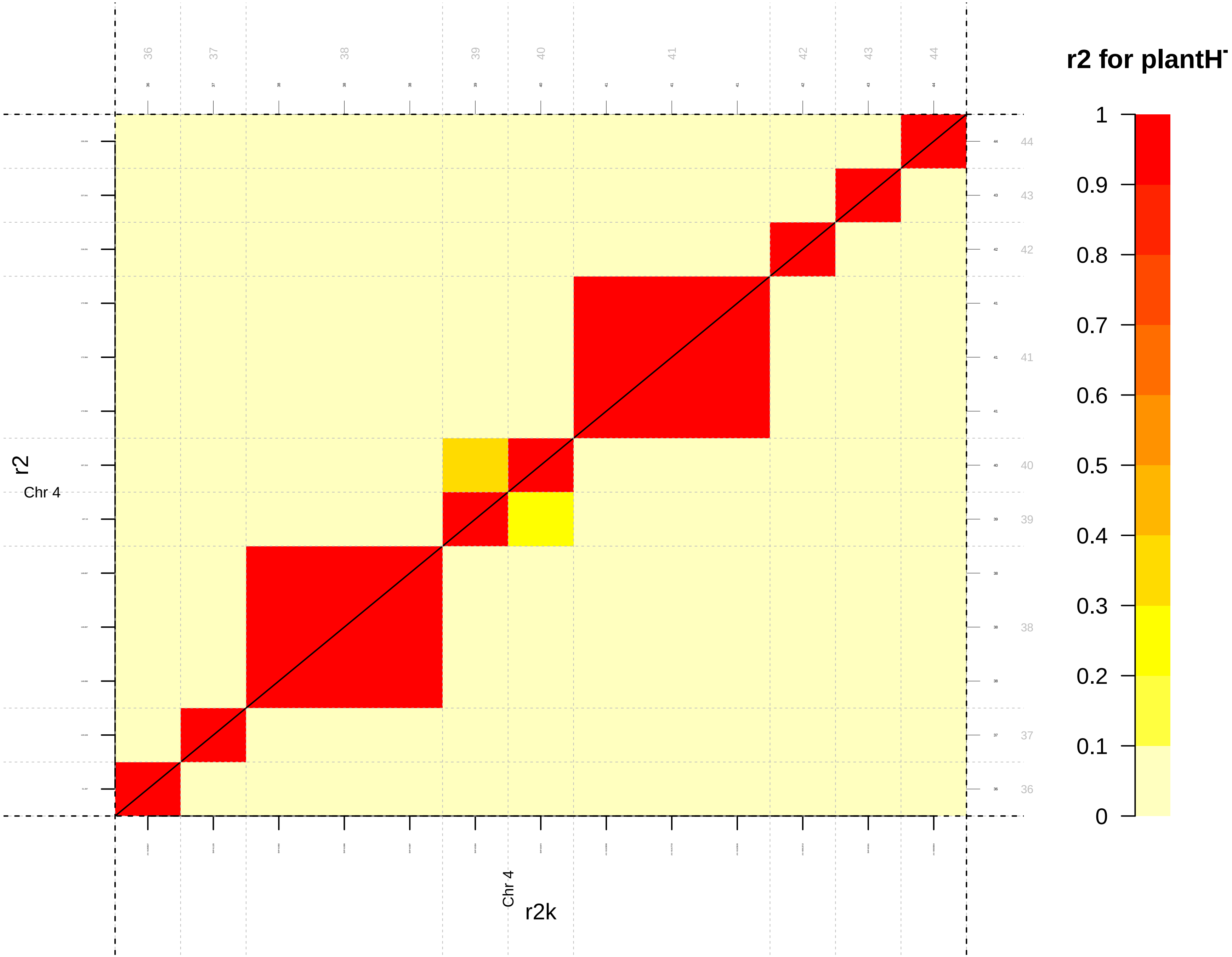
r2 for plantH
1
0.9
0.8
0.7
0.6
0.5
0.4
0.3
0.2
0.1
0
36
37
38
39
40
41
42
43
44
r2
Chr 4
r2k
Chr 4

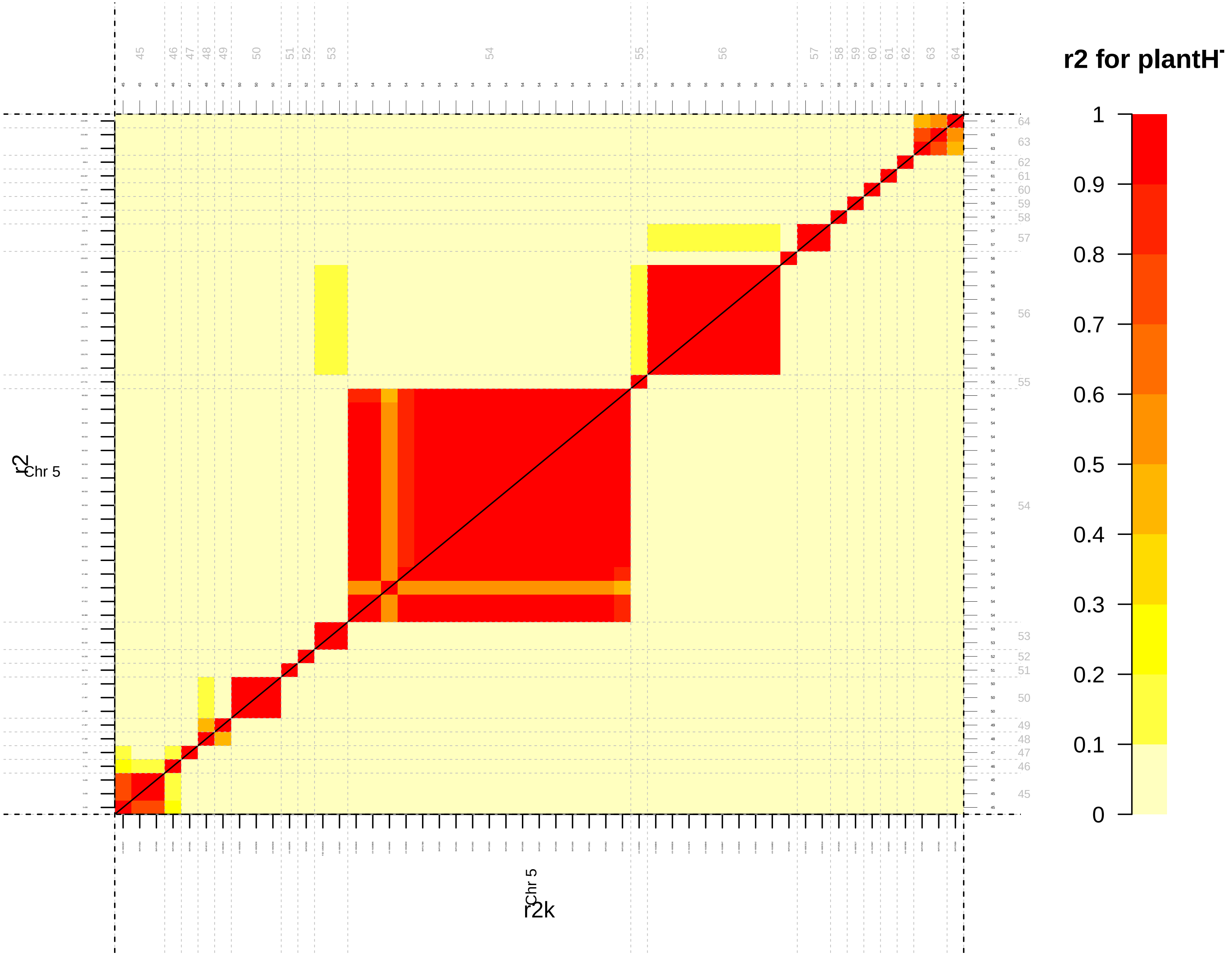
r2 for plantH
1
0.9
0.8
0.7
0.6
0.5
0.4
0.3
0.2
0.1
0
45 46 47 48 49 50 51 52 53 54 55 56 57 58 59 60 61 62 63 64
64 63 62 61 60 59 58 57 56 55 54 53 52 51 50 49 48 47 46 45
r2
Chr 5
Chr 5
r2k

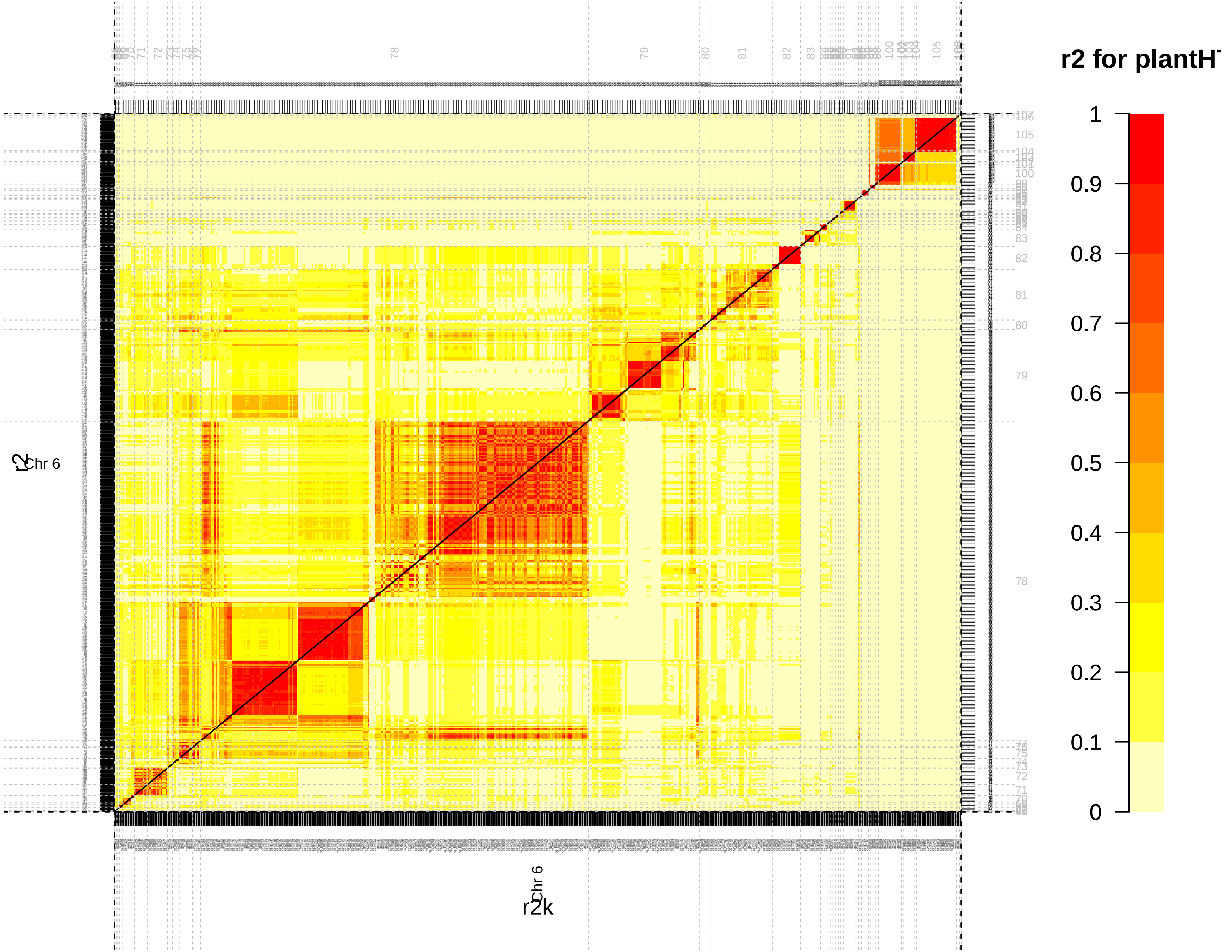

r2 for plantH
1
0.9
0.8
0.7
0.6
0.5
0.4
0.3
0.2
0.1
0
Chr 6
r2
r2k

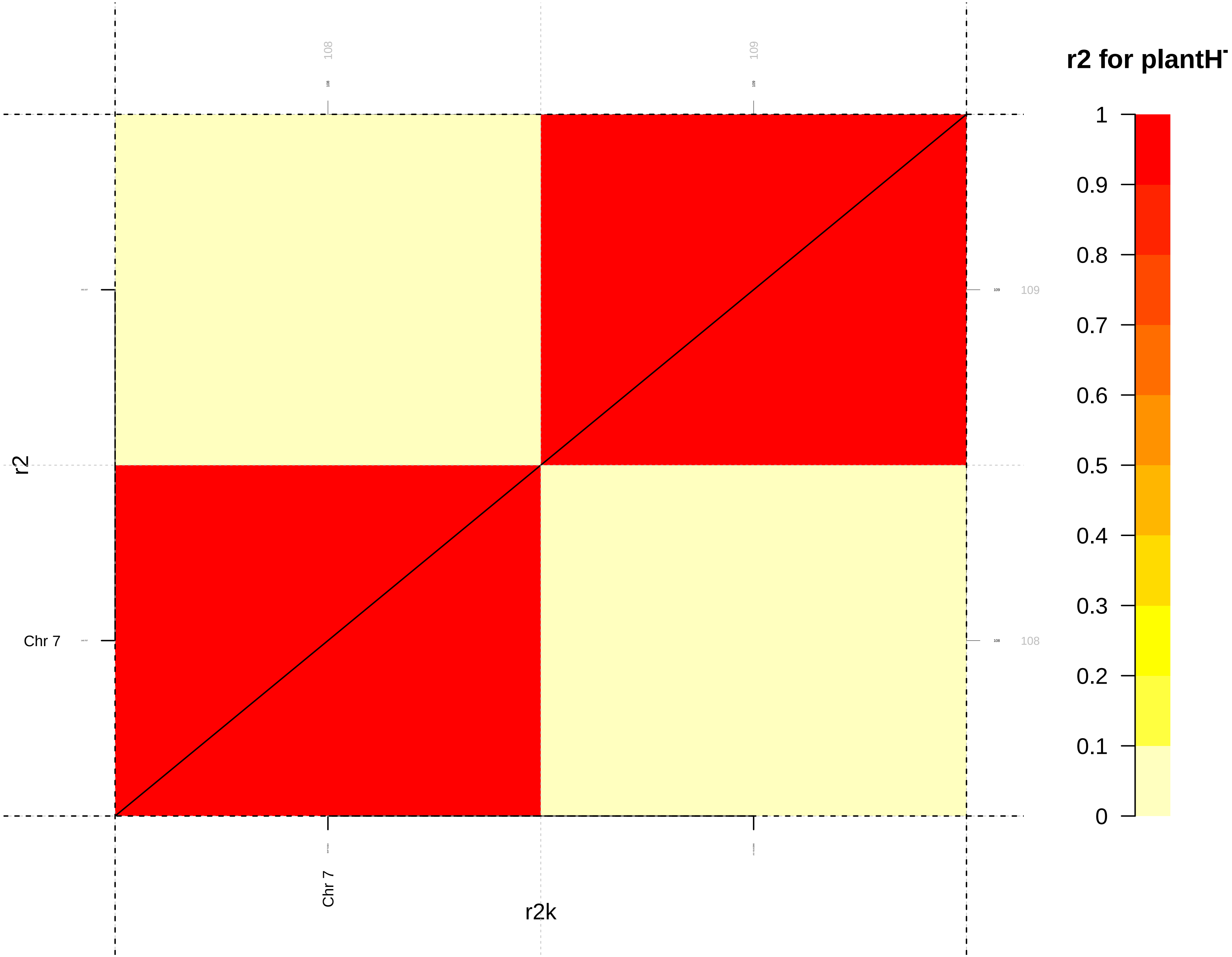
r2 for plantH
1
0.9
0.8
0.7
0.6
0.5
0.4
0.3
0.2
0.1
0
108
109
109
108
r2
Chr 7
Chr 7
r2k

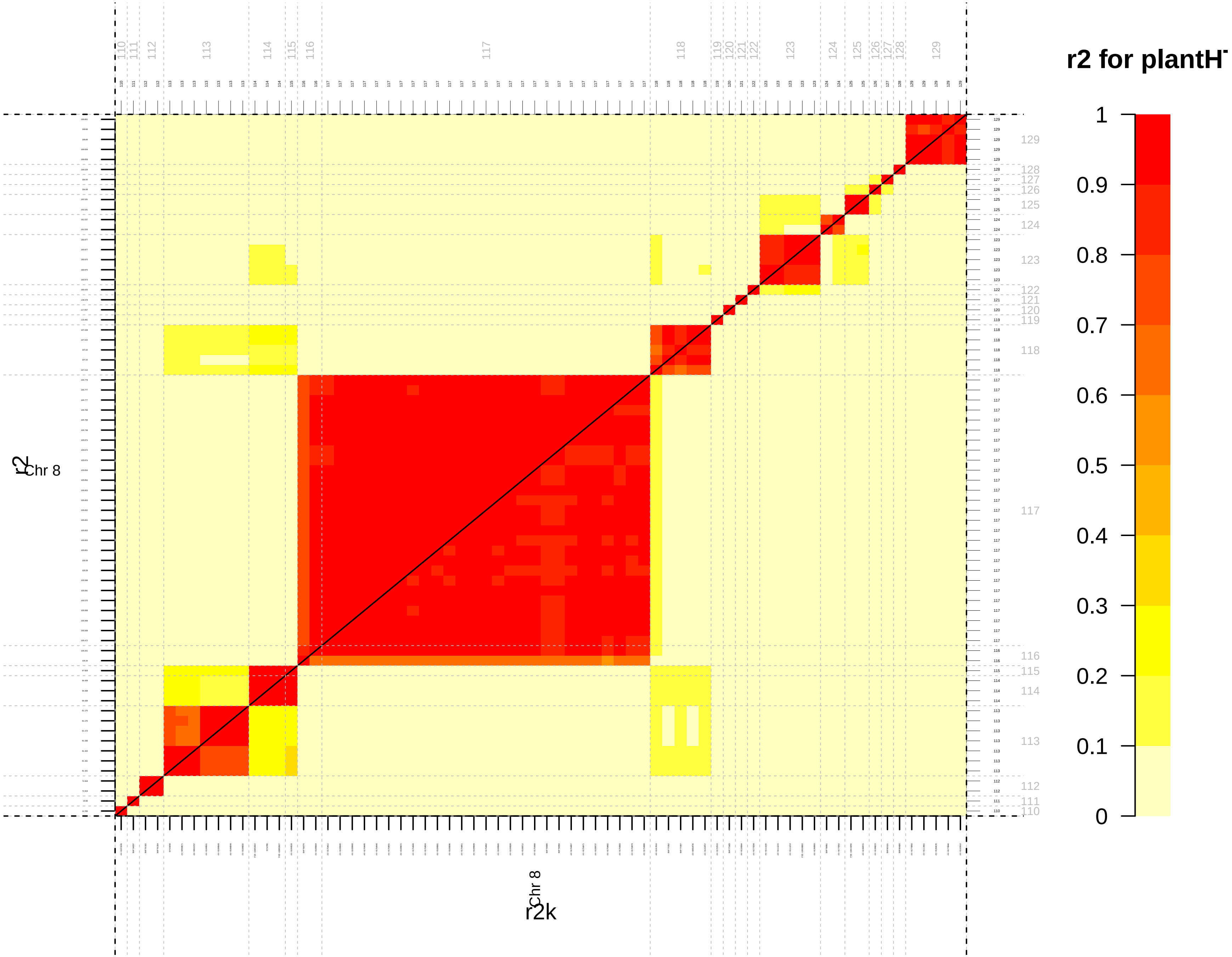

r2 for plantH
1
0.9
0.8
0.7
0.6
0.5
0.4
0.3
0.2
0.1
0
Chr 8
r2
Chr 8
r2k

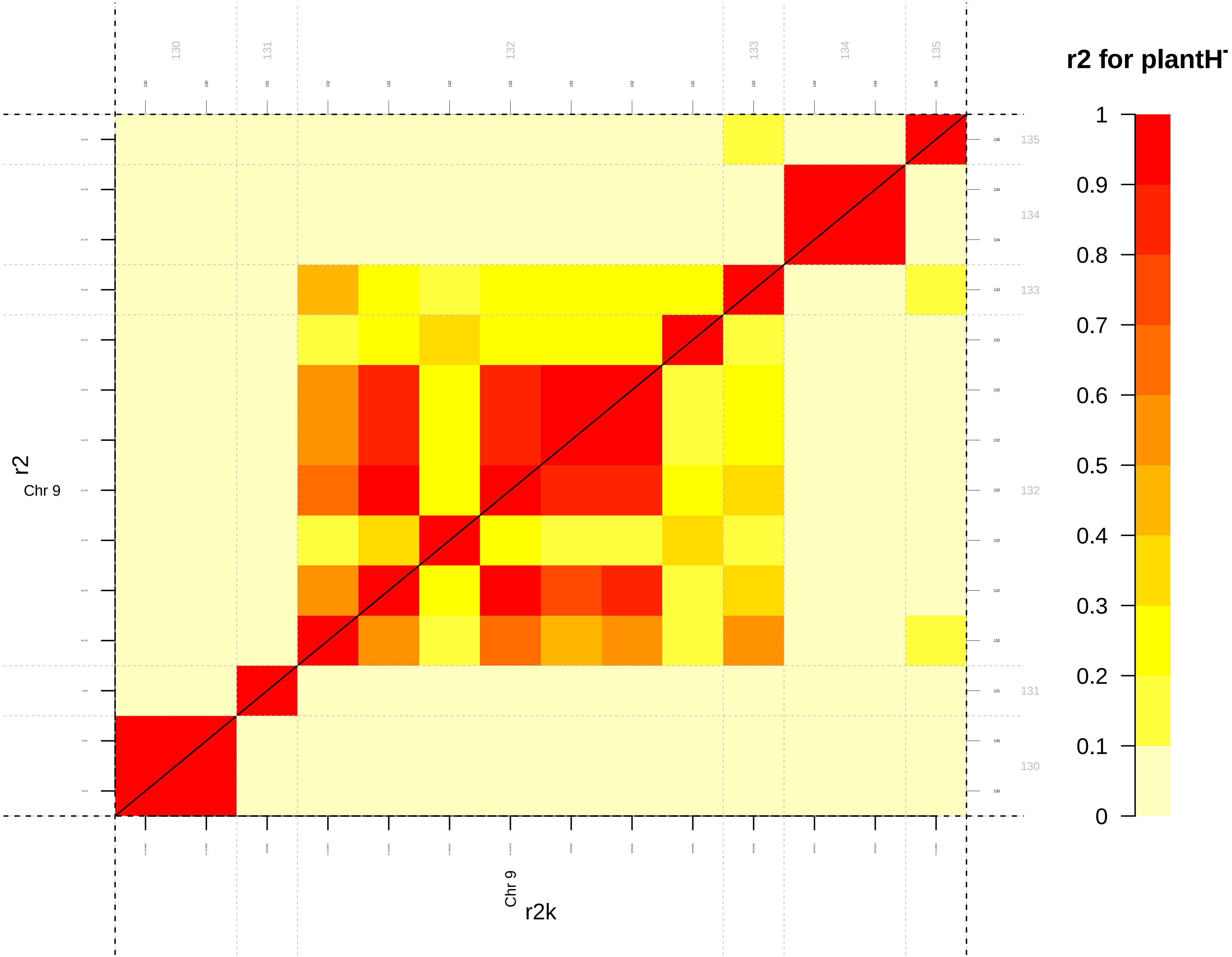

r2 for plantH
1
0.9
0.8
0.7
0.6
0.5
0.4
0.3
0.2
0.1
0
130
131
132
133
134
135
r2
Chr 9
r2k

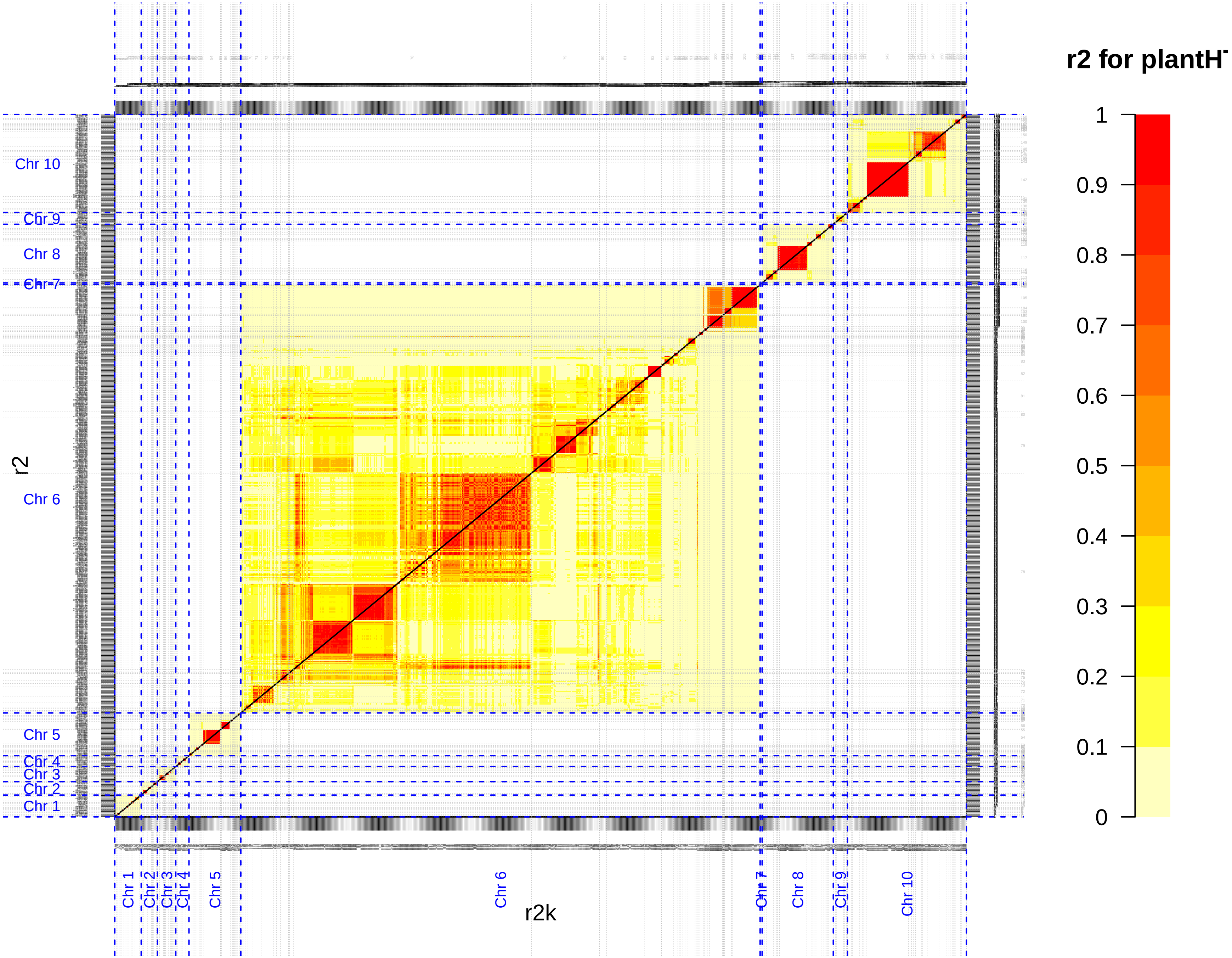

r2 for plantH
1
0.9
0.8
0.7
0.6
0.5
0.4
0.3
0.2
0.1
0
r2
r2k
Chr 10
Chr 9
Chr 8
Chr 7
Chr 6
Chr 5
Chr 4
Chr 3
Chr 2
Chr 1

**Additional file 7 (.pdf):**

**Figure S7:** QTL limits obtained by the *LD_Adj* approach projected on heatmaps representing the level of LD between associated SNPs for each trait (DTA: male flowering time, plantHT: plant height and GY: grain yield) for each chromosome. Upper and lower triangles on the heatmaps represent the $r^2$ and $r^2K$ values between associated SNPs, respectively. Linkage disequilibrium between loci was colored according to values from weak LD (yellow) to high LD (red). The significant markers were ordered according to their physical positions on the chromosome and were represented by ticks on the four sides of the heatmaps. Limits of QTLs were displayed by gray dotted lines. QTL numbers were indicated in gray on the top and the right of each heatmap.
