## Additional file 8: Fig S8 for "Genotyping-by-sequencing and SNP-arrays are complementary for detecting quantitative trait loci by tagging different haplotypes in association studies"

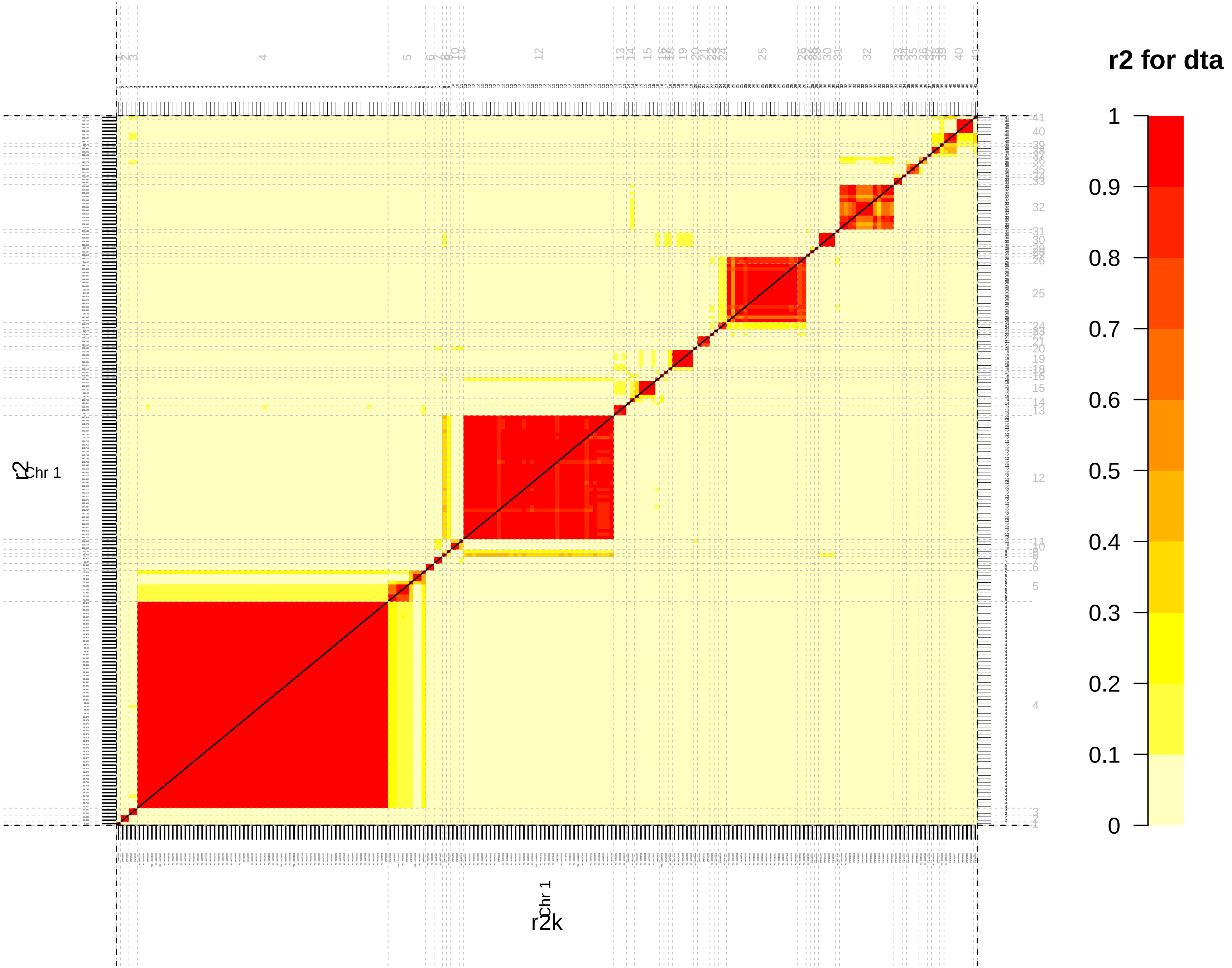

r2 for dta
1
0.9
0.8
0.7
0.6
0.5
0.4
0.3
0.2
0.1
0
Chr 1
r2k
Chr 1
r2

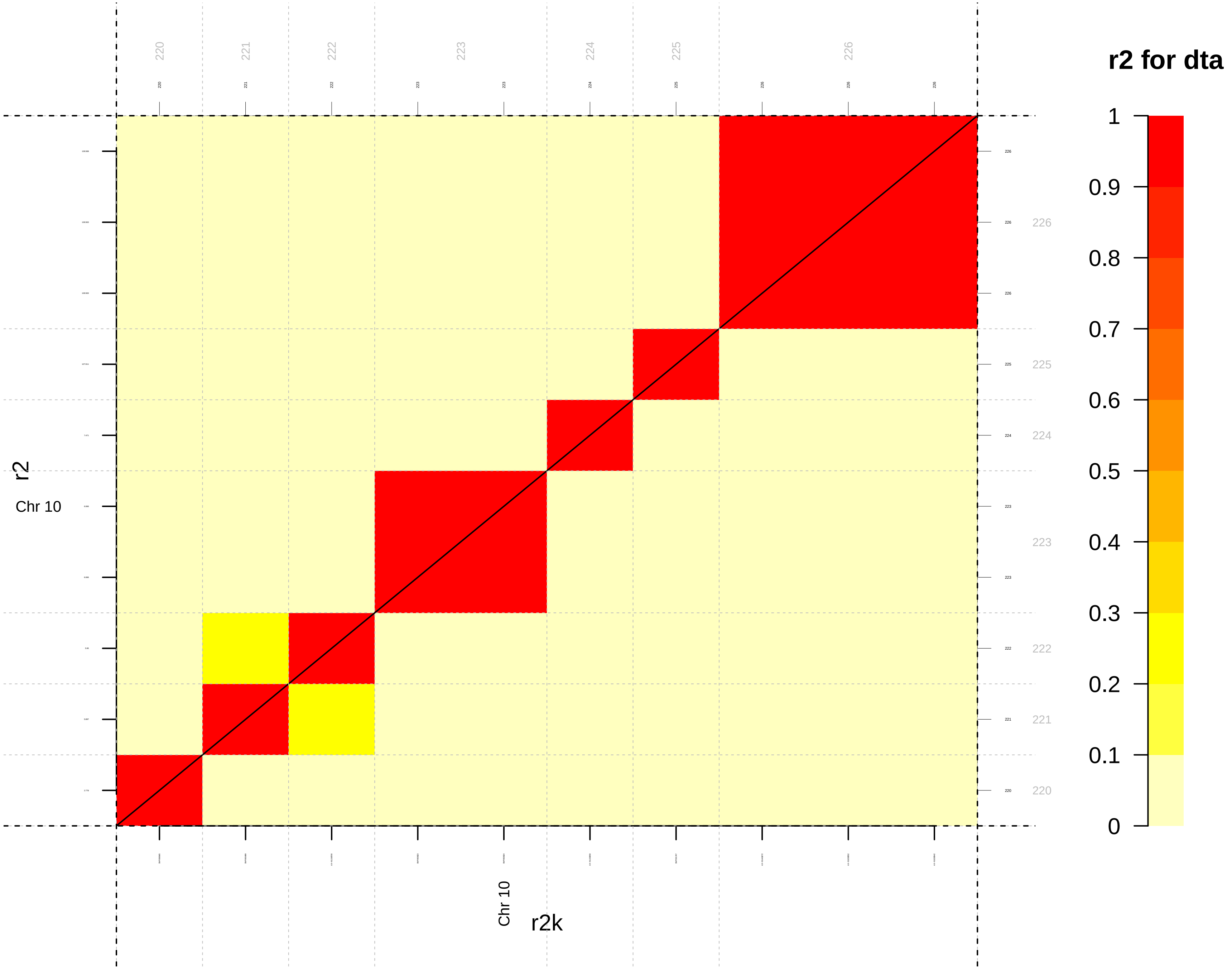
r2 for dta
1
0.9
0.8
0.7
0.6
0.5
0.4
0.3
0.2
0.1
0
r2
Chr 10
Chr 10
r2k
220
221
222
223
224
225
226

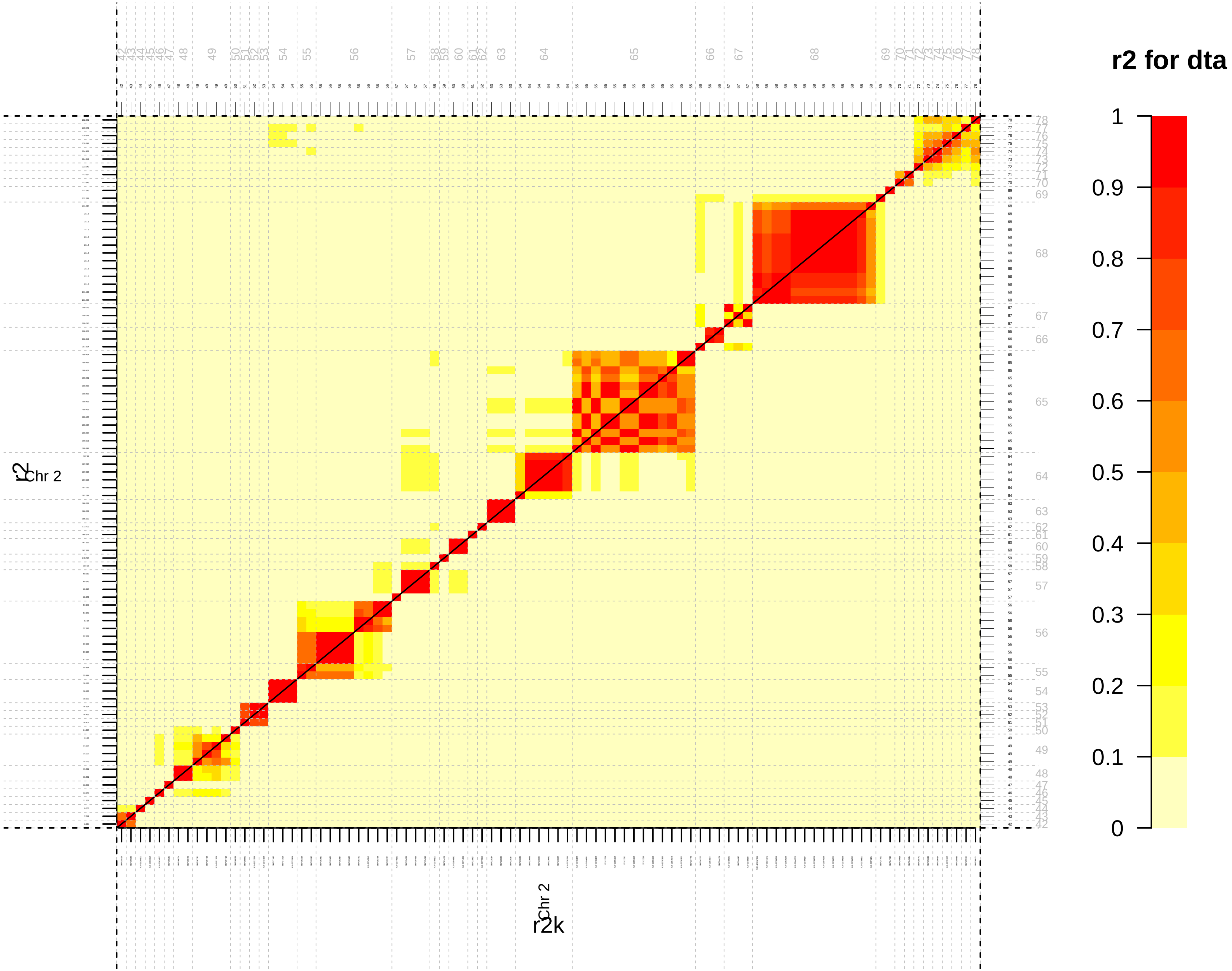
r2 for dta
1
0.9
0.8
0.7
0.6
0.5
0.4
0.3
0.2
0.1
0
Chr 2
r2
Chr 2
r2k

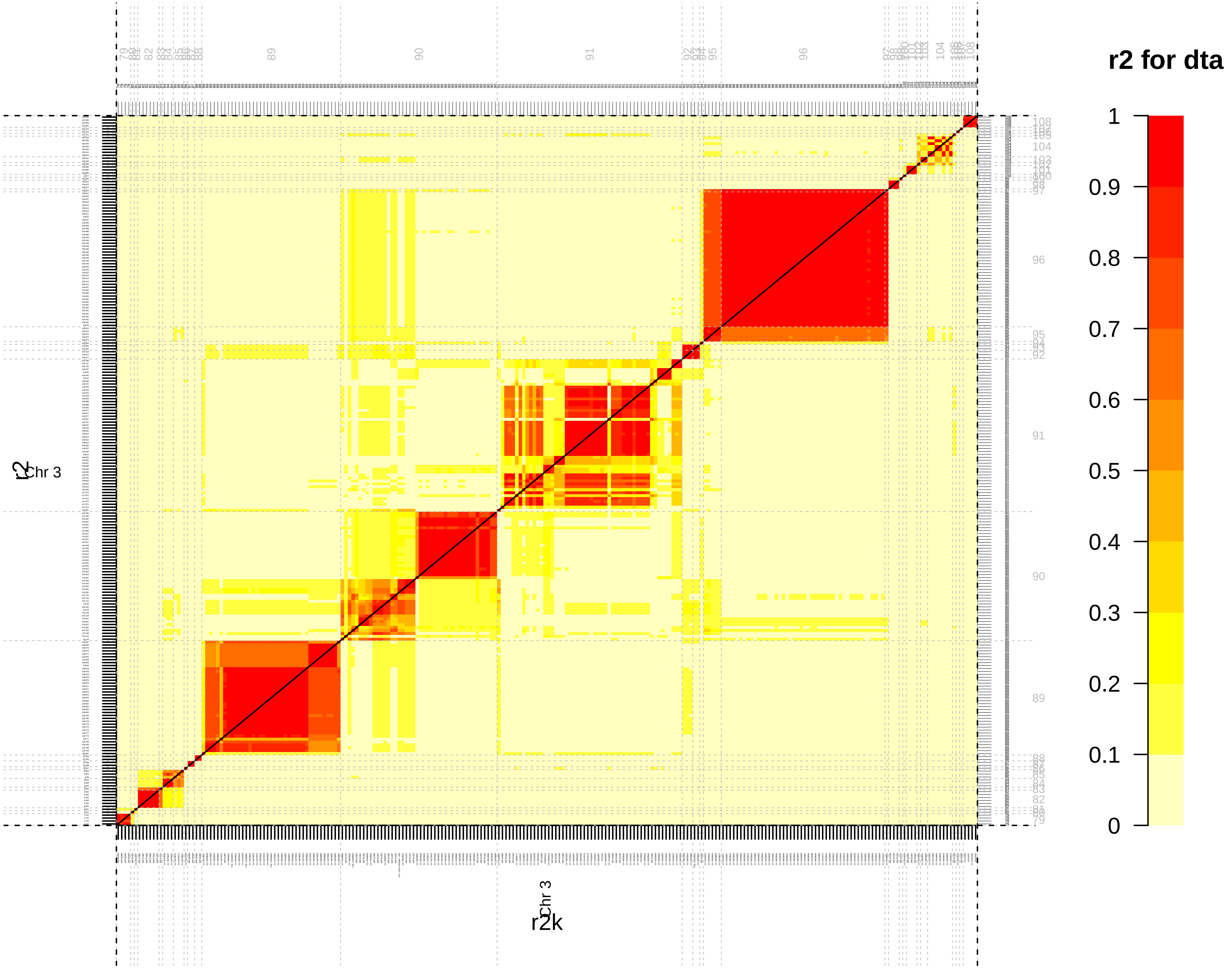
r2 for dta
1
0.9
0.8
0.7
0.6
0.5
0.4
0.3
0.2
0.1
0
Chr 3
r2
Chr 3
r2k

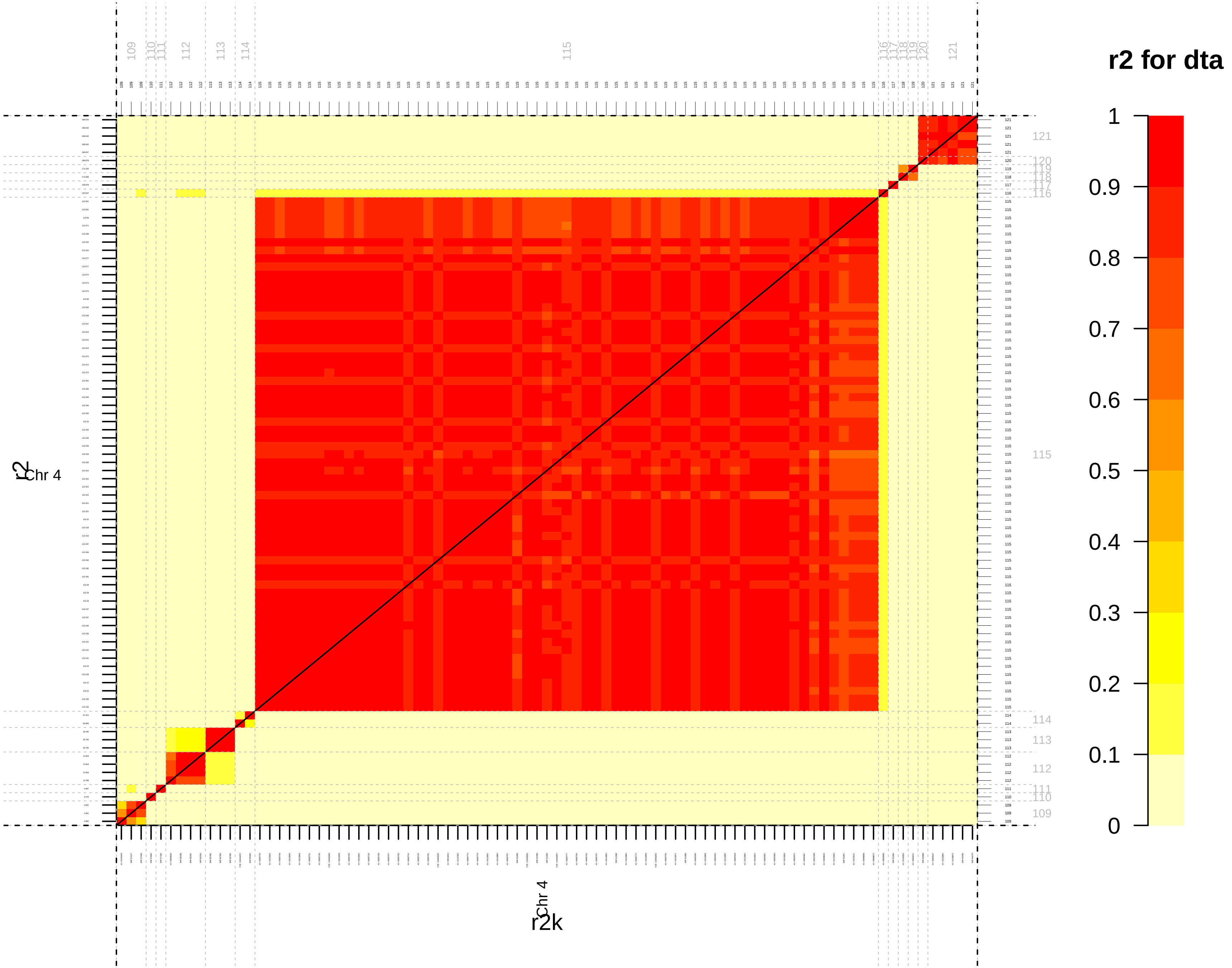

r2 for dta
1
0.9
0.8
0.7
0.6
0.5
0.4
0.3
0.2
0.1
0
Chr 4
r2
Chr 4
r2k

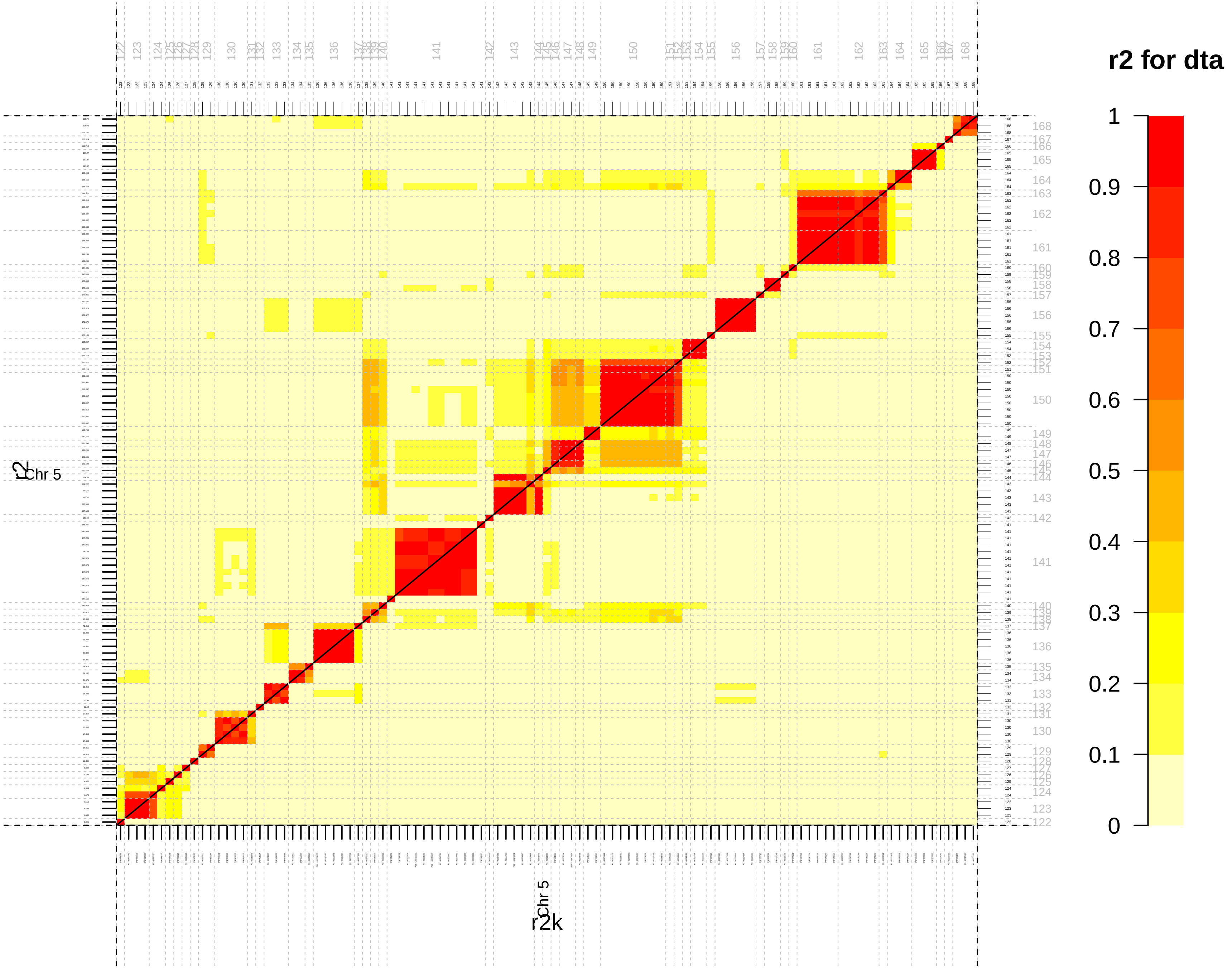
r2 for dta
1
0.9
0.8
0.7
0.6
0.5
0.4
0.3
0.2
0.1
0
Chr 5
r2
Chr 5
r2k

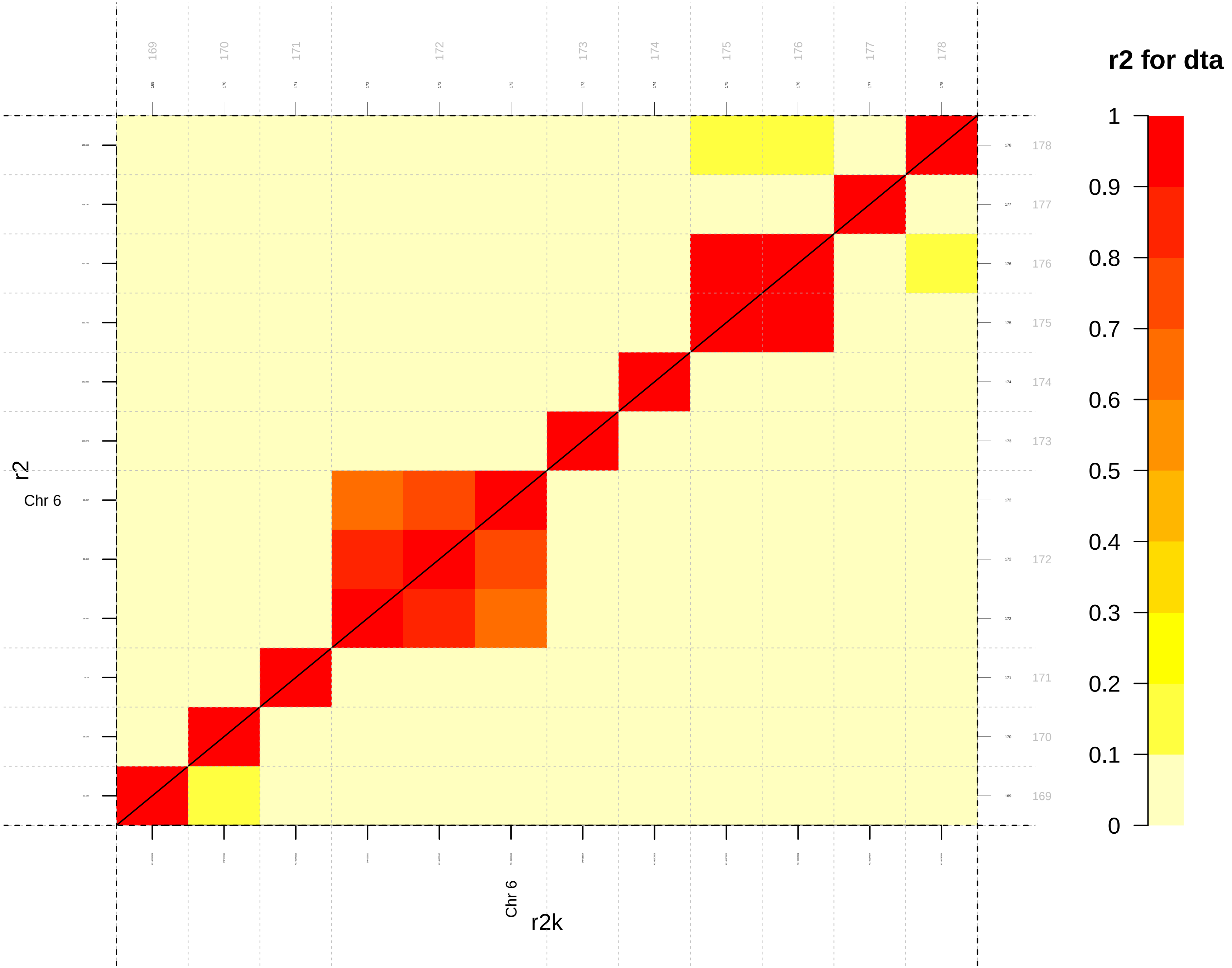

r2 for dta
r2
Chr 6
r2k
1
0.9
0.8
0.7
0.6
0.5
0.4
0.3
0.2
0.1
0

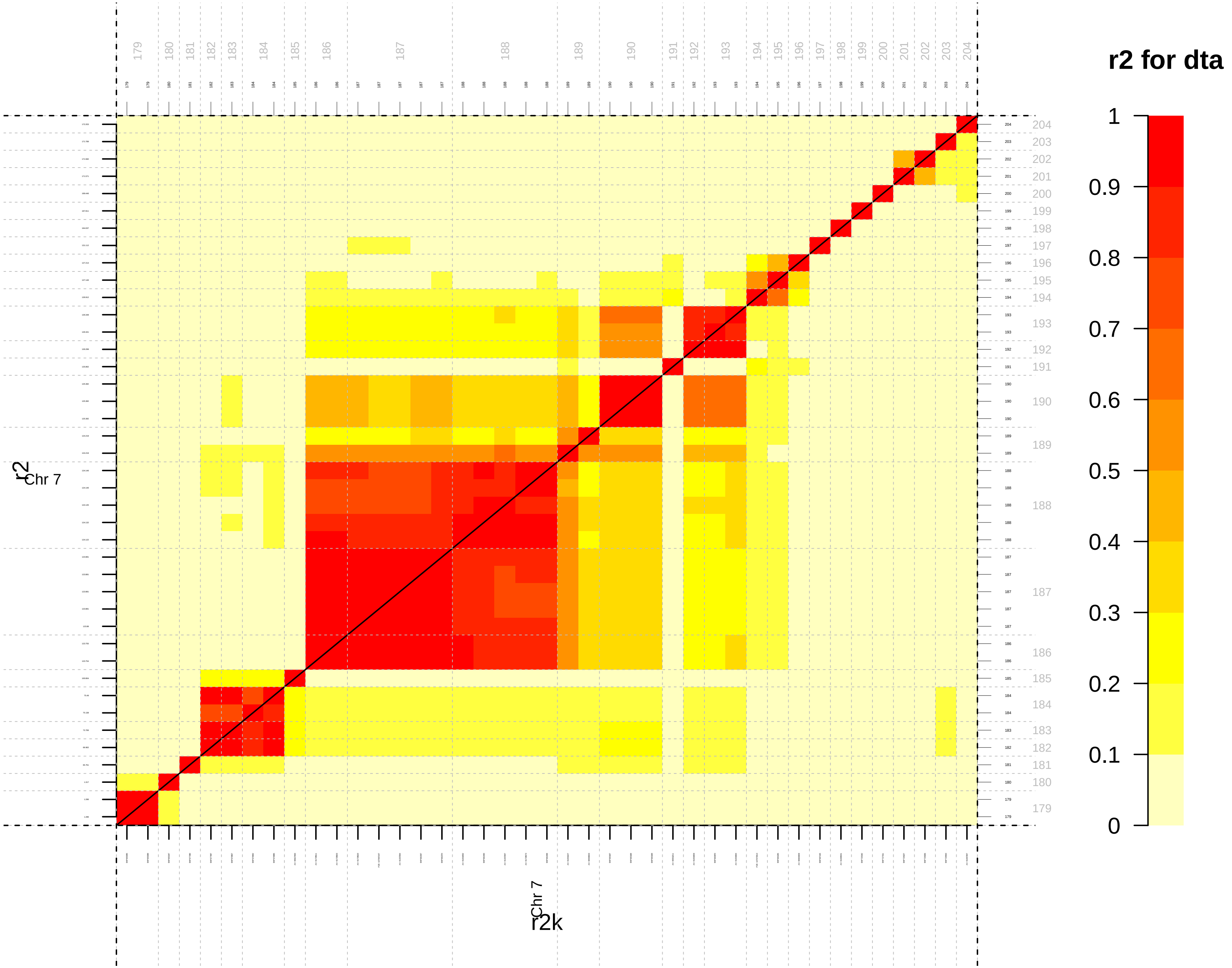

r2 for dta
1
0.9
0.8
0.7
0.6
0.5
0.4
0.3
0.2
0.1
0
r2
Chr 7
r2k
Chr 7

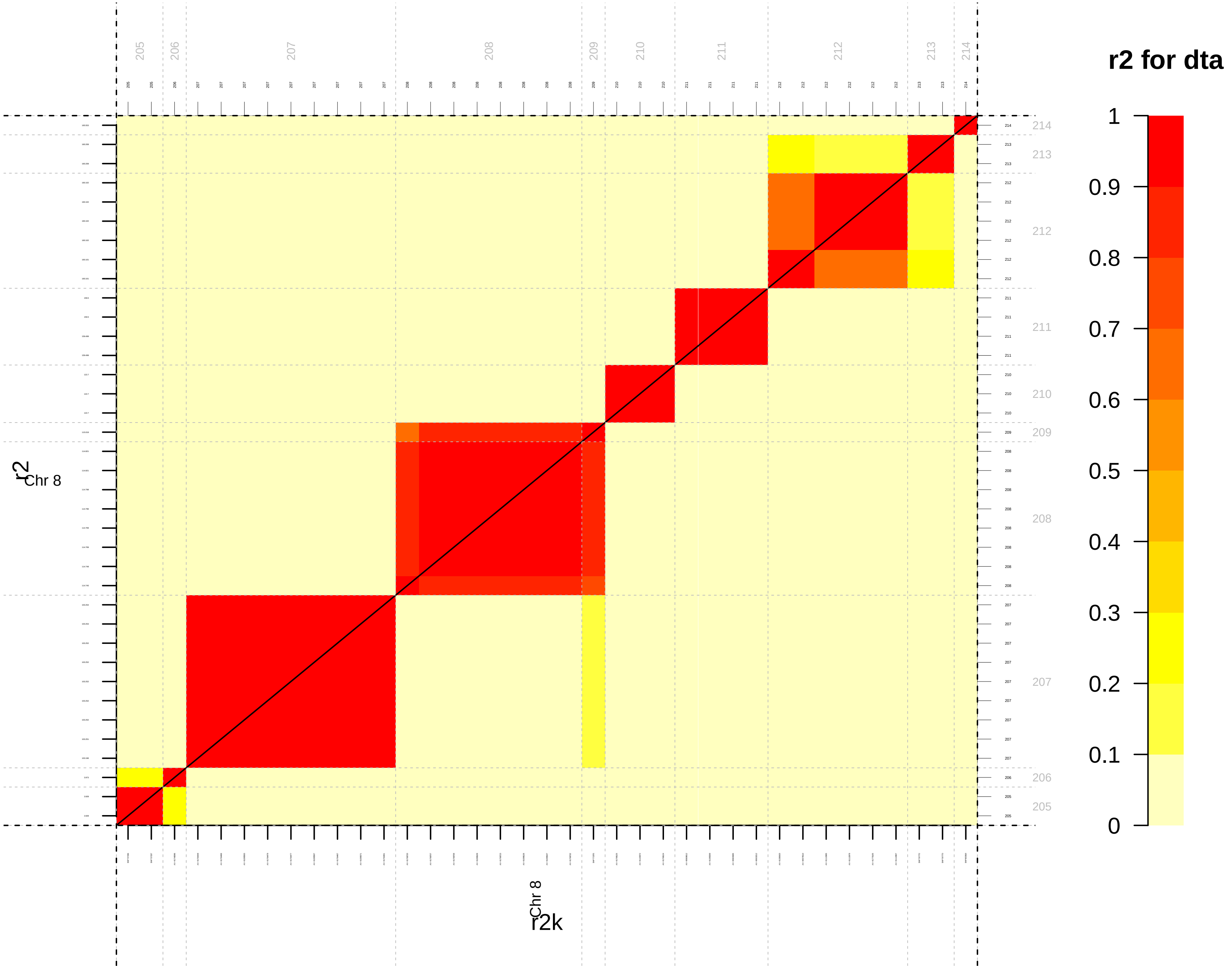
r2 for dta
1
0.9
0.8
0.7
0.6
0.5
0.4
0.3
0.2
0.1
0
r2
Chr 8
r2k
Chr 8

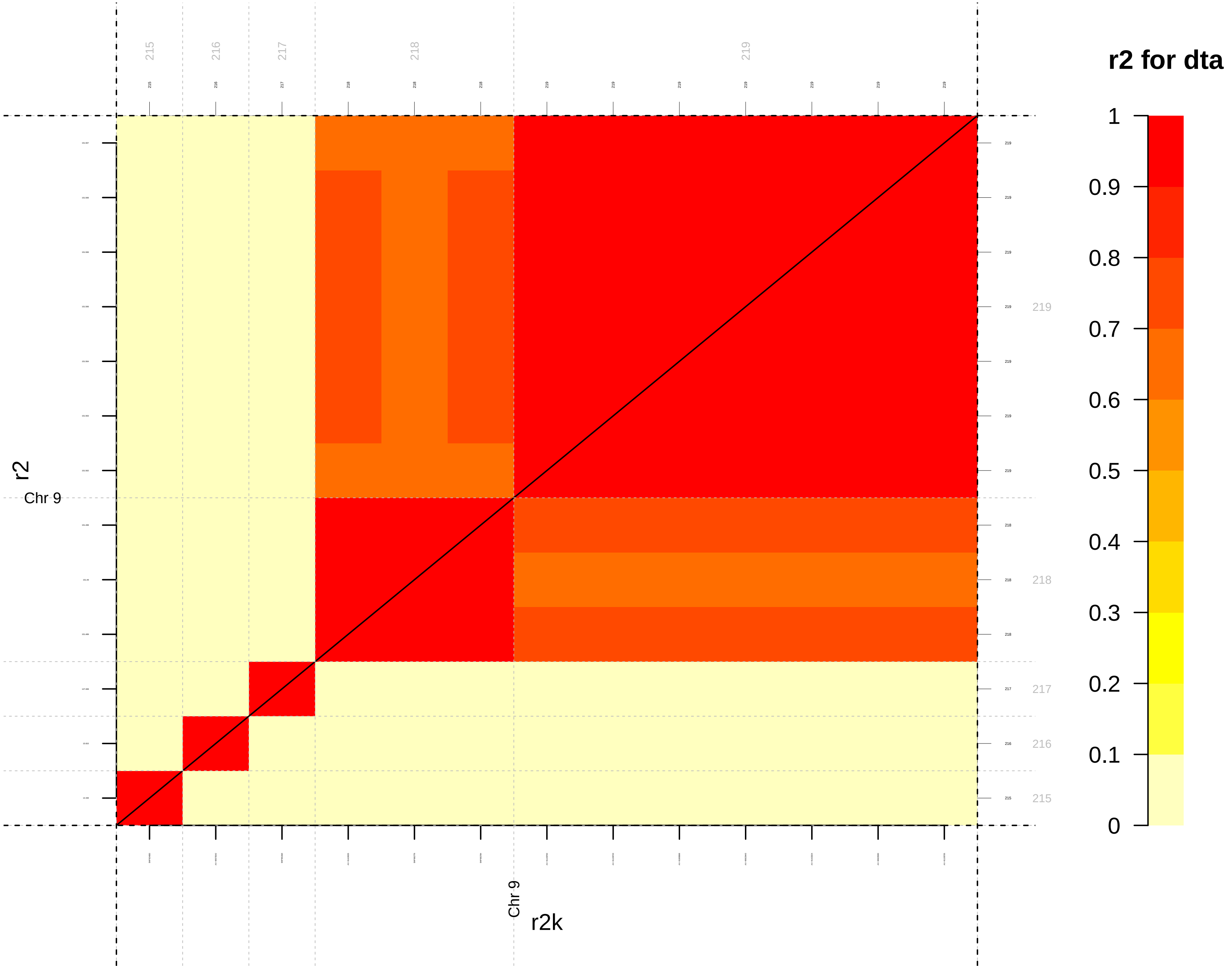
r2 for dta
1
0.9
0.8
0.7
0.6
0.5
0.4
0.3
0.2
0.1
0
215
216
217
218
219
r2
Chr 9
r2k
Chr 9

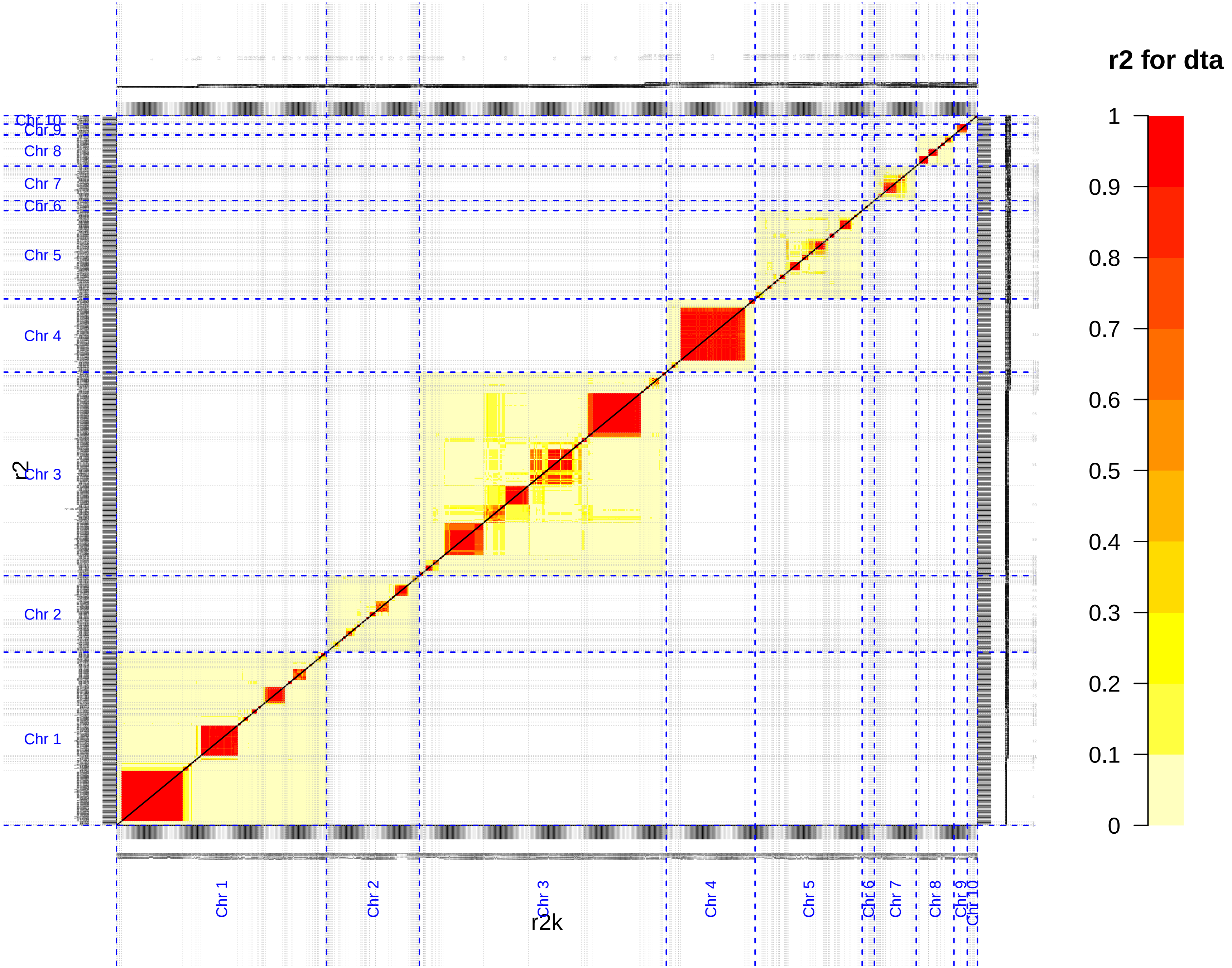

r2 for dta
1
0.9
0.8
0.7
0.6
0.5
0.4
0.3
0.2
0.1
0
Chr 10
Chr 9
Chr 8
Chr 7
Chr 6
Chr 5
Chr 4
Chr 3
Chr 2
Chr 1
r2
r2k

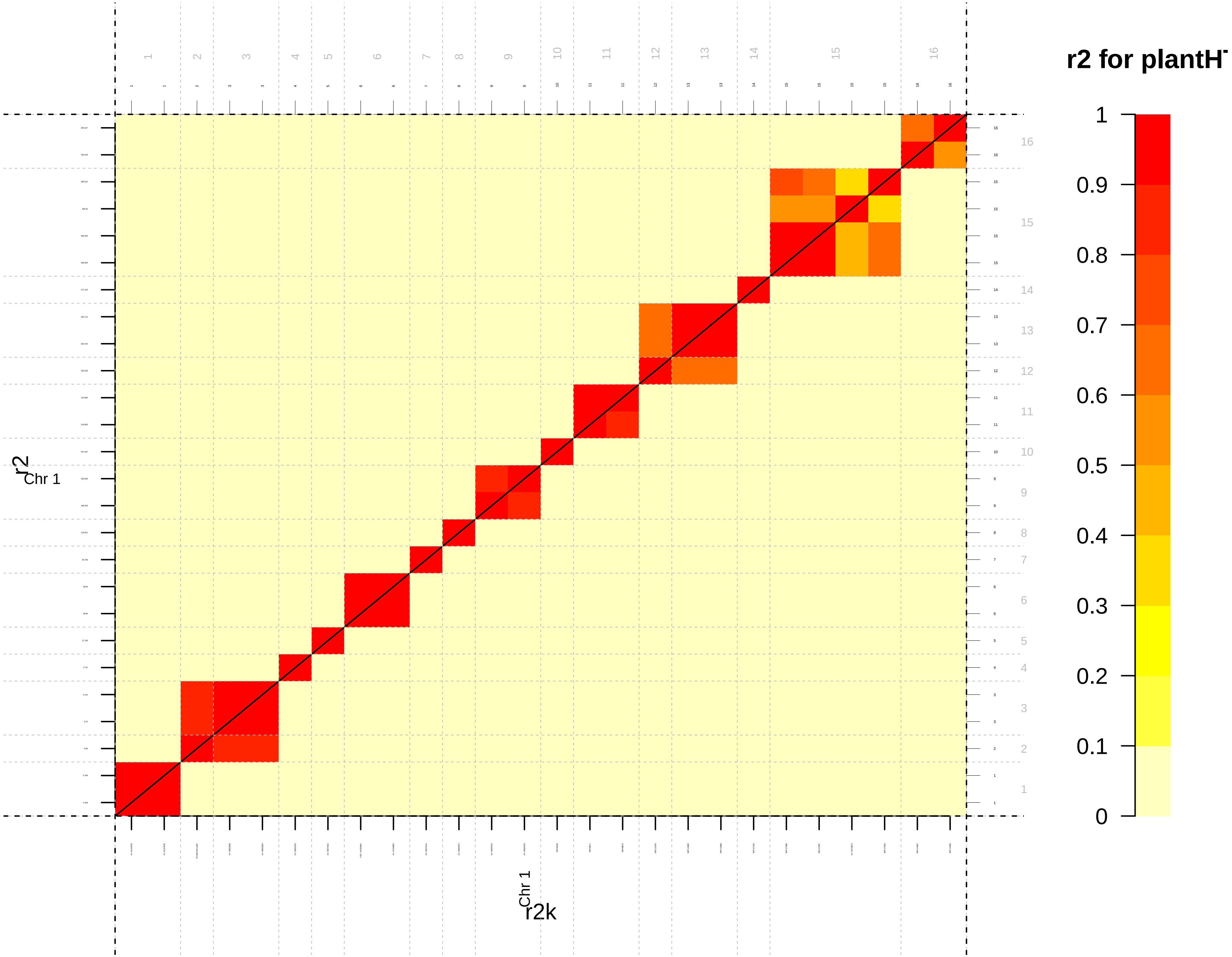
r2 for plantH
1
0.9
0.8
0.7
0.6
0.5
0.4
0.3
0.2
0.1
0
r2
Chr 1
r2k
Chr 1

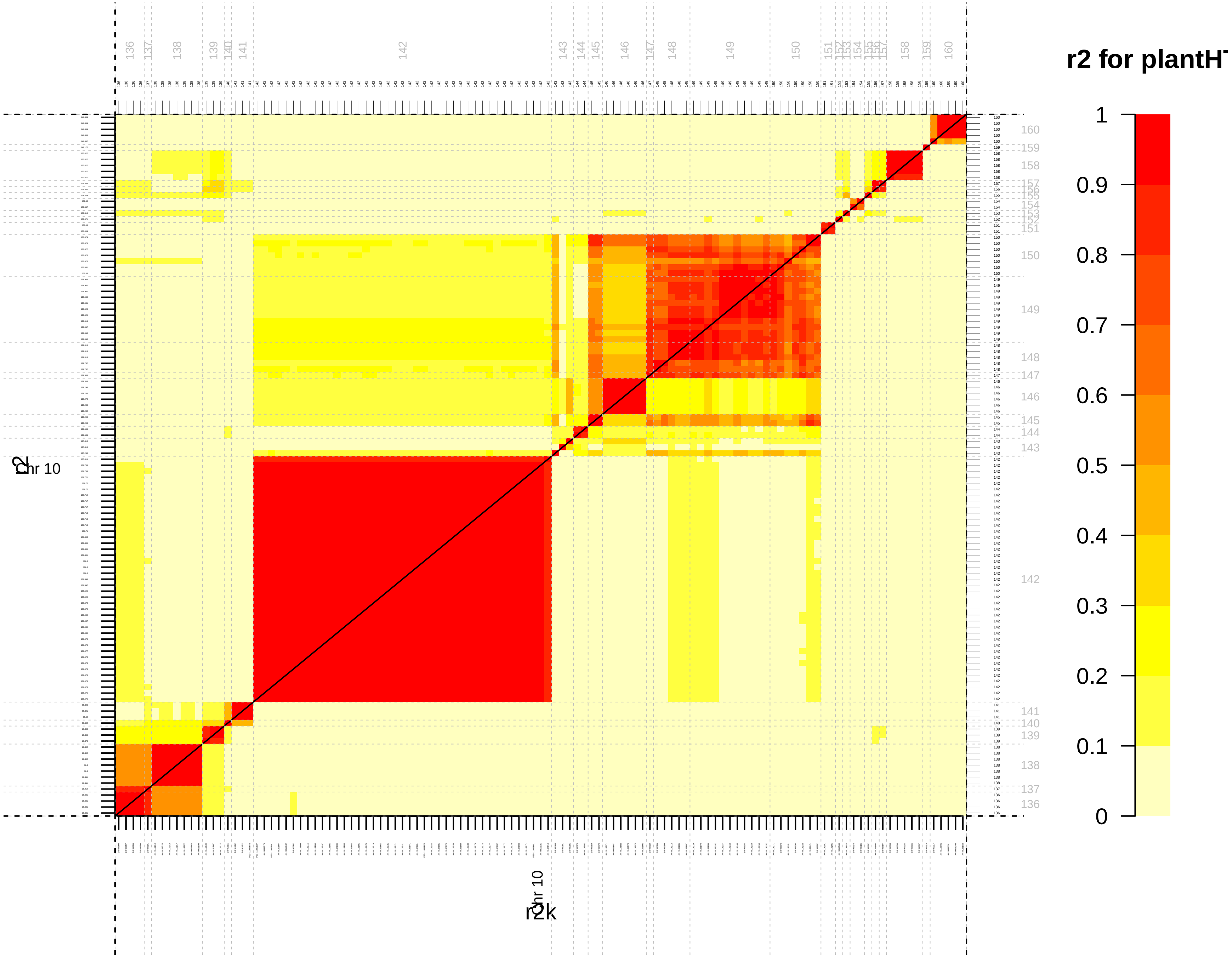

r2 for plantH
1
0.9
0.8
0.7
0.6
0.5
0.4
0.3
0.2
0.1
0
Chr 10
r2
r2k

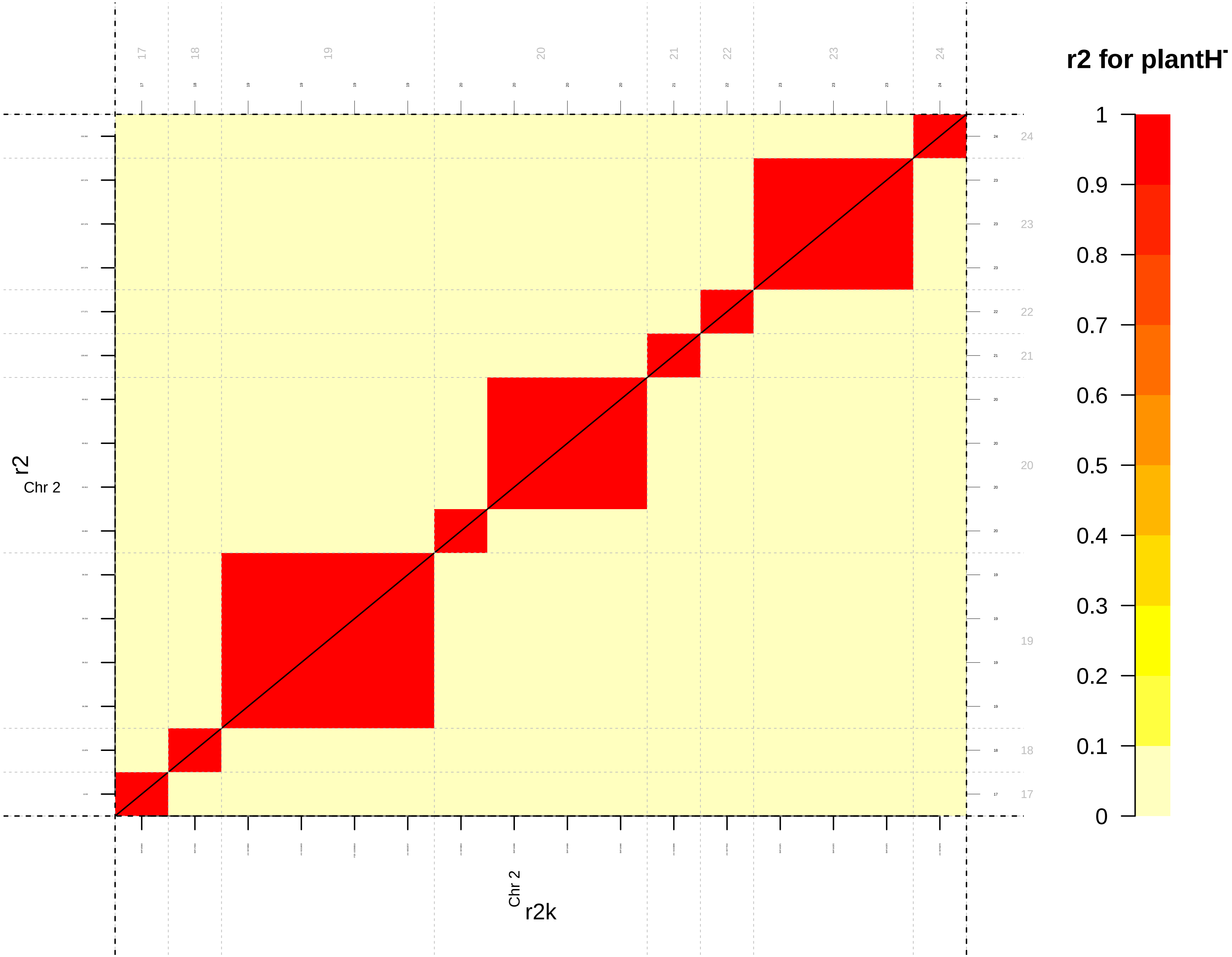
r2 for plantH
1
0.9
0.8
0.7
0.6
0.5
0.4
0.3
0.2
0.1
0
r2
Chr 2
r2k
Chr 2
17
18
19
20
21
22
23
24

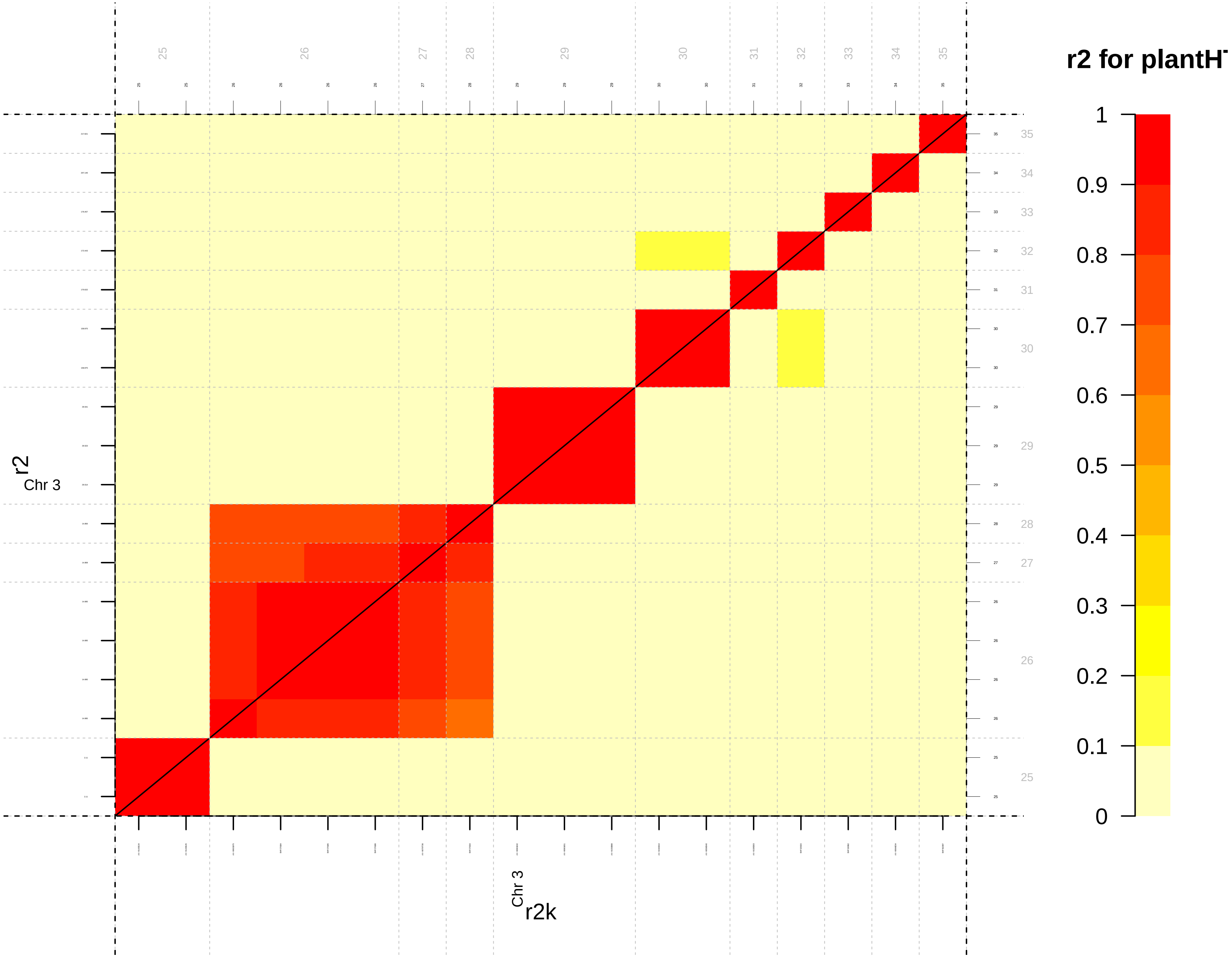

r2 for plantH
1
0.9
0.8
0.7
0.6
0.5
0.4
0.3
0.2
0.1
0
r2
Chr 3
r2k
Chr 3

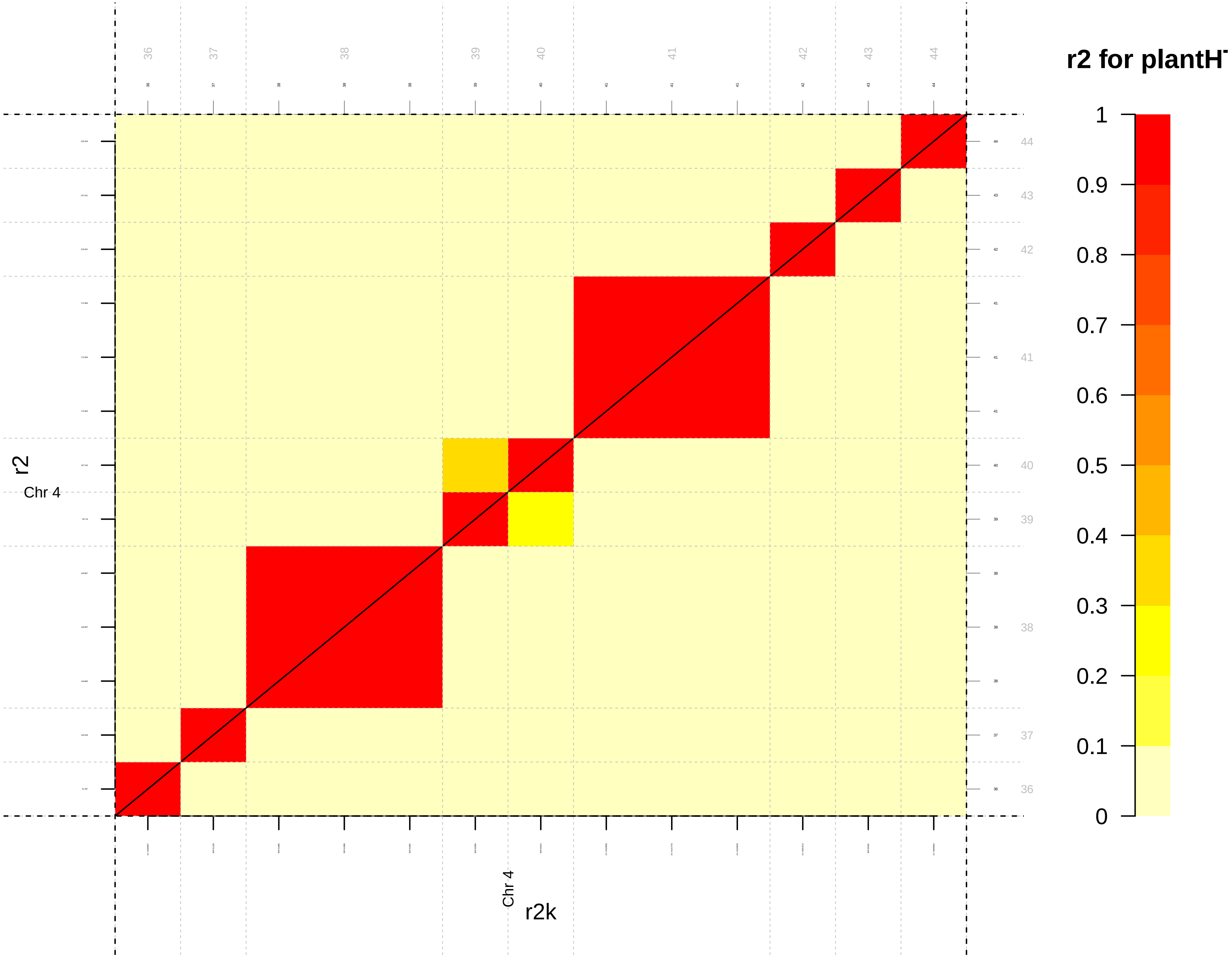

r2 for plantH
1
0.9
0.8
0.7
0.6
0.5
0.4
0.3
0.2
0.1
0
36
37
38
39
40
41
42
43
44
r2
Chr 4
r2k

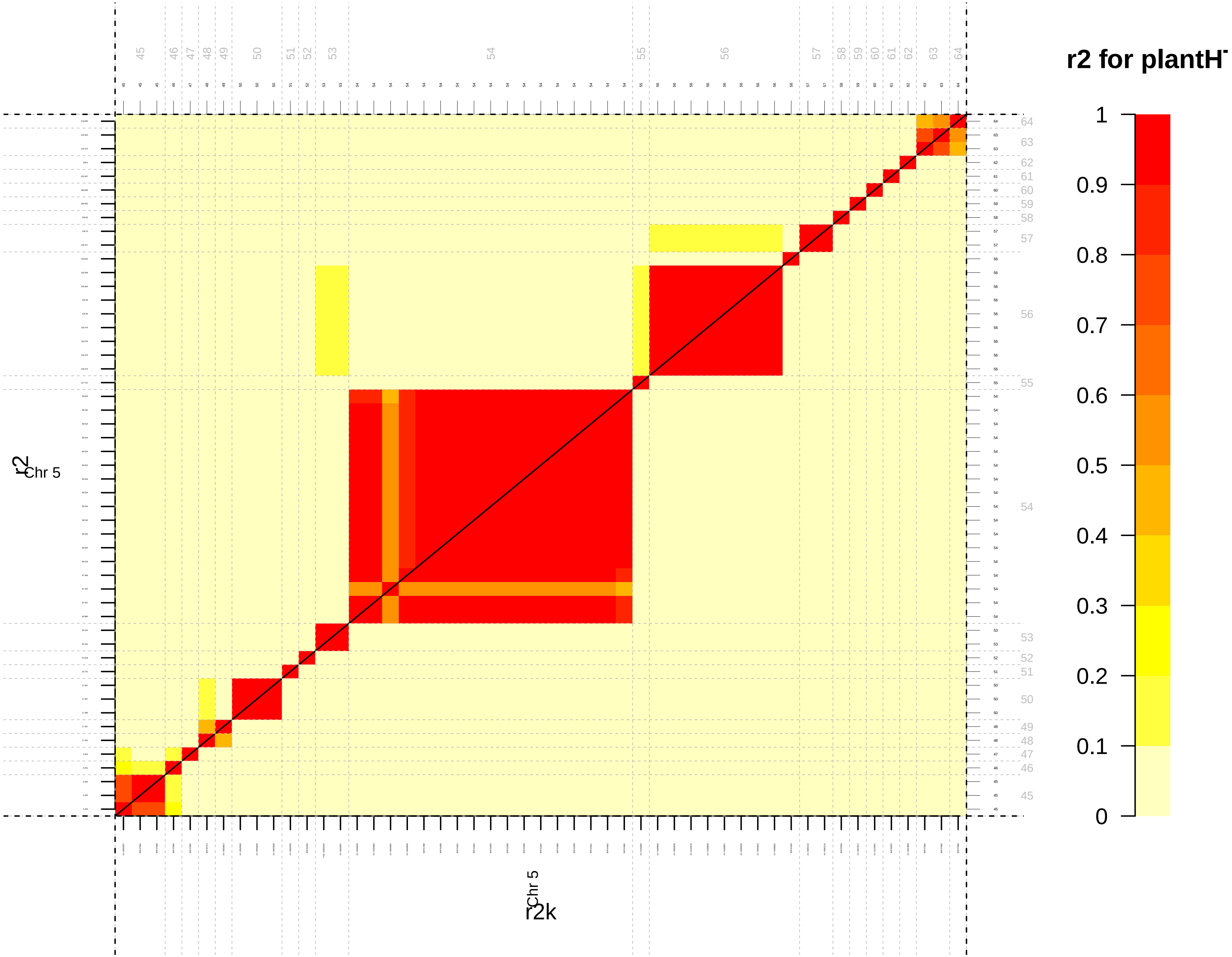
r2 for plantH
1
0.9
0.8
0.7
0.6
0.5
0.4
0.3
0.2
0.1
0
r2
Chr 5
r2k
Chr 5

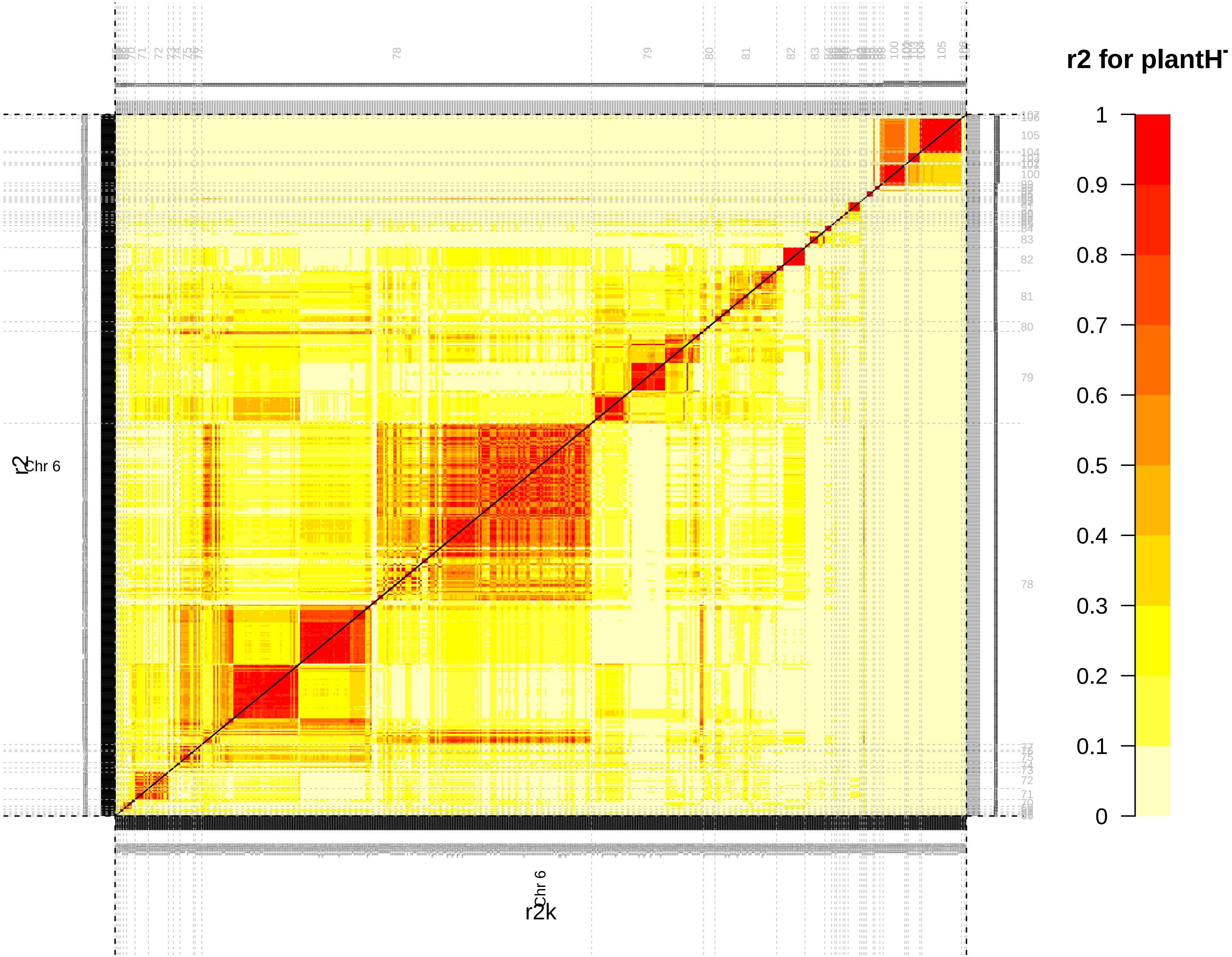
r2 for plantH
1
0.9
0.8
0.7
0.6
0.5
0.4
0.3
0.2
0.1
0
Chr 6
r2
Chr 6
r2k
78
79
80
81
82
83
100
105

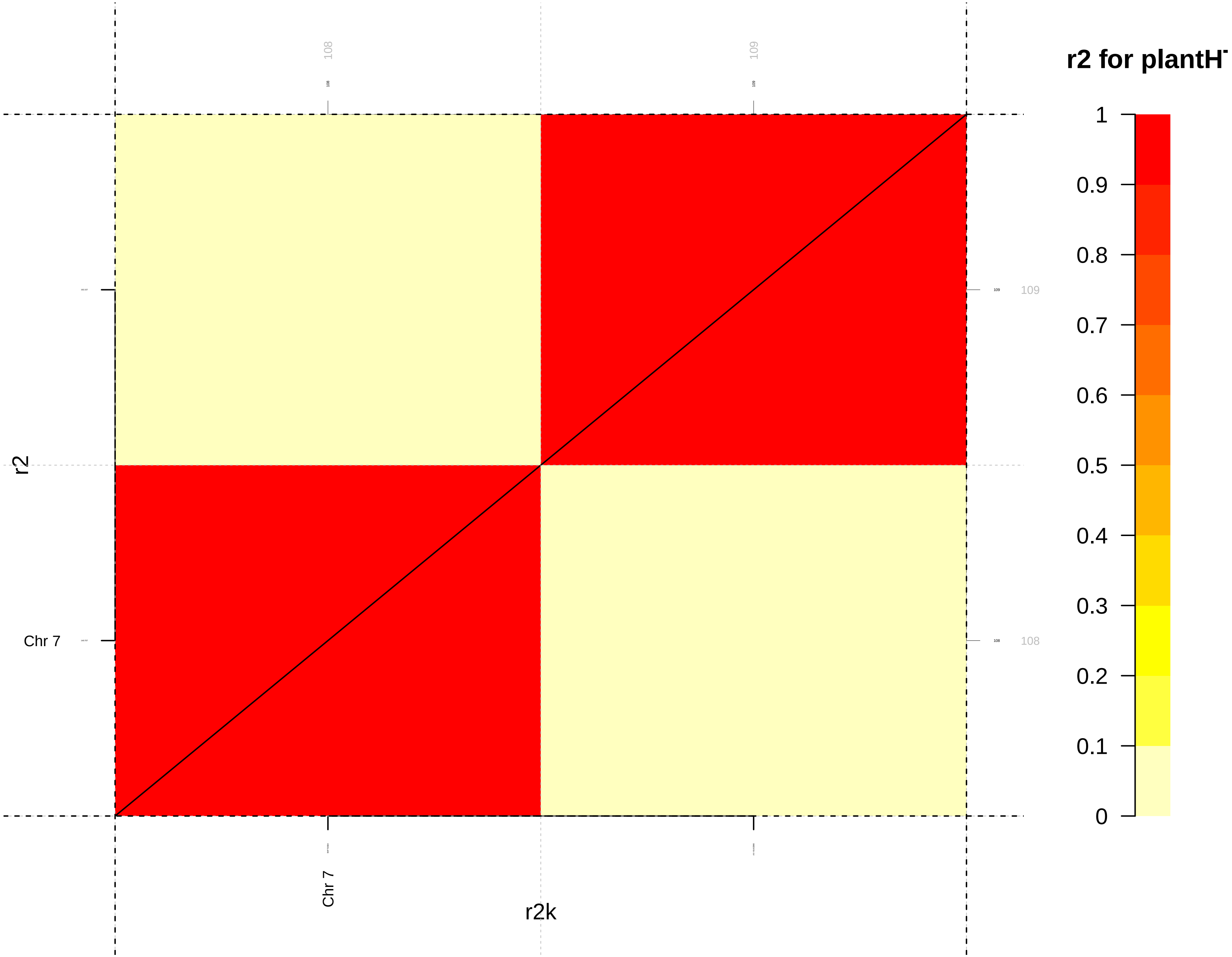

r2 for plantH
1
0.9
0.8
0.7
0.6
0.5
0.4
0.3
0.2
0.1
0
108
109
109
108
Chr 7
Chr 7
r2
r2k

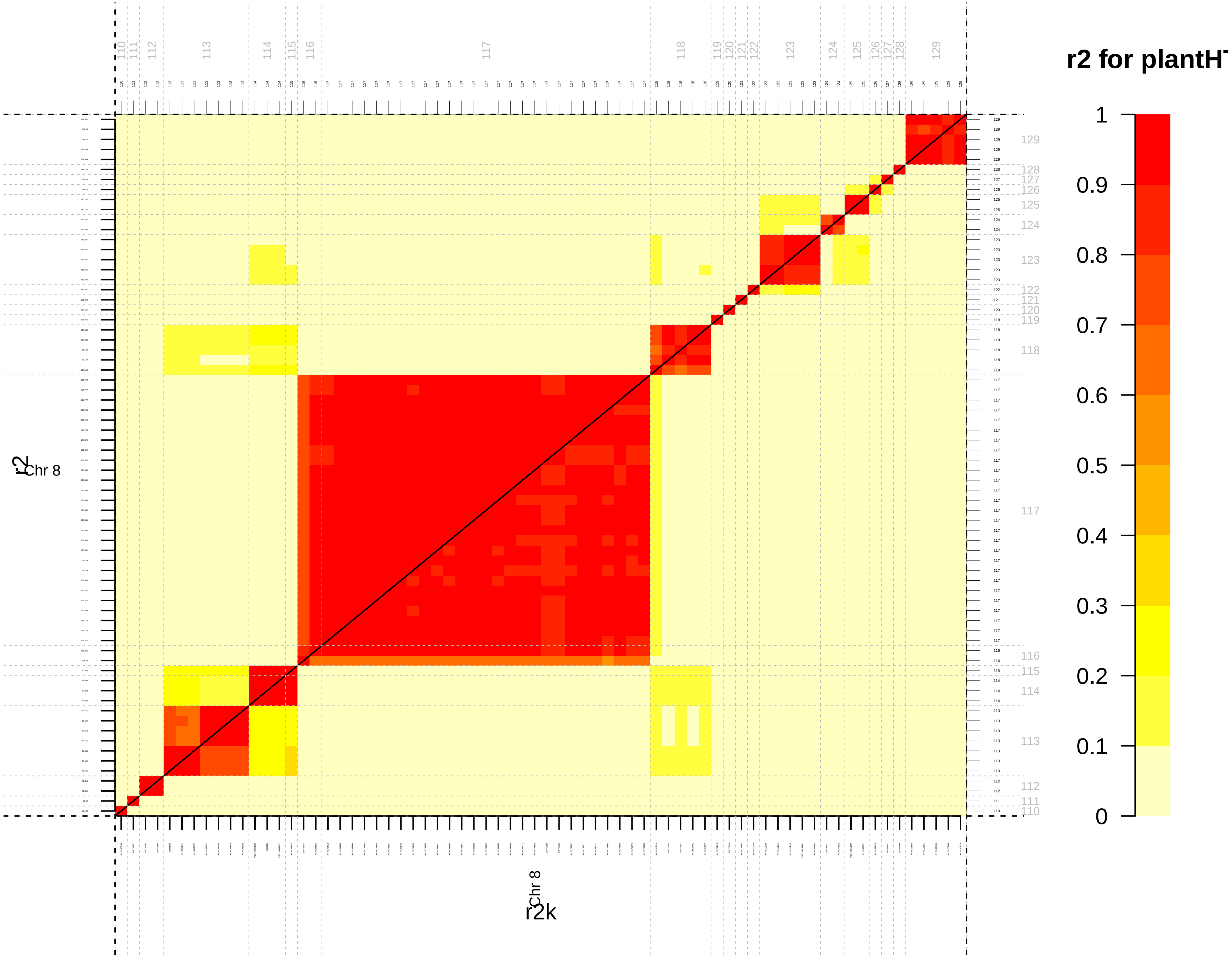

r2 for plantH
1
0.9
0.8
0.7
0.6
0.5
0.4
0.3
0.2
0.1
0
Chr 8
r2
Chr 8
r2k

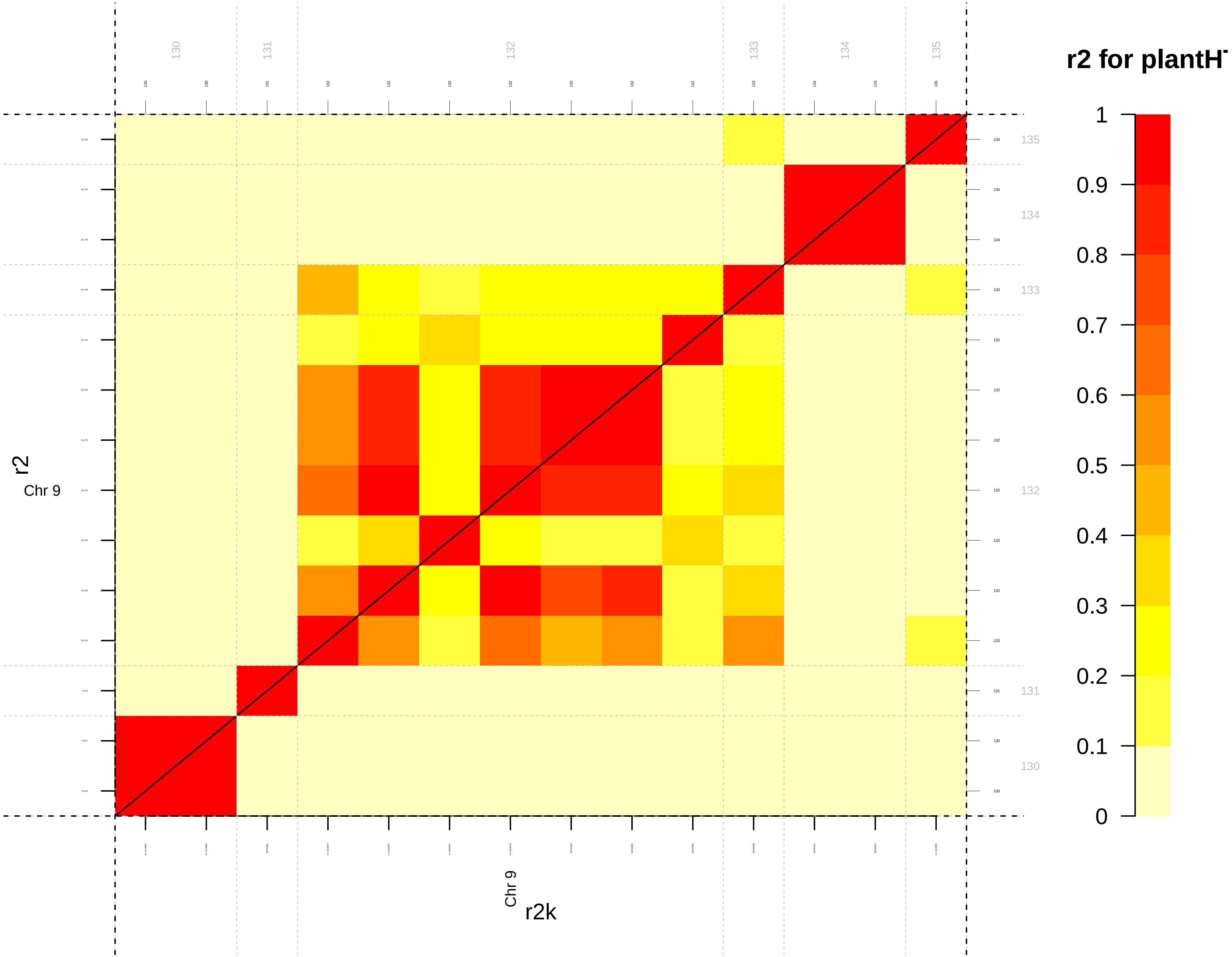

r2 for plantH
1
0.9
0.8
0.7
0.6
0.5
0.4
0.3
0.2
0.1
0
130
131
132
133
134
135
r2
Chr 9
Chr 9
r2k

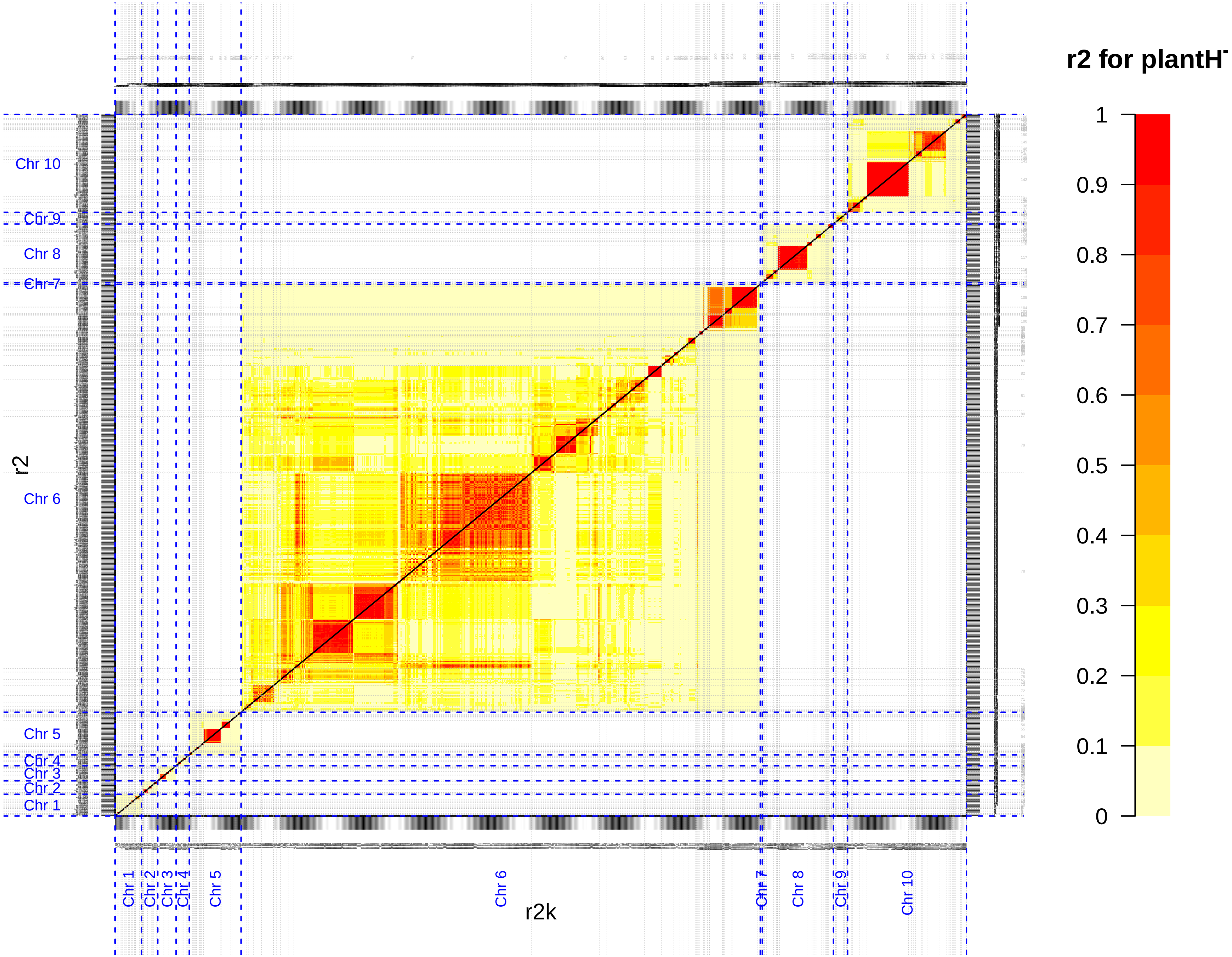

r2 for plantH
1
0.9
0.8
0.7
0.6
0.5
0.4
0.3
0.2
0.1
0
r2
Chr 10
Chr 9
Chr 8
Chr 7
Chr 6
Chr 5
Chr 4
Chr 3
Chr 2
Chr 1
r2k

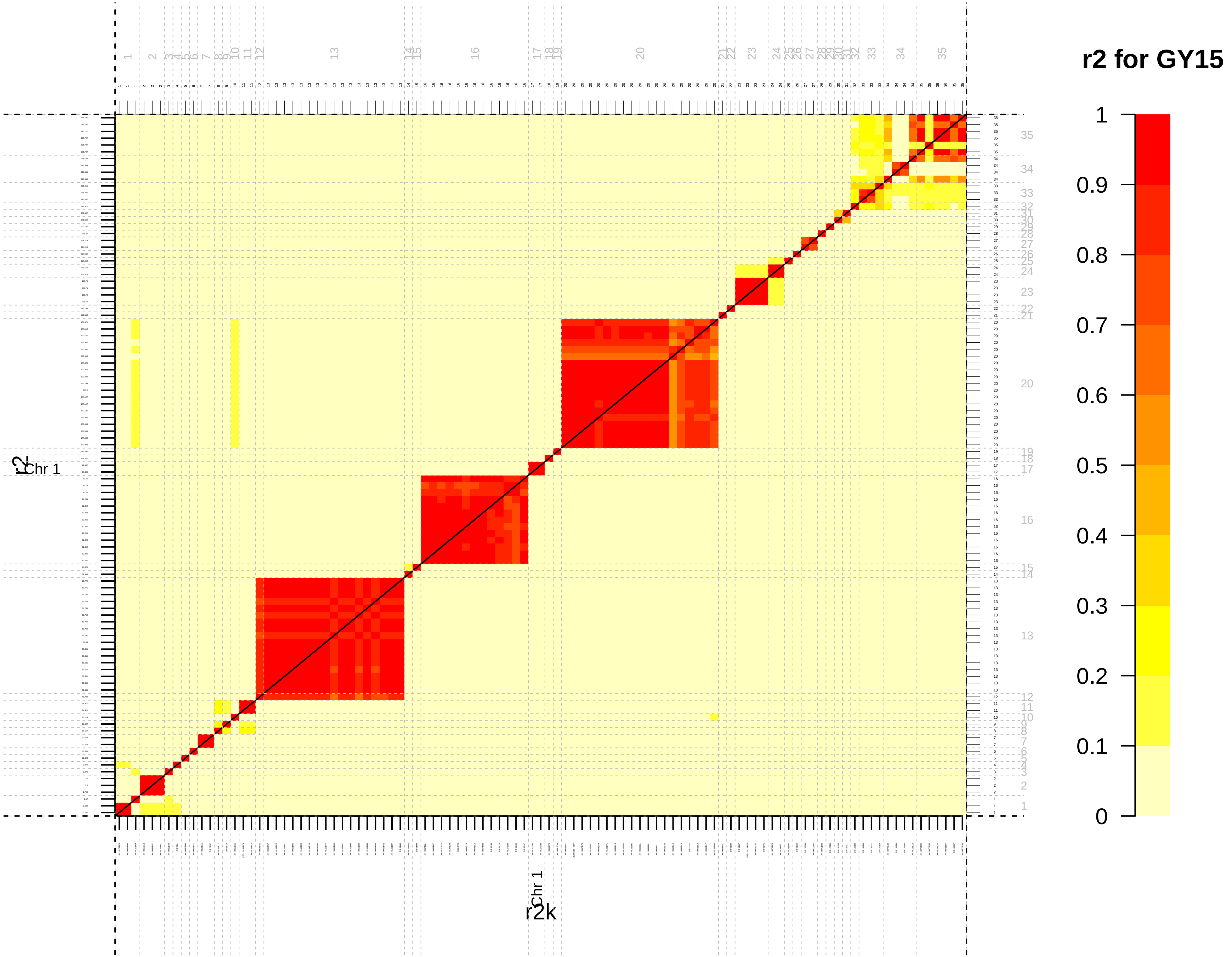
r2 for GY15
1
0.9
0.8
0.7
0.6
0.5
0.4
0.3
0.2
0.1
0
Chr 1
r2
Chr 1
r2k

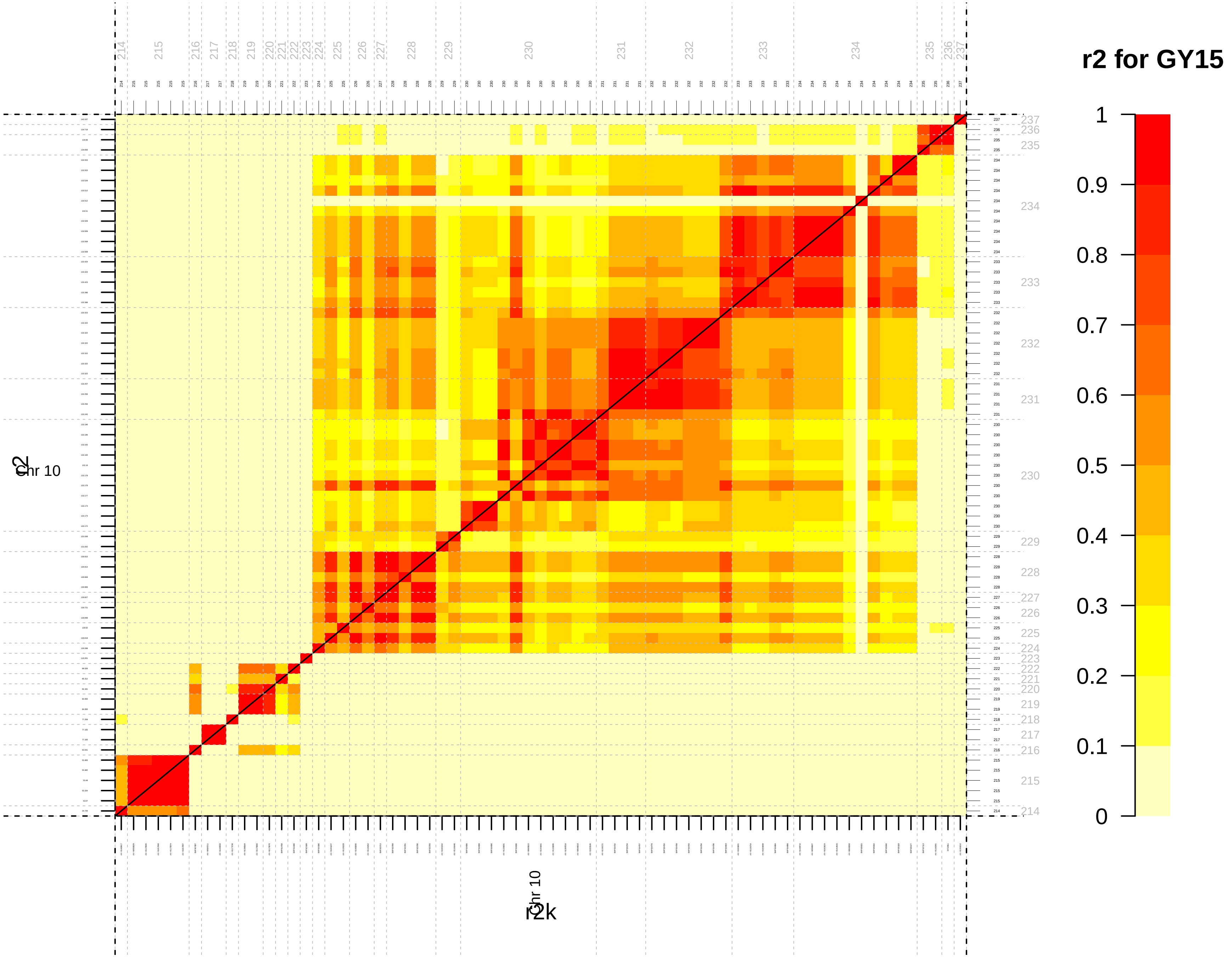
r2 for GY15
1
0.9
0.8
0.7
0.6
0.5
0.4
0.3
0.2
0.1
0
Chr 10
r2
Chr 10
r2k

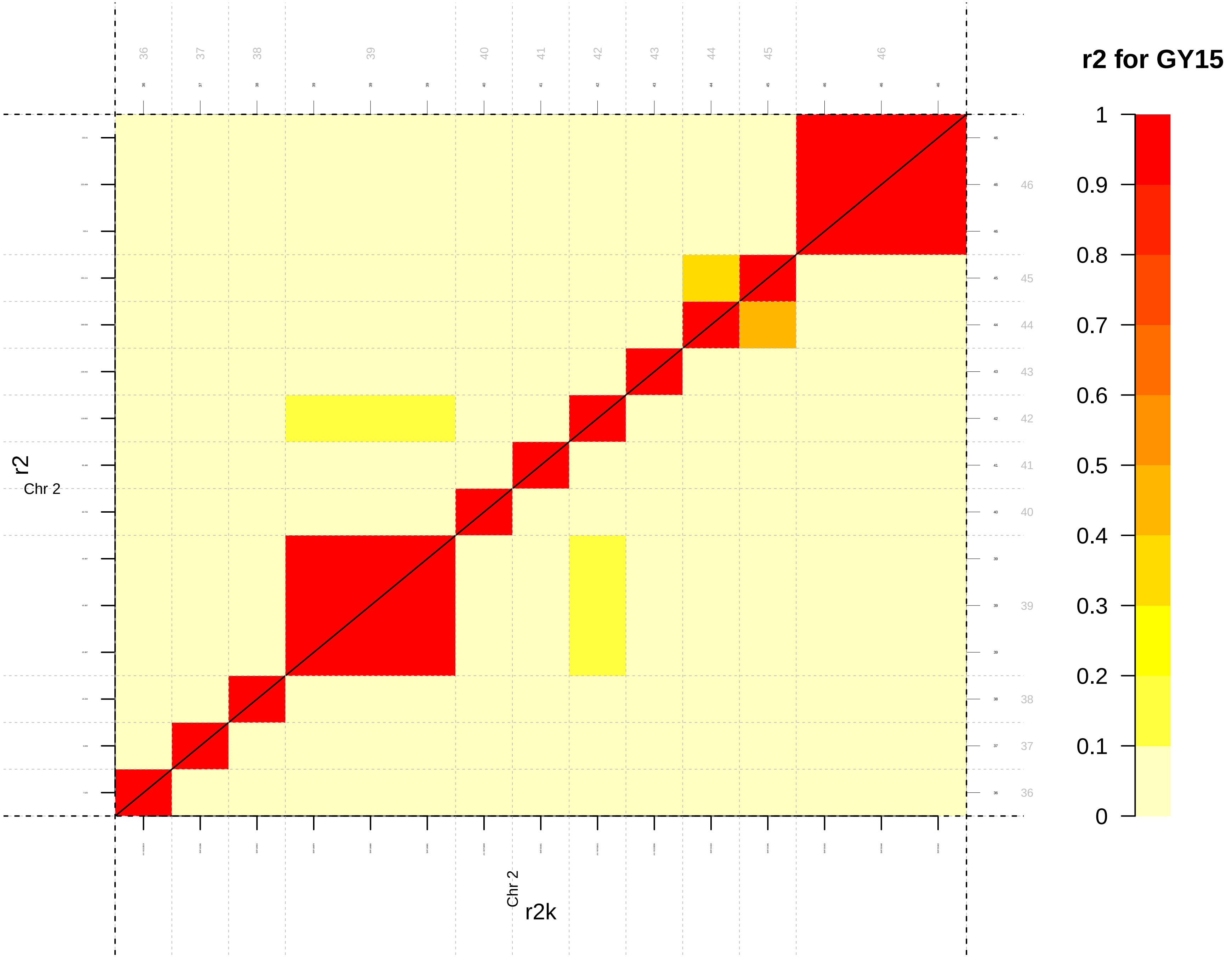

r2 for GY15
1
0.9
0.8
0.7
0.6
0.5
0.4
0.3
0.2
0.1
0
r2
Chr 2
r2k

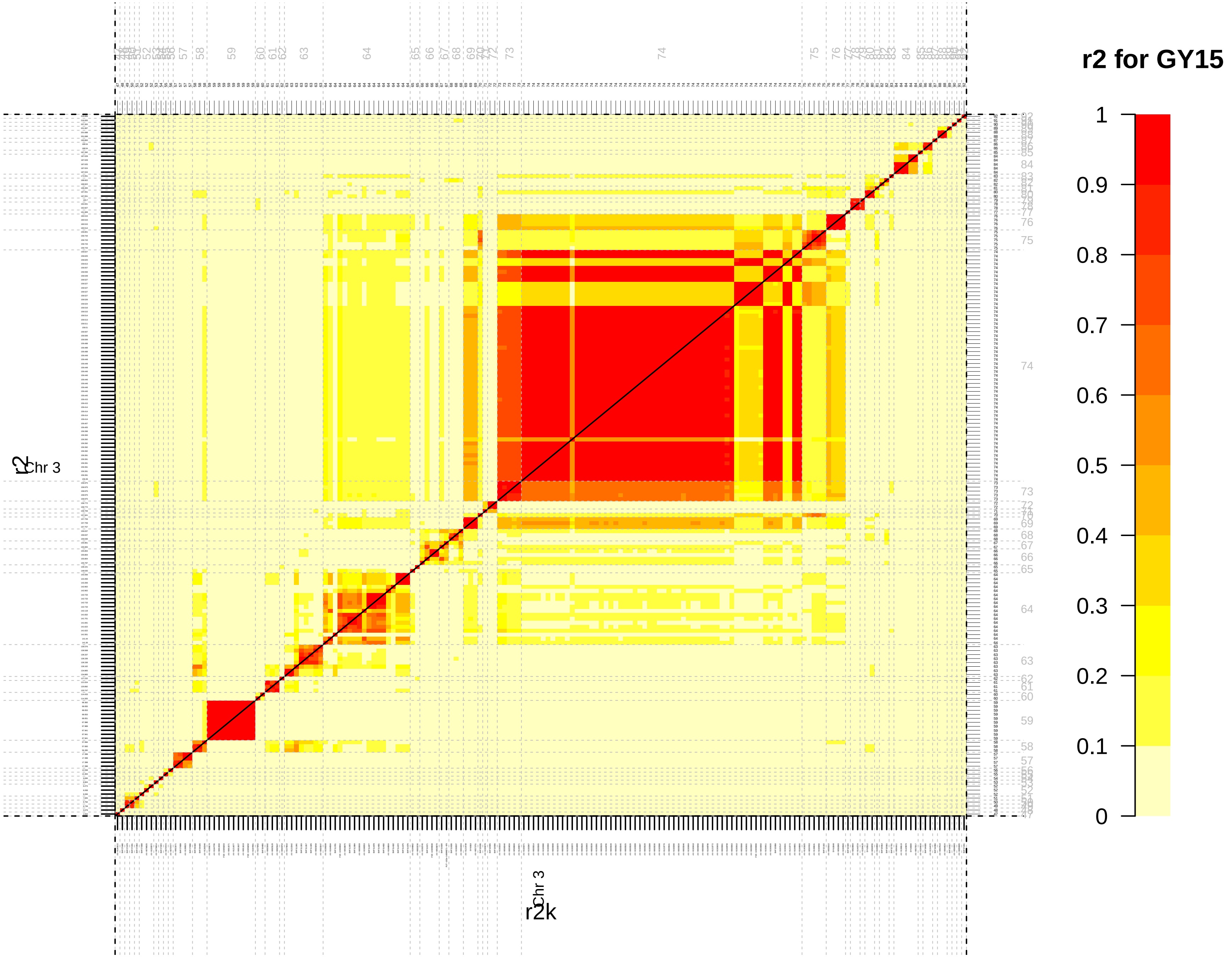
r2 for GY15
1
0.9
0.8
0.7
0.6
0.5
0.4
0.3
0.2
0.1
0
Chr 3
r2
r2k

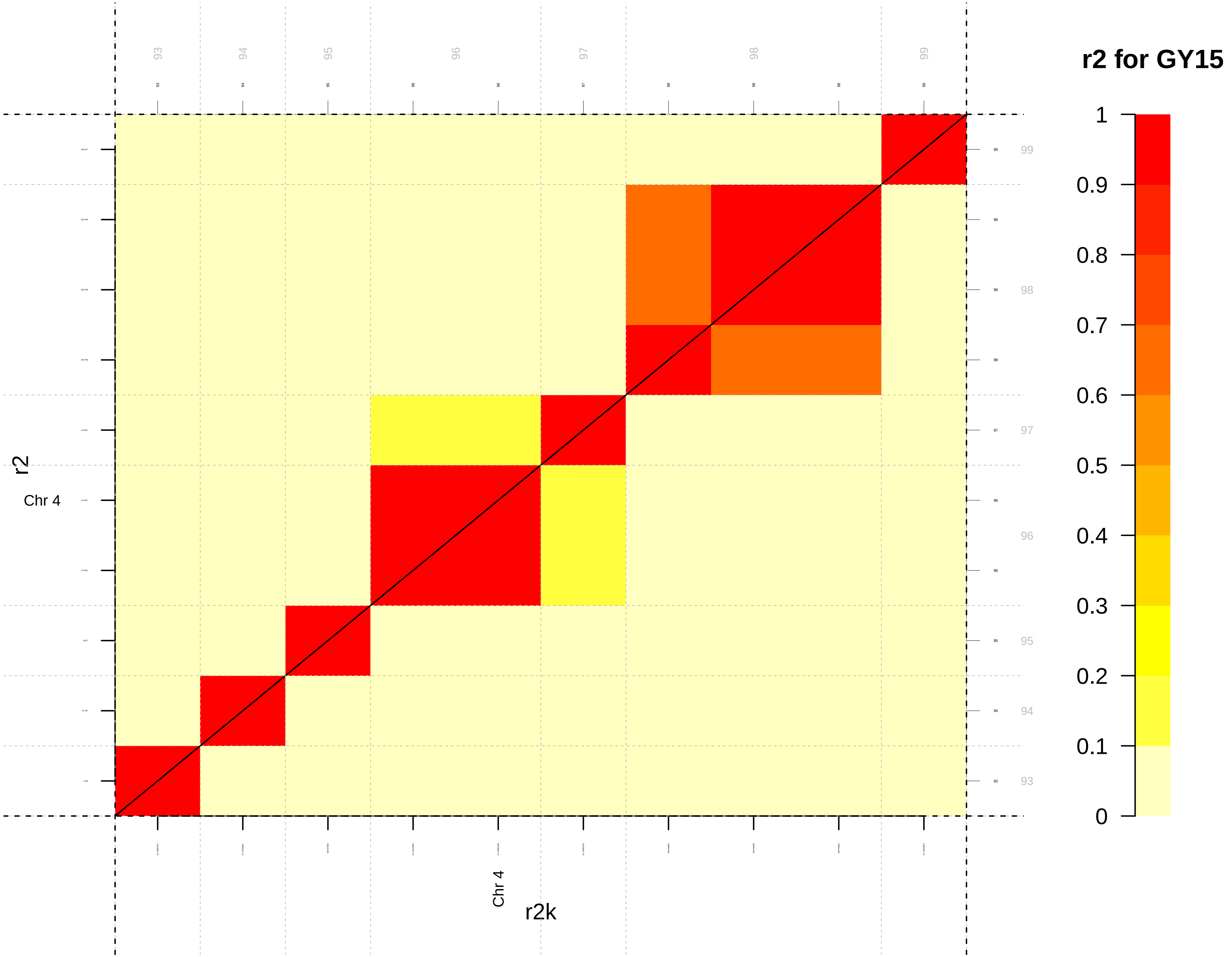

r2 for GY15
1
0.9
0.8
0.7
0.6
0.5
0.4
0.3
0.2
0.1
0
93
94
95
96
97
98
99
r2
Chr 4
r2k

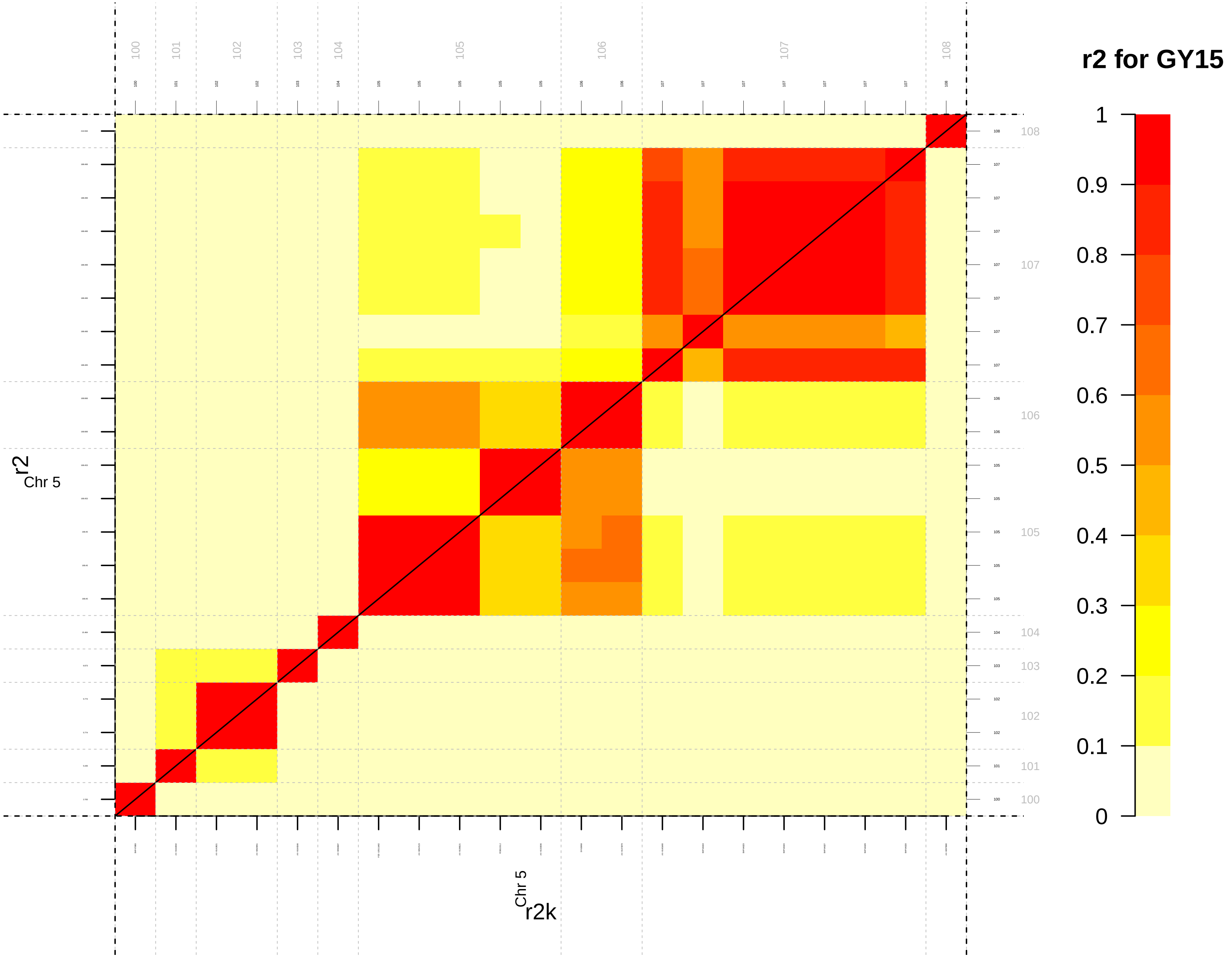

r2 for GY15
1
0.9
0.8
0.7
0.6
0.5
0.4
0.3
0.2
0.1
0
r2
Chr 5
r2k
Chr 5
100
101
102
103
104
105
106
107
108

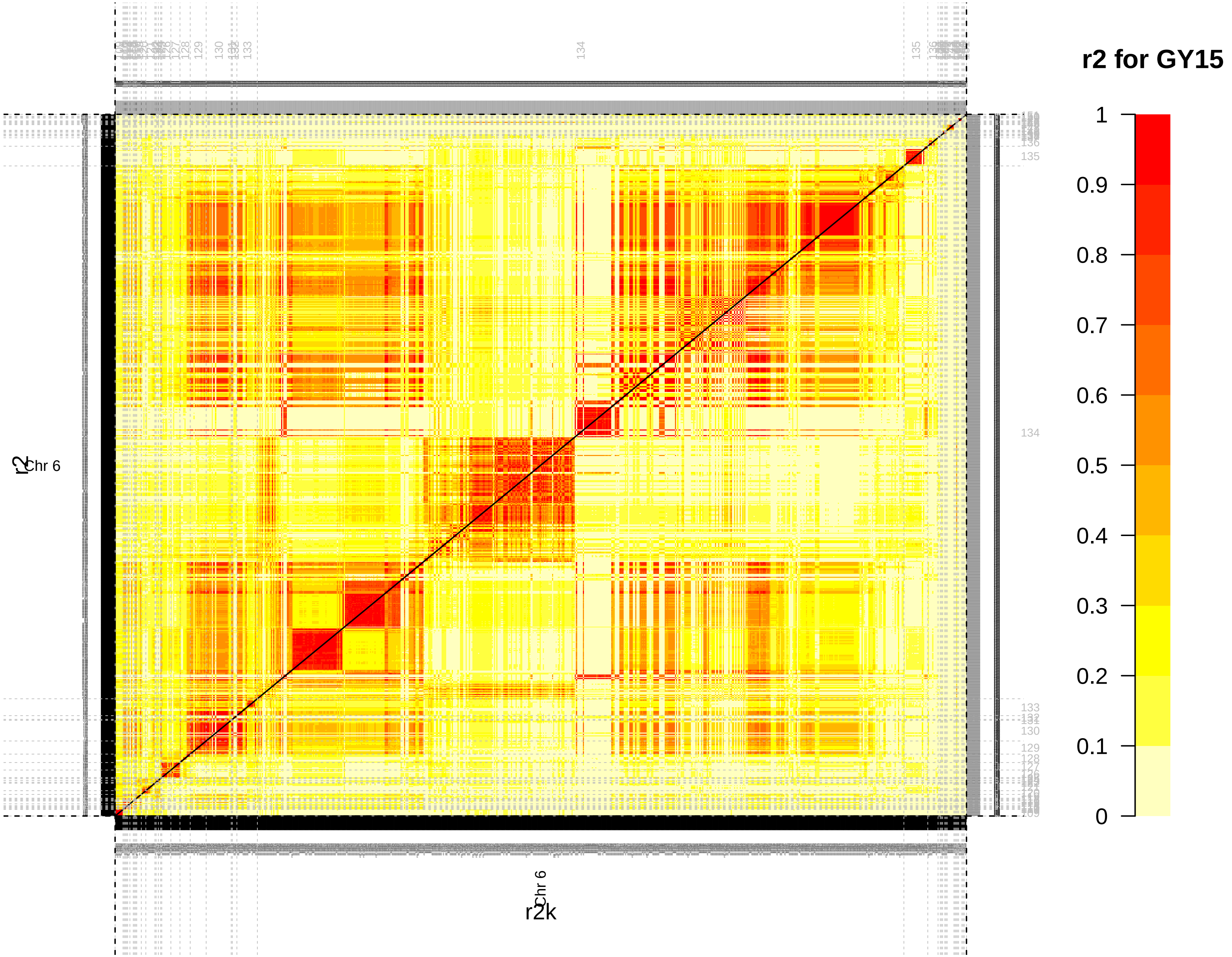

r2 for GY15
1
0.9
0.8
0.7
0.6
0.5
0.4
0.3
0.2
0.1
0
Chr 6
r2
r2k
134
135
130
133

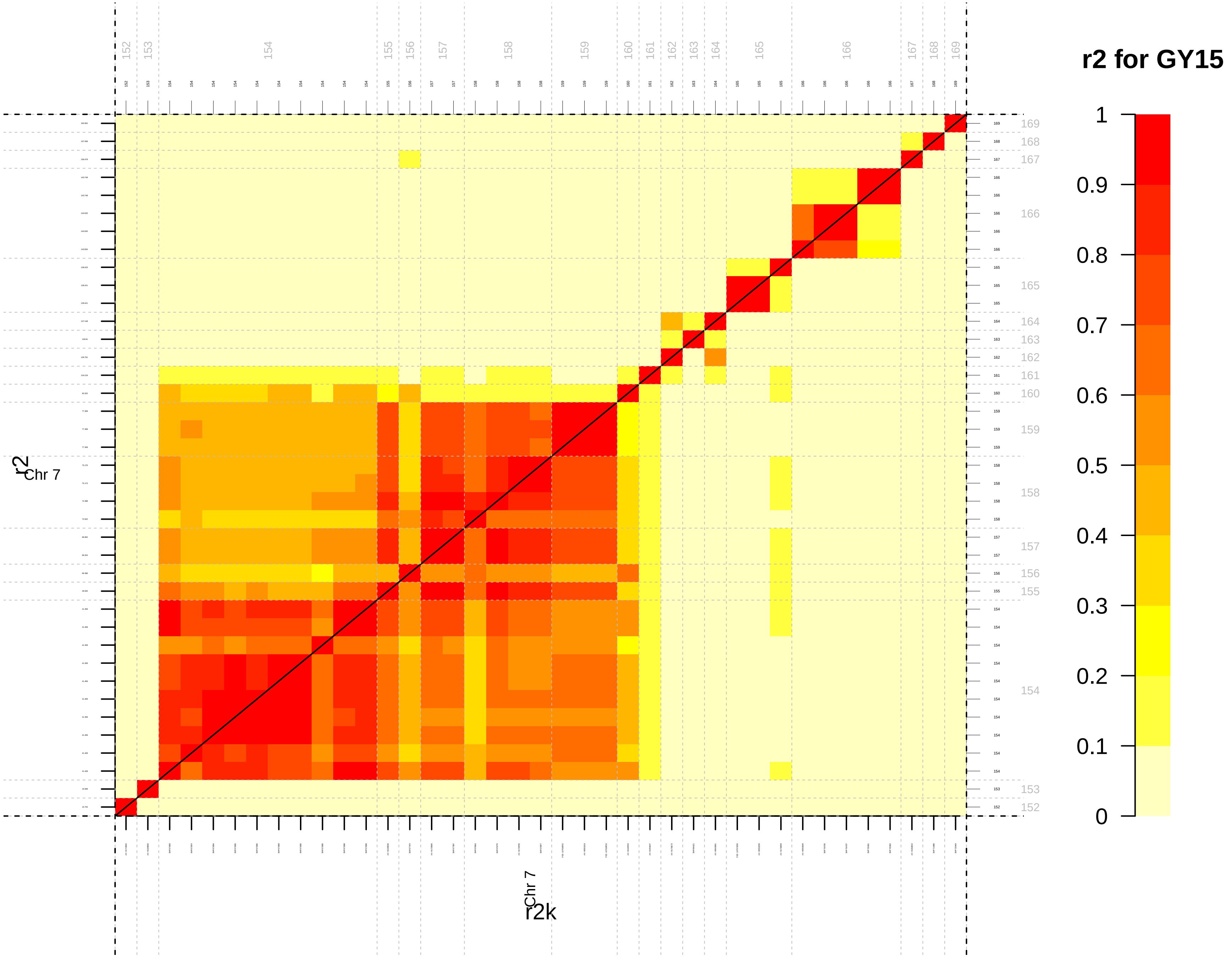

r2 for GY15
1
0.9
0.8
0.7
0.6
0.5
0.4
0.3
0.2
0.1
0
r2
Chr 7
r2k
Chr 7

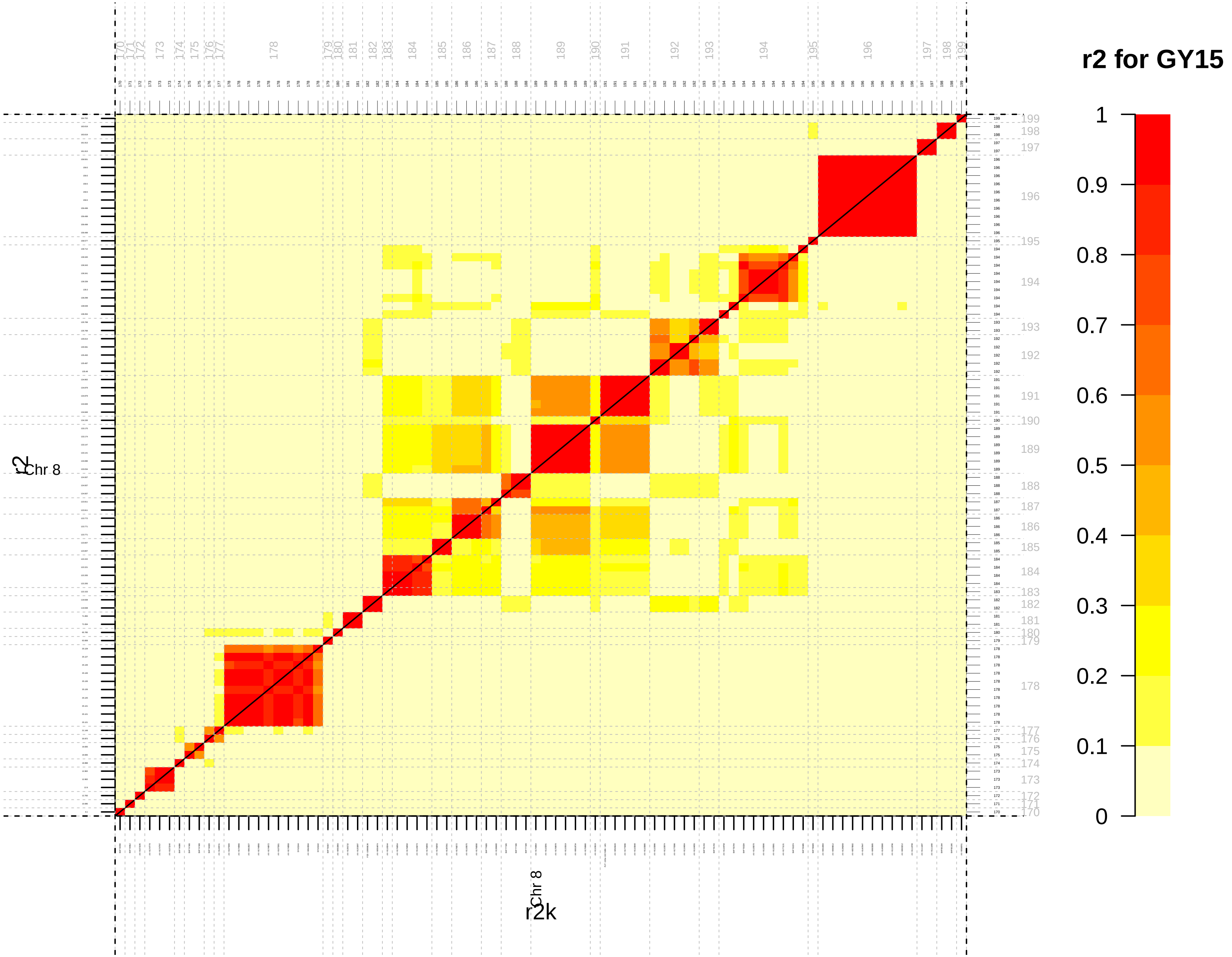

r2 for GY15
1
0.9
0.8
0.7
0.6
0.5
0.4
0.3
0.2
0.1
0
r2
Chr 8
Chr 8
r2k

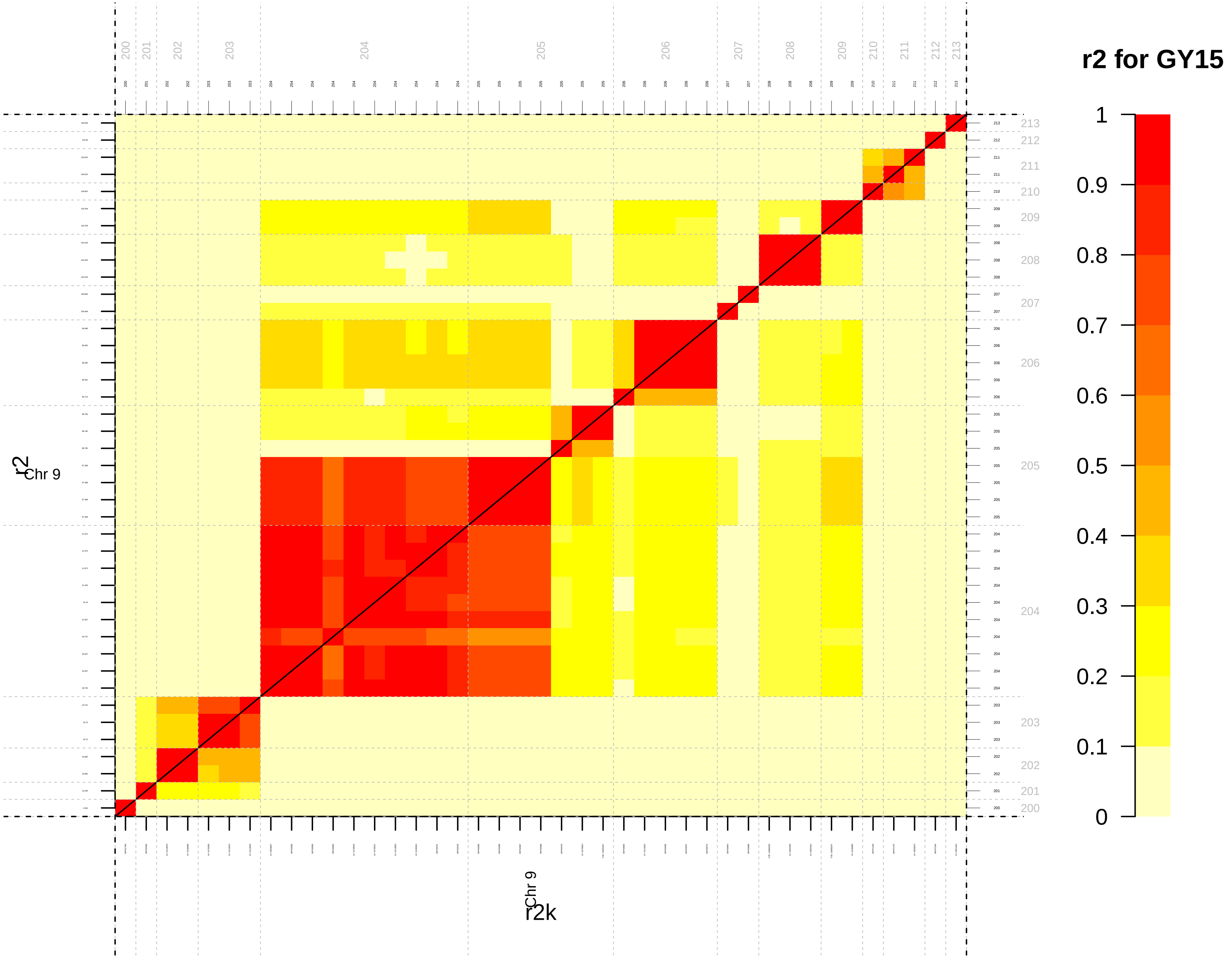
r2 for GY15
1
0.9
0.8
0.7
0.6
0.5
0.4
0.3
0.2
0.1
0
Chr 9
r2
Chr 9
r2k

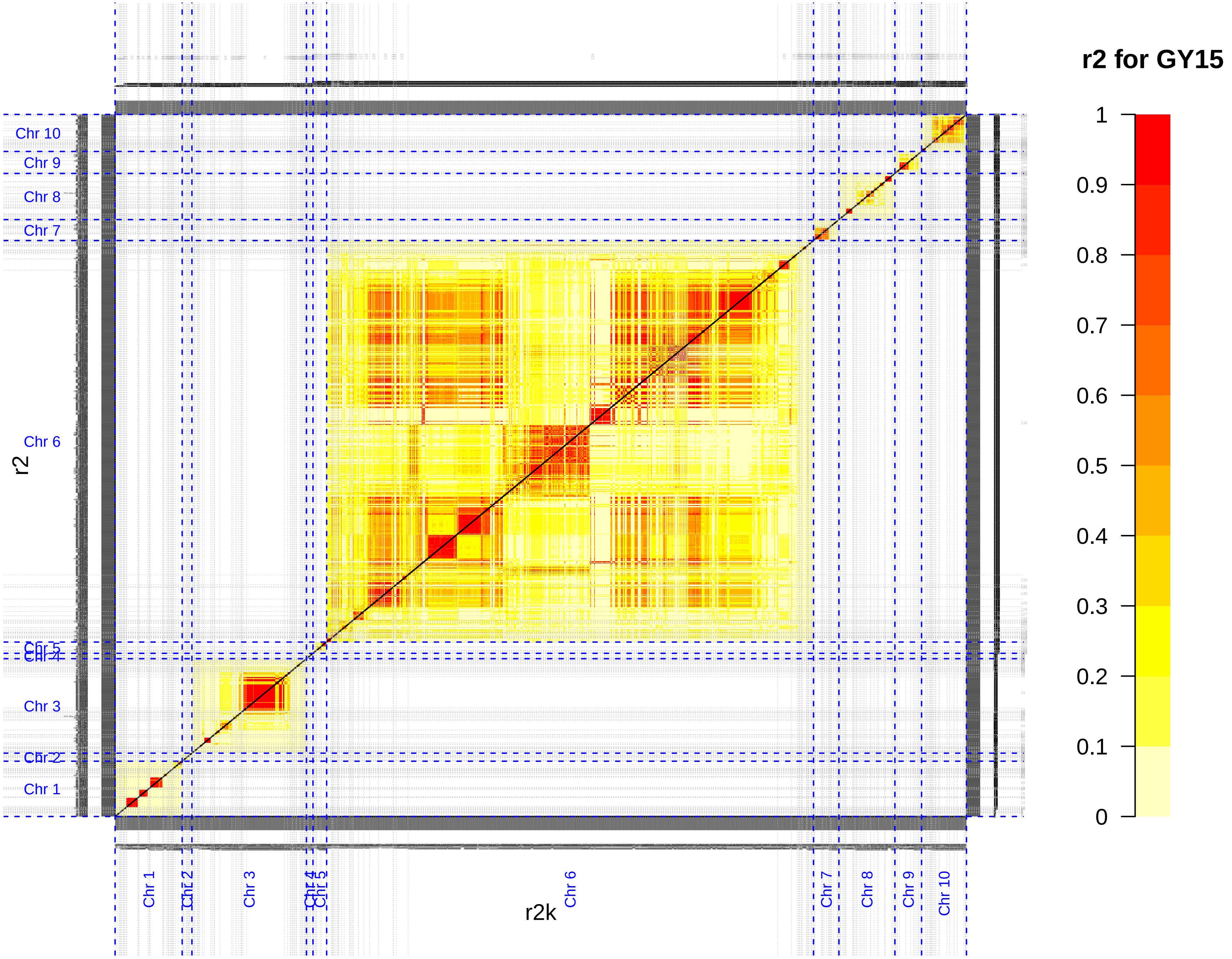

r2 for GY15
1
0.9
0.8
0.7
0.6
0.5
0.4
0.3
0.2
0.1
0
Chr 10
Chr 9
Chr 8
Chr 7
Chr 6
Chr 5
Chr 4
Chr 3
Chr 2
Chr 1
r2
r2k

**Additional file 8 (.pdf):**

**Figure S8-LD_Windows:** QTL limits obtained by the *LD_win* approach projected on heatmaps representing the level of LD between associated SNPs for each trait (DTA: male flowering time, plantHT: plant height and GY: grain yield) and each chromosome. Upper and lower triangles on the heatmaps represented the $r^2$ and $r^2K$ values between associated SNPs, respectively. Linkage disequilibrium between loci was colored according to values from weak LD (yellow) to high LD (red). The significant markers were ordered according to their physical positions on the chromosome and were represented by ticks on the four sides of the heatmaps. Limits of QTLs were displayed by gray dotted lines. QTL numbers were indicated in gray on the top and the right of each heatmap.
