## Additional file 9: Fig S9 for "Genotyping-by-sequencing and SNP-arrays are complementary for detecting quantitative trait loci by tagging different haplotypes in association studies"

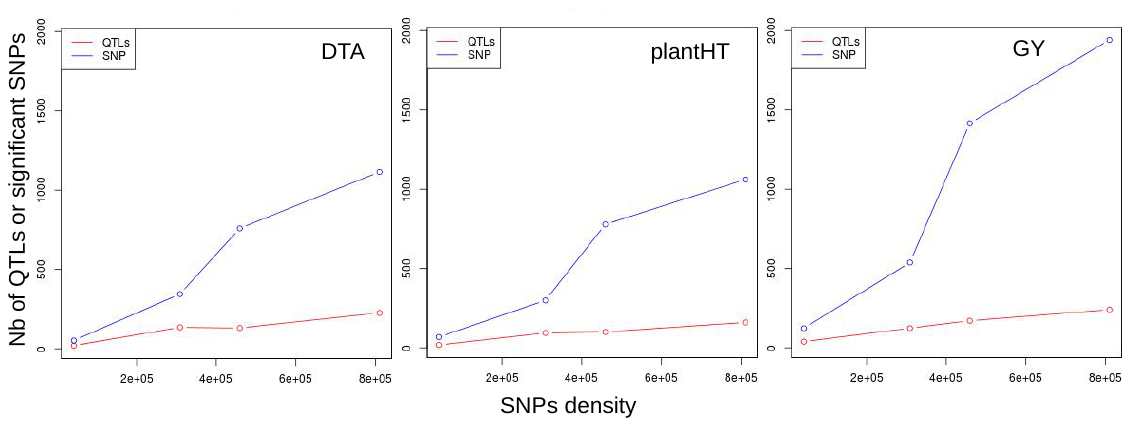

Figure S9: Number of significant SNPs (blue line) and QTLs (red line) identified as a function of SNP density (x-axis) for the three traits (DTA, male flowering time; plantHT, plant height; GY, grain yield).
