## Additional file 11: Fig S10 for "Genotyping-by-sequencing and SNP-arrays are complementary for detecting quantitative trait loci by tagging different haplotypes in association studies"

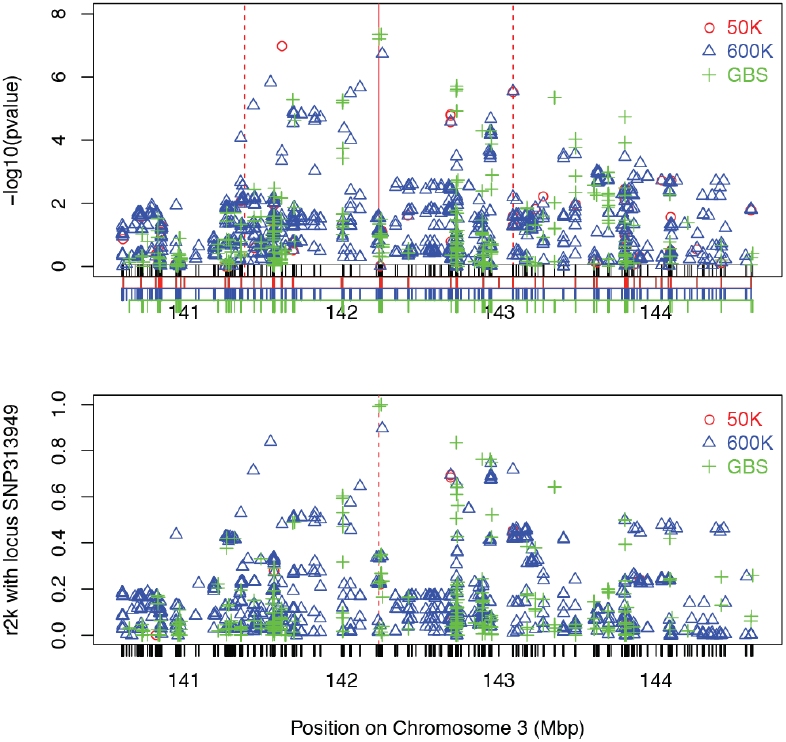

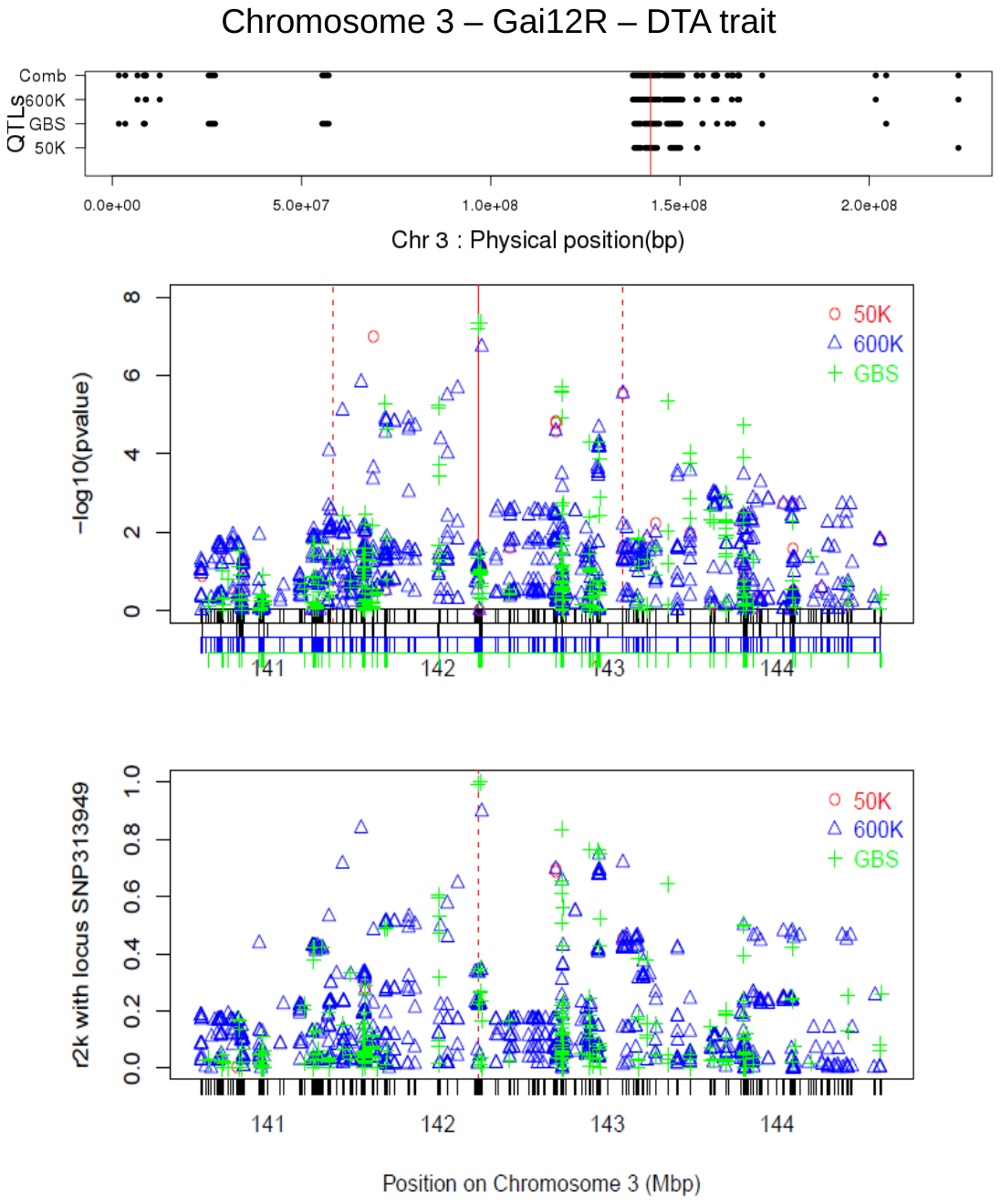

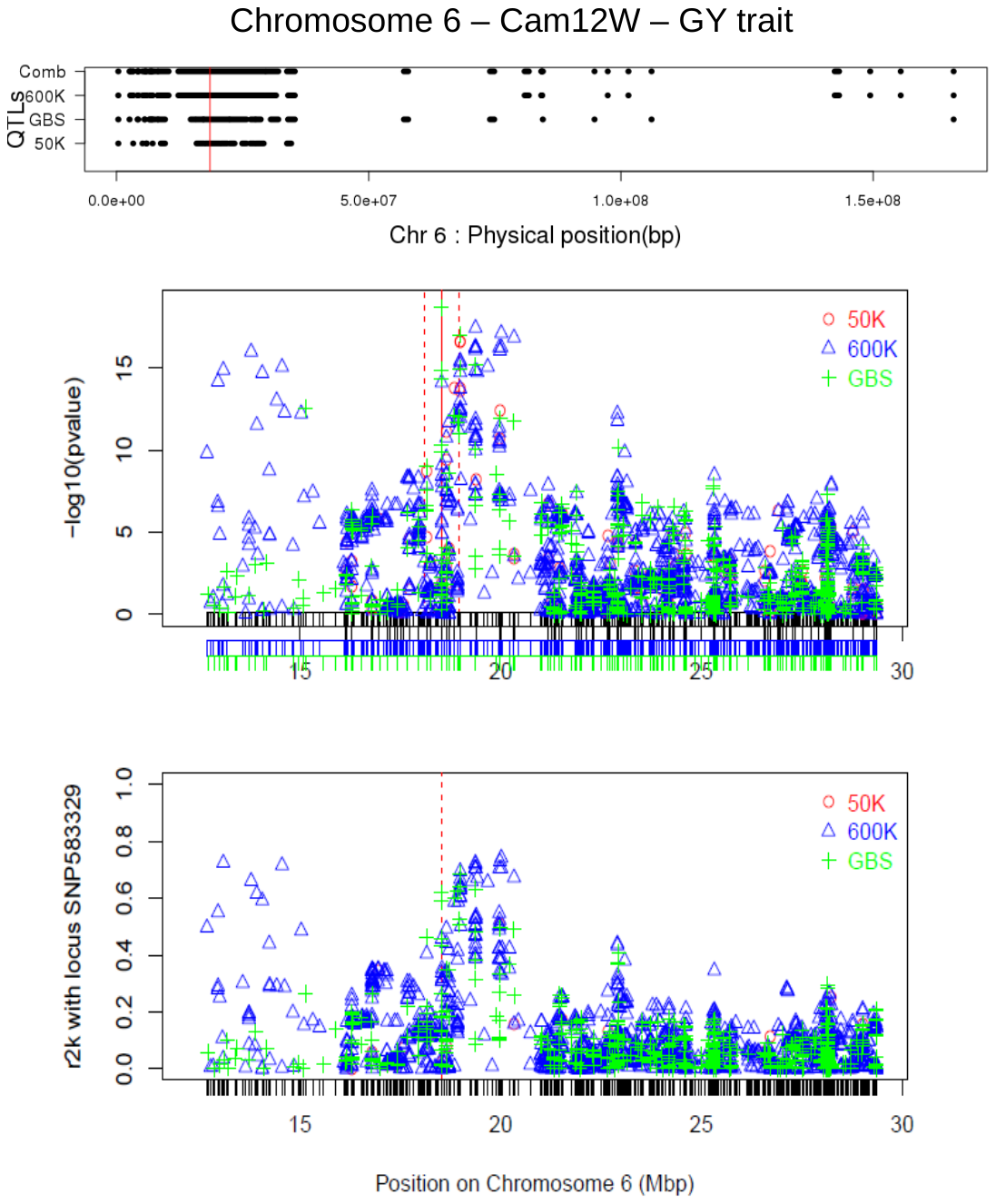

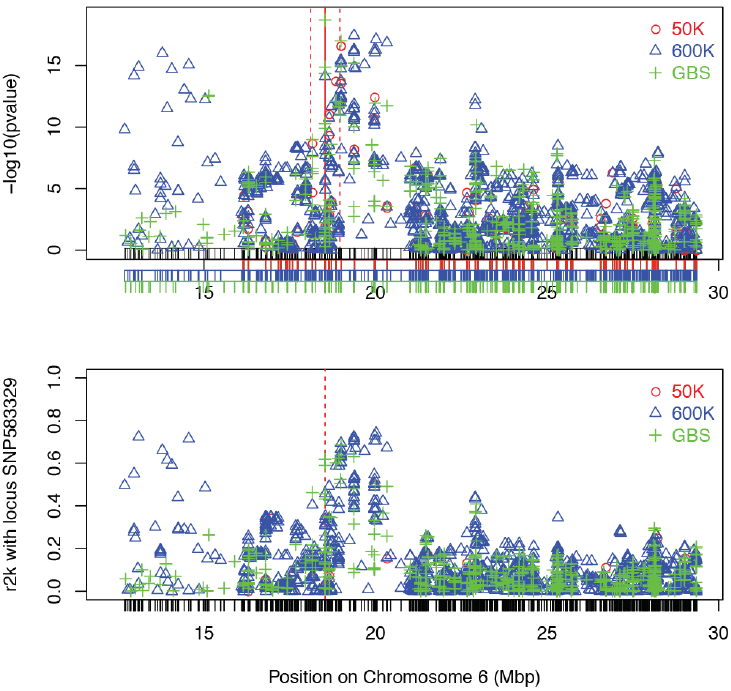

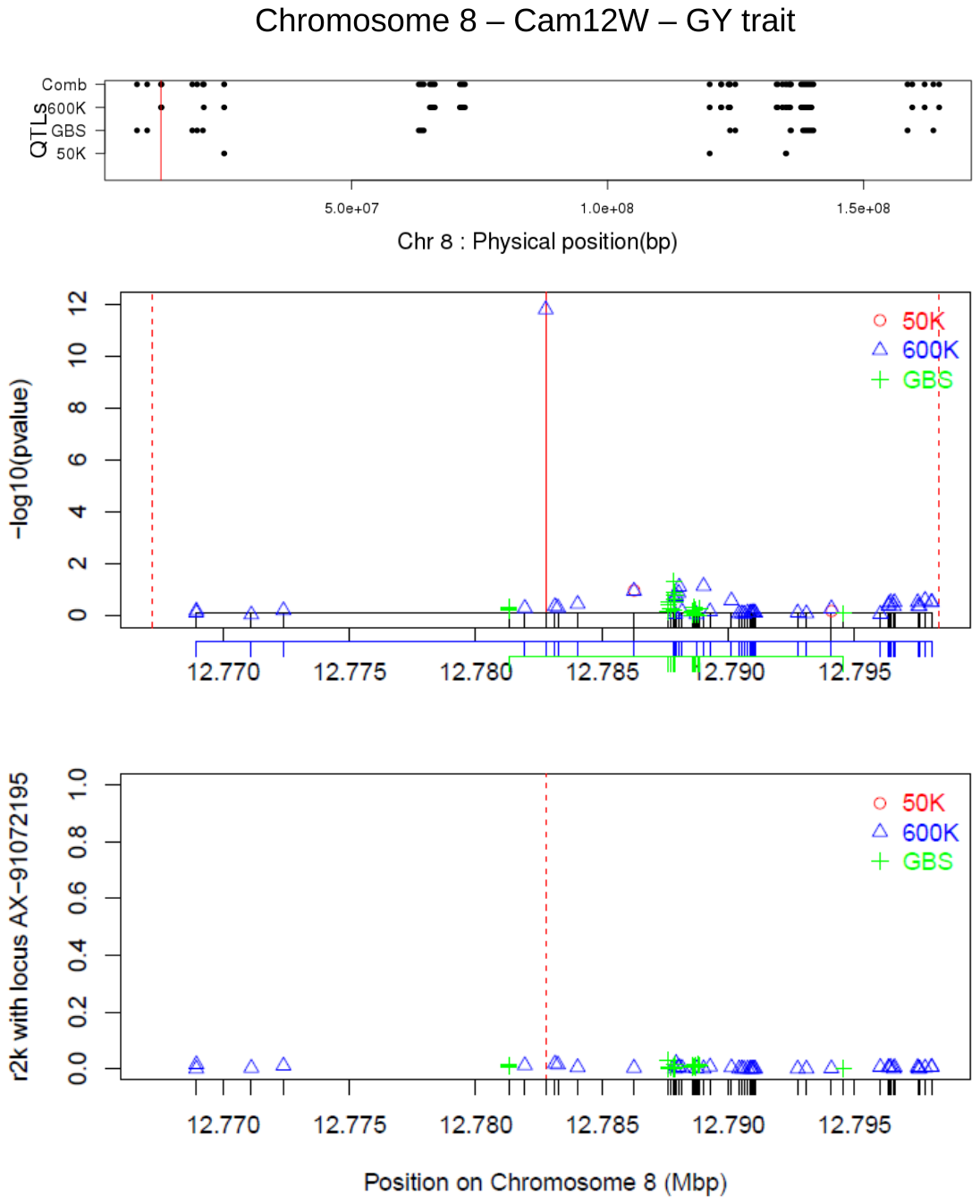

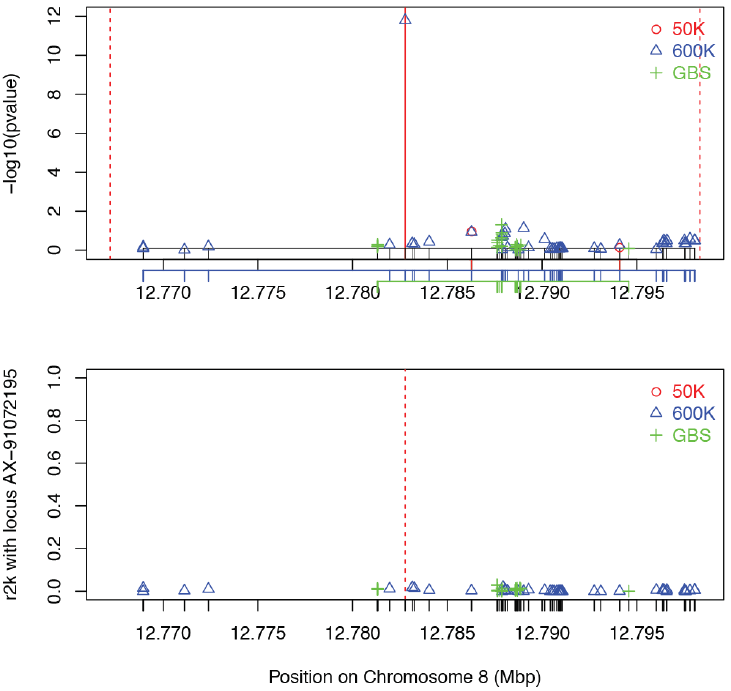

**Figure S10**: Examples of comparison of QTLs detection on Chromosome 3, 6 and 8 for the different traits. The top panel represents the distribution of the QTLs along the chromosome of interest. The vertical red line in this panel localized the chosen QTL. The middle panel is a zoom in the chromosome of the chosen QTL showing the Local distribution of the -*log_10_*(*p-value*). The bottom panel is the same zoom as the middle panel and shows the local linkage disequilibrium corrected by the kinship (*r^2^k*) of all SNPs, within this region, with the strongest associated marker within the chosen QTL. Ticks on different x-axes show the marker density of the three technologies (red for the 50K, blue for the 600K and green for the GBS). The vertical red line spots the position of the SNP with the maximum -*log_10_*(*p-value*) within the QTL.
