## Additional file 12: Fig S11 for "Genotyping-by-sequencing and SNP-arrays are complementary for detecting quantitative trait loci by tagging different haplotypes in association studies"

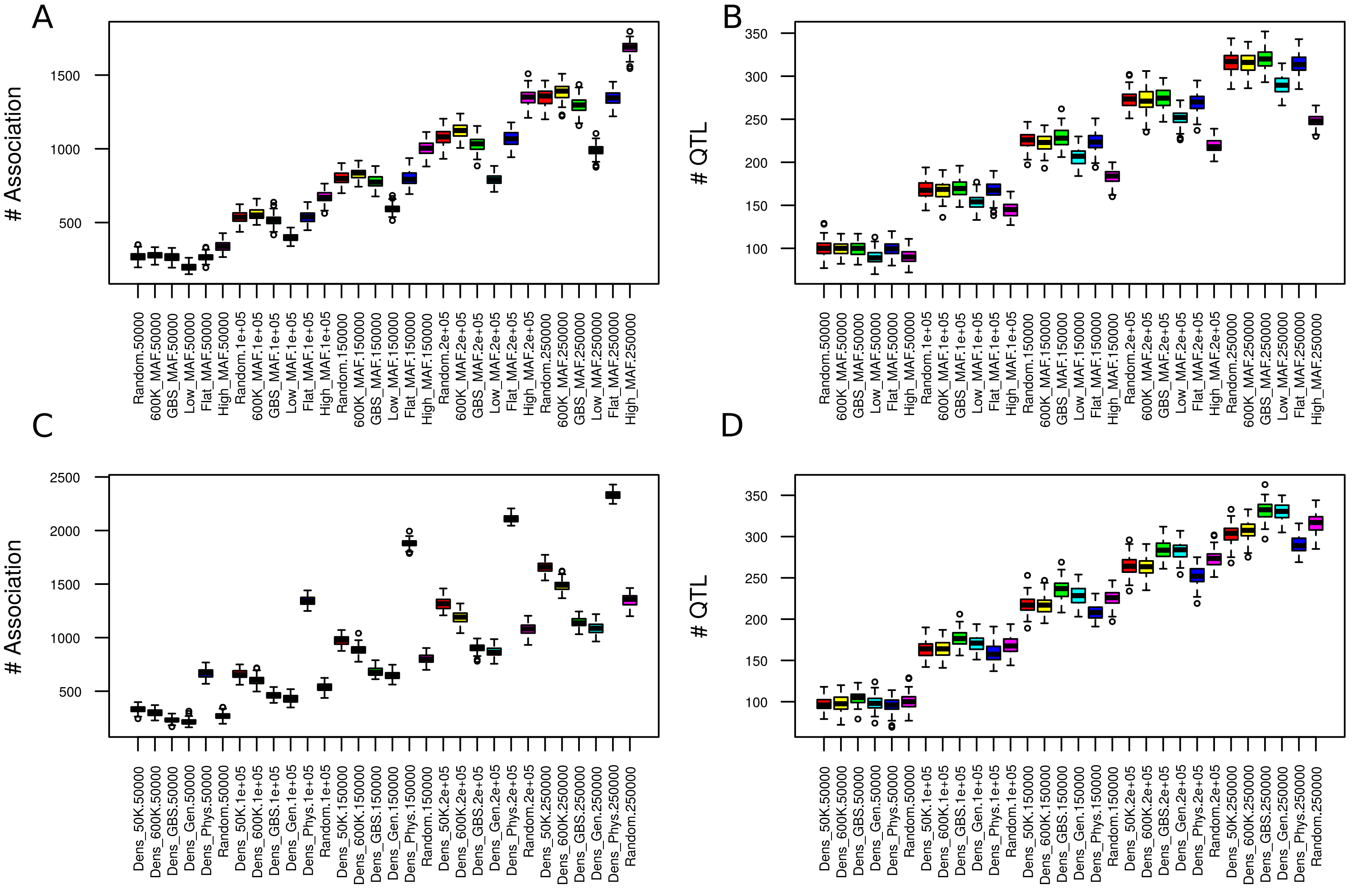

A
B
C
D
### Association
### QTL
Random.50000
600K_MAF.50000
GBS_MAF.50000
Low_MAF.50000
Flat_MAF.50000
High_MAF.50000
Random.1e+05
600K_MAF.1e+05
GBS_MAF.1e+05
Low_MAF.1e+05
Flat_MAF.1e+05
High_MAF.1e+05
Random.150000
600K_MAF.150000
GBS_MAF.150000
Low_MAF.150000
Flat_MAF.150000
High_MAF.150000
Random.2e+05
600K_MAF.2e+05
GBS_MAF.2e+05
Low_MAF.2e+05
Flat_MAF.2e+05
High_MAF.2e+05
Random.250000
600K_MAF.250000
GBS_MAF.250000
Low_MAF.250000
Flat_MAF.250000
High_MAF.250000
Dens_50K.50000
Dens_600K.50000
Dens_GBS.50000
Dens_Gen.50000
Dens_Phys.50000
Random.50000
Dens_50K.1e+05
Dens_600K.1e+05
Dens_GBS.1e+05
Dens_Gen.1e+05
Dens_Phys.1e+05
Random.1e+05
Dens_50K.150000
Dens_600K.150000
Dens_GBS.150000
Dens_Gen.150000
Dens_Phys.150000
Random.150000
Dens_50K.2e+05
Dens_600K.2e+05
Dens_GBS.2e+05
Dens_Gen.2e+05
Dens_Phys.2e+05
Random.2e+05
Dens_50K.250000
Dens_600K.250000
Dens_GBS.250000
Dens_Gen.250000
Dens_Phys.250000
Random.250000

Additional File 12 (.pdf)

Figure S11: Effect of minor allelic frequency distribution, SNP distributions along the genome and SNP densities on the number of associated SNP and QTL detected. Boxplot were drawn on 100 sets of 50 000 to 250 000 markers sampled according to different MAF distributions (A, B) and different SNP distributions along the genome (C, D). A, C: number of SNP associated; B, D: Number of QTL detected. In A and B, 600K_MAF (yellow), GBS_MAF (green), Low_MAF (cyan), Flat_MAF (blue), High_MAF (pink) on x axis indicate boxplots corresponding to MAF distribution similar to 600K, similar to GBS, skewed towards low MAF, flat MAF and skewed toward high MAF, respectively. In C and D, Dens_50K (red), Dens_600K (yellow), Dens_GBS (cyan), Dens_Gen (blue), Dens_Phys (pink) on x axis indicate distribution of SNPs along the genome corresponding to 50K, GBS, 600K, even genetic and physical distances, respectively. For A, B, C and D, modalities indicated as “Random” in x axis correspond to random sample of SNP. Number of markers for each boxplot are indicated after the point.
