## Additional file 13: Fig S12 for "Genotyping-by-sequencing and SNP-arrays are complementary for detecting quantitative trait loci by tagging different haplotypes in association studies"

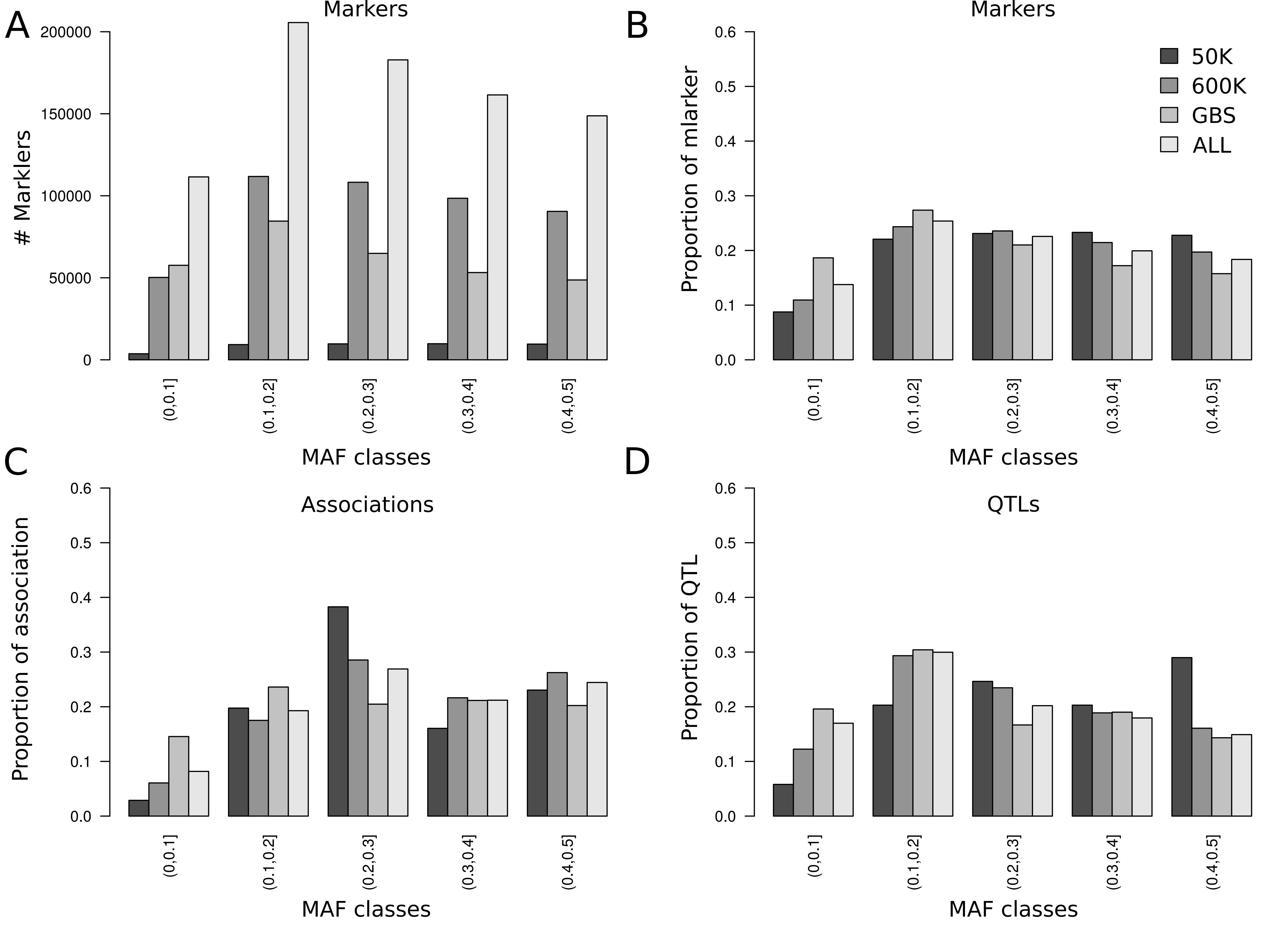

A
Markers
### Marklers
200000
150000
100000
50000
0
(0,0.1]
(0.1,0.2]
(0.2,0.3]
(0.3,0.4]
(0.4,0.5]
MAF classes
B
Markers
Proportion of mlarker
0.6
0.5
0.4
0.3
0.2
0.1
0.0
50K
600K
GBS
ALL
MAF classes
C
Associations
Proportion of association
MAF classes
D
QTLs
Proportion of QTL
MAF classes

Additional file 13 (.pdf)

Figure S12: Distribution of markers, associations and QTLs according to MAF for 50K, 600K GBS, and ALL technologies. A) Number of markers, B) Proportion of markers, C) Proportion of Association, D) Proportion of QTLs
