## Additional file 14: Fig S13 for "Genotyping-by-sequencing and SNP-arrays are complementary for detecting quantitative trait loci by tagging different haplotypes in association studies"

**Figure S13**: Colocalization of QTLs between the traits. Number of QTLs specific and shared by the three traits across all environments. Note that several QTLs from one trait were sometimes included in a single QTL of another trait.
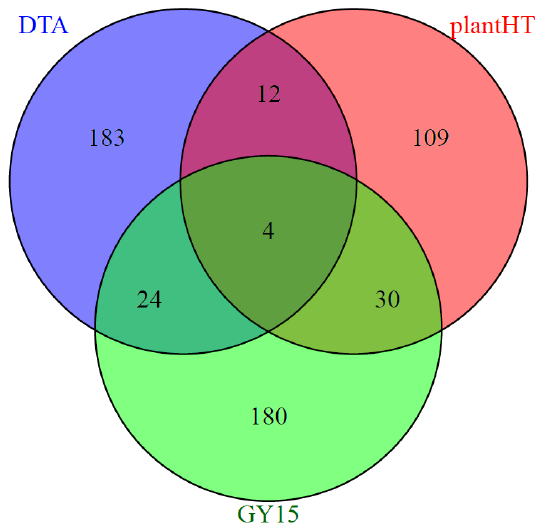
