## Additional file 15: Table S2 for "Genotyping-by-sequencing and SNP-arrays are complementary for detecting quantitative trait loci by tagging different haplotypes in association studies"

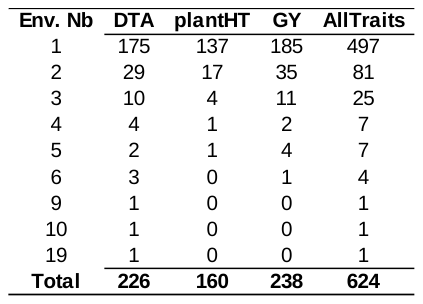

**Table S2:** Stability of QTLs across environments for the three traits (DTA: male flowering time, plantHT: Plant Height, GY: Grain Yield) and all traits. “Env. Nb” indicates the number of environment in which a QTL was detected. Next four columns indicate the number of QTL corresponding to each cartegory
