## Additional file 16: Table S3 for "Genotyping-by-sequencing and SNP-arrays are complementary for detecting quantitative trait loci by tagging different haplotypes in association studies"

**Table S3:** Proportion of low and high recombination regions, recombination rate and percentage of QTLs located in these regions for the three traits.

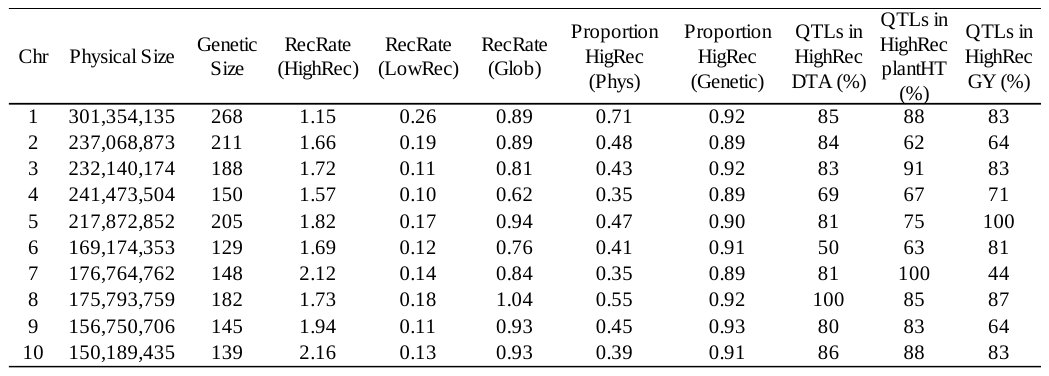

“Chr” indicates the chromosome. Physical and genetic size columns indicated the size of each chromosome in bp and cM, respectively. Average recombination rate (“RecRate”) and proportion of the physical (“Phys”) and genetic (“Genetic) map in high recombination regions (“HighRec”, >0.5 cM / Mbp) for each chromosome are shown. Percentage of QTL in high recombination regions were displayed for three traits (DTA PlantHT, GY)
