## Additional file 18: Table S5 for "Genotyping-by-sequencing and SNP-arrays are complementary for detecting quantitative trait loci by tagging different haplotypes in association studies"

**Table S5:** Narrow sense heritability (h²) and variance components (V_g_, genetic variance; V_e_, residual variance).

The heritability and variance components were estimated for all traits (grain yield, male flowering time and plant height) using the
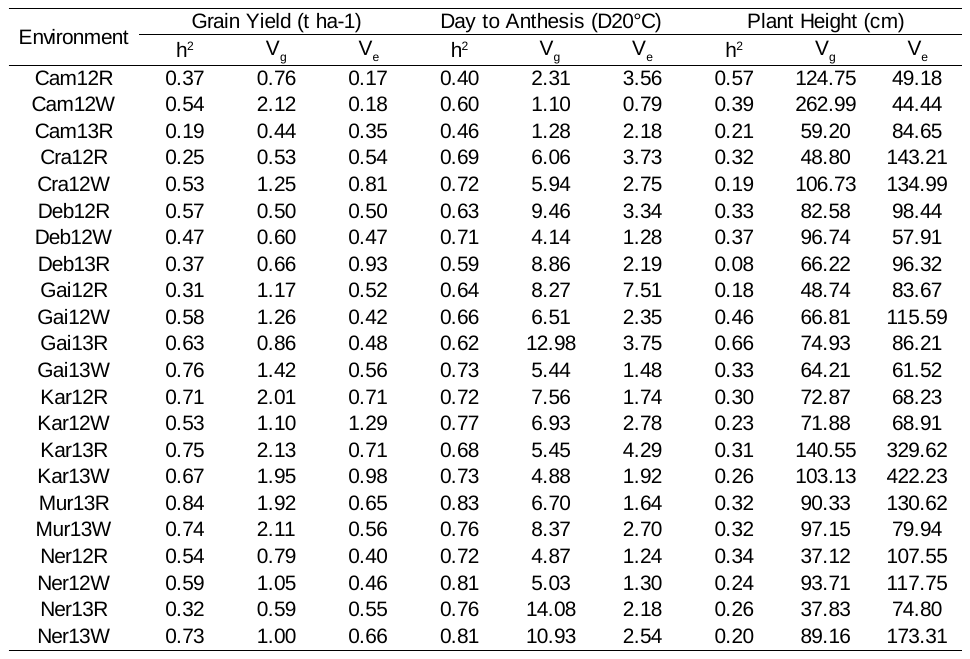
R package Heritability [1].
