## Additional file 19 Suppl Notes S1 for "Genotyping-by-sequencing and SNP-arrays are complementary for detecting quantitative trait loci by tagging different haplotypes in association studies"

**Supplementary Notes S1**

The two main differences between Axiom Affymetrix and Illumina Infinium HD array technologies are the way by which the probes on the array are produced (photolithography *vs* oligo-synthesis) and the way by which the alleles are revealed (ligation extension using labelled degenerated probes *vs* single end extension by polymerase chain reaction using stained nucleotides). The 50K array is composed of 49,574 SNPs (Single Nucleotide Polymorphism markers) previously discovered either by re-sequencing 26 inbred lines representing world-wide maize genetic diversity that are mapped on B73, so called PANZEA SNPs [1] or by aligning expressed sequence tags from Mo17 or other inbred lines as F2 against B73 genome [2]. Since 34,182 PANZEA SNPs were discovered from a broad genetic diversity panel, they are therefore suitable for studying genetic diversity of maize as ascertainment bias is much reduced [2-3]. The 600K Affymetrix array was developed following mapping whole genome sequencing reads of 30 representative temperate maize lines against B73 AGP_v2 [4]. This array includes 616,201 SNPs and 6759 indels, among these are 116,224 markers in coding regions. We selected the SNPs from the *PolyHighResolution* category (522,766 SNPs of high quality) classified by *SNP Polisher* provided by Axiom Affymetrix.

The Discovery pipeline uses the cumulative sequence data from all available samples run to date in a species to discover SNPs. We called 916,444 SNP using the Discovery Build in maize (AllZeaGBSv2.7) that included 97.5 million tags from the sequencing of 31,978 samples [5]. The Production pipeline uses a “production-ready TagsOnPhysicalMap” generated by the Discovery Pipeline to produce genotypes directly from input FASTQ files [5]. The “TagsOnPhysicalMap”, where the discovered SNPs are stored, contains all of the useful, unique, sequence tags, the genomic positions of the subset of the tags with unique best alignment positions, and the alleles that each useful tag represents for each discovered SNP. The *TASSEL-GBS* favors increasing the number of markers over a low sequencing depth (0.5 to 3X), thus the pipeline was designed with low coverage data. The subsequent missing data and under-calling of heterozygotes can be reduced by redundant coverage of haplotypes at high marker density. Related inbred lines can be locally Identical-By-Descent (IBD) because they share a common ancestor. Therefore, they can share locally a common haplotype.
