## Additional file 21 Suppl Notes S2 for "Genotyping-by-sequencing and SNP-arrays are complementary for detecting quantitative trait loci by tagging different haplotypes in association studies"

**Supplementary Notes S2**

Statistical models for GWAS

Model 1: Simple model;

*Y = Xβ + E*

Model 2:Q model;

*Y = Qv + Xβ + E*

Model 3: K model;

*Y = Xβ + Zu + E*

Model 4: Q/K model;

*Y = Qv + Xβ + Zu + E*

where *Y* is the vector of phenotypes; *X* is the vector of *N* genotypes at the tested marker; *β* includes the intercept and the additive marker effect; *Q* is the matrix containing the ancestral genomic fractions from ADMIXTURE; *v* is a vector of fixed population effect; *u ~ N(0, K.σ^2^_gl_)* is the vector of random polygenic background effects, *K* represents the kinship and *σ^2^_gl_* the residual polygenic variance; *E ~ N(0, I.σ^2^_el_)* is the vector of remaining residual effects with variance *σ^2^_el_*; *I* is the identity matrix of size *N; U* and *E* are assumed to be independent. K model and Q/K model are mixed-models where relatedness among individuals is taken into account by considering that the random polygenic effects are not independent, with a covariance matrix determined by *K*. We evaluated two different estimates of *K* in the mixed-models: *K_Freq* and *K_Chr* as described in the main text. This last estimator excludes markers in high LD with the tested SNP in the kinship and increases power [1-2].

The above four statistical models (M1-M4) were evaluated using ASReml to determine the model that controls best the confounding factors (i.e. population structure and relatedness or both) in GWAS. We compared the p-values obtained with different Q+K models. M3 models including only relatedness were sufficient to control false positive inflation. The comparison between mixed models in GWAS using different estimates of kinship (*K_IBS* and *K_Freq)* yielded closed results (data not shown). The kinship estimate that gives a higher weight to markers with low gene diversity (*K_Freq*) was chosen for the following analyses. We observed a gain of power in GWAS using *K_Chr* (kinship estimated by excluding the chromosome of the tested SNP) [2] compared to the mixed model using *K_Freq* (data not shown). Rincent *et al.* [2] approach appears to be a good compromise between control of false positives and power.

Different consequences in GWAS of using different kinships estimated from genetic data with different properties (e.g. allelic frequency profil) can be observed. We showed that the correlations between the *-log_10_(p-value) = 5* of the GWAS using a kinship estimated from the PANZEA 50K chip and a kinship estimated from the GBS (for different situations and traits) were high and most of the association peaks were found in both analyses (data not shown). To evaluate the consequences in GWAS of using different technologies, we used the mixed model (M3) with a kinship estimated from PANZEA markers of the 50K as we showed that the kinships estimated from the different technologies were highly correlated in our study. The false positives for the three technologies were well controlled (QQplot not shown).
